# Coordinated cellular neighborhoods orchestrate antitumoral immunity at the colorectal cancer invasive front

**DOI:** 10.1101/743989

**Authors:** Christian M. Schürch, Salil S. Bhate, Graham L. Barlow, Darci J. Phillips, Luca Noti, Inti Zlobec, Pauline Chu, Sarah Black, Janos Demeter, David R. McIlwain, Nikolay Samusik, Yury Goltsev, Garry P. Nolan

**Author notes:** These authors contributed equally (Christian M. Schürch, Salil S. Bhate, Graham L. Barlow, Darci J. Phillips).

## Abstract

Antitumoral immunity requires organized, spatially nuanced interactions between components of the immune tumor microenvironment (iTME). Understanding this coordinated behavior in effective versus ineffective tumor control will advance immunotherapies. We optimized CO-Detection by indEXing (CODEX) for para ffin-em bedded tissue microarrays, enabling profiling of 140 tissue regions from 35 advanced-stage colorectal cancer (CRC) patients with 56 protein markers simultaneously. We identified nine conserved, distinct cellular neighborhoods (CNs)–a collection of components characteristic of the CRC iTME. Enrichment of PD-1^+^CD4^+^ T cells only within a granulocyte CN positively correlated with survival in a high-risk patient subset. Coupling of tumor and immune CNs, fragmentation of T cell and macrophage CNs, and disruption of inter-CN communication was associated with inferior outcomes. This study provides a framework for interrogating complex biological processes, such as antitumoral immunity, demonstrating an example of how tumors can disrupt imm une functionality through interference in the concerted action of cells and spatial domains.

## INTRODUCTION

Recent efforts have highlighted how interactions between distinct components of the immune tumor microenvironment (iTME) play crucial roles in dictating cancer development, progression, rejection, and therapeutic response (Junttila and de Sauvage, 2013; Schreiber et al., 2011; Williams et al., 2016). Spatial complexity of tumor architecture is a fundamental feature of the iTME, and considerable local and regional variability are observed within individual tumors and in the surrounding tissue (Gillies et al., 2012; Junttila and de Sauvage, 2013). Pathologists have long recognized that conserved patterns of spatial organization and cellular features of tissue components are prognostic in many cancers, and recent studies have suggested that cellular spatial context and tissue sub-structure within the iTME play important roles in therapeutic response and patient outcome (Binnewies et al., 2018). How, then, can delineating the role of specific cell-cell interactions as well as higher order relationships between cancer, stromal, and immune components facilitate a richer understanding of cancer progression and lead to new therapeutic avenues?

Here we explore how the deep profiling of the iTME architecture in controlled clinical cohorts will facilitate the development of prognostic spatial signatures and lead to mechanistic understanding of biological programs underlying their formation. Multi-parameter tissue imaging technologies are primed to provide the raw data to achieve these objectives, as these technologies allow characterization of single cell phenotypes as well as mathematical access to every cell’s position and posture within intact tissues. Various optical multiplexing methods based on the detection of target-bound antibodies have recently been developed. The Vectra^®^ system, for instance, uses secondary detection by chromogenic molecules (Huang et al., 2013), with a current high limit of 10 simultaneous markers measured. Several cyclic immunofluorescence protocols were introduced that are based on fluorophore inactivation or antibody stripping followed by re-staining. These methods can visualize dozens of markers in single cells and include MELC (Schubert et al., 2006), MxIF (Gerdes et al., 2013), t-CyCIF (Lin et al., 2018) and 4i (Gut et al., 2018). A different approach uses DNA-barcoded antibodies that are visualized by cyclic addition and removal of fluorescently labeled DNA probes, as exemplified by exchange-PAINT (Agasti et al., 2017), DNA exchange imaging (DEI) (Wang et al., 2017b), and immuno-SABER (Saka et al., 2018). Other methods use mass spectrometry-based detection of isotope-labeled antibodies in tissue by raster laser ablation (imaging mass cytometry [IMC]) (Giesen et al., 2014) or ion beams (multiplexed ion beam imaging [MIBI]) (Angelo et al., 2014; Keren et al., 2018), detect and/or amplify endogenous nucleic acids *in situ* (Moffitt et al., 2018; Wang et al., 2018), or use vibrational signatures of chemical bonds to visualize molecules directly (Wei et al., 2017).

Our recently developed a method called CO-Detection by indEXing (CODEX) uses DNA-barcoded antibodies that are visualized by cyclic addition and removal of fluorescently labeled DNA probes (Goltsev et al., 2018) (Kennedy-Darling et al., in preparation). The commercial CODEX platform allows simultaneous visualization of 50 or more antigens in a single tissue section, thereby enabling a systems-level approach to the analysis of tissue architecture. In Goltsev et al., we introduced the concept of an “i-niche”, defined as a ring of immediate neighbors directly contacting an “index cell”. We demonstrated that the i-niche composition was associated with the index cell’s marker expression and tissue localization, as well as disease progression in a murine lupus model. Further demonstrating the clinical utility of high-parameter imaging methods for quantifying contacts between cells and their neighbors, Keren et al. reported that the spatial compartmentalization of tumor/immune cell-cell contacts is linked to patient outcome in triple negative breast cancer (Keren et al., 2018).

We reasoned that effective antitumoral immune responses require coordination of biological processes at multiple levels of abstraction, not just at the level of single cell behavior or pairwise cell-cell contacts (Schapiro et al., 2017; Zhu et al., 2018). This led us to devise descriptions of tissue at a more dialectic level, based on a philosophy that tissues self-organize according to spatial domain-based interactions that assemble via motivated communications initiated as individual cells, but collectively accomplishing functions only as an ensemble. In cancer, tumor cells could disrupt the emergent order created by these communication structures between spatial domains, thereby gaining another method by which to evade active immune assault. This premise was explored adopting the simplest possible approximation to such spatial domains, by defining cellular neighborhoods (CNs) as regions of the tissue within which each cell has a similar surrounding (its surrounding, in this case, defined by the relative proportions of various cell types within a fixed radius), and assessing how the CNs were structured by inter-cellular and inter-CN communication. As such, there is dynamic, coordinated behavior involving not only different combinations of cell types, but also involving different combinations of CNs, as well as an interplay between the biological processes operating at both levels of abstraction (cell types and CNs). In such a system, one would interrogate how the emergent behaviors of these cell types and CNs could model successful antitumoral immune responses. We focused on three complementary perspectives with which to understand these behaviors in the iTME: 1) how combinations of CNs and combinations of cell types are organized together to form the tissue; 2) variation in the functional states of CNs; and 3) how CNs communicate with each other with respect to key functional cellular subsets. We provide herein mathematical descriptions of such interactions and explore how they explain iTME structure correlations to patient outcomes in a cancer of extreme clinical importance.

A tumor with tractable, repeated, structural differences that are correlated to outcome and amenable to such an approach, is colorectal cancer (CRC) – one of the leading causes of cancer deaths in the Western world (Siegel et al., 2018). In a subset of CRC patients, organized tertiary lymphoid structures (TLS) form at the tumor invasive margin. Patients who exhibit this *de novo* formation of B cell follicles and surrounding T cell areas – the so-called “Crohn’s-like reaction” (CLR) – have better overall survival than patients with only diffuse inflammatory infiltration (DII) (Cyster, 2003; Di Caro et al., 2014; Graham and Appelman, 1990). Although the survival advantage of TLS formation is well established for many solid cancers (Dieu-Nosjean et al., 2016; Sautes-Fridman et al., 2016), the associated tissue programs contributing to spatial organization and cancer control remain elusive. The question becomes, then, are there immune structures, perceived as assemblies of cells with coordinated goals, that are required for a positive outcome in CLR, while in DII the tumor’s interference in the creation and concert of such structures mitigates appropriate immune action?

We performed multiplexed spatial imaging using an optimized CODEX methodology for formalin-fixed, paraffin-embedded (FFPE) tissue specimens to study the iTME in well-documented, archival CRC tissue samples. A pipeline for high-precision tissue microarrays (TMAs)was implemented to increase the throughput and enable imaging of numerous tumor samples simultaneously. Within a retrospective cohort of 715 patients, we identified 35 advanced-stage patients with distinct patterns of immune response to cancer (17 patients with CLR and 18 patients with DII) using stringent inclusion and exclusion criteria (**Figure 1A**). We leveraged FFPE-CODEX for the spatial dissection of 140 iTME tissue regions from the tumor-invasive front in these 35 patients, imaging 56 proteins simultaneously, including tumor, immune, immunoregulatory, and stromal antigens. We developed an analytical framework to describe the iTME architecture and identify how key alterations in it are associated with clinical outcomes.

**Figure 1.**
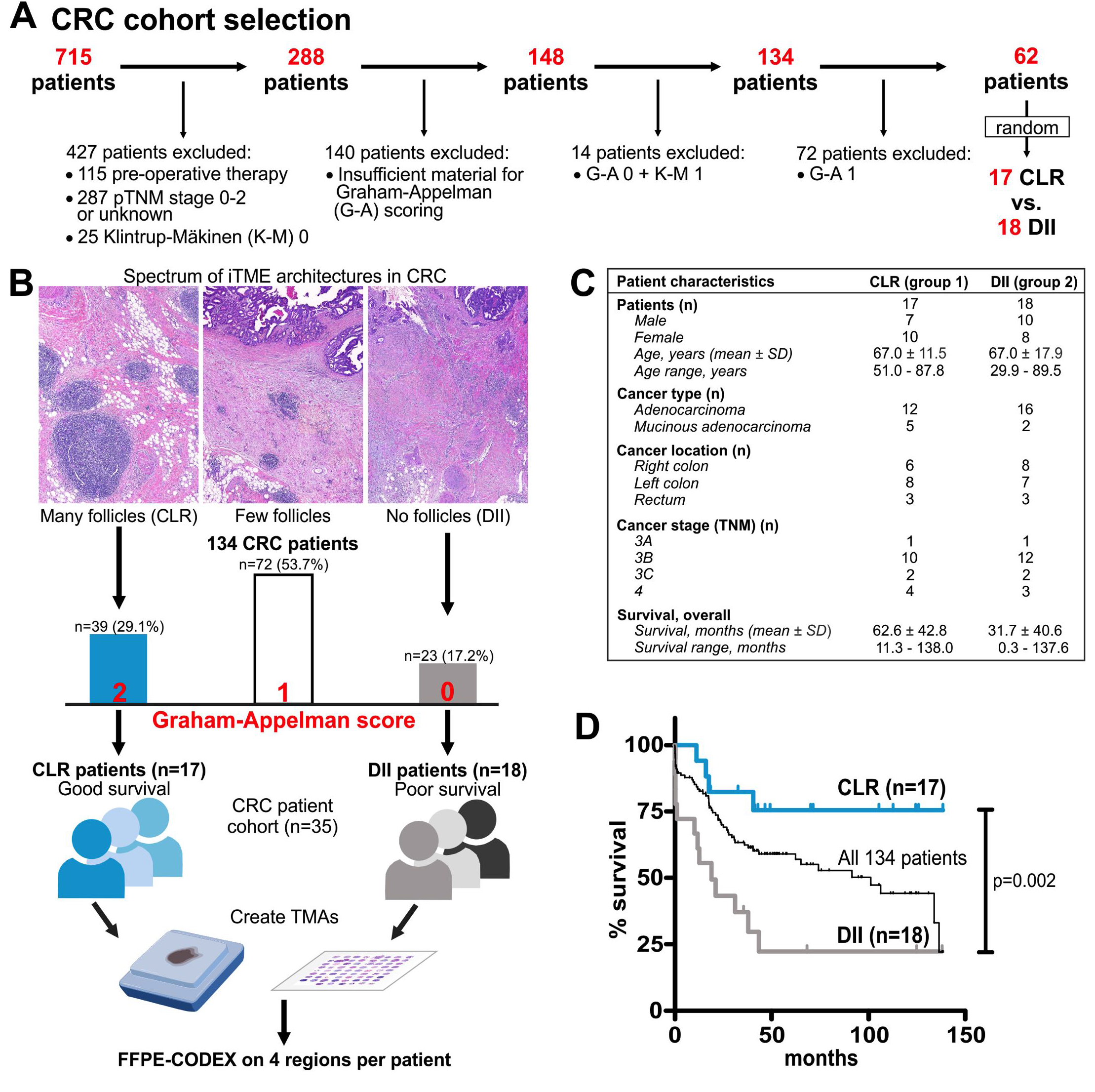
Colorectal cancer (CRC) study cohort. **(A)** From a database of 715 CRC patients, patients with pre-operative therapy, pathological tumor, nodes, metastasis (pTNM) score 0-2 or unknown, absent immune infiltration (Klintrup-Mäkinen [KM] score 0), insufficient material for Graham-Appelman (G-A) scoring, a combination of low immune infiltration (K-M 1) and absent follicles (G-A 0), or few follicles (G-A 1) were excluded (**see Methods**). From the remaining 62 patients, a matched study cohort of n=17 CLR and n=18 DII patients was randomly selected. **(B)** Spectrum of iTME architectures in 134 advanced-stage CRC patients. Representative hematoxylin and eosin (H&E)-stained sections from iTME areas with many follicles (G-A 2, characteristic of CLR), few follicles (G-A 1) and no follicles (G-A 0, characteristic of DII) are shown. **(C)** Characteristics of patients in the CRC study cohort. **(D)** Kaplan-Meier survival curve of the CRC study cohort. CLR and DII patients are compared (p determined with a log-rank test). All 134 patients from (B) are shown for comparison.

We found that the CRC iTME can be computationally decomposed into nine distinct CNs, each a characteristic microenvironment, that, with the exception of the CN corresponding to the TLS (i.e., the lymphoid follicle), were conserved in their abundance and local cell type composition across both patient groups. The functional state of a granulocyte-enriched CN, as assessed by the frequency of PD-1^+^CD4^+^ T cells within it, was positively correlated with survival in DII patients, whereas the overall frequency of these cells was not. The worse outcomes for DII patients were associated with crucial, subtle differences in the CNs’ organization, functional states, and communication network. We identified two “tissue modules” (coordinated tissue, CN and cell type components) in each patient group. In CLR patients, these corresponded to separate immune and tumor modules. In DII patients, these consisted of a coupled tumor/immune module and a distinct granulocyte module. An explanation for the coupled tumor/immune tissue module in DII patients is that immune CNs collectively compartmentalized in CLR were increasingly interspersed with tumor CNs in DII. This was associated with coupling between and fragmentation of T cell-enriched and macrophage-enriched CNs. Alongside these changes in organization, in DII patients, the T cell-enriched CN had decreased enrichment of proliferating CD8^+^ T cells, and the macrophage-enriched CN had increased enrichment of regulatory T cells. In addition, in DII patients, but not CLR patients, the frequency of regulatory T cells (Tregs) in this macrophage-enriched CN was negatively correlated with the frequency of proliferating CD8^+^ T cells in the T cell-enriched CN (which we infer as immunosuppressive inter-CN “communication”). Finally, while the T cell composition in the tumor boundary CN of CLR patients was correlated with the T cell-enriched CN, in DII patients it was correlated with the macrophage-enriched CN.

These results collectively suggest that in DII patients, unlike CLR patients, the tumor modulates the organization, functional states and intercommunication of the T cell- and macrophage-enriched CNs as well as their ability to communicate with the tumor boundary CN. Our data support a model in which the organization and behavior of T cell and macrophage CNs facilitate effective immune responses in CLR patients and ineffective responses in DII patients—with an apparent goal of the tumor to expend energy ensuring that the immune system does not consolidate into an effective coordinated antitumoral activity module. The approach described here is generalizable towards other multiplexed imaging modalities and enables critical architectural features to be linked to patient outcome and a mechanistic understanding of the iTME, begging the question of whether other tumors disrupt immune action in similar manners.

## RESULTS

### Selection of patient samples by classically determined iTME structures in CRC

Colorectal cancer (CRC) patients exhibit a range of native immune infiltrates and associated histologic features (**Figure 1B**). On one end of the spectrum of CRC iTME phenotypes are the patients who have a Crohn’s-like reaction (CLR), characterized by the presence of numerous TLS; these structures are correlated with better-than-average survival. At the other end, patients with diffuse inflammatory infiltration (DII) and absence of TLS have poor survival outcomes. Between these two extremes are a spectrum of intermediate iTME phenotypes with a range of outcomes.

From a database of 715 CRC patients, patients with early-stage CRC, pre-operative chemotherapy, insufficient material, and absent immune infiltration were excluded (**Figure 1A**). Of the remaining 134 patients, 62 had either CLR or DII patterns, and from these a study cohort of 17 patients with CLR and 18 with DII was randomly selected (**Figure 1B**). The two groups were matched with regards to gender, age, cancer type, location, and stage (**Figure 1C** and **Table S1**). In line with previous reports, overall survival of CLR patients was significantly better than that of DII patients (**Figure 1D**).

We reasoned that, since the immune responses to CRC occupy a wide spectrum that is directly linked to patient outcome (**Figures 1B-D**), the components of the iTME architecture that are conserved between its extremes would enable the development of a model describing its function and dysfunction. We therefore determined what exactly these common components are, and how their organization restricts or facilitates tumor growth. The key question was: Was there simply “less structure” in the architecture of the DII patients, or were there crucial differences in the organization of these components that could explain their distinct antitumoral responses?

### FFPE-optimized CODEX enables highly multiplexed fluorescence microscopy of archival human cancer samples

We previously reported the development of CODEX, a fluorescence microscopy platform that uses DNA-barcoded antibodies and iterative visualization based on nucleotide addition by polymerase single-base extension, and chemical removal, of fluorescent nucleotides for highly multiplexed imaging of fresh-frozen tissues (Goltsev et al., 2018). The CODEX technology was subsequently simplified by implementing an iterative approach involving exchange of fluorescently labeled complementary DNA probes (Kennedy-Darling et al., in involves iterative annealing, imaging, and stripping of fluorescently labeled DNA probes complementary to the DNA barcodes on the tissue-bound antibodies using an automated microfluidics system and a conventional optical fluorescent microscope (**Figure 2A**). Hoechst nuclear stain is recorded in every iteration cycle to serve as a reference for computational image alignment.

**Figure 2.**
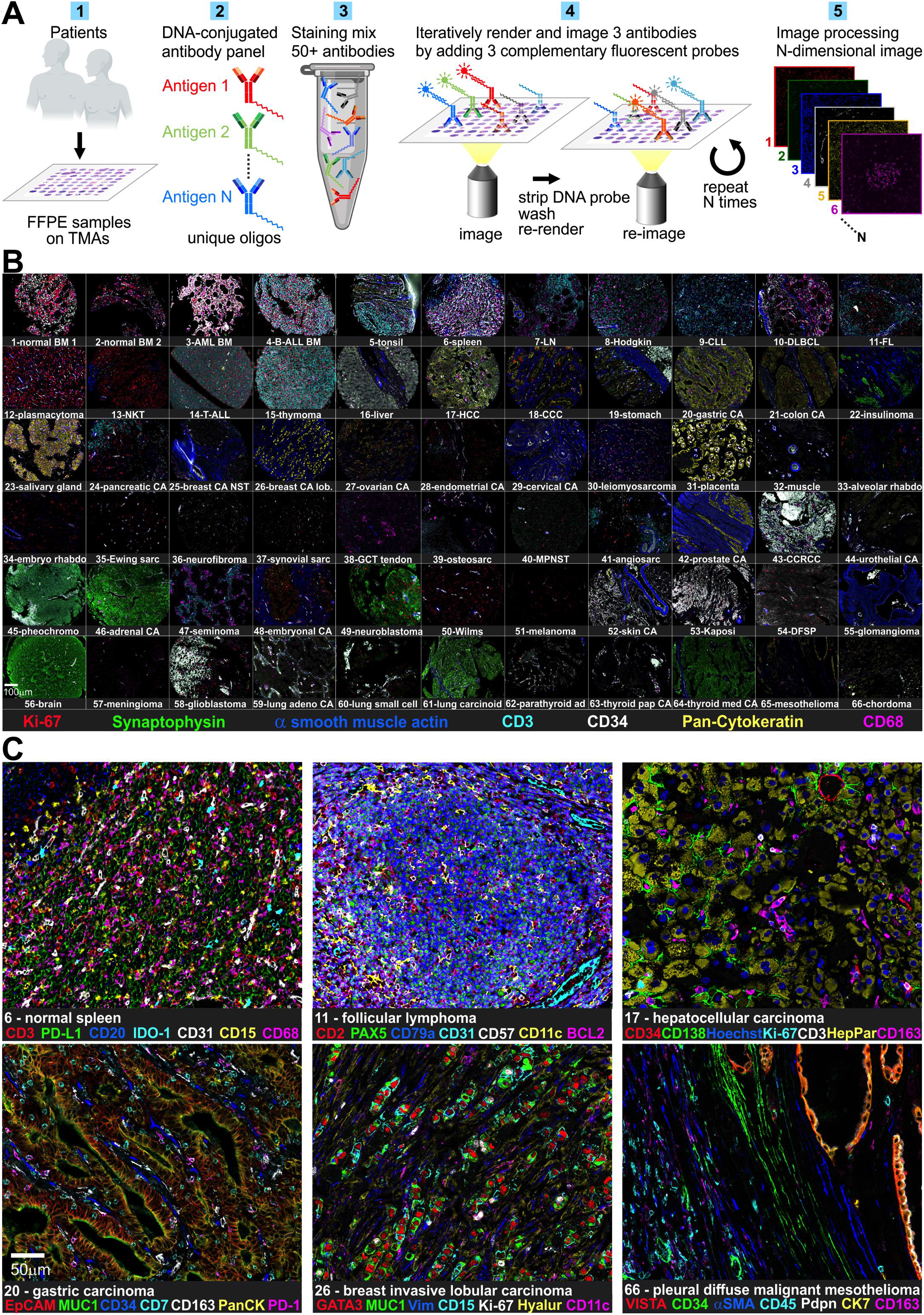
CODEX workflow and antibody validation in multi-tumor TMA. **(A)** CODEX workflow. (1) Patient samples from FFPE blocks are assembled into high-precision TMAs. (2) Each antibody is conjugated to a unique DNA-oligonucleotide. (3) Antibodies are assembled into a master mix, and tissues are stained overnight. (4) Antibodies are iteratively rendered visible in a multi-cycle reaction by adding complementary, fluorescently labeled DNA probes; in the workflow used here, two to three antibodies were imaged in each cycle. After imaging, the fluorescent probes are stripped off, and the process is repeated until all antibodies have been imaged. (5) The resulting images are computationally processed. **(B)** A multi-tumor TMA consisting of 55 different malignant and non-malignant tumors and 11 healthy tissues (**Figure S4 and Table S5**) was imaged using a 56-marker CODEX panel (**Table S4**). A seven-color overview image with Ki-67 (red), synaptophysin (green), α-SMA (blue), CD3 (cyan), CD34 (white), pan-cytokeratin (yellow), and CD68 (magenta) is shown. **(C)** Higher magnification and seven-color overlay images of normal spleen (CD3, PD-L1, CD20, IDO-1, CD31, CD15, CD68), follicular lymphoma (CD2, PAX5, CD79a, CD31, CD57, CD11c, BCL2), hepatocellular carcinoma (CD34, CD138, Hoechst, Ki-67, CD3, Hep-Par-1, CD163), gastric carcinoma (EpCAM, MUC-1, CD34, CD7, CD163, pan-cytokeratin, PD-1), breast invasive lobular carcinoma (GATA-3, MUC-1, vimentin, CD15, Ki-67, hyaluronan, CD11c), and pleural diffuse malignant mesothelioma (VISTA, CD34, α-SMA, CD45, podoplanin, cytokeratin 7, CD163). Abbreviations: BM, bone marrow; AML, acute myeloid leukemia; B-ALL, B cell acute lymphoblastic leukemia; LN, lymph node; CLL, chronic lymphocytic leukemia; DLBCL, diffuse large B cell lymphoma; FL, follicular lymphoma; NKT, natural killer/T cell lymphoma; T-ALL, T cell acute lymphoblastic leukemia; HCC, hepatocellular carcinoma; CCC, cholangiocarcinoma; CA, carcinoma; NST, no special type; rhabdo, rhabdomyosarcoma; sarc, sarcoma; GCT, giant cell tumor; MPNST, malignant peripheral nerve sheath tumor; CCRCC, clear cell renal cell carcinoma; DFSP, dermatofibrosarcoma protuberans. See also **Figures S1, S2, S3, S4, S5 and S6**

We optimized CODEX for use in FFPE tissue to study archival human tissue samples. FFPE-CODEX staining panels consisting of 50 or more markers were created by conjugating antibodies previously validated for immunohistochemistry (IHC) to unique short DNA oligonucleotides (**Tables S2, S3 and S4**). CODEX antibodies were screened and validated for correct staining patterns in parallel with standard manual IHC using identical staining protocols (**Figure S1**). Staining results were verified by comparison to online databases and published literature. As proof-of-principle for the FFPE-CODEX approach, final validation and titration of the complete antibody panels were performed in multi-cycle CODEX imaging experiments using FFPE human tonsil sections (**Figures S2 and S3, Table S4**). At the end of each CODEX run, the tissue sections were stained with hematoxylin and eosin (H&E) for morphological correlation.

We next performed an in-depth validation of our CODEX antibody panel by staining and analyzing a multi-tumor TMA (**Figures 2B and S4, Table S5**). Critically, since we could test multiple reagents across many tissue types in this experiment, use of co-staining and marker exclusion patterns (for instance, either CD4 or CD8 expression but not both on CD3^+^ T cells) greatly facilitated reagent validation. Expected antigen distribution patterns were observed in all tissues analyzed, exemplified by normal spleen, follicular lymphoma, hepatocellular carcinoma, gastric carcinoma, breast invasive lobular carcinoma, and pleural diffuse malignant mesothelioma (**Figure 2C**). For example, healthy spleen tissue showed a normal distribution of red and white pulp (CD3, CD20, CD68), presence of granulocytes (CD15), localization of indoleamine 2,3-dioxygenase 1 (IDO-1) in red pulp macrophages, and prominent PD-L1 expression in splenic sinusoids (CD31). Notably, we detected an unexpected strong and ubiquitous expression of the T cell checkpoint marker V domain Ig suppressor of T cell activation (VISTA) in mesothelioma, which is confirmed as a feature of mesothelioma in a recently described integrative genomic characterization study (Hmeljak et al., 2018).

The CODEX method is an iterative imaging approach with numerous washing and stripping steps that potentially degrade the tissue over the time of the multicycle reaction. To exclude this possibility, we performed a re-cycling experiment on FFPE tonsil using a panel of nine antibodies over 33 cycles. This experiment confirmed that the marker intensity remained constant and the tissue morphology was intact throughout the process (**Figures S5 and S6**).

Collectively, these data demonstrate that FFPE-optimized CODEX is suitable for highly multiplexed single-cell marker visualization, quantification and biomarker discovery in clinically relevant tissues.

### FFPE-CODEX enables *in situ* identification and quantification of major immune cell types in the iTME of the CRC invasive front

At the tumor invasive front of CRC, the iTME is usually seen as leukocyte-dense regions alternating with regions of sparse immune infiltration. In DII patients, TLS (follicles) are absent in the immune infiltrate but are abundant in tumors from CLR patients. We created two high-precision TMAs by selecting four representative leukocyte-dense iTME regions from the tumor invasive front for each patient, resulting in a total of 140 regions (**Figure S7**). For CLR patients, two regions per patient contained organized TLS and two regions contained diffusely inflamed tissue, whereas in the DII patient group all four regions selected contained diffusely inflamed tissue. FFPE-CODEX with 56 markers and two nuclear stains was performed on the TMAs, and staining results were validated for each marker (**Figures 3A-B and S8**).

**Figure 3.**
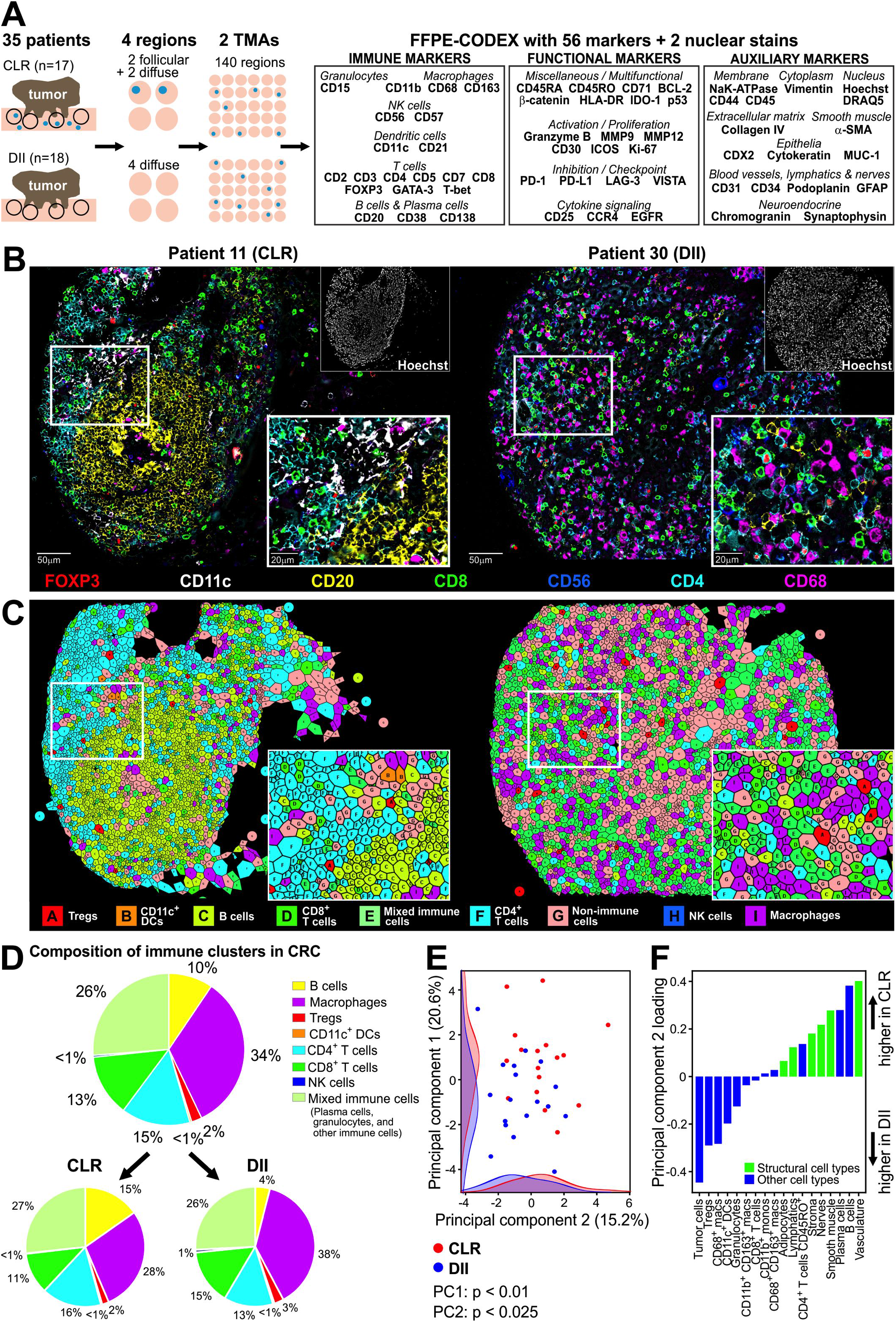
CODEX imaging reveals detailed spatial composition of immune infiltrates in CRC. **(A)** Schematic of CRC TMA assembly. For each of the 35 patients (CLR. n=17; DII, n=18). four cores were drilled at the tumor invasive front to create two 70-core TMAs were then subjected to CODEX with a 56-marker panel (**Table S4**). Blue dots represent follicles. **(B)** Representative TMA cores for CLR and DII patients are depicted as seven-color overlay images with FOXP3 (red), CD11c (white), CD20 (yellow). CD8 (green), CD56 (blue), CD4 (cyan) and CD68 (magenta). **(C-F)** Single-cell protein marker expression data from all spots of both TMAs (n=258,385 total cells) were subjected to X-shift clustering. Clusters were visually verified and manually merged based on morphology and marker expression profiles into 28 distinct clusters **(Figures S10 and S11). (C)** The 28 clusters were merged into eight different immune clusters and one cluster containing all non-immune cells (tumor cells, smooth muscle cells, stromal cells, vasculature, lymphatics, nenes and other cell types). These nine clusters are shown as Voronoi diagrams for both cores. The most important markers for cluster identification are listed. A: Tregs (CD3^+^CD4^+^FOXP3^+^). B: DCs (CD11c^+^). C: B cells (CD20^+^). D: CD8^+^T cells (CD3^+^CD8^+^). E: Other immune cells (H&E, CD45^+^, and other markers). F: CD4^+^T cells (CD3^+^CD4^+^). G: Non-immune cells (H&E, pan-cytokeratin^+^, CD31^+^, α-SMA^+^ and other markers). H: NK cells (CD56^+^GATA3^+^; cluster not appearing in images). I: Macrophages (CD68^+^. CD163^+^). **(D)** The eight immune clusters (n=132,437 cells) and their frequencies in (top) all CRC patients (bottom) and separated into CLR (n=57,894 cells) and DII patients (n=74,543 cells). **(E)** PCA correlating combinations of cell type abundances in CLR vs. DII patients. **(F)** Cell type loading in principal component 2. See also Figures S7, S8, S9, S10, S11, S12, S13, S14, S15 and S16A-B

Fifty-five of the 56 markers in the panel (all except collagen IV) are cell-surface or intracellular proteins. They are used as phenotypic and functional markers either alone or in combination to identify specific cell subpopulations and functional states. We approached the problem of cell type identification by two complementary approaches: one using a supervised gating strategy, and one using unsupervised clustering. After single-cell segmentation, marker quantification and spatial fluorescence compensation (Goltsev et al., 2018), manual cleanup gating for each individual TMA core was performed in CellEngine (www.cellengine.com) to positively identify cells (events double-positive for Hoechst and DRAQ5 nuclear stains), and to remove out-of-focus events in the Z plane. For unsupervised identification of cell types, presumed cell events were then exported and subjected to X-shift clustering using VorteX (Samusik et al., 2016). To assess the reliability of this unsupervised approach, cell events were additionally gated manually in CellEngine (**Figure S9**). VorteX clustering, followed by supervised merging of clusters based on marker expression profiles, tissue localization and morphology resulted in 28 unique clusters of definable cell subsets. These included 18 immune cell clusters, 6 stromal and vasculature clusters, 2 mixed clusters, 1 tumor cell cluster, and 1 undefined cluster (**Figures S10, S11 and S12**). Supervised manual gating and unsupervised X-shift clustering led to comparable results for absolute numbers and frequencies of defined major immune cell types (**Figures S13 and S14**). For downstream analyses, we chose cell types identified using unsupervised clustering.

Graphical representation of imaged samples as Voronoi diagrams colored by cell type can be used to visualize the spatial distribution of cell types in tissue and to verify the clustering results. Since the visual interpretation of 28 different CRC clusters on Voronoi diagrams was potentially subjective (**Figure S12**), we reduced the complexity manually. Six initial “macrophage” clusters (CD11b^+^ monocytes, CD11b^+^CD68^+^ macrophages, CD68^+^ macrophages, CD68^+^ granzyme B^+^ macrophages, CD68^+^CD163^+^ macrophages and CD163^+^ macrophages) were merged into a single cluster, and all tumor and stromal cell clusters were merged into a single cluster called “non-immune cells” (**Figures 3C and S15**). We quantified each immune cell type as a fraction of total immune cells on an overall, per group and per patient basis. The frequency of immune subsets across all patients revealed cell types with high (i.e., macrophages, 34%), medium (i.e., CD4^+^ T cells, 15%; CD8^+^ T cells, 13%; B cells, 10%) and low frequencies (i.e., Tregs, 2%; natural killer [NK] cells, <1%; CD11c^+^ dendritic cells [DCs], <1%) (**Figures 3D and S15**).

Differences in the composition of these immune cell clusters were observed between CLR and DII patients. Most notably, CLR patients had higher frequencies of B cells, whereas DII patients had higher frequencies of macrophages. Interestingly, the frequencies of CD8^+^ T cell and Treg clusters were not significantly different between CLR and DII patients (**Figures 3D and S15**). This is in contrast to prior studies in CRC and other cancers, including lung, breast, ovarian and melanoma, in which relative abundances of these two cell types were correlated with outcome (Fridman et al., 2012; Galon et al., 2014; Galon et al., 2012; Pages et al., 2018). Although there were notable differences in the composition of immune cell types across individual patients, the proportion of immune vs. non-immune cells was not correlated with the frequency of any specific immune cell type. (**Figure S15**).

We performed principal component analysis (PCA) to determine if combinations of cell types correlated with cancer state or clinical outcomes. The first two principal components were more prevalent in CLR patients than in DII patients (t test p<0.01 and p<0.025, respectively, **Figure 3E**). The first principal component contained cell subpopulations that are found in a classically defined follicle (B cells, plasma cells and CD4^+^ T cells) (**Figure S16A**). Interestingly, in the second principal component, non-immune and non-tumoral cell clusters (adipocytes, lymphatics, stroma, nerves, smooth muscle, and vasculature) had a positive weight (**Figure 3F**). These cell types make up structural components such as vessels and large percentage of the total variation (15.2%) and that the two patient groups differed in their projection on this component suggested that underlying factors preferentially promote the abundance of structural cell types in the iTME of CLR patients but not in DII patients. Therefore, these cell types may contribute to the improved survival of CLR patients.

A CODEX image can be analyzed for the spatial coordinates of each cell within the tissue. We computed pairwise cell-cell contact frequencies and frequency-normalized contact likelihood ratios (log-odds ratios) (Goltsev et al., 2018) for cell clusters in both groups of patients. The most dominant pairwise cell-cell interactions were homotypic (e.g., B cells with B cells; DCs with DCs), and we did not observe significant differences in these cell-cell contacts between CLR and DII patients (**Figure S16B**). Because no pairwise cell-cell contacts were sufficient to distinguish patient groups, we considered below whether features beyond pairwise contacts, such as higher-level features of tissue architecture and/or immune cell activation states, are attributes that define antitumoral immune responses and survival in this CRC cohort.

### Characteristic cellular neighborhoods of the CRC iTME are conserved across patient groups

In addition to the mere presence of tumor cells, immune cells and other microenvironmental components of the iTME, their spatial organization should provide insights into how they influence tumor development, progression, therapeutic response, or patient outcome. We reasoned that the dynamic spatial contexts of the tissue could be first approximated as cellular neighborhoods (CNs) consisting of tissue regions with a defined local composition of cell types. This approach was the simplest possible extension to treating the entire tissue as a homogeneous collection of cells. In addition, it made minimal assumptions (as far as could be said) regarding the permanence, shape or orientation of spatial contexts or their associated cellular dynamics, which we deemed important, not least because our data was a static, 2D approximation of a dynamic 3D tissue volume such as the iTME.

To identify CNs, cells were clustered based on the cell type composition of their 10 nearest spatial neighbors. The 10 nearest spatial neighbors for each individual cell, its so called “window” (**Figure 4A.1**), was identified using XY positions. The cell type composition, with respect to the 28 cell types that we had identified, was determined per window (**Figure 4A.2**), and windows with similar composition were grouped by *k*-means clustering (k=10) (**Figure 4A.3**). Granted, biologically, cells might exist in multiple neighborhoods simultaneously, or cell neighborhoods might “blend” one into another with no clear demarcation boundary. To simplify visualization, computation and interpretation of the spatial behavior of the tissue, any given cell was assigned here to a single CN. Each cell’s Voronoi representation was then colored according to the CN in which it resided (**Figure 4A.4**).

**Figure 4.**
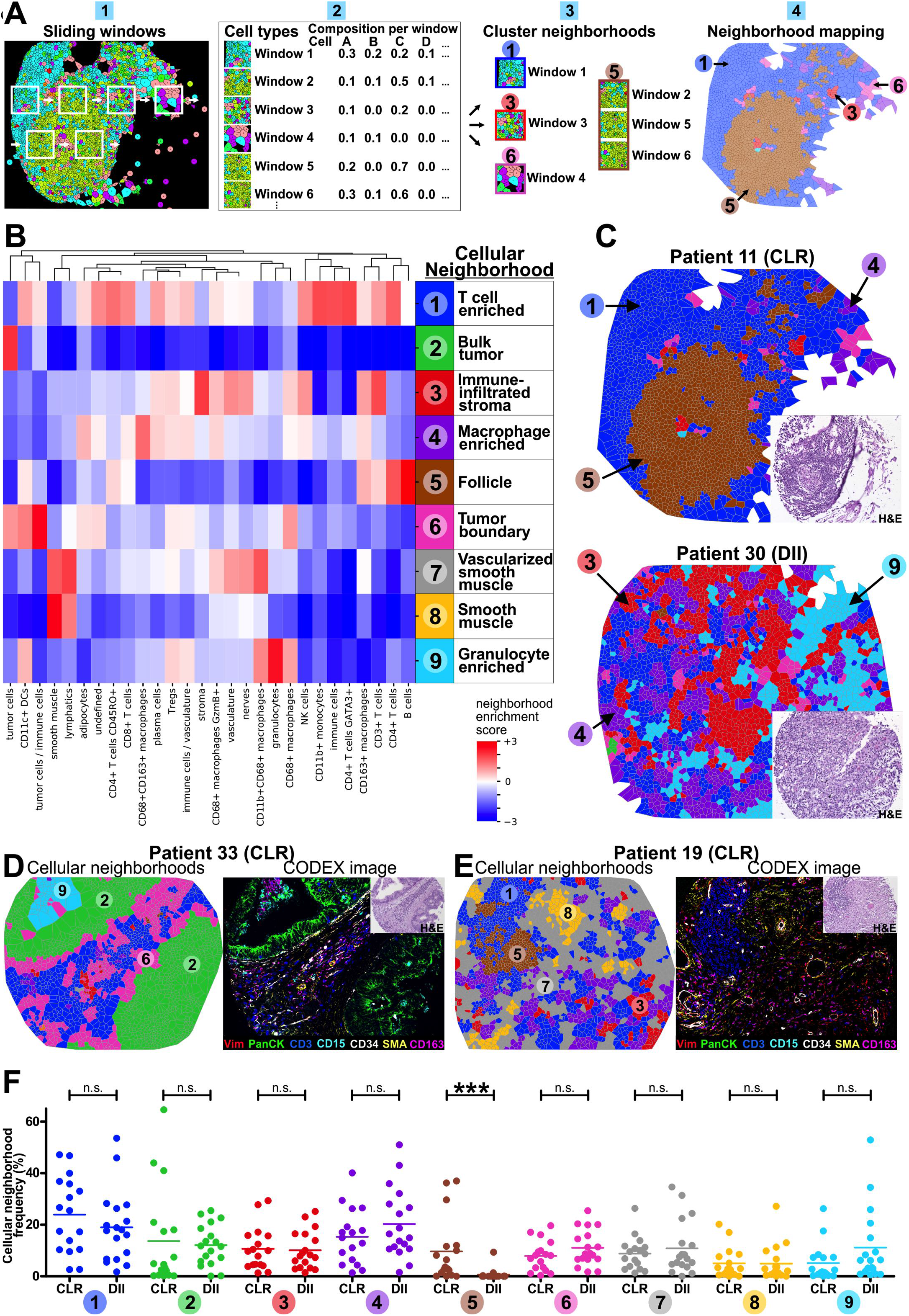
Identification of characteristic tissue CNs in the CRC iTME. **(A)** Schematic of automated computational CN identification. (1) The 10 nearest neighbors around every cell (its “window”) are identified. (2) Cell type composition per window is determined. (3) CNs are identified by clustering windows, and (4) each cell is assigned to a CN according to its window, and the Voronoi diagram of the cells in the tissue is colored by CN. **(B)** Identification of 9 distinct CNs based on the 28 original cell types and their respective frequencies (enrichment score) within each CN (pooled data from both patient groups). CN-1, T cells (blue); CN-2, bulk tumor cells (green); CN-3, immune-infiltrated stroma (red); CN-4, macrophages (purple); CN-5, follicle (brown); CN-6, tumor boundary (pink); CN-7, vascularized smooth muscle (gray); CN-8, smooth muscle (yellow); CN-9, granulocytes (cyan). **(C)** Representative Voronoi diagrams of CNs for CLR and DII patients. Insets, H&E images. **(D-E)** Representative Voronoi CN diagrams were selected to show the nine different CNs in two patients (left panels), with the corresponding seven-color overlay CODEX images (right panels). Vimentin (red); pan-cytokeratin (green); CD3 (blue), CD15 (cyan), CD34 (white), α-SMA (yellow), CD163 (magenta). Insets, H&E images. **(F)** Frequencies of CNs in CLR vs. DII patients. Each point represents the mean CN frequency from four TMA cores per patient (***p<0.001, Student’s *t*-test). See also Figures S16C-D, S17 and S18.

CNs were identified by clustering data from both patient groups together to maximize the recognition of common CNs. We found 10 distinct CNs in the CRC iTME that recapitulated the core tissue components, as validated on the original H&E-stained sections and fluorescent images in both patient groups (**Figures 4B-D** and **S17**). One of the 10 CNs was comprised mainly of imaging artifacts and therefore was removed from further analyses (data not shown). Surprisingly, except for a CN corresponding to the follicle, the remaining eight CNs were broadly present in both CLR and DII patients. When we applied this algorithm to each patient group separately, the identified CNs were still comparable across patient groups (**Figure S18**) and were therefore not an artifact of the data merging process. This led to the preliminary conclusion that the two extremes of the CRC iTME spectrum, while visually distinct as a result of the presence of TLS, shared underlying architectural components that could be defined by their characteristic local cell type frequencies.

The nine CNs recapitulated structures that directly related to components of the CRC iTME architecture as observed in H&E-stained tissue sections (**Figure 4B**). For example, CN-5 was enriched for B cells, CD4^+^ T cells, CD4^+^CD45RO^+^ T cells, CD11c^+^ DCs, and CD163^+^ macrophages and depleted of all other cell types. B cells and CD4^+^ T cells were the dominant cell types. When compared with H&E-stained images, CN-5 nearly perfectly aligns with the TLS (follicle) (**Figure 4B, CN-5 in heatmap; 4C, upper panel brown region**). Additionally, the analysis revealed previously unappreciated substructure in the remaining non-follicular leukocyte-dense tissue regions. Examples included T cell enriched CN-1 (**Figure 4B-C, blue regions**), macrophage enriched CN-4 (**Figure 4B-C, purple regions**), and granulocyte enriched CN-9 (**Figure 4B–4C, cyan regions**). The functional utility (to the immune system or tumor) of these repeatedly observed structures remains unknown, but the finding of such CNs underscores the unappreciated complexity, and coordination, of dynamic immune action against tumor presence as observed across multiple patients. In other words, a repeatable program is at play whether driven by the host, the tumor, or some interplay of the two.

The tumor itself was divided into two distinct CNs: CN-2 was mainly comprised of tumor cells (“bulk tumor”) (**Figure 4B and 4D, leftmost panel, green regions**), and CN-6 that contained tumor cells as well as CD11c^+^ DCs, CD68^+^ macrophages, T cell subsets and other immune and non-immune cell types (“tumor boundary”) (**Figure 4B and 4D, left panel, pink regions**). In the stroma, we discriminated three CNs: CN-3 was enriched in immune cells (**Figure 4B–4C, lower panel red regions**), CN-7 was vascularized smooth muscle (**Figure 4B and 4E, left panel gray regions**), and CN-8 mainly consisted of smooth muscle cells (**Figure 4B and 4E, left panel yellow regions**). Voronoi maps of CNs aligned well with fluorescent CODEX images (**Figures 4D-E**) and H&E images (**Figure S17**).

Since we observed a similar set of CNs in both CLR and DII, we determined whether the frequencies of any CNs differed in the CRC subtypes. We therefore computed the frequencies of each CN in each patient (**Figures 4F and S16C**). Except for CN-5 (follicle), which was highly enriched in CLR patients, none of the other CN frequencies differed significantly between patient groups. This indicates that the CNs we identified were common across patient groups and likely represent dynamic, conserved tissue compartments of CRC iTME. The question then becomes, are there other structures and relationships between these CNs that define the differences between CLR and DII, and, further, could these explain the respective patient outcome differences?

### Coupling of tumor and immune components in DII patients associated with coupling and fragmentation of T cell and macrophage CNs

The fact that the iTME could be decomposed into cell types and CNs found in both patient groups implied that despite the presence of follicles in CLR and their absence in DII, the core components of the CRC iTME are maintained across the spectrum of this cancer. What, then, are the differences in the organization of these components in each patient group towards such different outcomes?

Principal component analysis (**Figure 3E-F**) suggested that the biology driving the organization of the iTME coordinates distinct combinations of cell types to co-occur in each patient group. Presumably, this biology operates at multiple levels of abstraction, dynamically giving rise to combinations of cell types forming combinations of CNs, whose presence, behavior, and functions are mutually dependent, creating emergent function(s) from simpler components. Whereas PCA can identify explanatory axes of variation at one level of abstraction (for example, the cell type or the CN individually), it could not explicitly model coordinated coupling of cell types and CNs at multiple levels of abstraction. The question then becomes: How to describe a dynamically complex structure like the iTME, operating at these multiple levels, wherein the immune system, with its presumably regular order (i.e. the set rules by which it behaves), is in conflict with a tumor “strategizing” against it with genetic variation?

“Non-negative Tucker tensor decomposition” (Kim and Choi, 2007) was an attractive extension to PCA, because it enabled explicit modelling of the factors operating at the cell type level, CN level and the higher order organization of these factors. Variants of tensor decomposition have been previously applied to gene expression data collected across multiple tissues (Hore et al., 2016). Here, we applied this method to determine an optimal manner by which to decompose the tensors, higher-dimensional arrays representing the data across patients, into a collection of “CN modules” (combinations of CNs that partially contain similar combinations of cell types) and “cell type (CT) modules” (combinations of cell types that are found across similar neighborhoods). CN modules and CT modules “interact” in this specific approach to create “tissue modules” (interacting combinations of CT modules and CN modules that are found across similar patients) (**Figure 5A**, described below). The value of this approach is that it allows coordination at each level of abstraction (cell type, CN and tissue) to be viewed as interacting components that are combined, thereby giving a means to identify programs possibly driving the organization of the iTME in each patient group. The term “interacting” used here should be interpreted statistically and not as proof of physical cellular interaction or communication.

**Figure 5.**
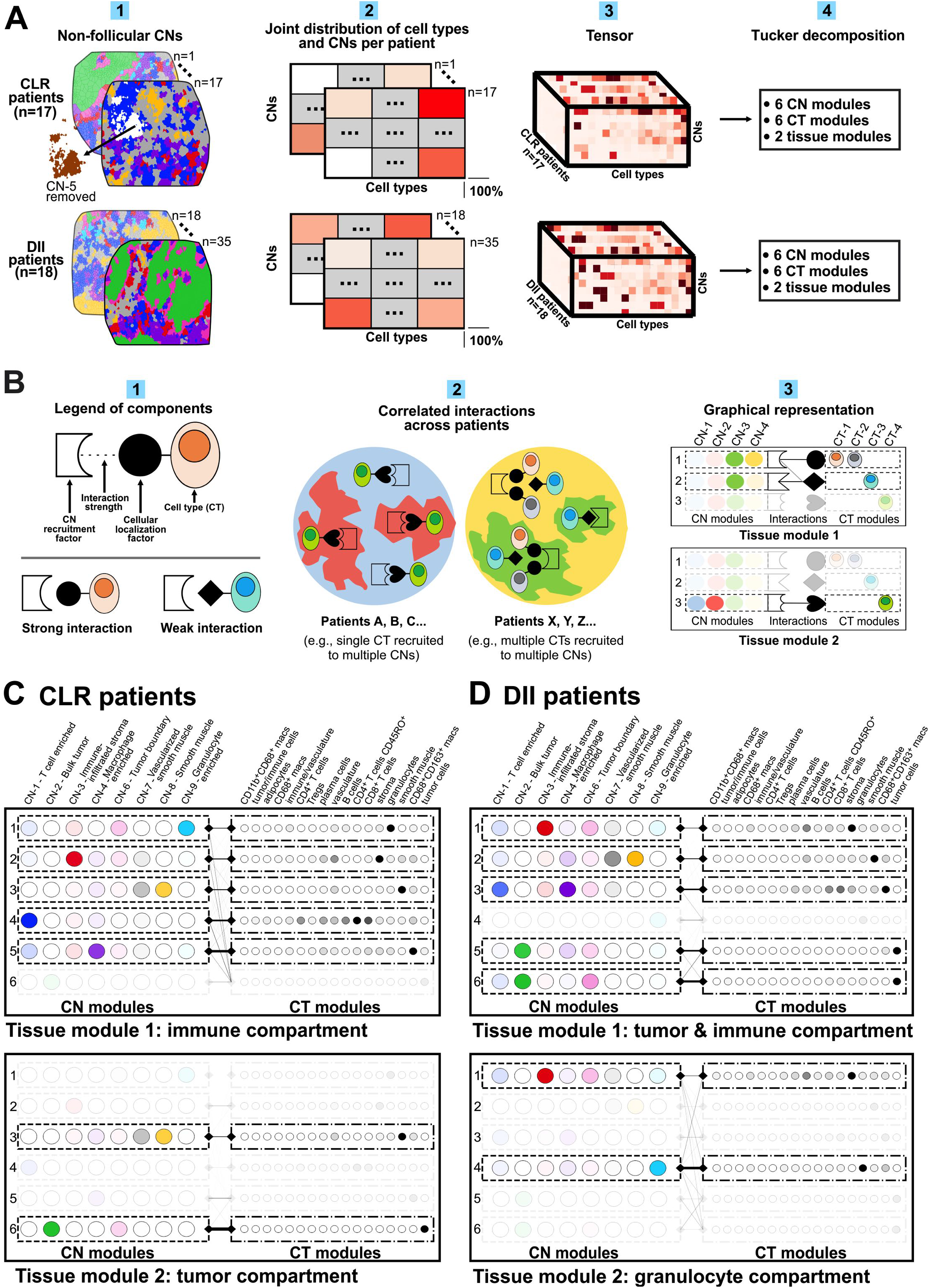
Tensor decomposition suggests differences in underlying programs spatially organizing the iTME. **(A)** Schematic of nonnegative Tucker tensor decomposition analysis. (1) The non-follicular CNs of the iTME were compared after removing CN-5. (2) For each patient, cell type (x axis) and CN (y axis) distribution are represented as a matrix. (3) Concatenating cell type and CN distributions from each patient along a third dimension (z axis) yields a tensor for each patient group. (4) Non-negative Tucker decomposition, applied to the tensors for each patient group individually, yielded 2 tissue modules, each comprised of six CN modules and six cell type (CT) modules. **(B)** Schematic illustrating the interpretation of the tensor decomposition output. (1) Legend of components: A CN module corresponds to a cell recruitment program utilized by the CNs comprising that module, and a CT module corresponds to a cell type localization program utilized by the cell types comprising that module. Different pairs of recruitment programs and localization programs interact to different strengths. (2) Different pairs of interacting recruitment programs and localization programs co-occur to form the tissue through balanced interactions between recruitment and localization factors. These combinations yield similar combinations of CNs and cell types within them across patients. (3) Graphical representation of tissue modules corresponding to combinations of interacting pairs, indicated by edges, of CN modules (left column) and CT modules (right column). CN modules and CT modules are common across both tissue modules. In each tissue module, the transparency of each CN module and CT module corresponds to the weight of the maximum edge of which it is part, i.e. indicating its contribution to that tissue module. **(C)** Decomposition results for CLR patients. Interacting pairs of CN modules and CT modules correspond to an “immune compartment” in tissue module 1 (top) and to a “tumor compartment” in tissue module 2 (bottom). **(D)** Decomposition for DII patients. Interacting pairs of CN modules and cell type modules correspond to an “immune & tumor compartment” in tissue module 1 (top) and to a “granulocyte compartment” in tissue module 2 (bottom). See also **Figure S19**.

DII patients do not have follicular structures; therefore, to remove any bias that such follicles alone might contribute to comparison, the CN-5 (follicle) data were excluded for the purposes of these analyses (**Figure 5A.1**). CLR and DII patient data were individually represented as a matrix of cell type distributions (x axis) and CNs (y axis) (**Figure 5A.2**). Two tensors (higher dimensional arrays – one for each patient group) were obtained by concatenating patients’ matrices along the z axis (**Figure 5A.3**), and the aforementioned tensor decomposition was performed (**Figure 5A.4**). The numbers of CN modules and CT modules (6 for each) were selected by visual identification of elbow points in the decomposition accuracy (**Figure S19**).

The relationships between CNs and cell types, as well as between CN modules and CT modules, determined by this decomposition, are then interpreted as an optimal description of the spatial iTME organization. Tensor decomposition was applied to patient groups individually, instead of combining patients and assessing ‘enrichment’ of a particular module. This is because we interpret the entire decomposition, including the relationships between modules, and not individual modules in isolation. This is then used to describe the differences between patient groups in the organization of the iTME.

One possible biological explanation for the tensor decomposition output is depicted as a schematic in **Figure 5B**: (1) The tissue is formed by the interaction of CN ‘recruitment factors’ (for example, cytokines) shared by multiple CNs to recruit cell types by interacting with cognate ‘cellular localization factors’ (for example, cytokine receptors) shared by multiple cell types (**Figure 5B.1, top aspect of the panel**). The term factor should be viewed in a statistical sense and could represent more complicated programs than a single ligand or receptor. Different factors can interact to different extents (**Figure 5B.1, lower aspect of the panel**). (2) Different interacting pairs of recruitment and localization factors are found together in the tissue, giving rise to the observed distribution of CNs and cell types (**Figure 5B.2)**. In the left region, the blue and red CNs share a recruitment factor (heart-shaped indentation), so share a common cell type (green) with a cognate localization factor (heart). In the right region, the orange and the gray cells share a localization factor (circle), so are found in multiple CNs. The green CN uses multiple recruitment factors, one shared with the yellow CN. Distinct interacting pairs of recruitment and localization factors co-occur across patients (red and blue found together, and yellow and green found together), each co-occurring collection of interacting pairs corresponding to a tissue module. These recruitment and localization factors are inferred from the tensor decomposition output, visualized as tissue modules comprised of CN modules and cell type (CT) modules, with interactions between them represented as edges (**Figure 5B.3).** Note that there is a common collection of CT modules and CN modules that are present to different extents in each tissue module. The contribution of each CN module and CT module to each tissue module is represented by its shading (**Figure 5B.3**). In tissue module 1 (top box), the CN module in the first row is interpreted as the recruitment factor with a circular indentation. This is because it contains yellow and green CNs, and there is a strong edge with the CT module containing the orange and grey cell types, and a weak edge with the CT module containing the blue cell type. The CN module with just the green CN (row 2) is interpreted as the recruitment factor with the square indentation. This is because that CN module does not contain any other CNs and has only one edge with one CT module containing the blue cell type. Since the red and green CNs are not found in the same patients, the CN module with the red and blue CNs and its cognate CT module with just the green cell type are faint in tissue module 1 and form tissue module 2. Note that the CN modules and the cell type modules are identified by their mutual dependence.

The identified CN modules and CT modules were common to each tissue module but contributed in the decomposition to different extents. We begin by describing the CN modules and CT modules, which are depicted twice in **Figure 5C-D** within the two tissue modules for each patient group. In CLR patients, the six CN modules that were identified corresponded to: 1) a module mainly containing CN-9 (granulocyte enriched) and CN-6 (tumor boundary); 2) a module mainly containing CN-3 (immune infiltrated stroma); 3) a module mainly containing CN-7 (vascularized smooth muscle) and CN-8 (smooth muscle); 4) a module mainly containing CN-1 (T cell enriched); 5) a module mainly containing CN-4 (macrophage enriched) (**Figure 5C, upper panel, left column, first five rows**), and 6) a module mainly containing CN-2 (bulk tumor) (**Figure 5C, lower panel, left column, last row**). The CT modules that were identified in CLR patients corresponded to: 1) a module mainly containing granulocytes; 2) a module mainly containing stroma and vasculature; 3) a module mainly containing smooth muscle and vasculature; 4) a module mainly containing CD8^+^ and CD4^+^ T cells, B cells, plasma cells and vasculature; 5) a module mainly containing CD68^+^CD163^+^ macrophages and plasma cells (**Figure 5C, upper panel, right column, first five rows**), and 6) a module containing tumor cells (**Figure 5C, lower right panel, last row**).

These CN modules and CT modules are combined to form two distinct tissue modules in each patient group. In CLR patients, the “immune compartment” tissue module consists of immune CN modules having their strongest edges with their cognate immune CT modules. For instance, CN module 5, consisting predominantly of CN-4 (macrophage enriched) had a strong edge with (obviously) CT module 5, consisting predominantly of macrophages (**Figure 5C, upper panel, fifth row, both columns**). Notably, in the immune compartment, the tumor neighborhood module and the tumor cell type module only had weak edges with the other neighborhood and cell type modules, and were therefore represented faintly (**Figure 5C, upper panel, sixth row, weak edges**). The second tissue module in the CLR patients was the “tumor compartment”. This primarily consisted of CN module 6, consisting predominantly of CN-2 (bulk tumor) and CN-6 (tumor boundary) that had a strong edge with CT module 6, consisting primarily of tumor cells (**Figure 5C, lower panel, sixth row**). Moreover, in this tissue module there was an additional edge between CN module 3, consisting of smooth muscle CNs and CT module 3, consisting of smooth muscle cell types (**Figure 5C, lower panel, third row**).

In DII patients, the six CN modules that were identified corresponded to: 1) a module mainly containing CN-3 (immune infiltrated stroma) and CN-6 (tumor boundary), 2) a module mainly containing CN-7 (vascularized smooth muscle) and CN-8 (smooth muscle), 3) a module mainly containing CN-1 (T cell enriched) and CN-4 (macrophage enriched), 4) a module mainly containing CN-9 (granulocyte enriched); 5) a module mainly containing CN-2 (tumor), CN-4 (macrophage enriched) and CN-6 (tumor boundary), and 6) a module mainly containing CN-1 (T cell enriched), CN-2 (tumor) and CN-6 (tumor boundary) (**Figure 5D, top panel, left column, rows 1-3 and 5-6; and lower panel, rows 1 and 4**).

The CT modules that were identified in DII patients corresponded to: 1) a module mainly containing stroma and vasculature; 2) a module mainly containing smooth muscle and vasculature; 3) a module mainly containing CD8^+^ and CD4^+^ T cells and CD68^+^CD163^+^ macrophages; 4) a module mainly containing granulocytes; 5) a module mainly containing tumor cells and CD68^+^CD163^+^ macrophages, and 6) a module mainly containing tumor cells (**Figure 5D, right columns**). We therefore labeled the tissue modules in DII patients 1) “tumor & immune compartment” and 2) “granulocyte compartment”.

Unlike in CLR patients, where the tumor compartment was distinct from the immune compartment, in DII patients there was a single compartment containing both tumor and immune components, and a distinct granulocyte compartment. This finding indicates that in DII patients there is a greater coupling between the formation of the tumor and the immune tissue compartments than in CLR patients. We speculate that in DII patients, as compared to CLR patients, the tumor might interfere with, or regulate, immune processes – although interestingly not those involving the granulocyte compartment. Furthermore, in CLR patients, the primary edges were between CN modules and their cognate CT modules. Only the tumor CT module had (weak) interactions with all the other CN modules (**Figure 5C, upper panel, right column, sixth row**). However, in DII patients, in addition to these weak edges connecting the tumor CT module to the immune CN modules, the tumor is itself part of the immune tissue module and other immune CNs are part of the two CN modules containing the tumor CNs. This could be interpreted as the tumor, in DII patients, extending the interactions present in CLR patients, to more deeply consolidate itself within the immune processes.

We noted interesting differences between the two patient groups in the CN modules and CT modules that interacted to form the tissue modules. In DII patients, one CN module had a high weight for both CN-1 (T cell enriched) and CN-4 (macrophage enriched) (**Figure 5D, upper left panel, row 3**), with its corresponding cell type module having a high weight for both T cells and macrophages (**Figure 5D, upper right panel, row 3**). In contrast, in CLR patients, no CN module had a high weight for both CN-1 and CN-4 and there existed no cell type module with a high weight for both T cells and macrophages (**Figure 5C**). This suggested that in DII patients, a common program drives the spatial organization of macrophages and adaptive immune cells into CN-1 (T cell enriched) and CN-4 (macrophage enriched), whereas in CLR patients, there were distinct programs driving the formation of these CNs. That these two CNs were part of the same CN module in DII patients also suggested that these two modules might be more interdigitated in DII patients than in CLR patients. We therefore computed the number of cells in CN-1 that had a cell in CN-4 as a nearest neighbor and vice versa. We divided this number by the total number of cells in the appropriate CN, thereby estimating their contact. We found that DII patients had a significantly higher contact between CN-1 (T cell enriched) and CN-4 (macrophage enriched) than did CLR patients (p = 0.046, Student’s t-test, **Figure S16D**). Despite this increased surface area, CN-1 and CN-4 are distinct CNs and are represented within each patient group individually (**Figure S18**).

In DII patients, two CN modules had a high weight of CN-2 (bulk tumor): one alongside CN-6 (tumor boundary) and CN-1 (T cell enriched), and one alongside CN-6 and CN-4 (macrophage enriched) (**Figure 5D**). In contrast, in CLR patients, no CN modules contained a high weight for both tumor CNs and other CNs (**Figure 5C**). This suggests that the molecular programs driving the recruitment of cells to the DII tumor neighborhoods (CN-2 and CN-6) could also drive the recruitment of cells to CN-1 (T cell enriched) and CN-4 (macrophage enriched). Therefore, the interference of the tumor in the immune processes of the DII iTME may restrict the spatial compartmentalization of T cells and macrophages.

Taken together, the tensor decomposition indicates that there are differences in the underlying programs that result in a distinct spatial organization of the iTME in CLR and DII patients. Moreover, it suggested that in DII patients, there was an increased coupling between the tumor and immune processes associated with increased coupling and spatial contact between T cell and macrophage CNs.

### Neighborhood-specific expression of functional markers on T cell subsets shows altered T cell processes in the bulk tumor

Understanding the iTME requires appreciating cells not only in terms of simple phenotypic descriptors, but also in terms of their functional states, and we would expect the same to be true for CNs. T cells exhibit diverse functional states in the iTME, often indicated by their expression of functional markers. Since the balance of these functional states is essential for a successful antitumoral immune response (Wherry, 2011), we would expect that the relative proportions of T cells expressing different functional markers is an indicator of the functional state of a given CN that could be relevant for patient outcomes.

The CD4^+^/CD8^+^ T cell ratio is often used as a simple measure to determine the overall balance of T cell function in cancer, providing prognostic information when measured in the tumor and tumor-immune interface (Shah et al., 2011; Wang et al., 2017a). The “tumor” and the “tumor-immune interface” are both likely composed of finer sub-structures that were not addressed in previous studies. We therefore computed and visualized the frequencies of CD4^+^ and CD8^+^ T cells in each CN for each patient (**Figures 6A**). Although the tensor decomposition identified structure by assessing variation present over multiple patients, overlaying T cell frequencies onto the CN Voronoi diagrams provides an intuitive visual interpretation of the tensor decomposition results. For instance, CD4^+^ and CD8^+^ T cells appear in similar combinations of CNs, which is expected since they were part of the same CT module. Similarly, while we see both CN-1 and CN-4 in the bottom panels, amongst immune neighborhoods we see predominantly CN-1 in the top Voronoi diagrams. This reflects that in the decomposition there is a neighborhood module with only CN-1, and another one with predominantly CN-4 and a weak presence of CN-1 (as defined in **Figure 5C, top left panel, rows 4-5**). Additionally, in **Figure 6A** there is a higher frequency of CD4^+^ and CD8^+^ T cells in CN-1 (blue) compared to CN-6 (pink) or CN-2 (green) in the top panels, which reflects that in **Figure 5C, top panel**, there is no edge between the cell type modules containing the T cells, and the neighborhood module containing CN-2 (bulk tumor). Furthermore, in the lower panels of **Figure 6A**, many of the immune neighborhoods appear in the same tissue section, as expected, since these are part of the same tissue module, the “immune compartment” (as defined in **Figure 5C, top left panel**).

**Figure 6.**
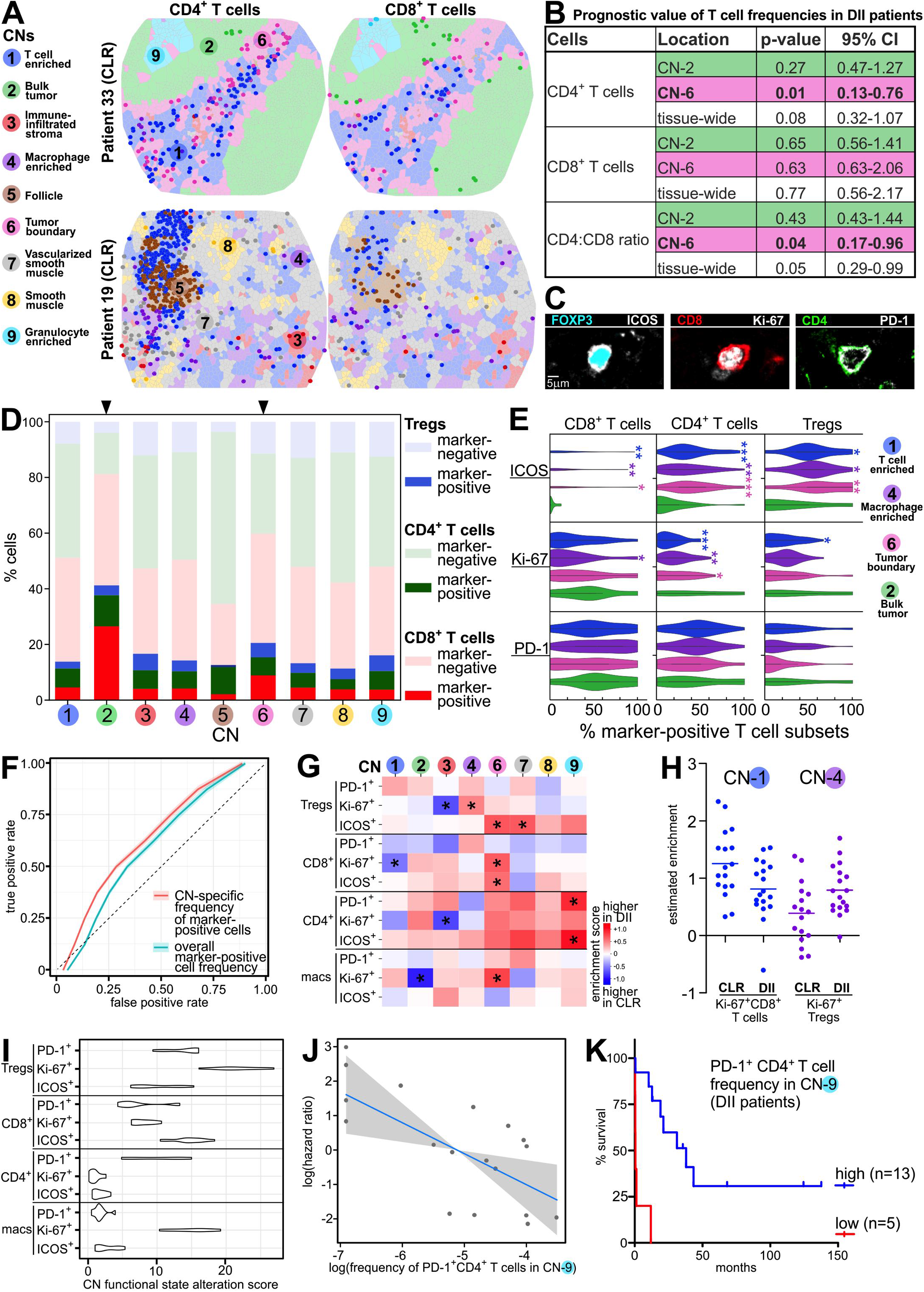
CN functional states are indicators of antitumoral immunity. **(A)** Example Voronoi diagrams of TMA spots, colored by CN. with CD4^+^ (left) and CD8^+^T cells (right) overlaid in each CN as points of the corresponding CN color. **(B)** Table of Cox proportional hazards regression results for T cell frequencies in indicated CNs. Each CN-specific frequency was tested individually in a distinct model (DII patients; n=18, 13 deaths). **(C)** Example staining for ICOS, Ki-67 and PD-1 on different T cell subsets. **(D)** Relative proportions of CD4^+^ (FOXP3^−^) Tcells, CD8^+^T cells, and Tregs positive for at least one of the functional markers ICOS, Ki-67. and PD-1 in each CN. Pooled data from all patients are shown (cell numbers for CN-1,17,822; CN-2,735; CN-3,4031; CN-4,11753; CN-5,4695; CN-6.2681; CN-7,4504; CN-8,1368; CN-9, 2884). **(E)** Violin plots of CN-specific cell type frequencies of marker-positive CD4^+^ T cells, CD8^+^T cells, and Tregs in CN-1, CN-2, CN-4, and CN-6. Asterisks indicate significant differences in the CN-specific cell type frequency underneath compared to the frequency in CN-2 (bulk tumor), tested across patients (*p<0.05, **p<0.01, ***p<0.001. Student’s *t*-test). **(F)** Receiver operating characteristic curves comparing the performance of L1-regularized logistic regression classifiers (CN-specific cell type frequency vs. overall frequency of marker-positive cells) over repeated hold-out samples to classify patients by group. **(G)** Heatmap of estimated differential enrichment coefficients (*p<0.05, not adjusted for multiple tests). A positive coefficient (red) indicates that the corresponding cell type is more enriched in DII patients in the given CN. (H) Estimated enrichment of Ki-67^+^ CDS^+^T cells in CN-1 and Ki-67^+^ Tregs in CN-4 for each patient. (I) Estimated CN activity alteration score for each cell type. Variation corresponds to the distribution of the score across 10 resampling iterations. (J) Partial residual plot of the log frequency of PD-1^+^ CD4^+^T cells in CN-9 vs. the estimated log hazard ratio with respect to overall survival in DII patients (p = 0.006; n=18, 13 deaths; Cox proportional hazards regression). A pseudocount of 0.001 was added to the frequency for all patients when logarithms were computed. (K) Kaplan-Meier curves for overall survival in DII patients corresponding to the best splitting of DII patients into two groups along the CN-9-specific frequency of PD-1^+^ CD4^+^T cells. See also **Figures S20 and S21**.

Are the frequencies of CD4^+^ and CD8^+^ T cells or the CD4^+^/CD8^+^ T cell ratio in CN-2 and CN-6 (bulk tumor and tumor boundary, respectively), or the tissue-wide frequencies of these cell types, associated with overall survival? We performed these tests only for DII patients, because there was insufficient mortality (only four deaths) in the CLR patient group to perform the analysis. Of these conditions, the CD4^+^ T cell frequency and the CD4^+^/CD8^+^ T cell ratio in CN-6 were significant prognostic factors (**Figure 6B, significant p-values in rows 2 and 8**), whilst the tissue-wide showed a prognostic trend. This highlighted the importance of T cell activity in CN-6 (tumor boundary) in the antitumoral immune response.

In addition to quantifying frequencies and ratios of T cell subsets, CODEX visualization allowed the simultaneous investigation of cellular functional states based on measured levels of key activation, co-stimulatory, and checkpoint molecules. We manually gated the functional markers PD-1, Ki-67 (proliferation marker), and inducible costimulator (ICOS) on the T cell subsets (**Figures 6C**, **S9, and S20**). We then quantified the relative proportions of marker-positive (PD-1^+^, Ki-67^+^, ICOS^+^) T cell subsets across the nine CNs, pooling cells from all patients (**Figures 6D**). These functional markers were not included during prior identification of cell types or CNs.

In CN-2 (bulk tumor), the proportion of T cells expressing at least one of these three markers was approximately twice as high as in any other CN (**Figure 6D, CN-2, second column**). The extent of this enrichment intratumorally, even in comparison to the tumor boundary (**Figure 6D, CN-6, sixth column**), was striking. We also observed that amongst T cells expressing any of the three functional markers, the CD4^+^/CD8^+^ T cell ratio was lower in tumor neighborhoods CN-2 and CN-6. In addition to there being changes in the frequency of “functional” CD4^+^ and CD8^+^ T cells and Tregs (as indicated by marker positivity) between CN-2 and other CNs, we would also expect there to be differences in the specific markers expressed by these marker-positive cells. For each of these subsets, and within each patient, we computed the relative proportion of ICOS^+^, Ki-67^+^ and PD-1^+^ cells out of marker-positive cells in CN-1, −2, −4 and −6, and visualized these as violin plots (**Figure 6E**). We observed that the proportion of marker-positive cells that were ICOS^+^ was significantly higher in CN-1, −4 and −6 compared to CN-2 (**Figure 6E, top row, asterisks**). This indicates that T cell subsets expressing the activation marker ICOS are less frequent in the bulk tumor as a proportion of marker-positive cells. In contrast, we found that the frequency of Ki-67^+^CD4^+^ T cells was highest in the bulk tumor (**Figure 6E, center panel**). No significant differences were observed for the PD-1^+^ subsets (**Figure 6E, bottom row**). These data point towards a prominent and sudden change in the inflammatory milieu within the bulk tumor compared to the tumor boundary and other iTME CNs and raises the question as to what tumor signals maintain this discrepant relationship.

### CN functional states with respect to T cells are distinct between CLR and DII patients and are correlated with survival

If the biological processes related to antitumoral immunity co-occurring in each CN are altered between patient groups, we would expect to observe concurrent changes in the frequencies of the relevant functional cell types therein. We first built two statistical models to test whether we can classify patients as CLR vs. DII using frequencies of T cells and macrophages expressing the functional markers PD-1, Ki-67 and ICOS. In the first model, we used only the overall frequency of each cell type across the non-follicular CNs. In the second model, we used the frequencies of these cells in each of the non-follicular CNs. Evaluating these models across repeated hold-outs (repeatedly “holding out” a randomly selected subset of the data, training the model on its complement, and evaluating it on the held-out subset), we observed that including spatial information (CN-specific cell type frequencies) improved the classification accuracy (**Figure 6F, red line**). Thus, the frequencies of cell types within certain compartments (we refer to these in the following as “CN-specific cell type frequencies”) could contain additional information distinguishing patient groups beyond what is contained in their overall frequencies.

This opened the question of whether CN-specific cell type frequencies are more different between patient groups than would be expected by the differences in the overall frequencies of the corresponding cell types (i.e. when was a cell type “differentially enriched” in a given CN between the patient groups?). This would be expected if there were differences in the functional state of a given CN, and not just changes in overall cellular composition of the iTME. Therefore, for each CN-specific cell type frequency, we estimated a linear model including as covariates both patient group and overall frequency of the corresponding cell type and visualized the estimated effects of the patient group as a heatmap, described below (**Figure 6G and see Methods**). According to this model, a significant coefficient (as indicated by asterisks in the heatmap) indicates that the given cell type is more enriched in a given CN in one group; i.e. that the CN-specific cell type frequency is higher in one group than what can be explained by changes in the overall frequency of that cell type (**Figure 6G**) (see next).

All CNs, except CN-8 (smooth muscle), exhibited significant differential enrichment of at least one functional cell subset between patient groups (**Figure 6G**). In CLR patients, there was an increased enrichment of Ki-67^+^ CD8^+^ T cells in CN-1, Ki-67^+^ macrophages in CN-2, and Ki-67^+^ Tregs and Ki-67^+^ CD4^+^ T cells in CN-3 (**Figure 6G, first 3 columns**). In DII patients, there was an increased enrichment of Ki-67^+^ Tregs in CN-4, ICOS^+^ Tregs, ICOS^+^ and Ki-67^+^ CD8^+^ T cells, and Ki-67^+^ macrophages in CN-6; ICOS^+^ Tregs in CN-7; and PD-1^+^ and ICOS^+^ CD4^+^ T cells in CN-9 (**Figure 6G, columns 4-8**).

That there was differential enrichment of four cell types in CN-6 compared to only one in CN-2 suggested that changes in immune activity in the tumor boundary CN could be implicated in the impaired survival of DII patients (**Figure 6G, compare CN-2 and CN-6**). In the tensor decomposition, we observed increased coupling between CN-1 (T cell enriched) and CN-4 (macrophage enriched) in DII patients. In addition, Ki-67^+^ CD8^+^ T cells were less enriched in CN-1, and Ki-67^+^ Tregs were more enriched in CN-4 in DII compared to CLR patients (**Figure 6H**), and activated ICOS^+^ Tregs were enriched in CN-6 in DII patients (**Figure 6G**). These data suggest that in DII patients, immunosuppressive activity is increased in CN-4 and CN-6, and that in CN-6 this could oppose the cytotoxic activity from Ki-67^+^ and ICOS^+^ CD8^+^ T cells. In contrast, in CLR patients, there is increased, unopposed cytotoxic activity in CN-1.

We noted that all the marker-positive cell subsets that were more enriched in any CN in CLR patients were Ki-67^+^ (**Figure 6G, first 3 columns, blue squares with asterisks**). This indicated that CN functional states are more altered between patient groups with respect to certain cell types than others, suggesting that these cell types could have different roles in iTME function in each patient group. We therefore computed a “CN functional state alteration score” for each marker-positive cell type. We compared the improvement in classification accuracy for a linear model trained to classify patients by group when the CN-specific frequencies for that cell type in the non-follicular CNs were included in addition to its overall frequency across repeated hold-outs (**see Methods**). We found that Ki-67^+^ Tregs were the most CN-activity altering cell type, followed by Ki-67^+^ macrophages (**Figure 6I**). Within CD4^+^ T cells, the PD-1^+^ subset was most activity altering, suggesting a role for this T cell subset in explaining the differences in the antitumoral immune response between patient groups.

The granulocyte enriched CN-9 stood out in the tensor decomposition analysis as being uniquely present in a tissue module distinct from the tumor in both patient groups. We also observed that PD-1^+^ and ICOS^+^ CD4^+^T cells were more enriched in CN-9 in DII than in CLR patients (**Figure 6G, rightmost column**). In addition, CD11c^+^ DCs were present in CN-9 and were differentially enriched between patient groups (**Figures 4B and S21**). These results suggested that certain processes occurring in CN-9, such as antigen presentation, could play key roles in the antitumoral immune response. Could this antitumoral response be driven by changes in the activity of CN-9 with respect to the cell types already identified above?

We assessed whether the frequencies of PD-1^+^ and ICOS^+^ CD4^+^ T cells in CN-9 were prognosticators of survival in DII patients. Neither the overall frequency of PD-1^+^ CD4^+^ T cells, nor the overall amount of CN-9 was a significant prognosticator. Notably, however, the PD-1^+^ CD4^+^ T cell frequency in CN-9 was a significant prognostic factor for overall survival (p = 0.006, Cox proportional hazards likelihood ratio test; 18 patients, 13 deaths) (**Figure 6J-K**). The positive association of the frequency of PD-1^+^ CD4^+^ T cells in CN-9 with overall survival (log(hazard ratio)) is visible in **Figure 6J**, and this count can be used to stratify patients as shown in **Figure 6K**. These findings could imply that specific events occurring in CN-9, related either to the production, maintenance, or function of PD-1^+^ CD4^+^ T cells are critical for the antitumoral immune response in CRC. Furthermore, although we could only compute the association with survival in DII patients, CN-9 is present in both groups, demonstrating its relevance to all CRC patients. The only other features tested for association with survival (excluding those in **Figure 6B**) were the frequencies of Ki-67^+^ Tregs in CN-4, ICOS^+^ Tregs in CN-6 and CN-7, as well as Ki-67^+^CD68^+^CD163^+^ macrophages in CN-6, because these, in addition to PD-1^+^ CD4^+^ T cells in CN-9, were the five most predictive features in the classification model of **Figure 6F (Figure S22).**

Taken together, these data indicate that the functional states of CNs are different between CLR and DII patients, and that CN functional states are potentially functionally relevant for the antitumoral immune response. Specifically, the functional state of a granulocyte-enriched CN, indicated by its frequency of PD-1^+^CD4^+^ T cells, was associated with overall survival in DII patients.

### Correlated CN functional states suggest immunosuppressive inter-CN communication and altered communication network in DII patients

The tensor decomposition showed that different CNs can recruit similar combinations of cell types, which could be interpreted as a form of communication between these CNs. We expected therefore that there would be other forms of communication between CNs that would give rise to correlations in CN-specific cell type frequencies between multiple CNs. We had observed that the functional states of CNs, as approximated by the frequencies of functional T cell subsets, were associated with survival outcomes. Therefore, communication between CNs that gave rise to correlated functional states (as approximated by the frequencies of functional T cell subsets), and changes in it, could be particularly important in the antitumoral immune response. This communication could be mediated by biological processes, such as immune cell infiltration, antigen presentation, cytokine production, metastasis, or as yet to be determined processes.

In CN-1 and CN-4 (T cell enriched and macrophage enriched, respectively), we observed opposing changes in the enrichment of Ki-67^+^ Tregs and CD8^+^ T cells between patient groups (**Figure 6G-H**), that is, Ki-67^+^ Tregs were more enriched in CN-4 in DII patients, and Ki-67^+^ CD8^+^ T cells were more enriched in CN-1 in CLR patients. Given that Tregs are capable of suppressing CD8^+^ T cell activity (Chen et al., 2005), we looked into whether the frequency of Tregs in CN-4 was correlated with the frequency of proliferating (Ki-67^+^) CD8^+^ T cells in CN-1 in each patient group separately.

We observed a significant negative correlation only in DII patients (**Figure 7A**), suggesting that a suppressive program involving Tregs and CD8^+^ T cells across CN-1 and CN-4 was only active in this patient group (see discussion for implications). Moreover, this finding demonstrated that between CLR and DII patients, there were changes in the communication across multiple CNs, with respect to their functional states.

**Figure 7.**
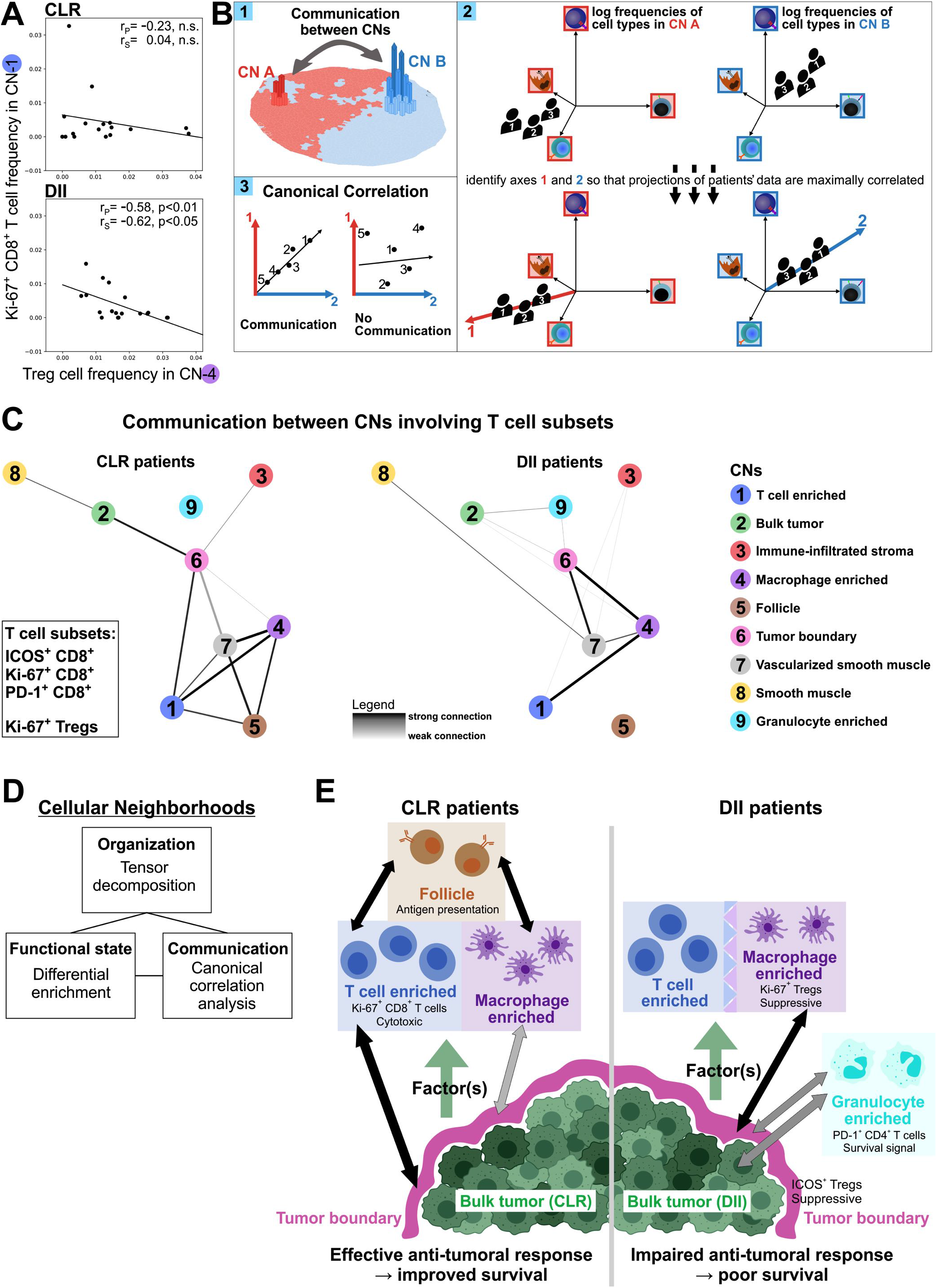
Altered inter-CN communication favors immunosuppression in DII patients. **(A)** Correlation of the frequency of Ki-67^+^ CD8^+^T cells in CN-1 (T cell enriched) and the frequency of Tregs in CN-4 (macrophage enriched) in each patient group. Spearman rank and Pearson correlation coefficients and p-values are shown. **(B)** Schematic illustrating canonical correlation analysis (CCA). (1) CN-specific cell-type frequencies in each CN are determined. (2) Linear combinations of pairs of CN-specific cell-type frequencies are identified that have maximized correlations. (3) If the canonical correlation between these linear combinations is higher than random, this is interpreted as communication between CNs in terms of the cell types assessed. **(C)** CCA was performed to identify communication between pairs of CNs involving T cell subsets. Specifically, canonical correlation with respect to the frequencies of ICOS^+^, Ki-67^+^, and PD-1^+^ CD8^+^T cells as well as Ki-67^+^ Tregs in each pair of CNs was compared to a permuted null distribution within each patient group. Those pairs of CNs whose observed canonical correlation with respect to these cell types was higher than 90% of permutations (permutation p-value <0.1) were connected by edges and visualized as a graph. **(D)** Conceptual framework for interpreting the CRC ITME architectural dynamics. **(E)** Model of differences in the iTME between CLR and DII patients with respect to CN organization (based on **Figure 5**), cellular function (based on **Figure 6**), and inter-CN communication (based on **Figures 7A-C**).

Should there not also be differences in the coordination of activity with respect to multiple cell subsets and across other pairs of CNs? We explicitly mapped the communication between different CNs involving T cell processes by performing canonical correlation analysis (CCA, **see Methods**) (Hardoon et al., 2004). CCA has been used to identify common signals across multiple data types, for example in multi-omic analyses (Witten and Tibshirani, 2009) and so we used it to find common signals across multiple CNs, with respect to their CN-specific frequencies. Briefly, for any given pair of CNs, the frequencies of the cell types of interest are computed within those CNs (**Figure 7B.1**). CCA then identifies the two combinations of CN-specific cell type frequency variations, one from each CN, such that their correlation is maximized (**Figure 7B.2**). This maximal correlation is the canonical correlation between two CNs, which we used as a proxy for functionally relevant communication (**Figure 7B.3**). We applied CCA to the frequencies of PD-1^+^, Ki-67^+^ and ICOS^+^ CD8^+^ T cells as well as Ki-67^+^ Tregs in each pair of CNs. In both patient groups, we identified pairs of CNs that likely communicate by comparing the observed canonical correlations to a permuted null distribution. These CN communication relationships were visualized as a graph of nodes corresponding to CNs, and edges indicative of communication for each patient group (**Figure 7C**).

There were interesting differences between patient groups in the communication networks of CNs with respect to functional T cell subsets. In both CLR and DII patients, CN-6 (tumor boundary) seems to play a central role. In CLR patients, CN-6 was strongly connected to CN-1 (T cell enriched), CN-2 (bulk tumor), and weakly connected to CN-4 (macrophage enriched) (**Figure 7C, left graph**). CN-4 and CN-1 were not directly connected to CN-2, indicating that functional T cell subsets in CN-1 and CN-4 could be communicating with the bulk tumor via the tumor boundary. In contrast, in DII patients, CN-6 was not connected to CN-1 but was strongly connected to CN-4 and weakly connected to CN-2 (**Figure 7C, right graph**). This is consistent with the communication between CN-1 and CN-6 having been rerouted via CN-4 in DII patients, which we showed was having immunosuppressive communication with CN-1, and had increased enrichment of Ki67+ Tregs (**Figure 6G**). The fact that the edge between CN-6 (tumor boundary) and CN-2 (bulk tumor) was weaker in DII patients suggests that communication with respect to T cells between the tumor and tumor boundary could be disrupted. In addition, only in DII patients were there connections between CN-9 (granulocyte enriched) and the tumor CNs. As the frequency of PD-1^+^ CD4^+^ T cells in CN-9 was associated with survival in DII patients (**Figure 6J-K**), our data are consistent with a role for CN-9 in T cell-mediated antitumoral responses. Finally, in CLR patients, CN-5 (follicle) was connected to CN-1, CN-4 and CN-7 (vascularized smooth muscle), indicating that the processes occurring in the follicle influence T cell activity in multiple CNs.

In summary, in DII patients, an immunosuppressive program was at play between CN-1 (T cell enriched) and CN-4 (macrophage enriched). Moreover, the network of communication between CNs with respect to functional T cell subsets was altered between patient groups, in a way that suggested that this immunosuppressive program could be affecting the phenotypes of functional T cell subsets in the tumor boundary and bulk tumor. Thus, the increased spatial contact between CN-1 and CN-4 as well as the changes in organization of CNs and cell types observed in the tensor decomposition could be playing a role in the impaired outcomes of these patients.

## DISCUSSION

The iTME is a dynamic system in which the combination of immune cell type, location and functional orientation leads to a tumor-rejecting or tumor-promoting environment (Chen and Mellman, 2017; Fridman et al., 2017). Naturally, the question arises as to how a tumor avoids immune action, such as by T cell exclusion (Joyce and Fearon, 2015) or the increased prevalence of certain cell types, including Tregs, macrophages and myeloid-derived suppressor cells (Gajewski et al., 2013; Hanahan and Weinberg, 2011; Kumar et al., 2016; Munn and Bronte, 2016). Several recent studies have characterized tumor-immune phenotypes in detail, but how spatial organization and the associated crosstalk between iTME components determine the effectiveness of the antitumoral immune responses has not yet been determined (Kather et al., 2017; Kather et al., 2018; Newell and Becht, 2018).

This current study probed the organization, functional states and communication within and between the CNs of the CRC iTME (**Figure 7D**). How do differences in these coordinated behaviors associate with survival outcomes? We developed FFPE-CODEX and computational approaches to describe the spatial organization and corresponding coordinated tissue behavior in a cohort of advanced-stage CRC patients with survival-associated iTME architectures: CLR (improved survival) vs. DII (poor survival).

In the iTME, T cells and macrophages are among the most abundant immune cells and are closely related to clinical outcome (Fridman et al., 2012; Fridman et al., 2017; Kather et al., 2017; Kather et al., 2018). This was supported by analysis of our CRC cohort, which identified T cell- and macrophage-enriched CNs. In DII patients, these two CNs were more interdigitated, with T cells and macrophages having closer physical contact, more mixing and increased information sharing between T cells. Consistent with a less spatially compartmentalized underlying organization of the iTME, immune and tumor compartments co-occurred in the tissue modules observed in the DII patients. These findings suggest an underlying molecular program whereby the tumor in DII patients directly interferes in the spatial organization of iTME compartments, highlighting different strategies for immune escape employed by tumors from DII and CLR patients. Critically, the differences in immune postures across the spectrum strongly imply that a “once size fits all” approach to immunotherapy in CRC would be shortsighted. After all, if different immune postures define different stages of how the tumor mitigates immune action, then distinct therapeutic modalities should be applied depending on the “stage” of the tumor’s interference. In other words, results such as those presented here provide a method of diagnostic subclassification based on mechanistic inferences from architecture, and could lead towards more nuanced therapeutic interventions.

A case in point is how the altered organization of the T cell and macrophage CNs in DII patients was accompanied by changes in their functional states as approximated by CN-specific frequencies of functional marker-positive T cell subsets. Unlike CLR patients, whose T cell-enriched CN was more cytotoxic (enrichment of Ki-67^+^CD8^+^ T cells), in DII patients the macrophage-enriched CN was more immunosuppressive (enrichment of Ki-67^+^ Tregs). Taken together, these alterations suggest an explanation for poor survival in DII patients: Tumor cells release factors that couple T cell- and macrophage-enriched CNs. This shifts the macrophage-enriched CN towards an immunosuppressive phenotype, which inhibits the cytotoxic activity of the T cell-enriched CN, limiting the antitumoral immune response (**Figure 7E**). Given this, as noted above, distinct modalities of intervention might be contemplated when this immune “module” is observed, or action taken prior to prevent its formation. Future studies that address the proteomic and metabolomic content of different iTME substructures, such as micro-dissected bulk tumor or tumor boundary regions, will be needed to identify the tumor-specific factors leading to the protumor environment observed in DII patients. Once such factors and their spatiotemporal distribution are identified, it will be necessary to determine whether manipulating these factors alters the immune response and leads to improved survival.

Recent studies of immunophenotypes in CRC have shown that the immune cell density is higher at the tumor invasive margin than in the tumor center (Bindea et al., 2013; Mlecnik et al., 2016). For this reason, and because TLS only occur at the invasive margin, we focused on the iTME at the invasive margin when creating our CRC TMAs. We found that the tumor boundary CN was an important site of antitumoral activity, with the frequency of CD4^+^ T cells and the CD4^+^/CD8^+^ T cell ratio having significant prognostic values in DII patients. Furthermore, DII patients showed increased enrichment of several unexpected cell types in the tumor boundary CN, including activated ICOS^+^ Tregs, which were recently shown to be increased in the hepatocellular carcinoma iTME and predictive of reduced survival (Tu et al., 2016). Interestingly, in DII patients there was direct communication between T cells in CN-6 (tumor boundary) and Tregs in CN-4 (macrophage enriched). This contrasts with CLR patients, where T cell frequencies were correlated in CN-6 and CN-1 (T cell enriched). These results suggest that ICOS^+^ Tregs and altered T cell exchange within the tumor boundary mediate the suppressed antitumoral immune response in DII patients (**Figure 7E**). However, the identities of the recruitment and retention signals that originate in the tumor boundary, how they act on ICOS^+^ Tregs, and whether blocking these signals improves survival, are unknown.

Intratumoral neutrophil granulocytes are generally associated with reduced survival of patients within solid tumors (Coffelt et al., 2016). However, the correlation of these cells with survival in CRC patients is controversial, with recent studies demonstrating both pro- and antitumoral immune activity for neutrophils (Berry et al., 2017; Rao et al., 2012). We identified CN-9 as a granulocyte enriched CN; it formed a distinct tissue module in DII patients, but was part of the immune tissue module in CLR patients. Within this granulocyte enriched CN, a higher frequency of PD-1^+^CD4^+^ T cells was observed in DII patients than in CLR patients, and a high frequency was associated with improved survival in this high-risk CRC patient group, whereas the overall frequencies of these cell types were not. Furthermore, in DII patients, T cell communication occurred between CN-9 and both the bulk tumor and tumor boundary CNs.

Taken together, our results imply that the granulocyte enriched CN in DII patients follows a distinct tissue molecule program that decouples it from the tumor/immune compartment and from information exchange with the tumor. A possible explanation for the improved survival in DII patients with higher frequencies of PD-1^+^CD4^+^ T cells in CN-9 is neutrophil-mediated destruction of tumor cells and antigen presentation to these PD-1^+^CD4^+^ T cells (**Figure 7E**). Future studies should therefore address the clinical implications of a granulocyte compartment for antigen presentation in CRC and other tumors and whether this spatial structure is targetable by therapeutics.

Our findings provided a model for how the iTME is organized to facilitate an effective antitumoral immune response in CLR patients, and how it is altered in DII patients. First, the tensor decomposition indicated that there are differences in the underlying biology organizing the iTME in DII patients compared to CLR patients. Specifically, in DII patients, there was increased coupling and interdigitation of T cell and macrophage CNs that could be attributed to recruitment factors provided by the tumor utilized by cell types occupying both of these CNs. In contrast, in CLR patients, the tumor appeared to interfere with the main immune processes to a lesser extent than observed in DII patients (**Figure 7E**).

Second, the CN-specific cell type frequencies indicated differences in the activities of functional T cell subsets in each CN. Specifically, in DII patients, CN-4 (macrophage enriched) had increased enrichment of Ki-67^+^ Tregs, whereas CN-1 (T cell enriched) had decreased enrichment of Ki-67^+^ CD8^+^ T cells. In addition, the activity of CN-9 (granulocyte enriched) was associated with improved survival outcomes.

Finally, CCA identified changes in the communication involving T cells across CNs. Specifically, in DII patients, CN-4 appeared to suppress immune activation in CN-1. Furthermore, in CLR patients there was direct communication between CN-1 and CN-6 (tumor boundary), suggesting T cell exchange between these CNs. In contrast, in DII patients, the more suppressive CN-4 was connected to the tumor boundary, suggesting a functional impact of this suppression on the antitumoral immune response.

These results are consistent with a model in which the tumor in DII patients provides a factor utilized by both macrophages and CD4^+^ T cells to promote the formation of their respective CNs, coupling these CNs. This coupling gives rise to more macrophages in the T cell CN, more T cells in the macrophage CN, more mixing of the two CNs, and a shift in the activity of the macrophage CN toward a more immunosuppressive phenotype. It also alters the exchange of T cells with the tumor from an otherwise activating adaptive immune CN. The activity of the granulocyte enriched compartment plays a functionally relevant role in antitumoral response, likely to be antigen presentation, that is altered between patient groups. Overall, an interpretation of the conclusion from **Figures 5C-D and S15** is that tumor-promoted fragmentation (dislocation) of immune contiguity is positively related to a worsening outcome for patients. In other words, a goal of the tumor’s immune evasion strategy is to prevent effective inter-CN communication which would culminate in the formation of functional immune modules, such as the frank appearance of follicular structures that are emblematic of CLR patients. Yet unknown are whether sub-clinical (and resolved) human CRC cases would provide insights into a completely effective immune architecture or posture. Considerations into the creation of animal models that mimic this situation would be informative, if possible.

Several caveats of course should be considered in the collection, interpretation, and bioinformatics analysis of imaging data. First, imaging and downstream analysis is affected by tissue quality; and, some tissue types may not be amenable for multiplexed fluorescence imaging due to autofluorescence. Poorly fixed tissues and non-specific antibodies can lead to low signal-to-noise ratios and potentially misleading staining patterns. These limitations can be improved by collaborating with pathologists to carefully select tissue regions and thoroughly validate antibodies across several positive and negative control tissues, but this process is costly. Here, we took extensive care to ensure reagent validity and patient selection. In addition, a large cohort of patients was initially screened for patient materials that fit strict inclusion and exclusion criteria. With regards to computational analyses, cell type identification and segmentation was performed with extensive validation and manual merging of identified clusters. This may be aided by computational correction for batch-dependent variation in staining intensity, as well as general-purpose neural network-based approaches for cell segmentation and cell type identification. Additionally, the performed neighborhood analysis was contingent upon a collection of CNs that could be readily identified and verified in the original imaging data. In other tissue types, CNs may be less well defined. For those situations, other statistical frameworks may improve identification of CNs and quantify their changes across different biological contexts. While we did not physically model CNs and the cell types within them, such models could eventually provide understanding(s) of tissue dynamics at the cell type and CN level, and the specific couplings between them. Other abstractions (for example, network models, field-based models or geospatial analyses) for articulating the spatial organization of tissue may provide complementary insights that better capture tissue behavior in other biological settings. Finally, while we treated the stroma and the bulk tumor as monolithic components this was due only to limitations in the markers applied. Accessing stromal or tumoral subtypes with additional markers, whether proteomic, genomic, or metabolic, will likely lead to a more fine-grained appreciation of substructures in these CNs as well.

In summary, our findings provide a model for how the CRC iTME is organized to facilitate an effective antitumoral immune response in CLR patients and interpretations of how it is impaired in DII patients. We provide a valuable dataset and conceptual framework for studying CRC spatial biology in a large, well-annotated cohort. Such a resource can be used to develop additional algorithms to identify clinically relevant factors and the molecular circuitry that underlies antitumoral immunity in CRC. Patient outcome was related to specific, repeated CN architectures, apparent dislocations of such CNs driven by tumor actions, appearance of tissue modules suggestive of immune suppressive states in some patients, and a coordinated change in the tumor interface comparing CLR to DII. The changing nature of the iTME across the CRC spectrum suggests manners by which antitumoral immunity might be enhanced through mechanistic understandings of how the emergent order links to function and dysfunction. The eventual linking of such favorable and unfavorable immunological attributes to patient outcomes will lead to the identification of prognostic spatial biomarkers and therapeutic strategies that shift high-risk CRC patients toward an antitumoral immune phenotype and amelioration of disease. The extension of such concepts to other cancers is warranted.

## Supporting information

Analysis code files

## ACKNOWLEDGMENTS

We thank Angelica Trejo, Han Chen, Kenyi Donoso, Nilanjan Mukherjee, Vishal Venkataraaman, and Gustavo Vazquez (Baxter Laboratory for Stem Cell Biology, Department of Microbiology and Immunology, Stanford University School of Medicine, Stanford, CA, USA) for excellent technical assistance and Sandrine Ruppen, Carmen Cardozo, and Dr. José Galván (Translational Research Unit and Compath, Institute of Pathology, University of Bern, Switzerland) for help with creating the TMAs. We are grateful to Dr. Anna Seigal (Mathematical Institute, University of Oxford, Oxford, UK University) for helpful discussions regarding tensor decomposition; to Dr. Julian Schardt (Department of Medical Oncology, Inselspital, University Hospital Bern, Switzerland) for helping obtain patient clinical information; and to Prof. Paul Bollyky (Department of Infectious Diseases, Stanford University School of Medicine, Stanford, CA, USA) for providing the biotinylated VG1 hyaluronan-detection reagent. We thank the patients for their consent to use their tissues for research. We would like to extend our gratitude to Dr. Sizun Jiang and Dr. Xavier Rovira-Clavé (Baxter Laboratory for Stem Cell Biology, Department of Microbiology and Immunology, Stanford University School of Medicine, Stanford, CA, USA) for critical comments on the manuscript. This work was supported by the US National Institutes of Health grants and sub awards: 2U19AI057229-16, 5P01HL10879707, 5R01GM10983604, 5R33CA18365403, 5U01AI101984-07, 5UH2AR06767604, 5R01CA19665703, 5U54CA20997103, 5F99CA212231-02, 1F32CA233203-01, 5U01AI140498-02, 1U54HG010426-01, 5U19AI100627-07, 1R01HL120724-01A1, R33CA183692, R01HL128173-04, 5P01AI131374-02, 5UG3DK114937-02, 1U19AI135976-01, IDIQ17X149, 1U2CCA233238-01, 1U2CCA233195-01; The Department of Defense (W81XWH-14-1-0180 and W81XWH-12-1-0591); The Food and Drug Administration (HHSF223201610018C and DSTL/AGR/00980/01); Cancer Research UK (C27165/A29073); The Bill and Melinda Gates Foundation (OPP1113682); The Cancer Research Institute; The Parker Institute for Cancer Immunotherapy; The Kenneth Rainin Foundation (2018-575); The Silicon Valley Community Foundation (2017-175329 and 2017-177799-5022); The Beckman Center for Molecular and Genetic Medicine; Juno Therapeutics, Inc. (122401), Pfizer, Inc. (123214); Celgene, Inc. (133826 and 134073); Vaxart, Inc. (137364); and the Rachford & Carlotta A. Harris Endowed Chair (G.P.N.). C.M.S. was supported by an Advanced Postdoc Mobility Fellowship from the Swiss National Science Foundation (P300PB_171189 and P400PM_183915), and an International Award for Research in Leukemia from the Lady Tata Memorial Trust, London, UK. D.J.P. was supported by an NIH T32 Fellowship through Stanford’s Department of Epithelial Biology (AR007422), an NIH F32 Fellowship (CA233203), a Stanford Dean’s Postdoctoral Fellowship, and Stanford’s Dermatology Department. S.S.B. was supported by a Bio-X Stanford Interdisciplinary Graduate Fellowship and Stanford’s Bioengineering Department. G.L.B was supported by an NIH T32 Fellowship through Stanford’s Molecular and Cellular Immunobiology Program (5T32AI007290-34).

## AUTHOR CONTRIBUTIONS

C. M.S. conceived and coordinated the study. C.M.S. and D. J.P. designed and performed experiments, analyzed and interpreted data, created figures and wrote the manuscript. S.S.B. coordinated computational analyses. S.S.B and G.L.B conceived and computationally implemented the conceptual framework, analyzed and interpreted data, performed statistical analyses, created figures and wrote the manuscript. C.M.S., L.N. and I.Z. designed and created the TMAs. L.N. and I.Z. obtained clinical and pathological information. I.Z. provided advice on statistical analysis. P.C. and S.B. performed experiments and analyzed data. J.D. and D.R.M. analyzed and interpreted data. N.S. and Y.G. conceived of CODEX, built the computational image processing pipeline and provided technical advice. G.P.N. supervised the study and wrote the manuscript. All authors revised the manuscript and approved its final version.

## METHODS

### CONTACT FOR REAGENT AND RESOURCE SHARING

Further information and requests for reagents and resources should be directed to and will be fulfilled by the lead contact, Garry P. Nolan.

### EXPERIMENTAL MODEL AND SUBJECT DETAILS

A cohort of 715 patients who underwent surgery for primary colorectal cancer between 2003 and 2014 at the University Hospital Bern, Switzerland, was screened. Clinicopathological data for all patients were extracted from clinical and pathological reports. The peritumoral inflammatory reaction was retrospectively assessed in all patients by L.N., under the supervision of I.Z. and C.M.S., in a blinded fashion using digitally scanned H&E-stained tumor sections. The tumor invasive margin was assessed for the presence of CLR according to the Graham-Appelman (G-A) criteria (Graham and Appelman, 1990). Cases were categorized as either GA 0 (lymphoid aggregates absent), G-A 1 (occasional lymphoid aggregates with rare or absent germinal centers) or G-A 2 (intense reaction with numerous lymphoid aggregates and germinal centers). The overall density of the peritumoral immune infiltrate was determined according to the Klintrup-Mäkinen (K-M) score (Klintrup et al., 2005), with low-grade (absent or mild and patchy inflammatory infiltrate; score 0-1) or high-grade (dense, linear inflammatory infiltrate with destruction of cancer cell islets; score 2-3) scorings. Cases with pre-operative therapy, pathological tumor, nodes, metastasis (pTNM) stage 0-2, absent peritumoral inflammatory infiltrate (K-M score 0), and those with insufficient material or information (total of 566 cases) were excluded. From the remaining 149 cases, 62 patients were identified based on their unique pattern of peritumoral inflammation and split into two groups: CLR group (G-A 2, any K-M grade) vs. DII group (GA 0, K-M high-grade). Subsequently, 35 stage-matched cases (17 CLR vs. 18 DII) were selected matched for gender, age, and cancer type, location, and stage. The use of patient tissue samples and data was approved by the local Ethics Committee of the Canton of Bern (KEK 200/2014) and by Stanford’s Institutional Review Board (HSR 48803).

### EXPERIMENTAL METHOD DETAILS

#### Construction of tissue microarrays

FFPE tissue blocks were retrieved from the tissue archive at the Institute of Pathology, University of Bern, Switzerland. For the multi-tumor TMA, 70 unique different tissues were selected (54 different cancers and non-malignant tumors as well as 16 normal tissues; for details see **Figure S5** and **Table S4**). Tumor and normal tissue regions were annotated on corresponding H&E-stained sections by a board-certified surgical pathologist (C.M.S.). A next-generation TMA (ngTMA^®^) with 0.6 mm diameter cores was assembled using a TMA Grand Master automated tissue microarrayer (3DHistech).

For the CRC study, two independent 70-core ngTMAs were created, containing four 0.6-mm cores per patient. TMA cores were digitally annotated by L.N., under the supervision of C.M.S and I.Z., as follows: CLR group, two regions containing a tertiary lymphoid structure and two diffuse immune infiltrate regions per patient; DII group, four diffuse immune infiltrate regions per patient. TMAs were sectioned at 3 μm thickness, stained with H&E, and digitized using a Pannoramic P250 digital slide scanner (3DHistech).

Square glass coverslips (Electron Microscopy Sciences) were pre-treated with Vectabond™ (Vector Labs) according to the manufacturer’s instructions. Briefly, coverslips were immersed in 100% acetone for 5 min and then incubated in a mixture of 2.5 ml Vectabond™ and 125 ml 100% acetone in a glass beaker for 30 min. Coverslips were washed in 100 % acetone for 30 sec and air dried, baked at 70°C for 1 h, and stored at room temperature. The 4-μm thick sections of the TMAs were mounted on Vectabond™-treated coverslips and stored in a coverslip storage box (Qintay) at 4°C in a vacuum desiccator (Thermo Fisher) containing drierite desiccant (Thermo Fisher) until analysis.

#### Buffers and solutions

Buffers and solutions were prepared, filtered sterile using 500-ml 0.2-μm pore size filters and stored at room temperature unless otherwise specified. Staining solution 1 (S1): 5 mM EDTA (Sigma), 0.5% w/v bovine serum albumin (BSA, Sigma) and 0.02% w/v NaN3 (Sigma) in PBS (Thermo Fisher Scientific); storage at 4°C. Staining solution 2 (S2): 61 mM NaH_2_PO_4_ · 7 H_2_O (Sigma), 39 mM NaH_2_PO_4_ (Sigma) and 250 mM NaCl (Sigma) in a 1:0.7 v/v solution of S1 and doubly-distilled H_2_O (ddH_2_O); final pH 6.8-7.0. Staining solution 4 (S4): 0.5 M NaCl in S1. TE buffer: 10 mM Tris pH 8.0 (Teknova), 1 mM EDTA and 0.02% w/v NaN3 in ddH_2_O. Tris stock solution (for conjugation buffer), 50 mM, pH 7.2 (at room temperature) was prepared in ddH_2_O using Trizma HCl and Trizma Base according to Sigma’s Trizma mixing table. Buffer C (for conjugation): 150mM NaCl, 2 mM Tris stock solution, pH 7.2, 1 mM EDTA and 0.02% w/v NaN3 in ddH_2_O. CODEX 2.0 buffer (H2): 150mM NaCl, 10 mM Tris pH 7.5 (Teknova), 10 mM MgCl_2_ · 6 H_2_O (Sigma), 0.1% w/v Triton™ X-100 (Sigma) and 0.02% w/v NaN3 in ddH_2_O. Blocking reagent 1 (B1): 1 mg/ml mouse IgG (Sigma) in S2. Blocking reagent 2 (B2): 1 mg/ml rat IgG (Sigma) in S2. Blocking reagent 3 (B3): Sheared salmon sperm DNA, 10 mg/ml in H2O (Thermo Fisher). Blocking component 4 (BC4): Mixture of 57 non-modified CODEX oligonucleotides (see Table S2) at a final concentration of 0.5 mM each in TE buffer. BS3 fixative solution (BS3): 200 mg/ml BS3 (Thermo Fisher) in DMSO from a freshly opened ampoule (Sigma); stored at −20°C in 15-μl aliquots. TCEP solution: 0.5 M TCEP (Sigma) in ddH_2_O, pH 7.0. Rendering buffer: 20% DMSO (v/v) in H2 buffer. Stripping buffer: 80% DMSO (v/v) in H2 buffer.

#### Generation of CODEX DNA-conjugated antibodies

All pipetting was performed using LTS filter tips (Rainin). Maleimide-modified short DNA oligonucleotides (for sequences, see **Table S2**) were purchased from TriLink. Maleimide groups were deprotected by heating in toluene at 90°C for 4h (with exchange of toluene after 2h). Deprotected oligonucleotides were repeatedly washed in 100% ethanol, resuspended in buffer C, and aliquoted at 50 μg in 0.2-ml 8-strip tubes (E&K Scientific). Oligonucleotides were flash-frozen in liquid N_2_, lyophilized overnight in 900 ml Labconco™ FastFreeze™ Flasks (Thermo Fisher) using a FreeZone^®^ 4.5 Plus lyophilizer (Labconco) and stored until conjugation at −20°C in an airtight box containing desiccant. Conjugations were performed at a 2:1 weight/weight ratio of oligonucleotide to antibody, with at least 100 μg of antibody per reaction. All centrifugation steps were at 12,000g for 8 min, unless otherwise specified. Purified, carrier-free antibodies (for details on clones and manufacturers, see Table S1) were concentrated on 50 kDa filters and sulfhydryl groups were activated using a mixture of 2.5 mM TCEP and 2.5 mM EDTA in PBS, pH 7.0, for 30 min at room temperature. After washing the antibody with buffer C, activated oligonucleotide was resuspended in buffer C containing NaCl at a final concentration of 400 mM. Oligonucleotide was then added to the antibody and incubated for 2 h at room temperature. The conjugated antibody was washed by resuspending and spinning down three times in PBS containing 900 mM NaCl. It was then eluted by centrifugation at 3,000g for 2 min in PBS-based antibody stabilizer (Thermo Fisher) containing 5 mM EDTA and 0.1% NaN3 (Sigma), and stored at 4 °C.

#### CODEX antibody staining of FFPE tissue and poststaining fixation

Coverslips were handled using Dumont coverslip forceps (Fine Science Tools). For deparaffinization, coverslips were baked at 70 °C for at least 1 h, followed by immersion in fresh xylene for 30 min. Sections were rehydrated in descending concentrations of ethanol (100% twice, 95% twice, 80%, 70%, ddH_2_O twice; each step for 3 min). Heat-induced epitope retrieval (HIER) was performed in a Lab Vision™ PT module (Thermo Fisher) using Dako target retrieval solution, pH 9 (Agilent) at 97 °C for 10 min. After cooling to room temperature for 30 min, coverslips were washed for 10 min in 1x TBS IHC wash buffer with Tween^^®^^ 20 (Cell Marque). Tissues were encircled using polyacrylamide gel (Bondic), and nonspecific binding was blocked for 1 h at room temperature using 100 μl of blocking buffer [S2 buffer containing B1 (1:20), B2 (1:20), B3 (1:20), and BC4 (1:15)]. For each coverslip, DNA-conjugated antibodies were added to 50 μl of blocking buffer on a 50-kDa filter unit, concentrated by spinning at 12,000 g for 8 min, and resuspended in blocking buffer to a final volume of 100 μl. This antibody cocktail was then added to the coverslip and staining was performed in a sealed humidity chamber overnight at 4 °C on a shaker. After staining, coverslips were washed for 4 min in S2 and fixed in S4 containing 1.6% paraformaldehyde for 10 min, followed by three washes in PBS. Then, coverslips were incubated in 100% methanol on ice for 5 min, followed by three washes in PBS. Fresh BS3 fixative was prepared immediately before final fixation by thawing and diluting one 15-μl aliquot of BS3 in 1 ml PBS. Final fixation was performed at room temperature for 20 min, followed by three washes in PBS. Thereafter, coverslips were stored in S4 in a 6-well plate at 4 °C for up to two weeks, or further processed for imaging.

#### Immunohistochemistry

Sections were cut to 4 μm thickness and placed on frosted histology glass slides (Thermo Fisher). H&E stained sections were obtained from each FFPE block. Deparaffinization, rehydration, and HIER were performed on an ST4020 small linear stainer (Leica) as described above. Nonspecific binding was blocked for 1 h at room temperature using 100 μl of serum-free protein block (Agilent). Antibodies were diluted in 100 μl antibody diluent (Agilent), and sections were stained overnight in a sealed humidity chamber at 4 °C on a shaker. After staining, slides were washed for 10 min in 1x TBS IHC wash buffer with Tween^^®^^ 20 (Cell Marque), and specimens were covered with dual endogenous enzyme-blocking reagent (Agilent) for 5-10 min at room temperature, followed by washing for 10 min. Bound antibodies were then visualized using the HRP/liquid DAB+ substrate chromogen system (Agilent) according to the manufacturer’s instructions. Sections were counterstained with hematoxylin, followed by dehydration, mounting, and imaging in brightfield mode on a BZ-X710 inverted fluorescence microscope (Keyence).

#### CODEX antibody screening, validation and titration

Antibodies were first screened and validated in CODEX single-stainings on tonsil tissue or a multi-tumor TMA, with cross-validation by manual IHC (Figures S1 and Table S4). Briefly, after antibody staining and fixation, 100 nM of fluorescent DNA probe was added to the tissue in rendering buffer, containing 0.7 mg/ml sheared salmon sperm DNA, and was incubated at room temperature for 5-10 min, followed by washing with rendering buffer and S4 buffer. Coverslips were mounted onto glass slides with nail polish (Sally Hansen) or Cytoseal XYL (Thermo Fisher), dried in the dark and imaged on a BZ-X710 inverted fluorescence microscope. All validation was performed by or under the supervision of a board-certified surgical pathologist (C.M.S.) and confirmed with online databases (The Human Protein Atlas, Pathology Outlines) and the published literature.

#### CODEX multi-cycle reaction and image acquisition

Coverslips were removed from S4, and the tissue was covered with a small piece of cling film. The non-tissue containing parts of the coverslips were rinsed in ddH_2_O to remove salt residues and thoroughly dried using vacuum. Then, coverslips were mounted onto custom-made CODEX acrylic plates (Bayview Plastic Solutions; blueprints available upon request) using coverslip mounting gaskets (Qintay), creating a well in the acrylic plate above the tissue section for liquid storage and exchange. The cling film was removed, and the tissue was stained with Hoechst nuclear stain at a dilution of 1:1000 in H2 buffer for 1 min, followed by three washes with H2 buffer. The CODEX acrylic plate was mounted onto a custom-designed plate holder (blueprints available upon request) and secured onto the stage of a BZ-X710 inverted fluorescence microscope. Fluorescent oligonucleotides (concentration: 400 nM) were aliquoted in Corning™ black 96-well plates in 250 μl H2 buffer containing Hoechst nuclear stain (1:600) and 0.5 mg/ml sheared salmon sperm DNA, according to the multi-cycle reaction panels. For details on the order of fluorescent oligonucleotides and microscope light exposure times, see **Table S4**. Black plates were sealed with aluminum sealing film (VWR Scientific) and kept at room temperature during the multi-cycle reaction. The final concentration of fluorescent oligonucleotides in the tissue-containing imaging well corresponded to 80 nM (1:5 dilution in rendering buffer).

Automated image acquisition and fluidics exchange were performed using an Akoya CODEX instrument and CODEX driver software (Akoya Biosciences). During imaging, the tissue was kept in H2 buffer. Hybridization of the fluorescent oligonucleotides was performed in rendering buffer. After imaging, fluorescent oligonucleotides were removed using stripping buffer. Overviews of the TMA were acquired manually using a CFI Plan Apo λ 2x/0.10 objective (Nikon), and automated imaging was performed using a CFI Plan Apo λ 20×/0.75 objective (Nikon). For multicycle imaging of the TMA spots, the multipoint function of the BZ-X viewer software (Keyence) was manually programmed to the center of each TMA spot and set to 17 Z-stacks. Hoechst nuclear stain (1:3000 final concentration) was imaged in each cycle at an exposure time of 1/175s. Biotinylated VG1 hyaluronan-detection reagent was produced as previously described (Clark et al., 2011), used at a dilution of 1:500, and visualized in the last imaging cycle using streptavidin-PE (1:2500 final concentration). DRAQ5 nuclear stain (1:500 final concentration) was added and visualized in the last imaging cycle. After each multi-cycle reaction, H&E-stainings were performed according to standard pathology procedures, and tissues were reimaged in brightfield mode.

A multi-cycle experiment performed the using multitissue TMA with 55 different antibodies, two nuclear markers and H&E took about 36 h to run and resulted in 3,630 tissue protein expression readouts (approximately 2,000 cells per 0.6-mm diameter core; total of 7.26 × 10^6^ single-cell protein readouts).

#### Figure creation

Parts of Figures 1B, 2A and 3A were created using templates from Biorender (https://biorender.com). Parts of Figures 1, 4, 6, S12, S13, S14, S15, S16, S20 were created and corresponding statistical analyses were performed using GraphPad Prism^®^ 5.0 (GraphPad Software).

## QUANTIFICATION AND STATISTICAL ANALYSIS

### Computational image processing

Raw TIFF image files were processed using the CODEX Toolkit uploader (Goltsev et al., 2018). Briefly, this software computationally concatenates and drift-compensates the images using Hoechst nuclear stain as a reference, removes out-of-focus light using the Microvolution deconvolution algorithm (Microvolution), subtracts the background (using blank imaging cycles without fluorescent oligonucleotides), and creates hyperstacks of all fluorescence channels and imaging cycles of the imaged TMA spots. The following settings in the CODEX uploader were used for the TMA experiments: Microscope: Keyence BZ-X710. Deconvolution: Microvolution. Objective type: Air. Magnification (x): 20. Numerical aperture: 0.75. Lateral resolution (nm/pixel): 377.442. Z pitch (nm): 1500. Number of Z-slices: 17. Color mode: grayscale. Drift compensation channel index: 1. Drift compensation reference cycle: 1. Best focus channel: 1. Best focus cycle: 1. Number of cycles / Range: 1-23 (multi-tumor TMA), 1-24 (CRC TMA). Tiling mode: snake. Region size X and Y: both 1. Tile overlap X and Y: both 0. H&E staining: yes (no in the case of background subtraction). Focusing fragment: no. Background subtraction: yes (no if H&E staining was co-processed). Use blind deconvolution: yes. Use bleach-minimizing cropping: no. Processing only, export as TIFF.

After uploading, all spots of each TMA were stitched together into a single 10×7 spots file using the grid/collection stitching plugin (Preibisch et al., 2009) in ImageJ software (Fiji, version 2.0.0). Antibody stainings were visually assessed for each channel and cycle in each spot, and seven-color overlay figures for selected markers were generated.

Hyperstacks from the CRC TMA spots were segmented based on DRAQ5 nuclear stain, pixel intensities were quantified, and spatial fluorescence compensation was performed using the CODEX toolkit segmenter, with the following settings: Radius: 7. Max. cutoff: 1.0. Min. cutoff: 0.07. Relative cutoff: 0.2. Cell size cutoff factor: 0.4. Nuclear stain channel: 4. Nuclear stain cycle: 23. Membrane stain channel: 1. Membrane stain cycle: −1 (i.e., not used). Use membrane: false. Inner ring size: 1.0. Delaunay graph: false. Anisotropic region growth: false. Comma-separated value (CSV) and flow cytometry standard (FCS) files were generated from each TMA spot and used for further downstream analysis.

### Cleanup gating, unsupervised hierarchical clustering and cluster validation

All 140 background-subtracted FCS files from both CRC TMAs were imported into CellEngine (www.cellengine.com). Gates were tailored for each file individually in a blinded manner by two experts in flow and mass cytometry (C.M.S and D.R.M.). Nucleated cells were positively identified, and artifacts were removed by gating on Hoechst1/DRAQ5 doublepositive cells, followed by gating on focused cells in the Z plane (**Figure S5**). After cleanup gating, FCS files were re-exported and subsequently imported into VorteX clustering software, where they were subjected to unsupervised hierarchical X-shift clustering using an angular distance algorithm (Samusik et al., 2016). The following data import settings were applied: Numerical transformation: none. Noise threshold: no. Feature rescaling: none. Normalization: none. Merge all files into one dataset: yes. Clustering was based on all antibody markers except CD57, CD71, CD194 (CCR4), CDX2, Collagen IV, MMP9 and MMP12. The following settings were used for clustering: Distance measure: Angular distance. Clustering algorithm: X-shift (gradient assignment). Density estimate: N nearest neighbors (fast). Number of neighbors for density estimate (K): From 150 to 5, steps 30. Number of neighbors: determine automatically.

The optimal cluster number was determined using the elbow point validation tool and was at K=40, resulting in 143 clusters. Clusters and corresponding data were exported as a CSV file and were manually verified and assigned to cell types by overlaying the single cells from each cluster onto the stitched TMA images in ImageJ, based on the unique cluster identifiers and cellular X/Y position, using custom-made ImageJ scripts (available upon request). Clusters with similar morphological appearance in the tissue and similar marker expression profiles were merged, and artifacts were removed, resulting in 28 final clusters. Minimal spanning trees of the clusters were generated in VorteX, based on angular distance, and were exported for each marker (**Figure S11**).

### Manual gating of cell types and checkpoint-positive cell subsets

After cleanup gating, the frequencies of major immune cell types (**Figure S9**) and their expression of Ki-67 and checkpoint molecules (**Figure S20**) were manually gated in a blinded manner for each TMA spot in CellEngine, and statistics were exported for further analysis. For checkpoint-positive cell subsets, quantified checkpoint expression was compared to the raw image for each spot and gates were adjusted to best represent the raw image.

### Generation of Voronoi diagrams and contact matrices

FCS files were exported from VorteX and subjected to a custom-made Java algorithm to create Voronoi diagrams and cell-to-cell contact matrices (code available upon request).

### Computation of pairwise cell-cell contacts

Direct neighbors of each cell were determined by Delaunay triangulation as implemented in the *deldir* R package, using the default settings. From the original 28 cell clusters, two clusters were removed (undefined and tumor/immune cells), and the remaining clusters were merged into 14 clusters (**Figure S16B**). To represent associations of cells from various clusters, likelihood ratios or relative frequencies were calculated between the various clusters for each group of patients, according to the following formulas:

Likelihood ratios:

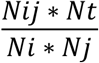

Relative frequencies:

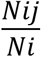

where

Nij is the number of edges between cells in cluster 1 and cluster 2

Nt is the total number of edges, Σ_*i*,j_ *Nij*

Ni is = Σ_*j*_ *Nij*

Nj is = Σ_*i*_ *Nij*

Log_2_ ratios of these metrics for CLR and DII patients were generated from the resulting matrices. A heatmap of the resulting values was plotted using the Heatmap function from the *ComplexHeatmap* R package after removing contacts between clusters where the number of unique adjacent cells was <100 in both patient groups.

### Neighborhood identification

For each of the 258,385 cells across these tissues across all spots and patient groups, a ‘window’ was captured consisting of the 10 nearest neighboring cells (including the center cell) as measured by Euclidean distance between X/Y coordinates. These windows were then clustered by the composition of their microenvironment with respect to the 29 cell types that had previously been identified by X-shift clustering and supervised annotation and merging (**Figure S12**). Of these, 28 cell clusters were assigned to biological cell types and one corresponded to imaging artifacts. This latter was included to identify poor quality regions of the image. Specifically, each window was converted to a vector of length 29 containing the frequency of each of the 29 cell types amongst the 10 neighbors. We then clustered these windows using Python’s *scikit-learn* implementation of MiniBatchKMeans with *k* = 10. Each cell was then allocated to the same CN as the window in which it was centered. To validate the CN assignment, these allocations were overlaid on the original tissue H&E-stained and fluorescent images. During this process, the CN cluster that contained the imaging artifacts (cellular cluster 29) was removed. The original code is available upon request.

### Non-negative Tucker tensor decomposition

The tensor of CN-cell type distributions for each patient, with dimensions patients × cell types × CNs, was produced by computing the frequency of each cell type in each CN in the non-follicular compartments (i.e., all CNs except CN-5). This tensor was split along the patient direction by patient group (CLR and DII). Non-negative Tucker decomposition as implemented in the *Tensorly* Python package was applied to each tensor (Kossaifi et al., 2019). The ranks in each dimension (2,6,6) were selected by a visual elbow point method assessing the decomposition loss (**Figure S19**). Several random-starts were utilized to ensure stability.

The cell type modules correspond to the factors in celltype space, and these are the values indicated in Figure 5. The CN modules correspond to the factors in CN space. The interactions comprising a tissue module correspond to each 6×6 slice of the 2×6×6 core tensor.

### Differential enrichment analyses

Linear models *Y_n,c_* = β_0_ + β_1_*X* + β_3_*Y*_c_ + ε were estimated, where *Y_c_* is the log overall frequency of cell type c, *X* is an indicator variable for patient group, *Y_n,c_* is the log frequency of cell type c in CN n, β_i_ are coefficients, and ε is mean zero Gaussian noise. A pseudocount of 1e^−3^ was added prior to taking logs. These were estimated using the *statsmodels* Python package (Seabold and Perktold, 2010). The coefficient estimates and p values were extracted and visualized.

### Classification of groups

Classification models were L1 regularized logistic regression models, fit using the *glmnet* R package (Friedman et al., 2010). Features were computed under the transformation x −> log(1e-3+ x) and z-normalized across the dataset prior to inclusion in any models.

Repeated hold out (RHO) was utilized to estimate prediction error, which utilized 10 training samples from each patient group. The L1 regularization parameter was chosen for each sampled train-test split by cross-validation on the sampled training set, and a model using this regularization parameter was fit on the sampled training set. The model was evaluated on the sampled test set, and ROC curves were estimated on the test set (of size 15). This was repeated 1000 times. The feature importance was computed as the z-score of the absolute value of the coefficient across resampling, as reported in (Laurin et al., 2016).

### CN activity alteration score

The CN activity alteration score was computed for each cell type individually. Specifically, for each cell type, the classification performance (AUC) was estimated using 200 RHO samples for two models. The first model included as features the overall frequency of that cell type. The second model included as features the overall frequency as well as the CN-specific cell type frequencies of that cell type in all CNs except CN-5. Each cell type had a different accuracy of classification with respect to the first model. To account for this, the CN activity alteration score was defined as the improvement in classification between the two models. Specifically, the negative log (base 10) of the p value of a one-sided *t*-test for a greater mean in the AUC between the second and first model was used. Since this entire procedure contained randomness, it was repeated 10 times to estimate the variability across the dataset of the CN activity alteration score.

### Survival analysis

We tested the log (1e-3 + frequency in CN-9) for each of PD-1^+^CD4^+^ and ICOS^+^CD4^+^ T cells individually, in Cox proportional hazards regression models, estimated using the *survival* R package (Therneau, 2015). The p value displayed was from the Cox regression model, and the Kaplan-Meier curve displayed was computed using the optimal split of the samples along the PD-1^+^CD4^+^ T cell frequency in CN-9 variable. The partial residual plot in **Figure 6J** was created using the *visreg* R package (Brehen and Burchett, 2013).

### Canonical correlation analysis

For each CN, the log CN-specific cell type frequency of each of ICOS^+^, Ki-67^+^, and PD-1^+^ and CD8^+^T cells as well as Ki-67^+^ Tregs was computed. For each pair of CNs, estimated canonical directions for the frequencies of these cells in each CN was estimated using the *scikit-learn* Python package (Pedregosa et al., 2011). For each pair of CNs, patients with no cells assigned to either CN were not included in the analysis. The correlation of the projections along these canonical directions was compared to a permutation distribution, corresponding to 5000 random permutations of the data. The permutation p value, i.e. the percentage of permutations whose estimated canonical correlation exceeded the observed one, was interpreted as the strength of communication.

## SUPPLEMENTAL INFORMATION

Supplemental information includes 22 figures, 6 tables and 12 scripts and can be found with this article online.

Table S1: Detailed patient characteristics

Table S2: Antibodies, clones, manufacturers

Table S3: CODEX oligonucleotides

Table S4: CODEX multi-cycle panels

Table S5: Multi-tumor TMA tissues

Table S6: Key resources

