## Supplementary material for "Coordinated cellular neighborhoods orchestrate antitumoral immunity at the colorectal cancer invasive front": Analysis code files: alteration_submission.ipynb

```
# %% [code]
library('glmnet')
library('pROC')
library('survival')
rename <- function(X){
    rownames(X) <- X$patients
    print(dim(X[,-1]))
    return(X[,-1])
}

# %% [code]
X_all <- rename(read.csv('X_all'))
X_cond_all <- rename(read.csv('X_cond_all'))
X_condnb_all <- rename(read.csv('X_condnb_all'))
X_chk_density <- rename(read.csv('X_chk_density'))

X_cts <- rename(read.csv('X_cts'))
X_chk_all <- rename(read.csv('X_chk_all'))
X_chk_overall_parent <- rename(read.csv('X_chk_overall_parent'))
X_chk_overall_pat <- rename(read.csv('X_chk_overall_pat'))
X_neighs <- rename(read.csv('X_neighs'))

X_cts_onlynb <- rename(read.csv('X_cts_onlynb'))
X_cts_onlynb_all <- rename(read.csv('X_cts_onlynb_all'))


group <- read.csv('group',header = FALSE)[,-1]
time <- read.csv('Y',header = FALSE)[,-1]
censor <- read.csv('censor',header = FALSE)[,-1]
status = censor#rep(1,length(time))#censor

# %% [code]
#the following function performs repeated hold-out
bs.cv.logit <- function(X,target,n = 500,num_train = 10,nfolds = 3){
    n_lam <- 0
    lams <- exp(-4*(c(1:50)/50))
    n_lam <- length(lams)
    
    print(n_lam)
    res <- list()
    T0 <- (1:length(target))[target==0]
    T1 <- (1:length(target))[target==1]
    coefmat <- matrix(0,nrow =n,ncol = 1+dim(X)[2])
    for (i in 1:n){
        #draw a permuted dataset; note the without replacement
        t0 <- sample(T0, length(T0), replace = F)
        t1 <- sample(T1, length(T1), replace = F)
        
        
        #split into train and test
        samp0 <- sample(1:length(t0),length(t0), replace = F)
        samp1 <- sample(1:length(t1),length(t1), replace = F)
        tr <- c(t0[samp0[1:num_train]],t1[samp1[1:num_train]])
        te <- c(t0[samp0[(1+num_train):length(samp0)]], t1[samp1[(1+num_train):length(samp1)]])
        
        #cv glmnet fit on train
        fit <- cv.glmnet(data.matrix(X[tr,]), target[tr], family = "binomial", maxit = 20000,nfolds = nfolds,lambda = lams)
        
        #predict on test
        y <- predict(fit, data.matrix(X[te,]), s= 'lambda.min')
        res[[i]] <- pROC::roc(response = target[te], predictor = c(y),auc = T)
        coefmat[i,1:(dim(X)[2]+1)]<- matrix(coef(fit, s = 'lambda.min'),nrow = 1)

    }
    
    return(list(res=res, coef = coefmat))
}
    

# %% [code]
#get the relative performance of the classification models for the main cell types
all_cell_names <- c('granulocytes', 'vasculature', 'CD4..T.cells.CD45RO.', 'tumor.cells', 'stroma', 'CD68.CD163..macrophages', 
'adipocytes', 'plasma.cells', 'CD8..T.cells', 'Tregs', 'CD4..T.cells', 'CD11c..DCs', 'B.cells', 'CD11b.CD68..macrophages', 
'smooth.muscle','nerves','lymphatics')


allr_overall = list()
allr_nb = list()
for (i in 1:10){
    print(i)
    allr_overall[[i]] <- list()
    allr_nb[[i]]<-list()
for (chk in all_cell_names){
    X_o <- data.frame(con = 1,o = scale(X_cts_onlynb[,colnames(X_cts_onlynb)[grepl(chk,colnames(X_cts_onlynb))]]))
    X_nb <- data.frame(scale(X_condnb_all[,colnames(X_condnb_all)[grepl(chk,colnames(X_condnb_all))]]))
    X_nb$o <- X_o$o
    allr_overall[[i]][[chk]] <- bs.cv.logit(X_o, group, n = 200)
    allr_nb[[i]][[chk]] <- bs.cv.logit(X_nb, group, n = 200)
}

}


# %% [code]
aucs_overall <- list()
aucs_nb <- list()
scores <- list()
sc_o <- list()
sc_nb <- list()
df <- NULL
for (j in 1:10){
    scores[[j]] <- list()
for (chk in all_cell_names){
    an <- NULL
    ao <- NULL
    for (i in 1:200){
        an <- c(an, allr_nb[[j]][[chk]]$res[[i]]$auc)
        ao <- c(ao, allr_overall[[j]][[chk]]$res[[i]]$auc)
    }
    aucs_overall[[chk]] <- ao
    aucs_nb[[chk]] <- an
    scores[[j]][[chk]] <- -log(t.test(an,ao,alternative = 'greater')$p.value,10)# mean(an)-mean(ao)
    }
    
}
js <- NULL
chks <-NULL
sc <- NULL


for (j in 1:10){
    for (chk in all_cell_names){
        js <- c(js, j)
        chks <- c(chks, chk)
        sc <- c(sc,scores[[j]][[chk]])
    }
}
df <- data.frame(it = js, chk = chks, score = sc)
options(repr.plot.width=12, repr.plot.height=6)
ggplot(df, aes(x = factor(chk), y = score)) + geom_jitter(width = 0.1,size = 0.5)+
theme_bw()+
theme(axis.text.x = element_text(angle = 90, hjust = 1))+
ylab('reorganization score')
ggsave('reorganization_score_all.pdf')

# %% [code]
#get the relative performance of the classification models for the functional subsets
chk.names = c( 'CD4.ICOS.','CD4.Ki67.','CD4.PD.1.','CD68.CD163.ICOS','CD68.CD163.Ki67.','CD68.CD163.PD.1.', 'CD8.ICOS.','CD8.Ki67.','CD8.PD.1.','Treg.ICOS.','Treg.Ki67.','Treg.PD.1.')


r_overall = list()
r_nb = list()
for (i in 1:10){
    print(i)
    r_overall[[i]] <- list()
    r_nb[[i]]<-list()
for (chk in chk.names){
    X_o <- data.frame(con = 1,o = scale(X_chk_overall_parent[,colnames(X_chk_overall_parent)[grepl(chk,colnames(X_chk_overall_parent))]]))
    X_nb <- data.frame(scale(X_chk_density[,colnames(X_chk_density)[grepl(chk,colnames(X_chk_density))]]))
    X_nb$o <- X_o$o
    r_overall[[i]][[chk]] <- bs.cv.logit(X_o, group, n = 200)
    r_nb[[i]][[chk]] <- bs.cv.logit(X_nb, group, n = 200)
}

}

# %% [code]
aucs_overall <- list()
aucs_nb <- list()
scores <- list()
sc_o <- list()
sc_nb <- list()
df <- NULL
for (j in 1:10){
    scores[[j]] <- list()
for (chk in chk.names){
    an <- NULL
    ao <- NULL
    for (i in 1:200){
        an <- c(an, r_nb[[j]][[chk]]$res[[i]]$auc)
        ao <- c(ao, r_overall[[j]][[chk]]$res[[i]]$auc)
    }
    aucs_overall[[chk]] <- ao
    aucs_nb[[chk]] <- an
    scores[[j]][[chk]] <- -log(t.test(an,ao,alternative = 'greater')$p.value,10)# mean(an)-mean(ao)
    }
    
}
js <- NULL
chks <-NULL
sc <- NULL
for (j in 1:10){
    for (chk in chk.names){
        js <- c(js, j)
        chks <- c(chks, chk)
        sc <- c(sc,scores[[j]][[chk]])
    }
}

# %% [code]
require(ggplot2)
library(repr)

# Change plot size to 4 x 3
options(repr.plot.width=4, repr.plot.height=3)

# %% [code]
df <- data.frame(it = js, chk = chks, score = sc)
goodchk <- c('Treg.PD.1.', 'Treg.Ki67.' ,'Treg.ICOS.' ,'CD8.PD.1.' ,'CD8.Ki67.', 'CD8.ICOS.', 'CD4.PD.1.', 'CD4.Ki67.' ,'CD4.ICOS.', 'CD68.CD163.PD.1.', 'CD68.CD163.Ki67.','CD68.CD163.ICOS' )
df <- df[df$chk%in%goodchk,]
df$chk <- factor(df$chk, levels = goodchk)
ggplot(df, aes(x = chk, y = score)) + geom_violin()+
theme_bw()+
theme(axis.text.x = element_text(angle = 90, hjust = 1))+
ylab('reorganization score')
ggsave('reorganization_score_violin.pdf')


# %% [code]

ggplot(df, aes(x = factor(chk), y = score)) + geom_boxplot()+
theme_bw()+
theme(axis.text.x = element_text(angle = 90, hjust = 1))+
ylab('reorganization score')
ggsave('reorganization_score_box.pdf')

# %% [code]

ggplot(df, aes(x = factor(chk), y = score)) + geom_jitter(size = 0.8,width = 0.2)+
theme_bw()+
theme(axis.text.x = element_text(angle = 90, hjust = 1))+
ylab('reorganization score')
ggsave('reorganization_score_jitter.pdf')
```
