## Supplementary material for "Coordinated cellular neighborhoods orchestrate antitumoral immunity at the colorectal cancer invasive front": Analysis code files: cca_submission.ipynb

```
# %% [code]
import numpy as np
import pandas as pd
from matplotlib import pyplot as plt
from scipy.stats import pearsonr,spearmanr
import itertools
from sklearn.cross_decomposition import CCA
import networkx as nx
import seaborn as sns
%pylab inline

# %% [code]
cells2 = pd.read_pickle('cells2_salil')
ct_frq = pd.read_pickle('cell_type_freqs_per_patient')
chk_frq = pd.read_pickle('checkpoint_to_parent_freqs_perneighborhood_perpatient')
neigh_frqs = pd.read_pickle('neigh_freqs_per_patient')
all_frqs = pd.read_pickle('allcell_type_counts_perneighborhood_perpatient')

# %% [code]
annotations = pd.read_csv('blocknumb_to_patient_no.csv')
survival = pd.read_csv('updated_survival.csv', header = 1)
annotations['B-Number'] = annotations['B-Number'].astype(str)
survival['Probenummer_Original'] = survival['Probenummer_Original'].astype(str)
# i checked that the indexes are the same (B numbers warped)
annotations['OS'] = survival['OS'].loc[annotations.index.values]
annotations['OS_censor'] = survival['OS_Censor'].loc[annotations.index.values]

annotations = annotations.set_index('Patient')
good_patients = annotations[~annotations['OS'].isna()].index.values

annotations['B-Number'].to_csv('BN')
group = annotations['Group'].loc[good_patients]

# %% [code]
#work out neighborhood counts
nbd_counts = cells2.groupby(['patients','neighborhood10']).size().unstack().fillna(0).loc[good_patients]
pat_counts = cells2.groupby(['patients']).size().fillna(0).loc[good_patients]


# %% [code]
# select which neighborhoods and functional subsets
nbs = [0,2,3,4,5,6,7,8,9]

chks = ['CD8+ICOS+','CD8+Ki67+','CD8+PD-1+','Treg-Ki67+']

# %% [code]
# make a df of neighborhood specific cell type densities
x = all_frqs.reset_index().iloc[:,1:]
df_list = []
for nb in nbs:
    arr = x.loc[x['neighborhood10']==nb,:].set_index('patients').loc[good_patients,chks].values
    arr /= nbd_counts[nb].values[:,None]
    df_list.append(pd.DataFrame(arr, index = good_patients, columns = [chk+'_'+str(nb) for chk in chks]))
    
X_density = pd.concat(df_list, axis = 1)

X_density = np.log(1e-3+X_density.loc[:, X_density.apply(np.std,axis = 0)>0])


# %% [code]
cca = CCA(n_components=1,max_iter = 1000)
def do_cca( n_perms,fun = 'pearson'):
    if fun == 'pearson':
        func = pearsonr
    if fun == 'spearman':
        func = spearmanr
    
    cols = [chk + '_' for chk in chks]
    stats = {}
    for gp in [1,2]:
        stats[gp] = {}
        for k,nb1 in enumerate(nbs):
            for l,nb2 in enumerate(nbs):
                if (k<l):
                    nb1_cols = [c +str(nb1) for c in cols]
                    nb2_cols = [c +str(nb2) for c in cols]
                    dat = X_density.loc[group==gp,nb1_cols+nb2_cols].dropna(axis = 1, how = 'all').dropna(axis = 0, how = 'any')
                    
                    a = dat.loc[:,[n for n in nb1_cols if n in dat.columns]].values
                    b = dat.loc[:,[n for n in nb2_cols if n in dat.columns]].values
#                     if (len(a)<2) or (a.shape[1]<1) or (b.shape[1]<1):
#                         print('continuing',nb1,nb2)
#                         continue

    
                    x,y = cca.fit_transform(a,b)
                    arr = np.zeros(n_perms)
                    #compute the canonical correlation
                    stats[gp][nb1,nb2] = (func(x,y)[0],arr)
                    for i in range(n_perms):
                        idx = np.arange(len(a))
                        np.random.shuffle(idx)
                        #compute over n_perms permutations
                        xt,yt = cca.fit_transform(a[idx],b)
                        arr[i] = func(xt,yt)[0]
        print(gp, 'done')


    return stats

# %% [code]
def draw_stats(stats_treg,cutoff = 0.1):
    g1 = nx.Graph()
    g2 = nx.Graph()
    for k,nb1 in enumerate(nbs):
        for l,nb2 in enumerate(nbs):
            if (k<l):
                p1 = np.mean(stats_treg[1][(nb1,nb2)][1]<stats_treg[1][(nb1,nb2)][0])
                p2 = np.mean(stats_treg[2][(nb1,nb2)][1]<stats_treg[2][(nb1,nb2)][0])
                if (1-p1)<cutoff:
                    g1.add_edge(nb1,nb2,weight = 1-p1)
                    print('gp1',nb1,nb2,p1,p2)
                if (1-p2)<cutoff:
                    g2.add_edge(nb1,nb2,weight = 1-p2)
                    print('gp2',nb1,nb2,p1,p2)

    nx.draw_networkx(g1) 
    plt.axis('off')
    plt.show()

    nx.draw_networkx(g2)
    plt.axis('off')
    plt.show()
    return g1,g2


# %% [code]
np.random.seed(145)
stats = do_cca( n_perms = 5000,fun = 'pearson')

# %% [code]
g1,g2 = draw_stats(stats)

# %% [code]

pal = sns.color_palette('bright',10)
dash = {True: '-', False: ':'}
pos=nx.drawing.nx_pydot.pydot_layout(g1,prog='neato')
for k,v in pos.items():
    x,y = v
    plt.scatter([x],[y],c = pal[k], s = 2000,zorder = 3)
    plt.text(x,y, k, fontsize = 10, zorder = 10,ha = 'center', va = 'center')


atrs = nx.get_edge_attributes(g1, 'weight')    
for e0,e1 in g1.edges():
    p = atrs[e0,e1]
    plt.plot([pos[e0][0],pos[e1][0]],[pos[e0][1],pos[e1][1]], c= 'black',alpha = 1-10*p,linewidth = 4)
    #plt.text(0.5*(pos[e0][0]+pos[e1][0]),0.5*(pos[e0][1]+pos[e1][1]), 'p = %.3f'%p)
    
plt.axis('off')

#plt.savefig('final_cca_gp1.pdf',transparent = True)
plt.show()

figsize(10,10)
pos=nx.drawing.nx_pydot.pydot_layout(g2,prog='neato')
for k,v in pos.items():
    x,y = v
    plt.scatter([x],[y],c = pal[k], s = 2000,zorder = 3)
    plt.text(x,y, k, fontsize = 10, zorder = 10,ha = 'center', va = 'center')
atrs = nx.get_edge_attributes(g2, 'weight')    
for e0,e1 in g2.edges():
    p = atrs[e0,e1]
    plt.plot([pos[e0][0],pos[e1][0]],[pos[e0][1],pos[e1][1]], c= 'black',alpha = 1-10*p,linewidth = 4)
    #plt.text(0.5*(pos[e0][0]+pos[e1][0]),0.5*(pos[e0][1]+pos[e1][1]), 'p = %.3f'%p)
plt.ylim(-100,350)
plt.axis('off')
#plt.savefig('final_cca_gp2.pdf',transparent = True)
    
plt.show()
                

# %% [code]
# to make graph with common node postiions  (in final paper, we use node positions of g1 however)
g_comb = g1.copy()
for source,dest in g2.edges:
    g_comb.add_edge(source,dest)
    
pos = nx.drawing.nx_pydot.graphviz_layout(g_comb,'neato')
f,ax = plt.subplots(ncols = 2,sharex=True,sharey=True,figsize = (10,5))
g1.add_node(9)
nx.draw_networkx(g1,pos = pos,ax = ax[0],node_color=[sns.color_palette('bright')[i] for i in g1.nodes])

g2.add_node(5)
nx.draw_networkx(g2,pos = pos,ax = ax[1],node_color=[sns.color_palette('bright')[i] for i in g2.nodes])


# %% [code]
```
