## Supplementary material for "Coordinated cellular neighborhoods orchestrate antitumoral immunity at the colorectal cancer invasive front": Analysis code files: voronoi.py

```
import numpy as np
import matplotlib.pyplot as plt
import seaborn as sns
# from shapely.ops import polygonize,unary_union
from shapely.geometry import MultiPoint, Point, Polygon
from scipy.spatial import Voronoi


        
def voronoi_finite_polygons_2d(vor, radius=None):
    """
    adapted from https://stackoverflow.com/questions/20515554/colorize-voronoi-diagram/20678647#20678647 3.18.2019
    
    
    Reconstruct infinite voronoi regions in a 2D diagram to finite
    regions.

    Parameters
    ----------
    vor : Voronoi
        Input diagram
    radius : float, optional
        Distance to 'points at infinity'.

    Returns
    -------
    regions : list of tuples
        Indices of vertices in each revised Voronoi regions.
    vertices : list of tuples
        Coordinates for revised Voronoi vertices. Same as coordinates
        of input vertices, with 'points at infinity' appended to the
        end.

    """

    if vor.points.shape[1] != 2:
        raise ValueError("Requires 2D input")

    new_regions = []
    new_vertices = vor.vertices.tolist()

    center = vor.points.mean(axis=0)
    if radius is None:
        radius = vor.points.ptp().max()

    # Construct a map containing all ridges for a given point
    all_ridges = {}
    for (p1, p2), (v1, v2) in zip(vor.ridge_points, vor.ridge_vertices):
        all_ridges.setdefault(p1, []).append((p2, v1, v2))
        all_ridges.setdefault(p2, []).append((p1, v1, v2))

    # Reconstruct infinite regions
    for p1, region in enumerate(vor.point_region):
        vertices = vor.regions[region]

        if all(v >= 0 for v in vertices):
            # finite region
            new_regions.append(vertices)
            continue

        # reconstruct a non-finite region
        ridges = all_ridges[p1]
        new_region = [v for v in vertices if v >= 0]

        for p2, v1, v2 in ridges:
            if v2 < 0:
                v1, v2 = v2, v1
            if v1 >= 0:
                # finite ridge: already in the region
                continue

            # Compute the missing endpoint of an infinite ridge

            t = vor.points[p2] - vor.points[p1] # tangent
            t /= np.linalg.norm(t)
            n = np.array([-t[1], t[0]])  # normal

            midpoint = vor.points[[p1, p2]].mean(axis=0)
            direction = np.sign(np.dot(midpoint - center, n)) * n
            far_point = vor.vertices[v2] + direction * radius

            new_region.append(len(new_vertices))
            new_vertices.append(far_point.tolist())

        # sort region counterclockwise
        vs = np.asarray([new_vertices[v] for v in new_region])
        c = vs.mean(axis=0)
        angles = np.arctan2(vs[:,1] - c[1], vs[:,0] - c[0])
        new_region = np.array(new_region)[np.argsort(angles)]

        # finish
        new_regions.append(new_region.tolist())

    return new_regions, np.asarray(new_vertices)

def plot_voronoi(points,colors,invert_y = True,edge_color = 'facecolor',line_width = .1,alpha = 1,size_max=np.inf):
    
# spot_samp = spot#.sample#(n=100,random_state = 0)
# points = spot_samp[['X:X','Y:Y']].values
# colors = [sns.color_palette('bright')[i] for i in spot_samp['neighborhood10']]

    if invert_y:
        points[:,1] = max(points[:,1])-points[:,1]
    vor = Voronoi(points)

    regions, vertices = voronoi_finite_polygons_2d(vor)

    pts = MultiPoint([Point(i) for i in points])
    mask = pts.convex_hull
    new_vertices = []
    if type(alpha)!=list:
        alpha = [alpha]*len(points)
    areas = []
    for i,(region,alph) in enumerate(zip(regions,alpha)):
        polygon = vertices[region]
        shape = list(polygon.shape)
        shape[0] += 1
        p = Polygon(np.append(polygon, polygon[0]).reshape(*shape)).intersection(mask)
        areas+=[p.area]
        if p.area <size_max:
            poly = np.array(list(zip(p.boundary.coords.xy[0][:-1], p.boundary.coords.xy[1][:-1])))
            new_vertices.append(poly)
            if edge_color == 'facecolor':
                plt.fill(*zip(*poly), alpha=alph,edgecolor=  colors[i],linewidth = line_width , facecolor = colors[i])
            else:
                plt.fill(*zip(*poly), alpha=alph,edgecolor=  edge_color,linewidth = line_width, facecolor = colors[i])
        # else:

        #     plt.scatter(np.mean(p.boundary.xy[0]),np.mean(p.boundary.xy[1]),c = colors[i])
    return areas
def draw_voronoi_scatter(spot,c,palette = sns.color_palette('bright'),X = 'X:X', Y = 'Y:Y',hue = 'neighborhood10',
        figsize = (5,5),
         voronoi_kwargs = {},
         scatter_kwargs = {}):
    '''
    plot voronoi of a region and overlay the location of specific cell types onto this
    
    spot:  cells that are used for voronoi diagram
    c:  cells that are plotted over voronoi
    palette:  color palette used for coloring neighborhoods
    X/Y:  column name used for X/Y locations
    hue:  column name used for neighborhood allocation
    figsize:  size of figure
    voronoi_kwargs:  arguments passed to plot_vornoi function
    scatter_kwargs:  arguments passed to plt.scatter()
    '''
    if len(c)>0:
        neigh_alpha = .3
    else:
        neigh_alpha = 1
        
    voronoi_kwargs = {**{'alpha':neigh_alpha},**voronoi_kwargs}
    scatter_kwargs = {**{'s':50,'alpha':1,'marker':'.'},**scatter_kwargs}
    
    plt.figure(figsize = figsize)
    colors  = [palette[i] for i in spot[hue]]
    a = plot_voronoi(spot[[X,Y]].values,
                 colors,#[{0:'white',1:'red',2:'purple'}[i] for i in spot['color']],
                 **voronoi_kwargs)
    
    if len(c)>0:
        colors  = [palette[i] for i in c[hue]]
        plt.scatter(x = c[X],y = (max(spot[Y])-c[Y].values),
                   c = colors,**scatter_kwargs
                   )
    plt.axis('off');
    return a
    
    
        
```
