## Supplementary material for "Coordinated cellular neighborhoods orchestrate antitumoral immunity at the colorectal cancer invasive front": Analysis code files: classification_submission.ipynb

```
# %% [code]
library('glmnet')
library('pROC')
library('survival')
library('ggplot2')


# %% [code]
rename <- function(X){
    rownames(X) <- X$patients
    print(dim(X[,-1]))
    return(X[,-1])
}

# %% [code]

X_chk_density <- rename(read.csv('X_chk_density'))
X_chk_overall_patient <- rename(read.csv('X_chk_overall_pat'))


group <- read.csv('group',header = FALSE)[,-1]

# %% [code]
#repeated hold out with nested cross-validation for selection of regularization parameter
bs.cv.logit <- function(X,target,n = 500){
    n_lam <- 0
    lams <- exp(-4*(c(1:50)/50))
    n_lam <- length(lams)
    
    print(n_lam)
    res <- list()
    T0 <- (1:length(target))[target==0]
    T1 <- (1:length(target))[target==1]
    coefmat <- matrix(0,nrow =n,ncol = 1+dim(X)[2])
    for (i in 1:n){
        #draw a permuted dataset
        t0 <- sample(T0, length(T0), replace = F)
        t1 <- sample(T1, length(T1), replace = F)
        
        
        #split into train and test
        samp0 <- sample(1:length(t0),length(t0), replace = F)
        samp1 <- sample(1:length(t1),length(t1), replace = F)
        tr <- c(t0[samp0[1:10]],t1[samp1[1:10]])
        te <- c(t0[samp0[11:length(samp0)]], t1[samp1[11:length(samp1)]])
        
        #cv glmnet fit on train
        fit <- cv.glmnet(data.matrix(X[tr,]), target[tr], family = "binomial", maxit = 20000,nfolds = 3,lambda = lams)
        
        #predict on test
        y <- predict(fit, data.matrix(X[te,]), s= 'lambda.min')
        res[[i]] <- pROC::roc(response = target[te], predictor = c(y),auc = T)
        coefmat[i,1:(dim(X)[2]+1)]<- matrix(coef(fit, s = 'lambda.min'),nrow = 1)

    }
    
    return(list(res=res, coef = coefmat))
}
    
get_coefs <- function(out,cols){
sd = apply(out$coef, 2, sd)
df <- data.frame(names = c("1", cols), zsc = colMeans(out$coef)/sd, mean = colMeans(out$coef))
return(df)
}


# %% [code]
set.seed(12345)
nsd_chks  <- colnames(X_chk_density)
of_chks  <- colnames(X_chk_overall_patient)
res.chk.overall.patient <- bs.cv.logit(scale(X_chk_overall_patient[,of_chks]), group, n = 1000)
res.chk.density <- bs.cv.logit(scale(X_chk_density[,nsd_chks]), group, n = 1000)

# %% [code]
df <- get_coefs(res.chk.density,cols = colnames(X_chk_density[,nsd_chks]))
df <- df[order(-abs(df$zsc))[1:20],]
df$names <- factor(df$names,levels = df$names)
ggplot(df, aes(x = names, y = abs(zsc)))+geom_col()+
theme_bw()+
ylab('coefficient importance')+
xlab('CN specific density')+
theme(axis.text.x = element_text(angle = 90, hjust = 1))
ggsave('feature_importance.pdf')

# %% [code]
get_roc_df <- function(out,name){
sens <- NULL
spec <- NULL
for (i in 1:1000){
    sens <- c(sens,out$res[[i]]$sensitivities)
    spec <- c(spec,out$res[[i]]$specificities)
}

df <- data.frame(se = sens, sp = spec)
df <- merge(aggregate(1-df$sp, by = list(df$se), function(x) return(sd(x)/sqrt(length(x)))),aggregate(1-df$sp, by = list(df$se), mean), by = 'Group.1')
colnames(df) <- c('TPR', 'SEM', 'FPR')
df$cond <- name
df$minSpec <- df$TPR- 3*df$SEM
df$maxSpec <- df$TPR + 3*df$SEM
if (max(df$TPR)<1){
df <- rbind(df, c(1.0, 0.0, 1.0,name, 1.0, 1.0))}
                      return(df)
}
                      
options(repr.plot.width = 5, repr.plot.height = 3, repr.plot.res = 300)
                      
df <- rbind(get_roc_df(res.chk.density,'checkpoint_density_in_nbd'),
get_roc_df(res.chk.overall.patient,'overall_checkpoints'))
ggplot(df, aes(fill = cond, x= FPR, y= TPR)) + geom_line(aes(col = cond))+
geom_ribbon(aes(ymin = minSpec, ymax = maxSpec),alpha = 0.2)+theme_bw()+
geom_abline(slope = 1, intercept = 0,linetype="dashed")+
xlim(0,1.2)+ylim(0,1.05)+
ggsave('revised_ROCs.pdf')

# %% [code]
```
