## Supplementary material for "Coordinated cellular neighborhoods orchestrate antitumoral immunity at the colorectal cancer invasive front": Analysis code files: differential_enrichment_submission.ipynb

```
# %% [code]
from tensorly.decomposition import non_negative_parafac,parafac,non_negative_tucker
from tensorly.regression import KruskalRegressor
import numpy as np
import pandas as pd
from matplotlib import pyplot as plt
from scipy.stats import ttest_ind
from sklearn.utils import resample
import itertools
from scipy.stats import spearmanr, ttest_ind
import seaborn as sns
%pylab inline

# %% [code]
cells2 = pd.read_pickle('cells2_salil')

# %% [code]
ct_frq = pd.read_pickle('cell_type_freqs_per_patient')
chk_frq = pd.read_pickle('checkpoint_to_parent_freqs_perneighborhood_perpatient')
neigh_frqs = pd.read_pickle('neigh_freqs_per_patient')
all_frqs = pd.read_pickle('allcell_type_counts_perneighborhood_perpatient')

annotations = pd.read_csv('blocknumb_to_patient_no.csv')
survival = pd.read_csv('updated_survival.csv', header = 1)
annotations['B-Number'] = annotations['B-Number'].astype(str)
survival['Probenummer_Original'] = survival['Probenummer_Original'].astype(str)
# i checked that the indexes are the same (B numbers warped)
annotations['OS'] = survival['OS'].loc[annotations.index.values]
annotations['OS_censor'] = survival['OS_Censor'].loc[annotations.index.values]
annotations['DFS'] = survival['DFS'].loc[annotations.index.values]
annotations['DFS_censor'] = survival['DFS_Censor'].loc[annotations.index.values]
annotations = annotations.set_index('Patient')
good_patients = annotations[~annotations['OS'].isna()].index.values
#good_patients = good_patients[annotations['DFS'].loc[good_patients]>0]
annotations['B-Number'].to_csv('BN')
group = (annotations['Group'].loc[good_patients]==2).astype(int) # 
(annotations['Group'].loc[good_patients]==2).astype(int).to_csv('group')

# %% [code]
# cells = ['CD4+ T cells CD45RO+',
#  'granulocytes',
#  'vasculature',
#  'plasma cells',
#  'stroma',
#  'tumor cells',
#  'CD68+CD163+ macrophages',
#  'CD8+ T cells',
#  'smooth muscle']
cells =  cells2['ClusterName'].unique()
parents = {'CD4+ICOS+':'CD4+',
 'CD4+Ki67+':'CD4+',
 'CD4+PD-1+':'CD4+',
 'CD68+CD163+ICOS+':'CD68+CD163+',
 'CD68+CD163+Ki67+':'CD68+CD163+',
 'CD68+CD163+PD-1+':'CD68+CD163+',
 'CD68+ICOS+':'CD68+',
 'CD68+Ki67+':'CD68+',
 'CD68+PD-1+':'CD68+',
 'CD8+ICOS+':'CD8+',
 'CD8+Ki67+':'CD8+',
 'CD8+PD-1+':'CD8+',
 'Treg-ICOS+':'CD25+FOXP3+',
 'Treg-Ki67+':'CD25+FOXP3+',
 'Treg-PD-1+':'CD25+FOXP3+'}

frq_chks = ['freq.CD4+ICOS+', 'freq.CD4+Ki67+',  'freq.CD4+PD-1+','freq.CD68+CD163+ICOS+', 
    'freq.CD68+CD163+Ki67+', 'freq.CD68+CD163+PD-1+','freq.CD68+ICOS+', 'freq.CD68+Ki67+','freq.CD68+PD-1+',
    'freq.CD8+ICOS+', 'freq.CD8+Ki67+',  'freq.CD8+PD-1+','freq.Treg-ICOS+', 'freq.Treg-Ki67+',
    'freq.Treg-PD-1+' ]
chks = [a.split('.')[1] for a in frq_chks ]


# %% [code]
nbd_counts = cells2.groupby(['patients','neighborhood10']).size().unstack().fillna(0).loc[good_patients]
pat_counts = cells2.groupby(['patients']).size().fillna(0).loc[good_patients]


# %% [code]
def normalize(X):
    arr = np.array(X.fillna(0).values)
    return pd.DataFrame(np.log2(1e-5 + arr/arr.sum(axis =1, keepdims = True)), index = X.index.values, columns = X.columns).fillna(0)


# %% [code]
# data prep
nbs = [0,2,3,4,6,7,8,9]
X_cts = normalize(ct_frq.reset_index().iloc[:,1:].set_index('patients').loc[good_patients,cells])

# X_neighs = add_constant(normalize(neigh_frqs.reset_index().iloc[:,1:].set_index('patients').loc[good_patients,nbs]))

x = all_frqs.reset_index().iloc[:,1:]
X_all = normalize(pd.concat([x.loc[x['neighborhood10']==nb,:].set_index('patients').loc[good_patients,cells].rename({col: col+'_'+str(nb) for col in cells},axis = 1) for nb in nbs], axis = 1))

# X_all.to_csv('X_all')
# X_cts.to_csv('X_cts')
# X_neighs.to_csv('X_neighs')

Y =  np.maximum(annotations['OS'].loc[good_patients],0.01)
censor = annotations['OS_censor'].loc[good_patients]

censor.to_csv('censor')
# Y.to_csv('Y')


x = all_frqs.reset_index().iloc[:,1:]
norm = (pd.concat([x.loc[x['neighborhood10']==nb,:].set_index('patients').loc[good_patients,cells].rename({col: col+'_'+str(nb) for col in cells},axis = 1) for nb in nbs], axis = 1))
ctf =all_frqs.reset_index().iloc[:,1:]
ctf = ctf[ctf['neighborhood10'].isin(nbs)].groupby('patients').apply(lambda x:x.sum(axis=0)).iloc[:,2:].loc[good_patients]
X_cond_all =np.vstack([(1+norm[col].values)/(1+ctf[col.split('_')[0]].values) for col in norm.columns]).T
X_cond_all = np.log(1e-3+pd.DataFrame(X_cond_all, columns = norm.columns, index = norm.index.values).fillna(0))
# X_cond_all.to_csv('X_cond_all')


nbs = [0,2,3,4,5,6,7,8,9]
norm = (pd.concat([x.loc[x['neighborhood10']==nb,:].set_index('patients').loc[good_patients,cells].rename({col: col+'_'+str(nb) for col in cells},axis = 1) for nb in nbs], axis = 1))
X_condnb_all =np.vstack([(1+norm[col].values)/(1+nbd_counts[int(col.split('_')[1])].values) for col in norm.columns]).T
X_condnb_all = np.log(1e-3+pd.DataFrame(X_condnb_all, columns = norm.columns, index = norm.index.values).fillna(0))
X_condnb_all = X_condnb_all.loc[:,X_condnb_all.apply(np.std, axis =0)>0]
# X_condnb_all.to_csv('X_condnb_all')


# %% [code]
# data prep
x = chk_frq[frq_chks].reset_index().iloc[:,1:].fillna(0)
X_chk_all = np.log(1e-3+pd.concat([x.loc[x['neighborhood10']==nb,:].set_index('patients').loc[good_patients,frq_chks].rename({col: col+'_'+str(nb) for col in frq_chks},axis = 1) for nb in nbs], axis = 1).fillna(0))
X_chk_all = X_chk_all.loc[:,X_chk_all.apply(np.std, axis =0)>0]
X_chk_all.to_csv('X_chk_all')


x = all_frqs.reset_index().iloc[:,1:]
totals = x.groupby('patients').apply(lambda x: x.sum(axis = 0)).loc[good_patients]
X_chk_overall =np.log(1e-3+ pd.DataFrame(np.stack([totals[chk].values/totals[parents[chk]].values for chk in chks]).T, columns = frq_chks, index = good_patients))
X_chk_overall.to_csv('X_chk_overall_parent')

x = all_frqs.reset_index().iloc[:,1:]
totals = x.groupby('patients').apply(lambda x: x.sum(axis = 0)).loc[good_patients]
X_chk_overall =np.log(1e-3+ pd.DataFrame(np.stack([totals[chk].values/pat_counts.values for chk in chks]).T, columns = frq_chks, index = good_patients))
X_chk_overall.to_csv('X_chk_overall_pat')


df_list = []
for nb in nbs:
    arr = x.loc[x['neighborhood10']==nb,:].set_index('patients').loc[good_patients,chks].values
    arr /= nbd_counts[nb].values[:,None]
    df_list.append(pd.DataFrame(arr, index = good_patients, columns = [chk+'_'+str(nb) for chk in chks]).fillna(0))
    
X_density = pd.concat(df_list, axis = 1)

X_density = np.log(1e-3+X_density.loc[:, X_density.apply(np.std,axis = 0)>0])
X_density.to_csv('X_chk_density')


# %% [code]
import statsmodels.api as sm
from statsmodels.stats.multitest import multipletests as mpt

# %% [code]
#differential enrichment for all cell subsets
changes = {}
nbs =[0, 2, 3, 4, 6, 7, 8, 9]
for col in X_cts.columns:
    for nb in nbs:
        #build a design matrix with a constant, group 0 or 1 and the overall frequencies
        X = pd.concat([X_cts[col], group.astype('int'),pd.Series(np.ones(len(group)), index = group.index.values)],axis = 1).values
        if col+'_%d'%nb in X_condnb_all.columns:
            Y = X_condnb_all[col+'_%d'%nb].values
            X = X[~pd.isna(Y)]
            Y = Y[~pd.isna(Y)]
            results = sm.OLS(Y,X).fit()
            changes[(col,nb)] = (results.pvalues[1], results.params[1])
        

dat = (pd.DataFrame(changes).loc[1].unstack())
dat = pd.DataFrame(np.nan_to_num(dat.values),index = dat.index, columns = dat.columns).T.sort_index(ascending=True).loc[:,X_cts.columns]
pvals = (pd.DataFrame(changes).loc[0].unstack()).T.sort_index(ascending=True).loc[:,X_cts.columns]

f, ax = plt.subplots(figsize = (20,10))
g = sns.heatmap(dat,cmap = 'bwr', vmin = -1, vmax = 1,cbar=False,ax = ax)
for a,b in zip(*np.where (pvals<0.05)):
    plt.text(b+.5,a+.55,'*',fontsize = 20,ha = 'center',va = 'center')
plt.tight_layout()
plt.savefig('all_cells_density_heatmap_uncut.pdf',transparent = True)

# %% [code]
frq_chks = ['freq.Treg-PD-1+','freq.Treg-Ki67+','freq.Treg-ICOS+','freq.CD8+PD-1+','freq.CD8+Ki67+','freq.CD8+ICOS+','freq.CD4+PD-1+','freq.CD4+Ki67+','freq.CD4+ICOS+','freq.CD68+CD163+PD-1+','freq.CD68+CD163+Ki67+','freq.CD68+CD163+ICOS+']
chks = [a.split('.')[1] for a in frq_chks ]
nbs = [0,2,3,4,6,7,8,9]
x = all_frqs.reset_index().iloc[:,1:]
x = x.loc[x['neighborhood10'].isin(nbs),:]
totals = x.groupby('patients').apply(lambda x: x.sum(axis = 0)).loc[good_patients]
X_chk_overall =np.log(1e-3+ pd.DataFrame(np.stack([totals[chk].values/pat_counts.values for chk in chks]).T, columns = frq_chks, index = good_patients))

df_list = []
for nb in nbs:
    arr = x.loc[x['neighborhood10']==nb,:].set_index('patients').loc[good_patients,chks].values
    arr /= nbd_counts[nb].values[:,None]
    df_list.append(pd.DataFrame(arr, index = good_patients, columns = [chk+'_'+str(nb) for chk in chks]))
    
X_density = pd.concat(df_list, axis = 1)

X_density = np.log(1e-3+X_density.loc[:, X_density.apply(np.std,axis = 0)>0])

changes = {}
for col in X_chk_overall.columns:
    for nb in nbs:
        X = pd.concat([X_chk_overall[col], group.astype('int'),pd.Series(np.ones(len(group)), index = group.index.values)],axis = 1).values
        if col.split('.')[1]+'_%d'%nb in X_density.columns:
            Y = X_density[col.split('.')[1]+'_%d'%nb].values
            X = X[~pd.isna(Y)]
            Y = Y[~pd.isna(Y)]
            results = sm.OLS(Y,X).fit()
            changes[(col,nb)] = (results.pvalues[1], results.params[1])
        

dat = (pd.DataFrame(changes).loc[1].unstack())
dat = pd.DataFrame(np.nan_to_num(dat.values),index = dat.index, columns = dat.columns).T.sort_index(ascending=True).loc[:,frq_chks]
pvals = (pd.DataFrame(changes).loc[0].unstack()).T.sort_index(ascending=True).loc[:,frq_chks]


ax = sns.heatmap(dat,cmap = 'bwr', vmin = -1, vmax = 1,cbar=False)

for a,b in zip(*np.where (pvals<0.05)):
    plt.text(b+.5,a+.55,'*',fontsize = 20,ha = 'center',va = 'center')
#plt.savefig('revised_heatmap.pdf')

# %% [code]


col = 'freq.CD8+Ki67+'
nb =0
X = pd.concat([X_chk_overall[col], group.astype('int'),pd.Series(np.ones(len(group)), index = group.index.values)],axis = 1).values
if col.split('.')[1]+'_%d'%nb in X_density.columns:
    Y = X_density[col.split('.')[1]+'_%d'%nb].values
    X = X[~pd.isna(Y)]
    Y = Y[~pd.isna(Y)]
    results = sm.OLS(Y,X).fit()

df0 = pd.DataFrame({'adj_den_cd8ki670':Y - results.params[0]*X_chk_overall[col],'group': [i+1 for i in group.values]})

col = 'freq.Treg-Ki67+'
nb =4
X = pd.concat([X_chk_overall[col], group.astype('int'),pd.Series(np.ones(len(group)), index = group.index.values)],axis = 1).values
if col.split('.')[1]+'_%d'%nb in X_density.columns:
    Y = X_density[col.split('.')[1]+'_%d'%nb].values
    X = X[~pd.isna(Y)]
    Y = Y[~pd.isna(Y)]
    results = sm.OLS(Y,X).fit()

df1 = pd.DataFrame({'adj_den_tregki674':Y - results.params[0]*X_chk_overall[col],'group': [i+1 for i in group.values]})
pd.concat([df0,df1],axis = 1)
#df.to_csv('adjusted_frequencies.csv')


sns.catplot(y = 'adj_den_cd8ki670',x = 'group', data =   df0)
sns.catplot(y = 'adj_den_tregki674',x = 'group', data =   df1)


# %% [code]
df

# %% [code]
```
