## Supplementary material for "Coordinated cellular neighborhoods orchestrate antitumoral immunity at the colorectal cancer invasive front": Analysis code files: fig6DE_submission.ipynb

```
# %% [code]
import pandas as pd
import numpy as np
import matplotlib.pyplot as plt
import seaborn as sns
from scipy.stats import ttest_ind
%matplotlib inline


# %% [code]
cells2 = pd.read_pickle('cells2_salil' )

# %% [code]
cells2.head()

# %% [code]
cells2['CD4notTreg'] = np.bitwise_and(cells2['CD4+']==1,cells2['CD25+FOXP3+']==0)

cells2['CD4notTregchkpt'] = np.bitwise_and(cells2['CD4notTreg'],cells2[['CD4+ICOS+','CD4+Ki67+','CD4+PD-1+']].sum(1)>0)
cells2['CD4notTregchklo'] = np.bitwise_and(cells2['CD4notTreg'],cells2[['CD4+ICOS+','CD4+Ki67+','CD4+PD-1+']].sum(1)==0)

cells2['Tregchkpt'] = cells2[['Treg-ICOS+','Treg-Ki67+','Treg-PD-1+']].sum(1)>0
cells2['Tregchklo'] = np.bitwise_and(cells2['CD25+FOXP3+']==1,cells2[['Treg-ICOS+','Treg-Ki67+','Treg-PD-1+']].sum(1)==0)

cells2['CD8chkpt'] = cells2[['CD8+ICOS+','CD8+Ki67+','CD8+PD-1+']].sum(1)>0
cells2['CD8+chklo'] = np.bitwise_and(cells2['CD8+']==1,cells2[['CD8+ICOS+','CD8+Ki67+','CD8+PD-1+']].sum(1)==0)

# %% [code]

chkpt3 = ['CD4notTregchkpt','CD8chkpt','Tregchkpt','CD4notTregchklo','CD8+chklo','Tregchklo']
densities = cells2[cells2['neighborhood10']!=1].groupby(['groups','patients','neighborhood10']).apply(lambda x:x[chkpt3].sum(0))
Dens = densities.dropna((0,1),how = 'all')


# %% [code]
Dens.head()

# %% [code]
f,ax = plt.subplots()
k = Dens.sum(level = 'neighborhood10').transform(lambda x: x/x.sum(),axis = 1)
k.loc[[0,2,3,4,5,6,7,8,9],:].iloc[:,[1,0,2]].plot(kind = 'bar',ax = ax,stacked = True,alpha = 1,
                                             color = ['#ED403C','#70BD46','#21409A'])

k.loc[[0,2,3,4,5,6,7,8,9],:].iloc[:,[4,3,5]].plot(kind = 'bar',stacked = True,alpha = .11,bottom =k.loc[[0,2,3,4,5,6,7,8,9],:].iloc[:,:3].sum(1),ax = ax,
                                             color = ['#ED403C','#70BD46','#21409A'])
plt.legend(loc='center left', bbox_to_anchor=(1.0, 0.5),)
# plt.ylim(0,.6)

# %% [code]
colss = [['CD4+ICOS+', 'CD4+Ki67+', 'CD4+PD-1+'],
 ['CD8+ICOS+', 'CD8+Ki67+', 'CD8+PD-1+'],
 ['Treg-ICOS+', 'Treg-Ki67+', 'Treg-PD-1+']]

# %% [code]
d = cells2.groupby(['groups','patients','neighborhood10']).apply(lambda x: x[[a for b in colss for a in b]].sum(0))

s

# %% [code]
d.head()

# %% [code]
def print_pvals(l):
    pvals = {}
    for i in [0,1,2]:
        chkpval = {}
        for n1 in [0,4,6,2]:
            for n2 in [0,4,6,2]:
                key = [n1,n2]
                key.sort()
                if tuple(key) not in chkpval:
                    i1 = l.loc[ix[:,:,n1],:].iloc[:,i]
                    i2 = l.loc[ix[:,:,n2],:].iloc[:,i]
                    pval = ttest_ind(i1.values,i2.values)[1]
                    chkpval[tuple(key)] = pval
        pvals[i] = chkpval
    return (pvals)

# %% [code]
ix = pd.IndexSlice
for i in [2,0,1]:
    l = d.loc[ix[:,:,[0,4,6,2]],colss[i]].transform(lambda x: x/x.sum(),1).dropna(how = 'all')
    print (print_pvals(l))
    lm = l.reset_index().melt(id_vars = ['groups','patients','neighborhood10'])
    sns.catplot(data = lm, hue_order = [0,4,6,2],hue = 'neighborhood10',
                dodge = True,x =  'variable',y = 'value' ,
#                 errwidth = 1,capsize = .05,saturation = 1,
                kind = 'violin',cut = 0,aspect = 2.5,linewidth = .5,
                palette = {i:sns.color_palette('bright')[i] for i in [0,4,6,2]}
                
            )
#     plt.savefig('raw_figs/'+'fig6chkviolin'+colss[i][0][:4]+'.pdf')
    plt.show()


# %% [code]
```
