## Supplementary material for "Coordinated cellular neighborhoods orchestrate antitumoral immunity at the colorectal cancer invasive front": Analysis code files: neigh_mixing script.py

```
'''
Graham Barlow, 6.27.2019

Code for computing the amount of shared surface area between two neighborhoods

requirements:

'tissues': saved in 'neighborhoodtemplate' python notebook.  List of length equal to the number of regions.  Each item is array shape (number of cells, number of nearest neighbors)
with the value being the index of the cell in cells2 (tissues = [get_windows(job,n_neighbors) for job in tissue_chunks])
IMPORTANT:  dataframe must be in same order as started with (and 'tissses' index refers to) or indexing would be incorrect

'cells2' (alias: cells).  Main dataframe output by neighborhoodtemplate with x,y,clusterID, and neighborhood allocation

'neigh1':  neighborhood name of 1st neighborhood
'neigh2':  neighborhood name of 2nd neighborhood


'''
neigh1 = 4
neigh2 = 0

#change column labels
neighborhood_name = 'neighborhood10'
group_name = 'groups'
patient_name = 'patients'


index = np.concatenate([tissues[i][:,0] for i in range(len(tissues))]) #index of original cell
neigh = np.concatenate([tissues[i][:,1] for i in range(len(tissues))]) #index of nearest neighbor of that original cell

nn = cells2.loc[neigh,neighborhood_name]
cells2['nearest_neighborhood'] = -1

#assign the neighborhood of the nearest cell to the original cell
cells2.loc[index,'nearest_neighborhood'] = nn.values
cells2['nearest_neighborhood'] = cells2['nearest_neighborhood'].astype('category')

#compute for each patient, in each group and neighborhood, the number of cells in that neighborhoood
counts = cells2.groupby([group_name,patient_name,neighborhood_name]).apply(lambda x: len(x)).unstack()

#compute for each patient, in each group and neighborhood:  the count of how many of the cells in that neighborhood are next to a cell in the other neighborhood
neighs = cells2.groupby([group_name,patient_name,neighborhood_name]).apply(lambda x: pd.concat([x['nearest_neighborhood'].value_counts(sort = False))


#index the values for the neighborhoods of interest
ix = pd.IndexSlice
inters = pd.concat([neighs.loc[ix[:,:,[neigh1]],[neigh2]],neighs.loc[ix[:,:,[neigh2]],[neigh1]]],1).mean(level = [group_name,patient_name]).mean(1)
wholes = neighs.sum(1).loc[ix[:,:,[neigh2,neigh1]]].unstack().sum(1)

combo = pd.concat([inters,wholes],1).dropna((0))
combo['ratio'] = combo[0]/combo[1]
```
