## Supplementary material for "Coordinated cellular neighborhoods orchestrate antitumoral immunity at the colorectal cancer invasive front": Analysis code files: neighborhood_template_submission.ipynb

```
# %% [code]
import pandas as pd
import numpy as np
from sklearn.neighbors import NearestNeighbors
import time
import sys
import matplotlib.pyplot as plt
from sklearn.cluster import MiniBatchKMeans
import seaborn as sns


def get_windows(job,n_neighbors):
    '''
    For each region and each individual cell in dataset, return the indices of the nearest neighbors.
    
    'job:  meta data containing the start time,index of region, region name, indices of region in original dataframe
    n_neighbors:  the number of neighbors to find for each cell
    '''
    start_time,idx,tissue_name,indices = job
    job_start = time.time()
    
    print ("Starting:", str(idx+1)+'/'+str(len(exps)),': ' + exps[idx])

    tissue = tissue_group.get_group(tissue_name)
    to_fit = tissue.loc[indices][[X,Y]].values

#     fit = NearestNeighbors(n_neighbors=n_neighbors+1).fit(tissue[[X,Y]].values)
    fit = NearestNeighbors(n_neighbors=n_neighbors).fit(tissue[[X,Y]].values)
    m = fit.kneighbors(to_fit)
#     m = m[0][:,1:], m[1][:,1:]
    m = m[0], m[1]
    

    #sort_neighbors
    args = m[0].argsort(axis = 1)
    add = np.arange(m[1].shape[0])*m[1].shape[1]
    sorted_indices = m[1].flatten()[args+add[:,None]]

    neighbors = tissue.index.values[sorted_indices]
   
    end_time = time.time()
   
    print ("Finishing:", str(idx+1)+"/"+str(len(exps)),": "+ exps[idx],end_time-job_start,end_time-start_time)
    return neighbors.astype(np.int32)


# %% [markdown]
### ------------------------

# %% [markdown]
### Fill In:
    -ks = individual window sizes to collect.  
        -This accelerates the process of trying different window sizes by collecting them all in one step.
    -path_to_data:  file path for csv file with x,y cluster information
    -X:  column to use for X location
    -Y:  column to use for Y location
    -reg:  column to use for region (should be different for every tissue)
    -file_type('csv','pickle'):  file type
    -cluster_col : column name of cell type cluster to be used for neighborhoods
    -cellhierpath:  path to cellhier libary (should only have to set once per computer
    -keep_cols:  columns to transfer from original dataframe to dataframe with windows and neighborhoods

# %% [code]
ks = [5,10,20] # k=5 means it collects 5 nearest neighbors for each center cell
path_to_data = '/media/TitanRAID/GRAHAM/Christian for Graham/Colorectal Cancer/CRC_clusters_final_with_clusternames.csv'
X = 'X:X'
Y = 'Y:Y'
reg = 'File Name'
file_type = 'csv'

cluster_col = 'ClusterName'
keep_cols = [X,Y,reg,cluster_col]
save_path = ''


# %% [markdown]
### ------------------------

# %% [code]
#read in data and do some quick data rearrangement
n_neighbors = max(ks)
assert (file_type=='csv' or file_type =='pickle') #


if file_type == 'pickle':
    cells = pd.read_pickle(path_to_data)
if file_type == 'csv':
    cells = pd.read_csv(path_to_data)

cells = pd.concat([cells,pd.get_dummies(cells[cluster_col])],1)
sum_cols = cells[cluster_col].unique()
values = cells[sum_cols].values

# %% [code]
#find windows for each cell in each tissue region
tissue_group = cells[[X,Y,reg]].groupby(reg)
exps = list(cells[reg].unique())
tissue_chunks = [(time.time(),exps.index(t),t,a) for t,indices in tissue_group.groups.items() for a in np.array_split(indices,1)] 
tissues = [get_windows(job,n_neighbors) for job in tissue_chunks]


# %% [code]
#for each cell and its nearest neighbors, reshape and count the number of each cell type in those neighbors.
out_dict = {}
for k in ks:
    for neighbors,job in zip(tissues,tissue_chunks):

        chunk = np.arange(len(neighbors))#indices
        tissue_name = job[2]
        indices = job[3]
        window = values[neighbors[chunk,:k].flatten()].reshape(len(chunk),k,len(sum_cols)).sum(axis = 1)
        out_dict[(tissue_name,k)] = (window.astype(np.float16),indices)

#concatenate the summed windows and combine into one dataframe for each window size tested.
windows = {}
for k in ks:
   
    window = pd.concat([pd.DataFrame(out_dict[(exp,k)][0],index = out_dict[(exp,k)][1].astype(int),columns = sum_cols) for exp in exps],0)
    window = window.loc[cells.index.values]
    window = pd.concat([cells[keep_cols],window],1)
    windows[k] = window


# %% [markdown]
### ------------------------

# %% [markdown]
### Fill In:
    -k:  window_size (needs to be in ks (saved above))
    -n_neighborhoods:  number of neighborhoods to find

# %% [code]
# 1 Follicular (Crohn's Like)
# 2 Diffuse

# %% [code]
k = 10
n_neighborhoods = 10
neighborhood_name = "neighborhood"+str(k)
k_centroids = {}

# %% [code]
windows2 = windows[10]
# windows2[cluster_col] = cells[cluster_col]

km = MiniBatchKMeans(n_clusters = n_neighborhoods,random_state=0)

labelskm = km.fit_predict(windows2[sum_cols].values)
k_centroids[k] = km.cluster_centers_
cells['neighborhood10'] = labelskm
cells[neighborhood_name] = cells[neighborhood_name].astype('category')
#['reg064_A','reg066_A','reg018_B','reg023_A']


# %% [code]
cell_order = ['tumor cells', 'CD11c+ DCs', 'tumor cells / immune cells',
       'smooth muscle', 'lymphatics', 'adipocytes', 'undefined',
       'CD4+ T cells CD45RO+', 'CD8+ T cells', 'CD68+CD163+ macrophages',
       'plasma cells', 'Tregs', 'immune cells / vasculature', 'stroma',
       'CD68+ macrophages GzmB+', 'vasculature', 'nerves',
       'CD11b+CD68+ macrophages', 'granulocytes', 'CD68+ macrophages',
       'NK cells', 'CD11b+ monocytes', 'immune cells',
       'CD4+ T cells GATA3+', 'CD163+ macrophages', 'CD3+ T cells',
       'CD4+ T cells', 'B cells']

# %% [code]
# this plot shows the types of cells (ClusterIDs) in the different niches (0-7)
k_to_plot = 10
niche_clusters = (k_centroids[k_to_plot])
tissue_avgs = values.mean(axis = 0)
fc = np.log2(((niche_clusters+tissue_avgs)/(niche_clusters+tissue_avgs).sum(axis = 1, keepdims = True))/tissue_avgs)
fc = pd.DataFrame(fc,columns = sum_cols)
s=sns.clustermap(fc.loc[[0,2,3,4,5,6,7,8,9],cell_order], vmin =-3,vmax = 3,cmap = 'bwr',row_cluster = False)
# s.savefig("raw_figs/celltypes_perniche_10.pdf")


# %% [code]
cells['neighborhood10'] = cells['neighborhood10'].astype('category')
sns.lmplot(data = cells[cells['groups']==1],x = 'X:X',y='Y:Y',hue = 'neighborhood10',palette = 'bright',height = 8,col = reg,col_wrap = 10,fit_reg = False)


# %% [code]
cells['neighborhood10'] = cells['neighborhood10'].astype('category')
sns.lmplot(data = cells[cells['groups']==2],x = 'X:X',y='Y:Y',hue = 'neighborhood10',palette = 'bright',height = 8,col = reg,col_wrap = 10,fit_reg = False)


# %% [markdown]
###### -------------------------------

# %% [code]


# %% [code]
#plot for each group and each patient the percent of total cells allocated to each neighborhood
fc = cells.groupby(('patients','groups')).apply(lambda x: x['neighborhood10'].value_counts(sort = False,normalize = True))

fc.columns = range(10)
melt = pd.melt(fc.reset_index(),id_vars = ['patients','groups'])
melt = melt.rename(columns = {'variable':'neighborhood','value':'frequency of neighborhood'})
f,ax = plt.subplots(figsize = (10,5))
sns.stripplot(data = melt, hue = 'groups',dodge = True,alpha = .2,x ='neighborhood', y ='frequency of neighborhood')
sns.pointplot(data = melt, scatter_kws  = {'marker': 'd'},hue = 'groups',dodge = .5,join = False,x ='neighborhood', y ='frequency of neighborhood')

handles, labels = ax.get_legend_handles_labels()
ax.legend(handles[:2], labels[:2], title="Groups",
          handletextpad=0, columnspacing=1,
          loc="upper left", ncol=3, frameon=True)

# %% [code]
#t-test to evaluate if any neighborhood is enriched in one group
from scipy.stats import ttest_ind
for i in range(10):
    n2 = melt[melt['neighborhood']==i]
    print (i,'    ',ttest_ind(n2[n2['groups']==1]['frequency of neighborhood'],n2[n2['groups']==2]['frequency of neighborhood']))

# %% [code]
#same as above except neighborhood 5 is removed from analysis.
fc = cells[cells['neighborhood10']!=5].groupby(('patients','groups')).apply(lambda x: x['neighborhood10'].value_counts(sort = False,normalize = True))

fc.columns = range(10)
melt = pd.melt(fc.reset_index(),id_vars = ['patients','groups'])
melt = melt.rename(columns = {'variable':'neighborhood','value':'frequency of neighborhood'})

f,ax = plt.subplots(figsize = (10,5))
sns.stripplot(data = melt, hue = 'groups',dodge = True,alpha = .2,x ='neighborhood', y ='frequency of neighborhood')
sns.pointplot(data = melt, scatter_kws  = {'marker': 'd'},hue = 'groups',dodge = .5,join = False,x ='neighborhood', y ='frequency of neighborhood')

handles, labels = ax.get_legend_handles_labels()
ax.legend(handles[:2], labels[:2], title="Groups",
          handletextpad=0, columnspacing=1,
          loc="upper left", ncol=3, frameon=True)

# %% [code]

for i in range(10):
    n2 = melt[melt['neighborhood']==i]
#n2 = n2[n2['Frequency']>.015]
    print (i,'    ',ttest_ind(n2[n2['groups']==1]['frequency of neighborhood'],n2[n2['groups']==2]['frequency of neighborhood']))

# %% [code]
```
