## Supplementary material for "Coordinated cellular neighborhoods orchestrate antitumoral immunity at the colorectal cancer invasive front": Analysis code files: pca_submission.ipynb

```
# %% [code]
import pandas as pd
import numpy as np
import matplotlib.pyplot as plt
import seaborn as sns
%matplotlib inline


# %% [code]
cells2 = pd.read_pickle('cells2_salil' )

# %% [code]
from sklearn.decomposition import PCA
def pca_mat(neigh_freqs,n_compon):
    '''
    main pca analysis.  Preprocessing includes adding 1e-5 to frequency data before taking logs.  
    These data are then z normalized for each feature.
    
    neigh_freqs: shape = (number_of_patients, number_of_celltypes), the frequency of each cell type in each patient.
    n_compon:  the number of components needed
    '''
    nf = np.log(1e-5+neigh_freqs).values
    zn  = (nf-nf.mean(0,keepdims = True))/nf.std(0,keepdims = True)
    pca = PCA(n_components=n_compon)
    loads = pca.fit_transform(zn)
    return loads,pca
def report(loads):
    for j in range(loads.shape[1]):
        a = loads[[i[0]==1 for i in neigh_freqs.index ]][:,j]
        b = loads[[i[0]==2 for i in neigh_freqs.index ]][:,j]
        print ()
        print ('{}:  {:.2f},{:.2f}, {}'.format(j,np.mean(a),np.mean(b),ttest_ind(a,b)))

# %% [code]
sum_cols2 = ['lymphatics', 'CD11c+ DCs', 'nerves',
       'CD11b+ monocytes', 'CD11b+CD68+ macrophages',
        'adipocytes', 'CD68+ macrophages',
         'Tregs', 
       'plasma cells', 'vasculature', 'B cells',
       'CD4+ T cells CD45RO+', 'CD8+ T cells', 'stroma', 'granulocytes','smooth muscle', 'CD68+CD163+ macrophages', 'tumor cells']


# %% [code]
#remove dirt neighborhood and compute PCA
cell_freqs_foll = cells2[~cells2['neighborhood10'].isin([1])].groupby(['groups','patients']).apply(lambda x: x[sum_cols2].mean(0)).dropna()

fol_l, fol_pc = pca_mat(cell_freqs_foll,10)
fol_pc.explained_variance_ratio_

# %% [code]
plt.scatter(fol_l[:,0],fol_l[:,1],c = [{True:'red',False:'blue'}[i[0]==1] for i in cell_freqs_foll.index])
plt.xlabel('PC 1')
plt.ylabel('PC 2')

# plt.savefig('celltypespcascatter.pdf')
# plt.close()

# %% [code]
from scipy.stats import ttest_ind

# %% [code]
g1 = [i[0]==1 for i in cell_freqs_foll.index]
g2 = [i[0]==2 for i in cell_freqs_foll.index]
plt.figure(figsize = (5,2))
sns.kdeplot(fol_l[g1,0],shade = True,c = 'red')
sns.kdeplot(fol_l[g2,0],shade = True,c = 'blue')

# plt.xlim(-4,4)
plt.yticks([], [])
plt.tight_layout()
ax = plt.gca()

ax.spines['right'].set_visible(False)
ax.spines['left'].set_visible(False)
ax.spines['top'].set_visible(False)
# plt.ylim(0,.3)
# plt.axis('off')
# plt.savefig('raw_figs/pc1kde.pdf')
# plt.savefig('raw_figs/pc1kde.svg')
# plt.show()

plt.figure(figsize = (5,2))
sns.kdeplot(fol_l[g1,1],shade = True,c = 'red')
sns.kdeplot(fol_l[g2,1],shade = True,c = 'blue')
plt.yticks([], [])
plt.tight_layout()

ax = plt.gca()

ax.spines['right'].set_visible(False)
ax.spines['left'].set_visible(False)
ax.spines['top'].set_visible(False)
# # plt.axis('off')
# plt.savefig('raw_figs/pc2kde.pdf')
# plt.savefig('raw_figs/pc2kde.svg')
plt.show()

# %% [code]
#PC1 difference between groups
print (ttest_ind(fol_l[g1,0],fol_l[g2,0]))

#PC2 difference between groups
print (ttest_ind(fol_l[g1,1],fol_l[g2,1]))

# %% [code]
arch = ['lymphatics','nerves','adipocytes','vasculature','stroma','smooth muscle']
vec = np.argsort(fol_pc.components_[1])
plt.bar(range(len(vec)),fol_pc.components_[1][vec],color = [{False:'blue',True:'lime'}[i in arch] for i in np.array(sum_cols2)[vec]])

plt.xticks(range(len(vec)),np.array(sum_cols2)[vec],rotation = 90);
```
