## Supplementary material for "Coordinated cellular neighborhoods orchestrate antitumoral immunity at the colorectal cancer invasive front": Analysis code files: survival submission.ipynb

```
# %% [code]
library('glmnet')
library('pROC')
library('survival')
rename <- function(X){
    rownames(X) <- X$patients
    print(dim(X[,-1]))
    return(X[,-1])
}

X_chk_density <- rename(read.csv('X_chk_density'))
X_chk_overall_patient <- rename(read.csv('X_chk_overall_pat'))

group <- read.csv('group',header = FALSE)[,-1]
time <- read.csv('Y',header = FALSE)[,-1]
censor <- read.csv('censor',header = FALSE)[,-1]
status = censor#rep(1,length(time))#censor
BN <- read.csv('BN',header = FALSE)[,-1]

# %% [code]
# survival tests
tests <- c('CD4.PD.1._9', 'Treg.ICOS._6','CD68.CD163.Ki67._6', 'Treg.ICOS._7', 'Treg.Ki67._4')
for (test in tests){
dat <-  data.frame(time = time, status = status,group = group,nbd = X_chk_density[,test])
mod0 <- coxph(Surv(time,status) ~nbd, data = dat[group==1,])
print(summary(mod0))}

# %% [code]

## overall frequency
dat <-  data.frame(time = time, status = status,group = group,nbd = X_chk_overall_patient$freq.CD4.PD.1.)
mod1 <- coxph(Surv(time,status) ~nbd, data = dat[group==1,])
print(summary(mod1))

# %% [code]
library(visreg)

# %% [code]
visreg(mod0, "nbd", ylab="log(Hazard ratio)",xlab = 'CD4+ PD-1+/nbd 9',main = 'Hazard ratio and CD4-PD1 density relationship',points=list(cex=1.5, pch=16))

# %% [code]
## ANOVA
dat <-  data.frame(time = time, status = status,group = group,overall = X_chk_overall_patient$freq.CD4.PD.1.,nbd = X_chk_density$CD4.PD.1._9)
mod2 <- coxph(Surv(time,status) ~nbd, data = dat[group==1,])
print(anova(mod1,mod2))
```
