## Supplementary material for "Coordinated cellular neighborhoods orchestrate antitumoral immunity at the colorectal cancer invasive front": Analysis code files: tensor_decomposition_submission.ipynb

```
# %% [code]
from tensorly.decomposition import non_negative_parafac,parafac,non_negative_tucker,partial_tucker,tucker
from tensorly.regression import KruskalRegressor
import tensorly as tl
import numpy as np
import pandas as pd
from matplotlib import pyplot as plt
from scipy.stats import ttest_ind
from sklearn.utils import resample
from scipy.stats import ttest_1samp
import itertools
%pylab inline

# %% [code]
cell_counts = pd.read_csv('cell_counts.csv',header=None)
ct_frq = pd.read_pickle('cell_type_freqs_per_patient')
chk_frq = pd.read_pickle('checkpoint_to_parent_freqs_perneighborhood_perpatient')
neigh_frqs = pd.read_pickle('neigh_freqs_per_patient')
all_frqs = pd.read_pickle('allcell_type_counts_perneighborhood_perpatient')

annotations = pd.read_csv('blocknumb_to_patient_no.csv')
survival = pd.read_csv('updated_survival.csv', header = 1)
annotations['B-Number'] = annotations['B-Number'].astype(str)
survival['Probenummer_Original'] = survival['Probenummer_Original'].astype(str)
# i checked that the indexes are the same (B numbers warped)
annotations['OS'] = survival['OS'].loc[annotations.index.values]
annotations['OS_censor'] = survival['OS_Censor'].loc[annotations.index.values]
annotations['DFS'] = survival['DFS'].loc[annotations.index.values]
annotations['DFS_censor'] = survival['DFS_Censor'].loc[annotations.index.values]
annotations = annotations.set_index('Patient')
good_patients = annotations[~annotations['OS'].isna()].index.values
#good_patients = good_patients[annotations['DFS'].loc[good_patients]>0]
annotations['B-Number'].to_csv('BN')
group = (annotations['Group'].loc[good_patients]==2).astype(int)

cells = ['CD4+ T cells CD45RO+',
 'granulocytes',
 'vasculature',
 'plasma cells',
 'stroma',
 'tumor cells',
 'CD68+CD163+ macrophages',
 'CD8+ T cells',
 'smooth muscle']

parents = {'CD4+ICOS+':'CD4+',
 'CD4+Ki67+':'CD4+',
 'CD4+PD-1+':'CD4+',
 'CD68+CD163+ICOS+':'CD68+CD163+',
 'CD68+CD163+Ki67+':'CD68+CD163+',
 'CD68+CD163+PD-1+':'CD68+CD163+',
 'CD68+ICOS+':'CD68+',
 'CD68+Ki67+':'CD68+',
 'CD68+PD-1+':'CD68+',
 'CD8+ICOS+':'CD8+',
 'CD8+Ki67+':'CD8+',
 'CD8+PD-1+':'CD8+',
'Treg-ICOS+':'CD4+',
 'Treg-Ki67+':'CD4+',
 'Treg-PD-1+':'CD4+'}
#  'Treg-ICOS+':'CD25+FOXP3+',
#  'Treg-Ki67+':'CD25+FOXP3+',
#  'Treg-PD-1+':'CD25+FOXP3+'}


all_cells =  ['granulocytes',
 'vasculature',
 'CD4+ T cells CD45RO+',
 'tumor cells',
 'stroma',
 'CD68+CD163+ macrophages',
 'adipocytes',
 'plasma cells',
 'CD8+ T cells',
 'Tregs',
 'CD4+ T cells',
 'CD11c+ DCs',
 'B cells',
 'CD11b+CD68+ macrophages',
 'smooth muscle',
 'tumor cells / immune cells',
 'immune cells / vasculature',
 'immune cells',
 'NK cells',
 'nerves',
 'CD68+ macrophages GzmB+',
 'CD68+ macrophages',
 'lymphatics',
 'CD11b+ monocytes']


# %% [code]
cells2 = pd.read_pickle('cells2_salil')

# %% [code]
#build the tensors

#overall_counts = x.groupby('patients').apply(lambda x: x.loc[x['neighborhood10'].isin(nbs),:].sum(axis=0))
chks = cell_counts[0]


nbs1 = [0,2,3,4,6,7,8,9]
T1 = np.zeros((len(group1_patients),len(nbs1),len(chks)))
nbs2 = [0,2,3,4,6,7,8,9]
T2 = np.zeros((len(group2_patients),len(nbs2),len(chks)))

nbd_counts1 = x.loc[:,all_cells+['patients','neighborhood10']].groupby(['patients','neighborhood10']).sum()[all_cells].sum(axis=1).unstack().loc[group1_patients,:]
nbd_counts2 = x.loc[:,all_cells+['patients','neighborhood10']].groupby(['patients','neighborhood10']).sum()[all_cells].sum(axis=1).unstack().loc[group2_patients,:]


for i,nb in enumerate(nbs1):
    for j,chk in enumerate(chks):
        #T1[:,i,j] = x.loc[x['neighborhood10']==nb,:].set_index('patients').loc[group1_patients,chk].fillna(0).values/nbd_counts1[nb].values
        T1[:,i,j] = x.loc[x['neighborhood10']==nb,:].set_index('patients').loc[group1_patients,chk].fillna(0).values
        
        
for i,nb in enumerate(nbs2):
    for j,chk in enumerate(chks):
        #T2[:,i,j] = x.loc[x['neighborhood10']==nb,:].set_index('patients').loc[group2_patients,chk].fillna(0).values/nbd_counts2[nb].values
        T2[:,i,j] = x.loc[x['neighborhood10']==nb,:].set_index('patients').loc[group2_patients,chk].fillna(0).values
        
        
        
# dat1 =np.log(1+np.nan_to_num(T1/T1.sum(axis = (0), keepdims =True)))
# dat2 = np.log(1+np.nan_to_num(T2/T2.sum(axis = (0), keepdims =True)))
# dat1 =np.log(1+np.nan_to_num(T1))
# dat2 = np.log(1+np.nan_to_num(T2))

#normalize so we have joint distributions each slice
dat1 =np.nan_to_num(T1/T1.sum((1,2), keepdims = True))
dat2 =np.nan_to_num(T2/T2.sum((1,2), keepdims = True))

# %% [code]
figsize(10,5)
def tensor_plots(dat,scale = 0.4, savename = None):
    pal = sns.color_palette('bright',10)
    palg = sns.color_palette('Greys',10)
    mat1 = np.zeros((5,15))
    #finding the elbow point
    for i in range(2,15):
        for j in range(1,5):
            facs_overall = non_negative_tucker(dat,rank=[j,i,i],random_state = 2336)
            mat1[j,i] = np.mean((dat- tl.tucker_to_tensor(factors = facs_overall[1], core = facs_overall[0]))**2)
    person_rank = 2
    rank = 6
    facs_overall = non_negative_tucker(dat,rank=[person_rank,rank,rank],random_state = 32)

    figsize(10,5)
    plt.plot(2+np.arange(13),mat1[2][2:],c = 'red',label = 'rank = (2,x,x)')
    plt.plot(2+np.arange(13),mat1[1][2:],c = 'blue',label = 'rank = (1,x,x)')
    plt.xlabel('x')
    plt.ylabel('error')
    plt.show()

    figsize(3.67*scale,2.00*scale)
    nb_scatter_size = scale*scale*45
    cel_scatter_size = scale*scale*15
    
    
    #script to draw the tissue modules (requires fine tuning for rescaling/positioning)
    for p in range(person_rank):
        for idx in range(rank):
            an = float(np.max(facs_overall[0][p][idx,:])>0.1) + (np.max(facs_overall[0][p][idx,:])<=0.1)*0.05
            ac = float(np.max(facs_overall[0][p][:,idx])>0.1) + (np.max(facs_overall[0][p][:,idx])<=0.1)*0.05

            nb_fac = facs_overall[1][1][:,idx]
            cel_fac = facs_overall[1][2][:,idx]

            cols_alpha = [(*pal[nb], an*np.minimum(nb_fac, 1.0)[i]) for i,nb in enumerate(nbs2)]
            cols = [(*pal[nb], np.minimum(nb_fac, 1.0)[i]) for i,nb in enumerate(nbs2)]
            cell_cols_alpha = [(0,0,0, an*np.minimum(cel_fac, 1.0)[i]) for i,_ in enumerate(cel_fac)]
            cell_cols = [(0,0,0, np.minimum(cel_fac, 1.0)[i]) for i,_ in enumerate(cel_fac)]
            
            plt.scatter(0.5*np.arange(len(nb_fac)), 5*idx + np.zeros(len(nb_fac)), c = cols_alpha, s = nb_scatter_size)#,edgecolors=len(cols_alpha)*[(0,0,0,min(1.0,max(0.1,2*an)))], linewidths= 0.05)
            offset = 9
            for i,k in enumerate(nbs2):
                pass
                #plt.text(0.5*i, 5*idx, k,fontsize = scale*2,ha = 'center', va = 'center',alpha = an)

            plt.scatter(-4.2+0.25*np.arange(len(cel_fac))+offset, 5*idx + np.zeros(len(cel_fac)), c = cell_cols_alpha, s = 0.5*cel_scatter_size)#,vmax = 0.5,edgecolors=len(cell_cols_alpha)*[(0,0,0,min(1.0,max(0.1,2*an)))], linewidths= 0.05)
            rect = plt.Rectangle((-0.5,5*idx-2 ),4.5,4,linewidth=scale*scale*1,edgecolor='black',facecolor='none',zorder = 0,alpha = an,linestyle = '--')
            ax = plt.gca()
            ax.add_artist(rect)
            plt.scatter([offset-5],[5*idx],c = 'black', marker = 'D', s = scale*scale*5, zorder = 5,alpha = an)
#             plt.text(offset-5,5*idx,idx,color = 'white',alpha = an, ha = 'center', va = 'center',zorder = 6,fontsize = 4.5)
            plt.scatter([offset-4.5],[5*idx],c = 'black', marker = 'D', s = scale*scale*5, zorder = 5,alpha = ac)
#             plt.text(offset-4.5,5*idx,idx,color = 'white',alpha = ac, ha = 'center', va = 'center', zorder = 6,fontsize = 4.5)

            rect = plt.Rectangle((offset-4.5,5*idx-2 ),4.5,4,linewidth=scale*scale*1,edgecolor='black',facecolor='none',zorder = 0, alpha = ac,linestyle = '-.')
            ax.add_artist(rect)

#        cols = lcols    
#         for i,chk in enumerate(cell_cols):
#                 plt.text(-4+offset+0.2*i, 27.5, chk, rotation = 45, color = 'black',ha = 'left', va = 'bottom',fontsize = scale*15,alpha = ac)
        for nb_i in range(rank):
            for cel_i in range(rank):
                plt.plot([-3+offset -2, -4+offset - 0.5],[5*nb_i, 5*cel_i], color = 'black', linewidth =2*scale*scale*1* min(1.0, max(0,-0.00+facs_overall[0][p][nb_i,cel_i])),alpha = min(1.0, max(0.000,-0.00+10*facs_overall[0][p][nb_i,cel_i])))#max(an,ac))


        plt.ylim(-5, 30)
        plt.axis('off')
        
        if savename:
            plt.savefig(savename+'_%d.pdf'%p)
            plt.savefig(savename+'_%d.svg'%p)
        plt.show()
    


# %% [code]
tensor_plots(dat1,scale = 1,savename = 'clr_updated')
tensor_plots(dat2,scale = 1,savename='dii_updated')

# %% [code]
```
