## Supplementary material for "Coordinated cellular neighborhoods orchestrate antitumoral immunity at the colorectal cancer invasive front": Analysis code files: voronoi_submission.ipynb

```
# %% [code]
%load_ext autoreload

%autoreload 2
import pandas as pd
import sys
sys.path.append('/media/compute/CellHier/graham/cellhier/')
from cellhier.general import *
cells2 = pd.read_pickle('cells2_salil')

# %% [code]
from cellhier.voronoi import draw_voronoi_scatter
import seaborn as sns

# %% [code]
spot = cells2[cells2['File Name']=='reg059_A']

_ = draw_voronoi_scatter(spot,[],)

# %% [code]

spot = cells2[cells2['File Name']=='reg021_A']


sizes = draw_voronoi_scatter(spot,[],voronoi_kwargs={'size_max':3000})

# %% [code]
spot = cells2[cells2['File Name']=='reg038_A']
cd4s = spot[spot['CD4+']==1]
_ = draw_voronoi_scatter(spot,cd4s)

# %% [code]
spot = cells2[cells2['File Name']=='reg065_A']
cd4s = spot[spot['CD4+']==1]
_ = draw_voronoi_scatter(spot,cd4s)
```
