## Supplemental Data 2 for "Coordinated cellular neighborhoods orchestrate antitumoral immunity at the colorectal cancer invasive front"

[1] Department of Microbiology & Immunology, Stanford University School of Medicine, Stanford, CA 94305, USA
[2] Department of Bioengineering, Stanford University School of Medicine, Stanford, CA 94305, USA
[3] Department of Dermatology, Stanford University School of Medicine, Stanford, CA 94305, USA
[4] Institute of Pathology, University of Bern, 3008 Bern, Switzerland
[5] Lead contact
[6] These authors contributed equally

#### SUPPLEMENTAL FIGURE LEGENDS

**Figure S1. Screening and validation of CODEX antibodies.** Antibodies were conjugated to DNA oligonucleotides and tested individually along with cross-validation in standard IHC using the same, non-conjugated antibody clone. Clones, manufacturers, and staining specifications are listed for each antibody, and examples of IHC staining, CODEX staining (false gray color fluorescence images), and similar areas on independent H&E-stained sections are shown. Scale bars, 100 µm.

**Figure S2. Validation and titration of CODEX antibody panel.** FFPE tonsil tissue was stained with a cocktail of 55 different DNA-conjugated antibodies (**Table S4**), and a multi-cycle experiment was performed followed by H&E staining. Images of identical tissue regions at the interface of a follicle (top left in each image) and epithelium (bottom right in each image) are depicted in false gray color for each antibody; H&E staining is also shown.

**Figure S3. CODEX antibody validation.** An FFPE tonsil section was stained with a 55-marker CODEX panel (**Table S4**), and a multi-cycle reaction was performed. Top left panel: Overview in five-color overlay image with Hoechst (blue; nuclei), CD31 (yellow; vasculature), CD3 (red; T cells), CD20 (green; B cells), and pan-cytokeratin (CK, white; epithelium). Inset: H&E staining. Regions 1-4 are indicated by white rectangles. Region 1: Six-color overlay and single-marker images of a follicle with CD57 (red), ICOS (also known as CD278, green), PD-1 (also known as CD279, blue), VISTA (cyan), LAG-3 (also known as CD223, white), and Ki-67 (magenta). Region 2: Six-color overlay and single-marker images of an inflamed epithelial-lymphoid parenchyma interface with CD15 (blue), CD68 (red), CD163 (cyan), CD56 (white), PD-L1 (also known as CD274, green), and EGFR (magenta). Region 3: Six-color overlay and single-marker images of a follicle with CD4 (red), CD8 (green), CD25 (yellow), CD45RA (blue), CD45RO (cyan), and FOXP3 (magenta). A $CD4^+CD25^{hi}FOXP3^+CD45RO^+$ regulatory T cell is indicated by the white arrow. Region 4: Six-color and two-color overlay images of an epithelial region with Pdpn (green), CD34 (yellow), EMA (also known as MUC-1, white), CD45 (blue), vimentin (cyan), and SMA (magenta).

**Figure S4. Multi-tumor tissue microarray (TMA).** Representative H&E-stained section. For details of tissues and tumor types, see **Table S5**. Scale bar, 1 mm.

**Figure S5. Intact tissue and DNA-antibody-antigen complexes during CODEX multicycle experiment. (A)** FFPE tonsil tissue was stained with a cocktail of nine different DNA-conjugated antibodies. Antibodies were repeatedly rendered visible using complementary fluorescent oligonucleotides in 33 cycles with blank cycles to measure autofluorescence (no fluorescent oligonucleotides added) at the beginning, between each rendering cycle, and at the end. The microscope light exposure times were kept constant for each antibody in each cycle. Hoechst nuclear stain was used as a reference. **(B)** Example images of nuclear marker Ki-67-Alexa488 and membrane markers CD20-ATTO550 and CD3-Alexa647 in cycles 4, 10, and 20 are shown. Images are representative of cycles and nuclear and membrane markers that are not shown. **(C)** Comparison of fluorescence intensity profiles from cycles 4 and 20, as measured by ImageJ software on the yellow lines in panel B.

**Figure S6. Marker signal strength, autofluorescence, and tissue integrity during CODEX multicycle experiment.** Data are from the FFPE tonsil tissue stained with nine DNA-conjugated antibodies as described in **Figure S5**. Cells were segmented using the CODEX toolkit and clustered using X-shift (VorteX). **(A)** Mean marker expression for CD45 (ATTO550), CD20 (ATTO550), and CD3 (Alexa647) on lymphocytes (combined $CD20^+$ cells and $CD3^+$ cells). **(B)** Mean marker expression for Na-K-ATPase (Alexa488) and pan-cytokeratin (ATTO550) on epithelial cells (pan-cytokeratin$^+$ cells). **(C)** Mean marker expression for Ki-67 (Alexa488) and CD45 (ATTO550) on proliferating cells (Ki-67$^+$ cells). **(D)** Mean marker expression for HLA-DR (Alexa488) and CD45 (ATTO550) on antigen-presenting cells (HLA-DR$^+$ cells). For lymphocytes, epithelial cells, and proliferating cells, 1500 cells were sampled; for antigen presenting cells, >250 cells were sampled. **(E)** Mean autofluorescence levels on all cells combined measured in each channel in blank cycles (no fluorescent DNA probes added). **(F)** Mean expression of the nuclear marker Hoechst per cell in cycle 20 vs. cycle 1. **(G)** Representative image of H&E staining performed after cycle 33.

**Figure S7. Colorectal cancer (CRC) TMAs.** Representative H&E-stained section for each CRC TMA. Cores are arranged according to patient number (1-35), with two cores per patient per TMA (4 cores per patient in total). **(A)**, TMA A and **(B)**, TMA B. See Table S1 for detailed patients' characteristics. Scale bar, 1 mm.

**Figure S8. CRC CODEX antibody panel.** Each marker of the CRC CODEX panel (**Table S4**) is depicted individually for one representative TMA spot (spot 36 of TMA 1; patient 18). CD30 and MMP12 were not detectable in any spots in either TMA and were therefore omitted. H&E and Hoechst nuclear stainings are shown for morphological reference.

**Figure S9. Comparison of unsupervised X-shift clustering vs. supervised manual gating of cell populations.** Flow cytometry standard (FCS) files from segmented images were imported into CellEngine (www.cellengine.com). Gates were tailored individually for each file and cell population. (1) Cleanup gating: Nucleated cells were selected for by gating on cells positive for Hoechst (cycle 1) and DRAQ5 (cycle 23), and out-of-focus events were removed by gating on the focused Z planes. (2) FCS files were exported for X-shift unsupervised clustering in VorteX and (3) were further analyzed for major immune cell types in CellEngine. The gating strategy to identify T cells ($CD3^{+}$), cytotoxic T cells ($CD3^{+}CD8^{+}$), T helper cells ($CD3^{+}CD4^{+}$), Tregs ($CD3^{+}CD4^{+}CD25^{+}FOXP3^{+}$), B cells ($CD3^{-}CD20^{+}$), NK cells ($CD3^{-}CD20^{-}CD56^{+}$), $CD68^{+}$ macrophages ($CD3^{-}CD20^{-}CD56^{-}CD68^{+}$), $CD163^{+}$ macrophages ($CD3^{-}CD20^{-}CD56^{-}CD163^{+}$), $CD68^{+}CD163^{+}$ double-positive macrophages ($CD3^{-}CD20^{-}CD56^{-}CD68^{+}CD163^{+}$), lymphatics ($CD3^{-}CD20^{-}CD56^{-}Podoplanin^{+}$), vasculature ($CD3^{-}CD20^{-}CD56^{-}CD31^{+}$), dendritic cells ($CD3^{-}CD20^{-}CD56^{-}CD11c^{+}$), granulocytes ($CD3^{-}CD20^{-}CD56^{-}CD15^{+}$), and plasma cells ($CD3^{-}CD20^{-}CD56^{-}CD68^{-}CD163^{-}CD38^{+}$) is shown.

**Figure S10. Supervised annotation of CRC clusters.** After X-shift clustering, single cells from the 143 resulting clusters were overlaid on the raw data fluorescent images and on H&E stains of TMAs based on X/Y positions and visually verified based on marker expression profiles, morphology, and localization within the tissue. Similar clusters were manually merged, resulting in 28 clusters. For each cluster, yellow crosses based on X/Y coordinates of the cells contained in that cluster were overlaid on stitched montages of all TMA cores. For each cluster, three examples of markers important for cluster identification (2 positive and 1 negative) and DRAQ5 nuclear stain (right panels) are shown as well as a global overview of cellular distribution of that cluster within a single TMA spot (yellow crosses on black background, left panel).

**Figure S11. Minimal spanning trees (MSTs) and mean maker expression of 28 CRC clusters from unsupervised X-shift clustering.** MSTs show the relationship between the clusters (edges and distances), their sizes, and their mean marker expression. MSTs were generated in VorteX for each marker analyzed.

**Figure S12. Voronoi diagrams and distribution of 28 CRC clusters. (A)** Five representative Voronoi diagrams are shown for each patient group. **(B)** The frequencies of clusters for all CRC patients and for each group. Significant differences are highlighted in bold (Mann-Whitney test). Data represent mean values from four biological replicates (TMA cores) per patient.

**Figure S13. Identification and quantification of major immune cell populations in CODEX data is comparable between supervised manual gating and unsupervised X-shift clustering.** Numbers of cells per TMA spot and relative cell frequencies for manual gating in CellEngine (left columns, blue background) vs. unsupervised clustering in VorteX (right columns, white background) of selected major cell subsets.

**Figure S14. Correlation of major immune cell populations quantified by supervised manual gating and unsupervised X-shift clustering.** Correlation diagrams based on individual TMA cores for each cell population. Data from **Figure S13**.

**Figure S15. Percentages and distributions of eight merged clusters. (A)** Distributions of eight merged clusters in all CRC patients and CLR and DII groups. **(B)** Cell numbers (left panel) and distributions of the eight merged clusters (right panel) in each individual patient.

**Figure S16. Pairwise cell -cell contacts and CNs. (A)** PCA correlating combinations of cell-type abundances in CLR vs. DII patients. Cell-type loading in principal component 1 is shown. **(B)** Heatmap of likelihood ratios of direct cell-to-cell interactions for 14 selected clusters is shown for clusters with at least 100 unique interacting cells. Gray boxes indicate less than 100 unique interacting cells; these data were omitted. Pooled data from all TMA cores are shown. **(C)** Frequencies of each CN in each patient are shown. Frequencies are z scored by column to highlight major differences between CLR patients (blue) and DII patients (orange). **(D)** The contacts between CN 1 (T cell-enriched) and CN 4 (macrophage-enriched) were computed (see Methods) and are displayed as "CN mixing" by patient group. For each patient, the mean mixing score of four TMA cores is shown (*p<0.05, Student's *t*-test).

**Figure S17. Neighborhoods and corresponding H&E stainings.** H&E-stained images **(A, C)** and the 9 identified CNs **(B, D)** are shown for all cores of TMA A (A-B) and TMA B (C-D).

**Figure S18. Neighborhood analysis independently identifies comparable cluster sets in each patient group.** Both CLR and DII patient groups were clustered separately. CNs were annotated manually. CN-0 corresponds to the imaging artifacts cluster; this cluster was omitted from the analysis shown in **Figure 4C**. Neighborhoods that did not have matching counterparts in the analysis of the combined groups are labeled "not defined".

**Figure S19. Elbow points for Tucker tensor decomposition.** Tensor decomposition loss for choices of rank in patient space, CN space, and cell-type space used for selection of decomposition rank in each patient group. Blue lines, one tissue module; red lines, two tissue modules. The elbow point was found at CN-6 and cell-type modules (red line).

**Figure S20. Gating strategy for analysis of checkpoint molecule expression on T cells and macrophage populations. (A)** Representative dot plots from CellEngine are shown for each marker and population. **(B)** Heatmap of marker-positive cell populations per patient. **(C)** Frequencies of marker-positive cell populations per patient. Data are mean values from four biological replicates (TMA cores) per patient (*p<0.05, **p<0.01, Student's *t*-test).

**Figure S21. Heatmap of estimated differential enrichment coefficients for cell types not shown in Figure 6.** Asterisks indicate CNs and cell types with a regression p-value <0.05 (not adjusted for multiple tests). A positive coefficient (red) indicates that the corresponding cell type is more enriched in DII patients than in CLR patients in the given CN.

**Figure S22. Feature importance for classification model.** Bar plot of absolute coefficient z-scores for CN-specific cell type frequencies, estimated from a model classifying patient groups using iterative resampling, as described in **Figure 6F**. The five CN-specific cell type frequencies with a coefficient importance of 0.3 or higher (left to red dotted line) were considered for assessment with respect to survival in DII patients.

#### SUPPLEMENTAL TABLES

**Table S1:** CRC patient characteristics and cell types per patient (see separate .xlsx file online)
**Table S2:** Antibodies, clones, manufacturers and corresponding CODEX oligonucleotides
**Table S3:** Sequences of CODEX oligonucleotides
**Table S4:** CODEX multi-cycle panels
**Table S5:** Tissue composition of the multi-tumor TMA
**Table S6:** Key resources

#### SCRIPTS

Folder containing 12 scripts for computational CODEX data analysis (see separate .zip file online)

### Figure S1a

#### Staining specifications | IHC | CODEX | H&E

**Antigen: CD1a**
**Clones: O10 + C1A/711**

Company:
Novus Biologicals (NBP2-34698)

Tissue: Skin

Dilution:
IHC: 1:600
CODEX: 1:100

CODEX oligo: 43-Alexa647

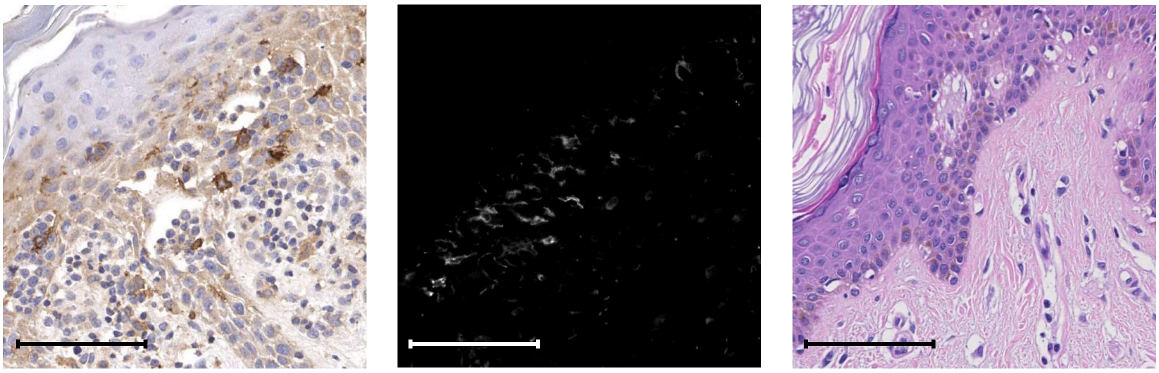

**Antigen: CD2**
**Clone: RPA-2.10**

Company:
BioLegend (300202)

Tissue: Skin T cell lymphoma

Dilution:
IHC: 1:5
CODEX: 1:25

CODEX oligo: 25-Alexa647

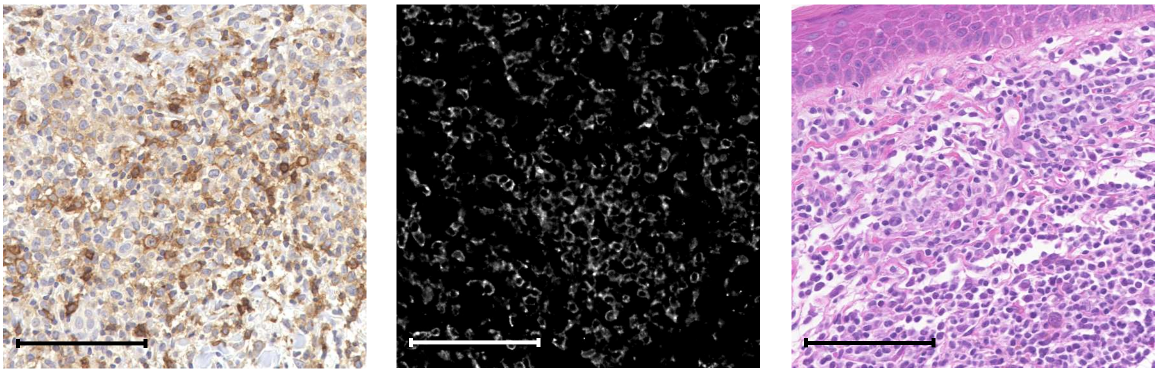

**Antigen: CD3**
**Clone: D7A6E or MRQ-39**

Companies:
Cell Signaling Technology (custom)
Cell Marque (custom)

Tissue: Tonsil

Dilution:
IHC: 1:100
CODEX: 1:100

CODEX oligo: 77-Alexa647

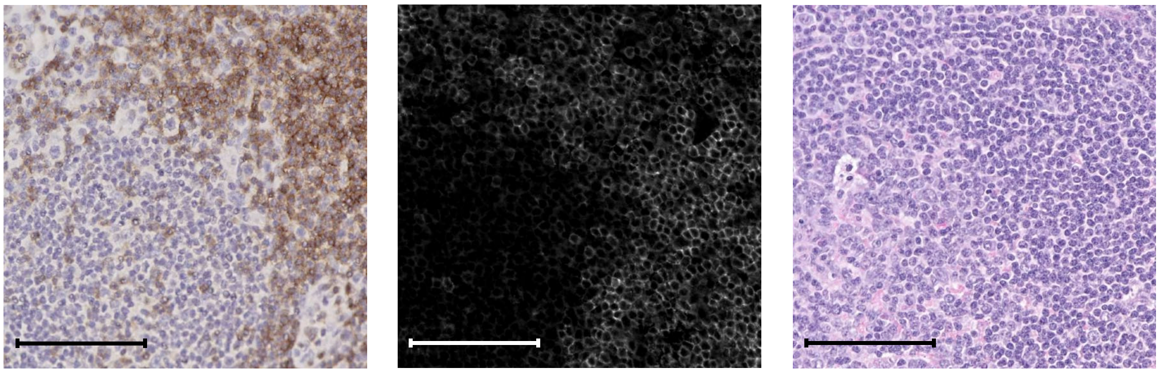

**Antigen: CD4**
**Clone: EPR6855**

Company:
Abcam (ab181724)

Tissue: Tonsil

Dilution:
IHC: 1:25
CODEX: 1:100

CODEX oligo: 20-ATTO550

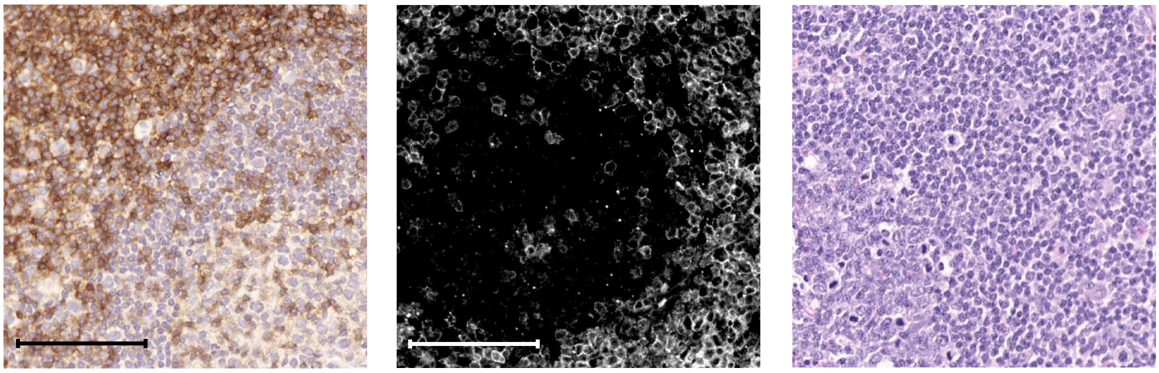

**Antigen: CD5**
**Clone: UCHT2**

Company:
BD Biosciences (555350)

Tissue: Tonsil

Dilution:
IHC: 1:5
CODEX: 1:25

CODEX oligo: 75-ATTO550

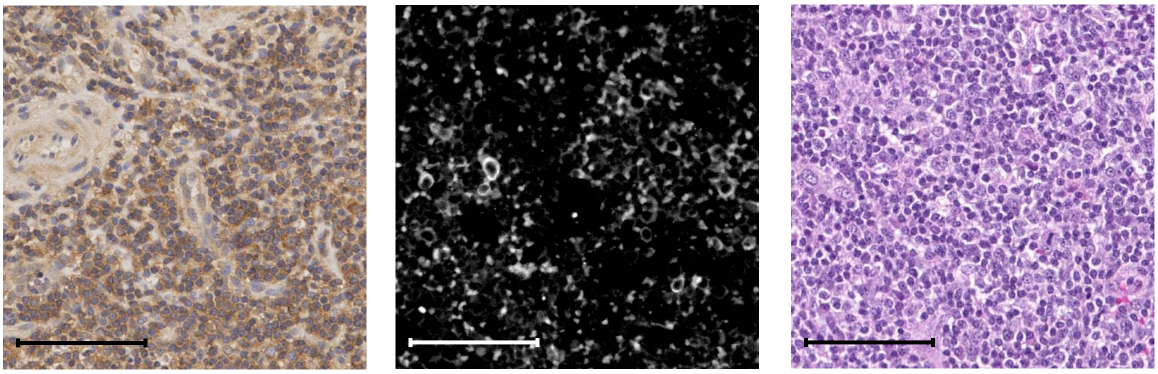

### Figure S1b

| Staining specifications | IHC | CODEX | H&E |
|---|---|---|---|

**Antigen: CD7**
**Clone: MRQ-56**

**Company:**
**Cell Marque (custom)**

**Tissue: Tonsil**

**Dilution:**
**IHC: 1:100**
**CODEX: 1:100**

**CODEX oligo: 63-Alexa488**

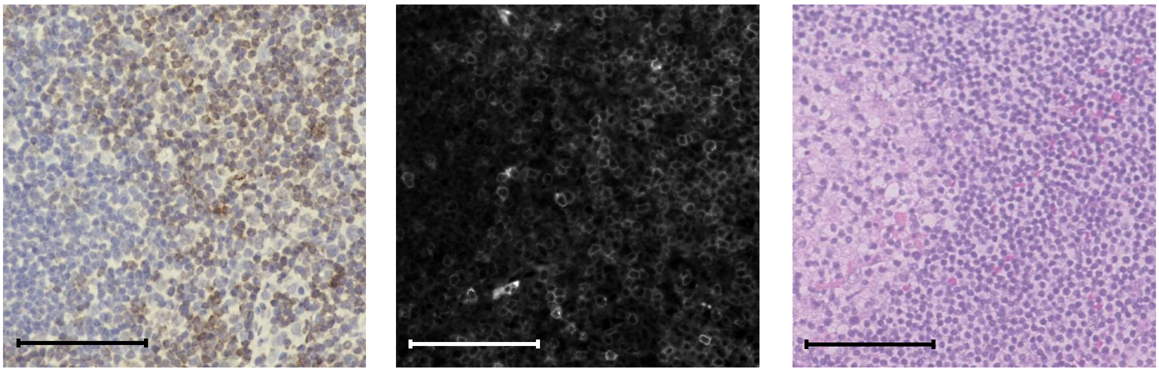

**Antigen: CD8**
**Clone: C8/144B**

**Company:**
**Cell Marque (custom)**
**Santa Cruz Bio (sc-53212)**

**Tissue: Tonsil**

**Dilution:**
**IHC: 1:100**
**CODEX: 1:50**

**CODEX oligo: 8-Alexa488**

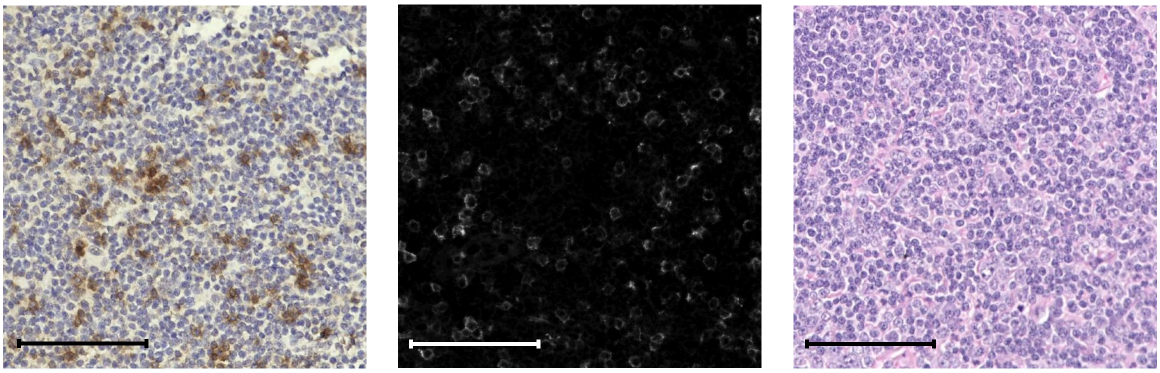

**Antigen: CD11b**
**Clone: EPR1344**

**Company:**
**Abcam (ab216445)**

**Tissue: Tonsil**

**Dilution:**
**IHC: 1:100**
**CODEX: 1:50**

**CODEX oligo: 28-Alexa647**

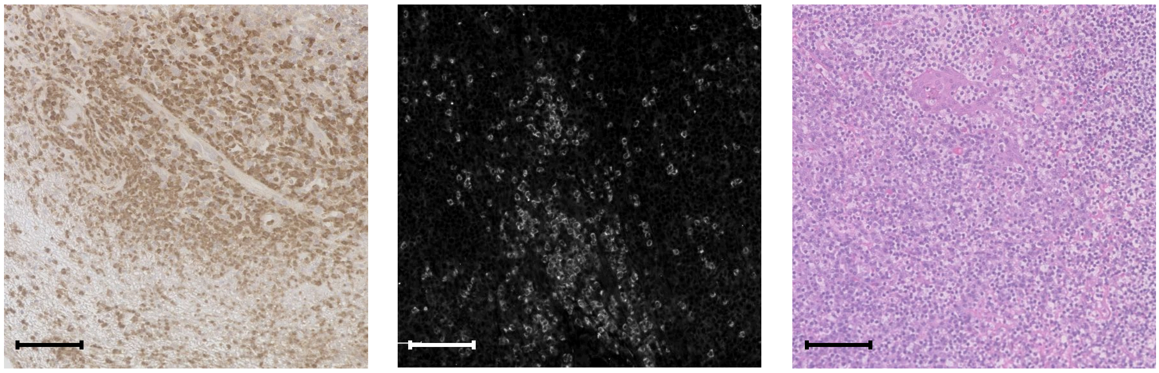

**Antigen: CD11c**
**Clone: EP1347Y**

**Company:**
**Abcam (ab216655)**

**Tissue: Tonsil**

**Dilution:**
**IHC: 1:100**
**CODEX: 1:50**

**CODEX oligo: 49-ATTO550**

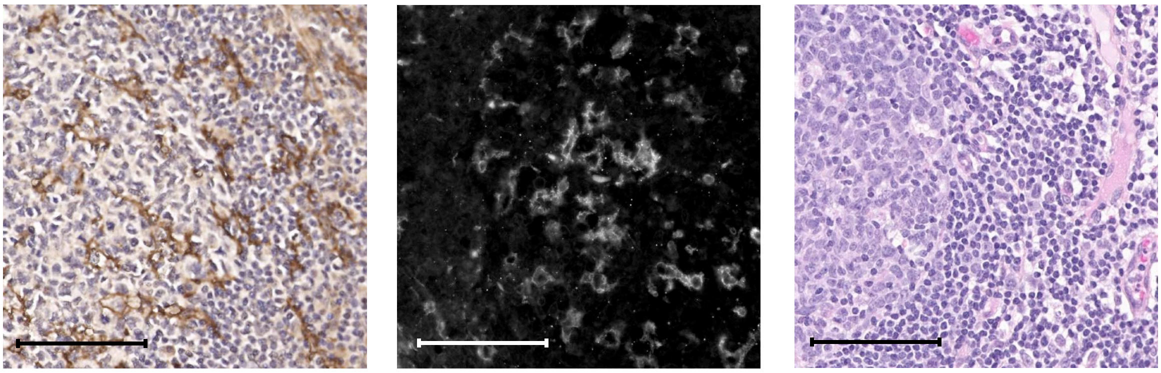

**Antigen: CD15**
**Clone: MMA**

**Company:**
**BD Biosciences (559045)**

**Tissue: Tonsil**

**Dilution:**
**IHC: 1:100**
**CODEX: 1:200**

**CODEX oligo: 14-ATTO550**

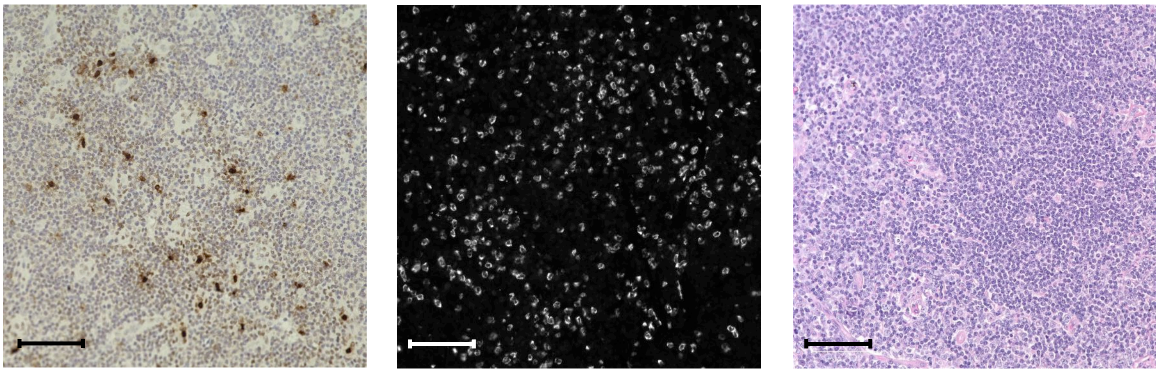

### Figure S1c

**Staining specifications**

**Antigen: CD15**
**Clone: HI98**

Company:
Biolegend (301902)

Tissue: Tonsil

Dilution:
IHC: 1:100
CODEX: 1:200

CODEX oligo: 15-Alexa488

**Antigen: CD16**
**Clone: D1N9L**

Company:
Cell Signaling Technology (custom)

Tissue: Tonsil

Dilution:
IHC: 1:100
CODEX: 1:100

CODEX oligo: 26-Alexa647

**Antigen: CD20**
**Clone: rIGEL/773**

Company:
Novus Biologicals (NBP2-54591)

Tissue: Tonsil

Dilution:
IHC: 1:100
CODEX: 1:200

CODEX oligo: 48-ATTO550

**Antigen: CD20**
**Clone: H1**

Company:
BD Biosciences (555677)

Tissue: Tonsil

Dilution:
IHC: 1:200
CODEX: 1:10

CODEX oligo: 48-ATTO550

**Antigen: CD21**
**Clone: Bu32**

Company:
Biolegend (354902)

Tissue: Tonsil

Dilution:
IHC: 1:100
CODEX: 1:50

CODEX oligo: 21-Alexa647

**IHC** | **CODEX** | **H&E**

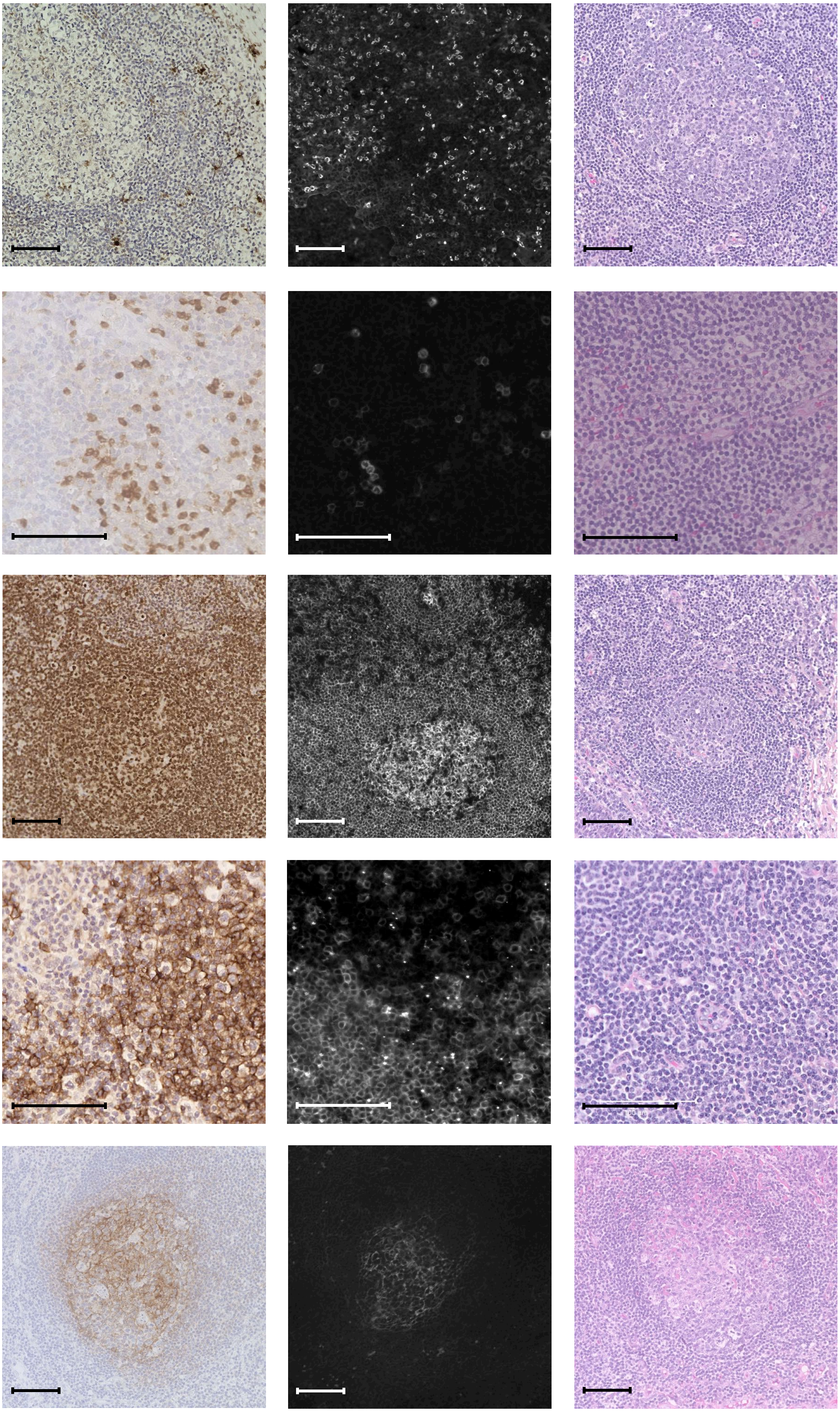

### Figure S1d

| Staining specifications | IHC | CODEX | H&E |
|---|---|---|---|

**Antigen: CD25**
**Clone: 4C9**

**Company:**
**Cell Marque (custom)**

**Tissue: Tonsil**

**Dilution:**
**IHC: 1:**
**CODEX: 1:100**

**CODEX oligo: 24-ATTO550**

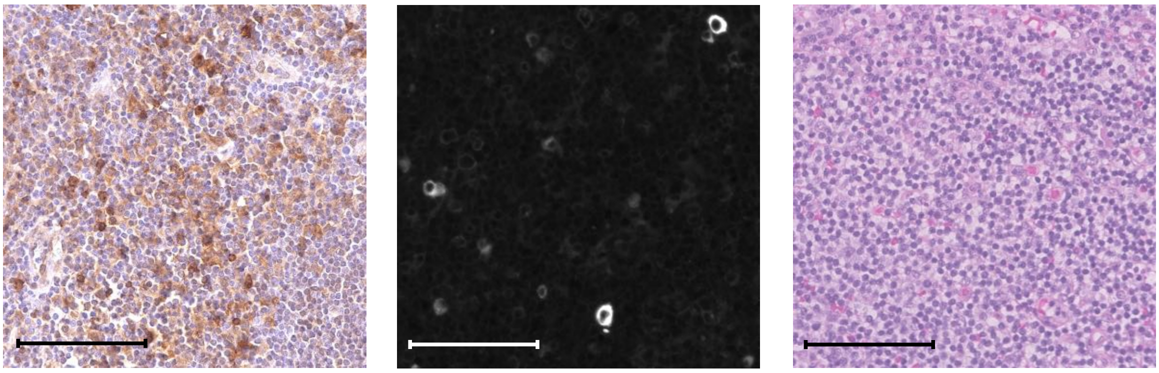

**Antigen: CD30**
**Clone: Ber-H2**

**Company:**
**Cell Marque (custom)**

**Tissue: Tonsil**

**Dilution:**
**IHC: 1:200**
**CODEX: 1:25**

**CODEX oligo: 57-ATTO550**

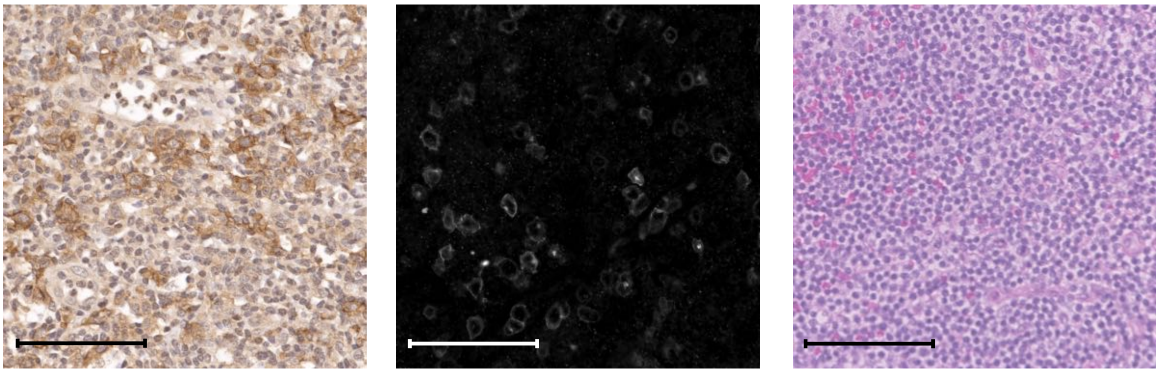

**Antigen: CD31**
**Clones:**
**C31.3 + C31.7 + C31.10**

**Company:**
**Novus Biologicals (NBP2-47785)**

**Tissue: Placenta**

**Dilution:**
**IHC: 1:100**
**CODEX: 1:200**

**CODEX oligo: 68-ATTO550**

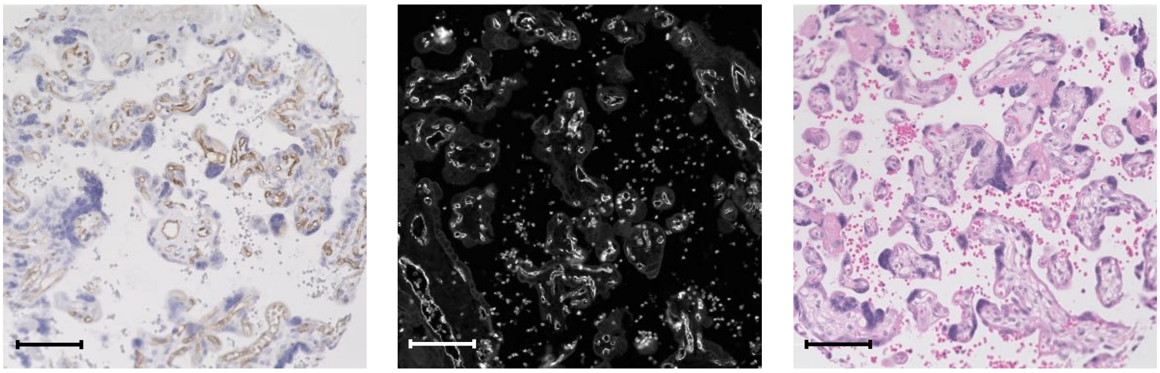

**Antigen: CD34**
**Clones:**
**QBEnd/10 + HPCA1/764**

**Company:**
**Novus Biologicals (NBP2-47909)**

**Tissue: Tonsil**

**Dilution:**
**IHC: 1:200**
**CODEX: 1:100**

**CODEX oligo: 38-ATTO550**

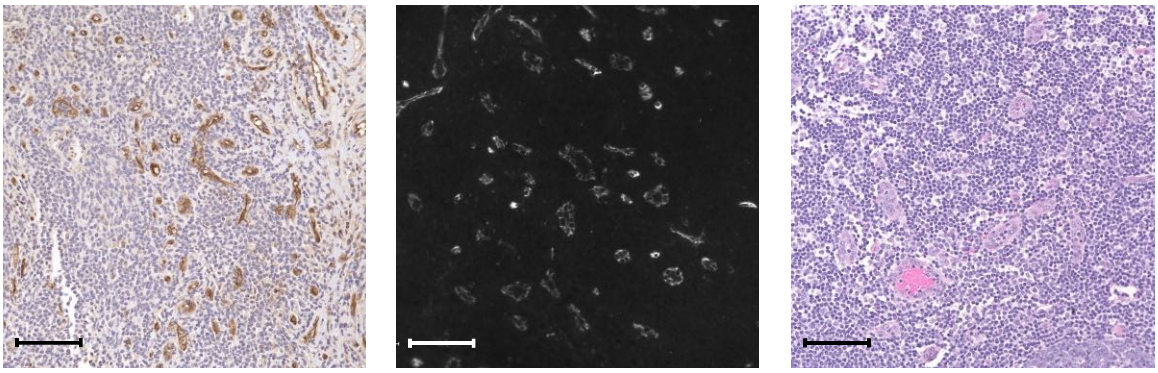

**Antigen: CD38**
**Clone: EPR4106**

**Company:**
**Abcam (ab176886)**

**Tissue: Plasmacytoma**

**Dilution:**
**IHC: 1:200**
**CODEX: 1:100**

**CODEX oligo: 66-ATTO550**

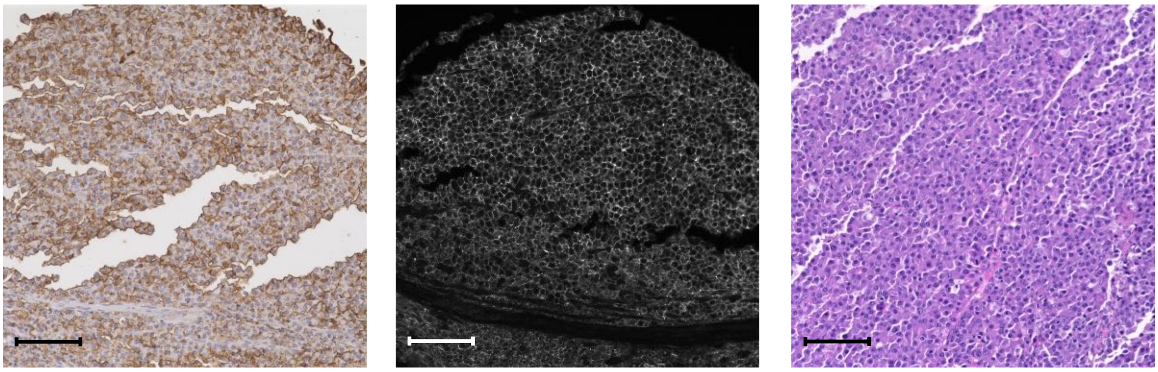

### Figure S1e

**Staining specifications**

**Antigen: CD44**
**Clone: IM-7**

**Company:**
**BD Biosciences (553131)**

**Tissue: Tonsil**

**Dilution:**
**IHC: 1:25**
**CODEX: 1:25**

**CODEX oligo: 44-Alexa488**

**Antigen: CD45**
**Clones: 2B11 + PD7/26**

**Company:**
**Novus Biologicals (NBP2-34528)**

**Dilution:**

**IHC: 1:200**
**CODEX: 1:400**

**CODEX oligo: 56-ATTO550**

**Antigen: CD45RA**
**Clone: HI100**

**Company:**
**BD Biosciences (555486)**

**Tissue: Tonsil**

**Dilution:**
**IHC: 1:50**
**CODEX: 1:50**

**CODEX oligo: 72-Alexa488**

**Antigen:  CD45RO**
**Clone: UCH-L1**

**Company:**
**Santa Cruz Bio (sc-1183)**

**Tissue: Tonsil**

**Dilution:**
**IHC: 1:50**
**CODEX: 1:25**

**CODEX oligo: 2-ATTO550**

**Antigen: CD56**
**Clone: MRQ-42**

**Company:**
**Cell Marque (custom)**

**Tissue: NK/T cell lymphoma**

**Dilution:**
**IHC: 1:200**
**CODEX: 1:100**

**CODEX oligo: 29-Alexa647**

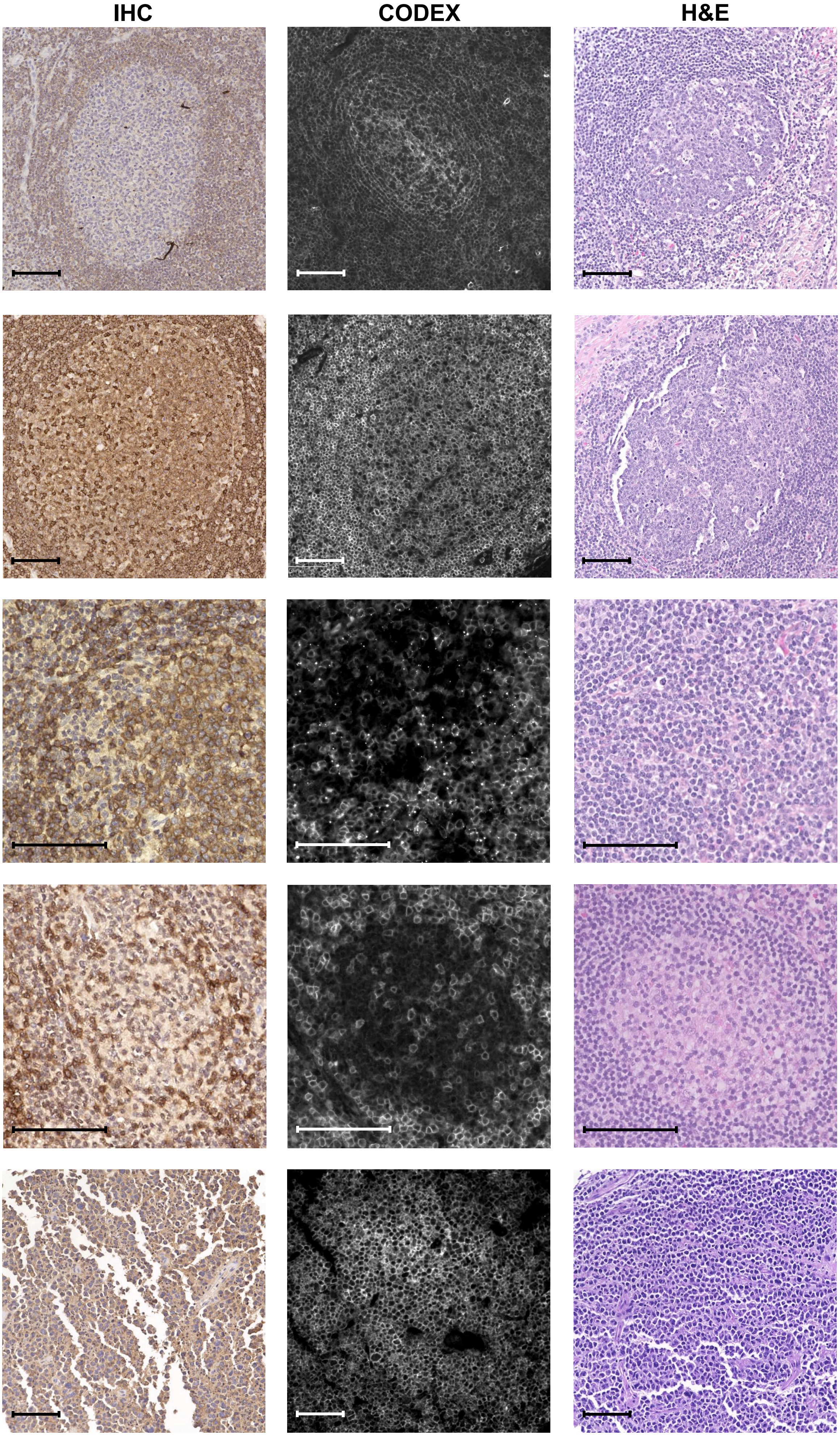

### Figure S1f

#### Staining specifications

**Antigen: CD57**
**Clone: HCD57**

**Company:**
**BioLegend (322325)**

**Tissue: Tonsil**

**Dilution:**
**IHC: 1:100**
**CODEX: 1:200**

**CODEX oligo: ST30-ATTO550**

**Antigen: CD66a**
**Clone: B1.1/CD66**

**Company:**
**BD Biosciences (551354)**

**Tissue: Tonsil**

**Dilution:**
**IHC: 1:200**
**CODEX: 1:200**

**CODEX oligo: 41-Alexa488**

**Antigen: CD68**
**Clone: D4B9C or KP-1**

**Companies:**
**Cell Signaling Technology (custom)**
**Biolegend (916104)**

**Tissue: Tonsil**

**Dilution:**
**IHC: 1:100**
**CODEX: 1:200**

**CODEX oligo: 70-Alexa647**

**Antigen: CD69**
**Clone: polyclonal**

**Company:**
**Novus Biologicals (AF2359)**

**Tissue: Tonsil**

**Dilution:**
**IHC: 1:100**
**CODEX: 1:200**

**CODEX oligo: 36-ATTO550**

**Antigen: CD71**
**Clone: MRQ-48**

**Company:**
**Cell Marque (custom)**

**Tissue: Myelolipoma**

**Dilution:**
**IHC: 1:400**
**CODEX: 1:100**

**CODEX oligo: 3-Alexa647**

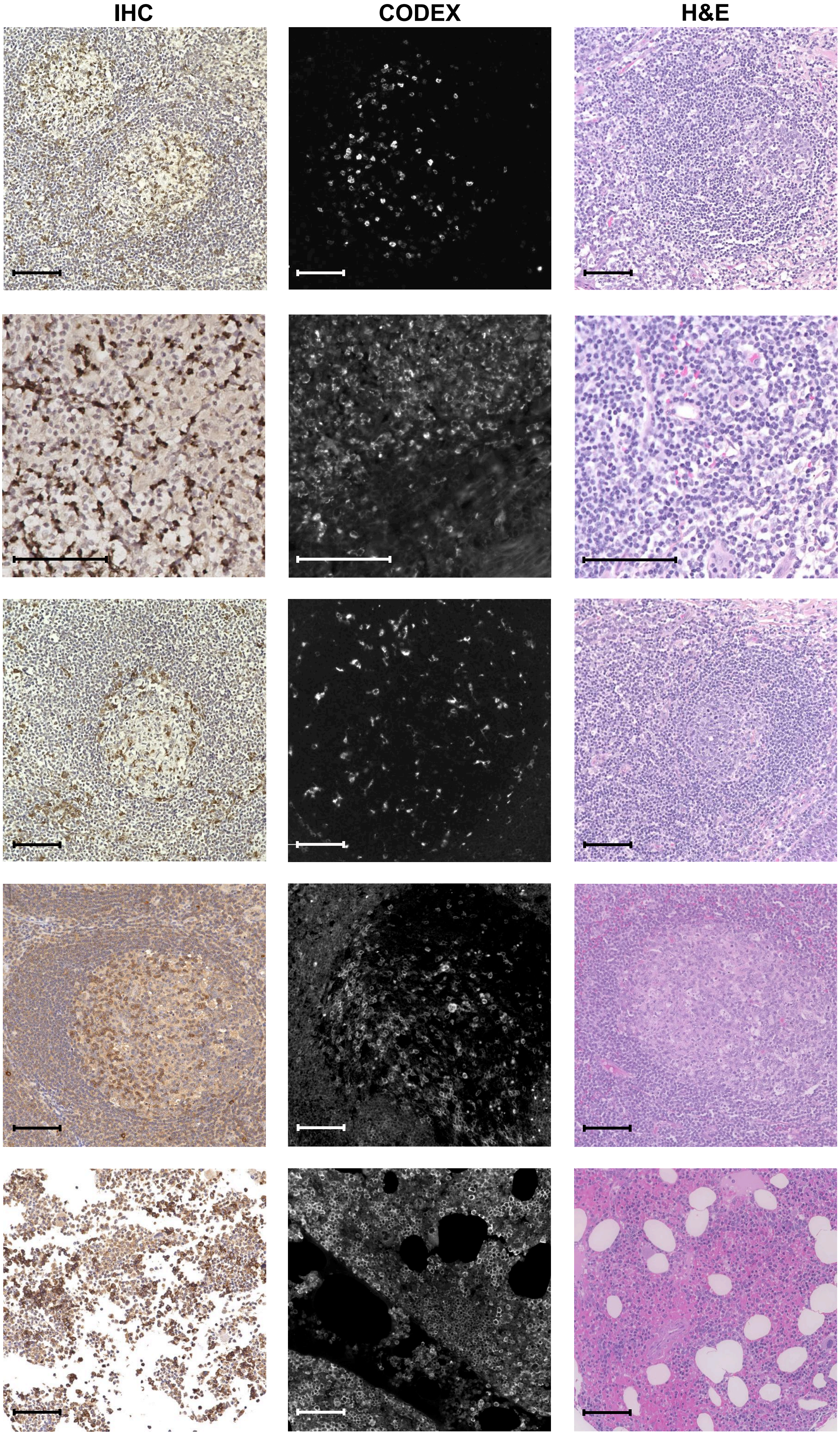

### Figure S1g

#### Staining specifications

**Antigen: CD79a**
**Clone: JBC117**

Company:
Cell Marque (custom)

Tissue: Follicular lymphoma

Dilution:
IHC: 1:300
CODEX: 1:20

CODEX oligo: 46-Alexa488

**Antigen: CD138** (Syndecan-1)
**Clone: B-A38**

Company:
Invitrogen (MA1-10091)

Tissue: Colorectal carcinoma

Dilution:
IHC: 1:100
CODEX: 1:50

CODEX oligo: ST76-Alexa647

**Antigen: CD162 (CLA)**
**Clone: HECA-452**

Company:
BD Biosciences (555946)

Tissue: Bone marrow

Dilution:
IHC: 1:600
CODEX: 1:200

CODEX oligo: 46-Alexa47

**Antigen: CD163**
**Clone: EDHu-1**

Company:
Novus Biologicals (NB110-48686)

Tissue: Tonsil

Dilution:
IHC: 1:200
CODEX: 1:200

CODEX oligo: 45-Alexa647

**Antigen: CD164**
**Clone: N6B6**

Company:
BD Biosciences (551296)

Tissue: Tonsil

Dilution:
IHC: 1:100
CODEX: 1:200

CODEX oligo: 69-Alexa488

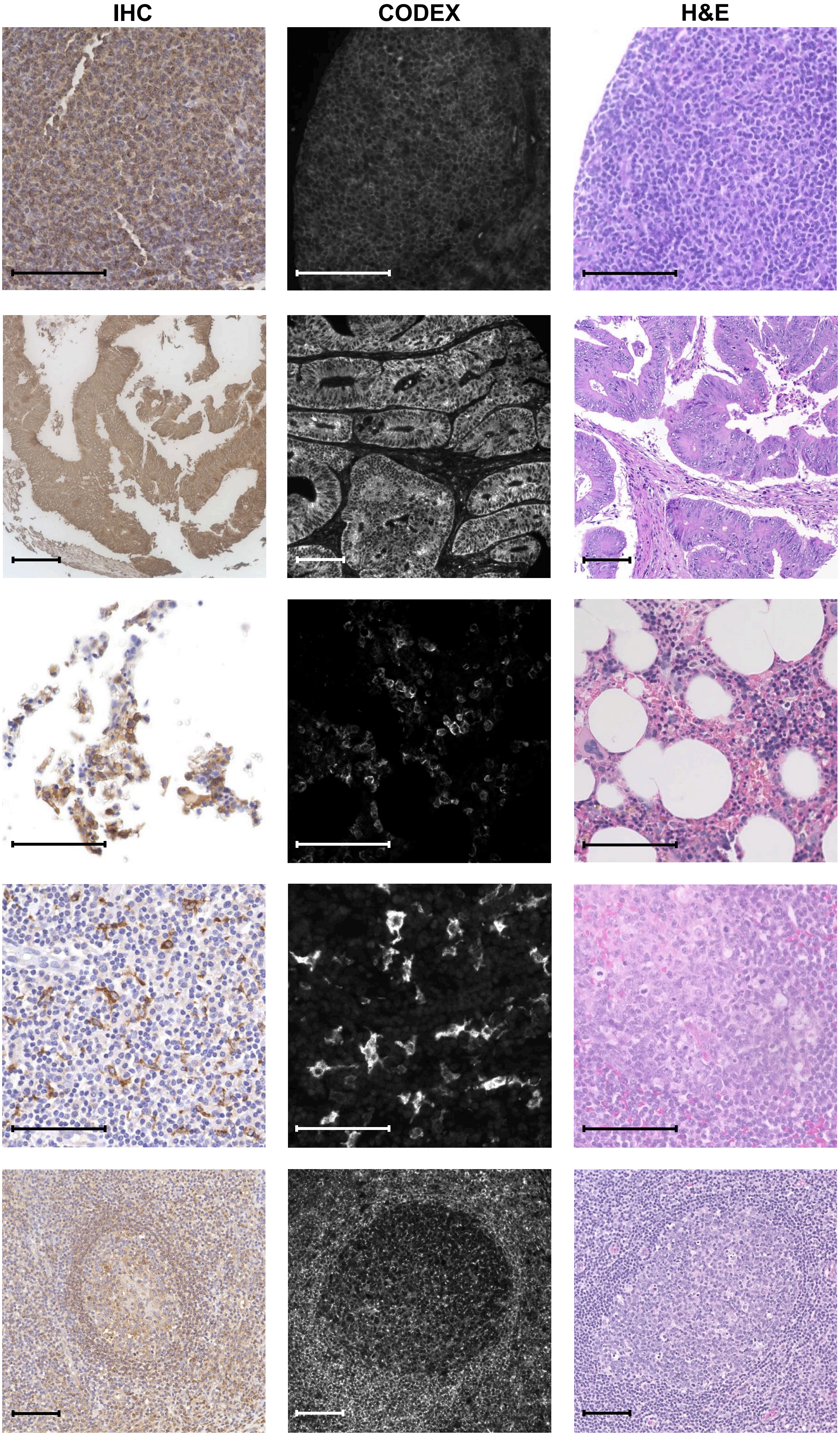

### Figure S1h

**Staining specifications** | **IHC** | **CODEX** | **H&E**

**Antigen: CD194 (CCR4)**
**Clone: L291H4**

Company:
BioLegend (359402)

Tissue: Cholangiocarcinoma

Dilution:
IHC: 1:50
CODEX: 1:10

CODEX oligo: ST2-ATTO550

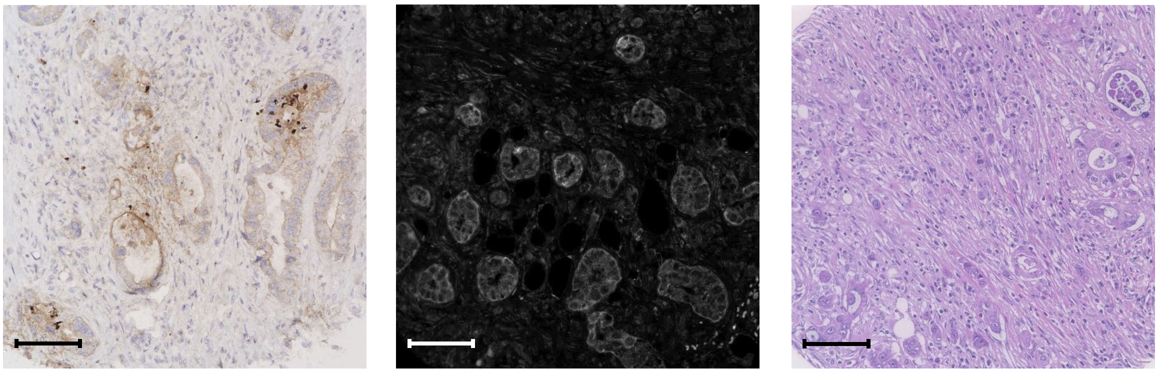

**Antigen: CD223 (LAG-3)**
**Clone: D2G4O or 17B4**

Companies:
Cell Signaling Technology (custom)
LSBio (LS-C18692-100)

Tissue: Tonsil

Dilution:
IHC: 1:300
CODEX: 1:20

CODEX oligo: 42-Alexa647

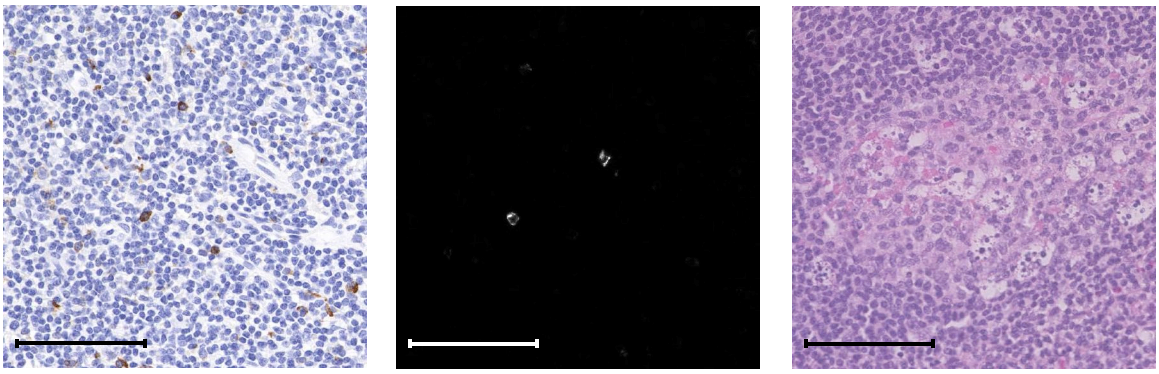

**Antigen: CD235a**
**Clone: GA-R2**

Company:
BD Biosciences (555569)

Tissue: Spleen

Dilution:
IHC: 1:400
CODEX: 1:200

CODEX oligo: 69-Alexa488

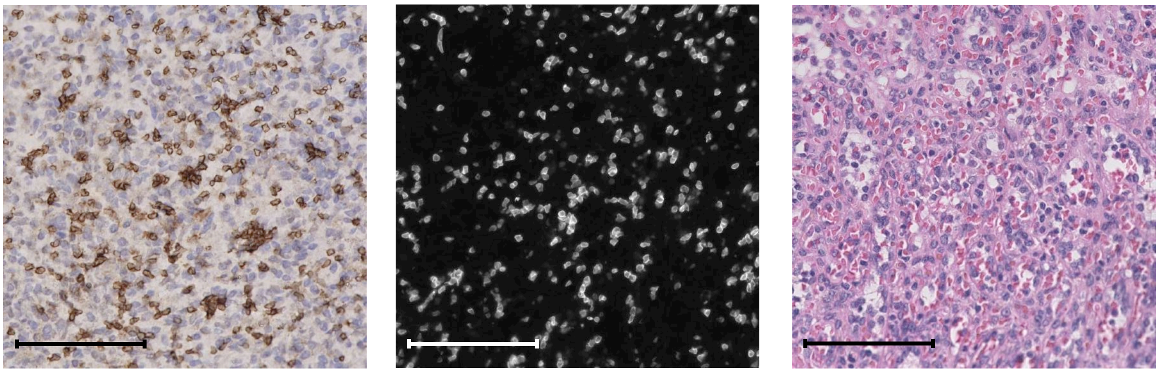

**Antigen: CD274 (PD-L1)**
**Clone: E1L3N**

Company:
Cell Signaling Technology (custom)

Tissue: Tonsil

Dilution:
IHC: 1:50
CODEX: 1:100

CODEX oligo: 11-ATTO550

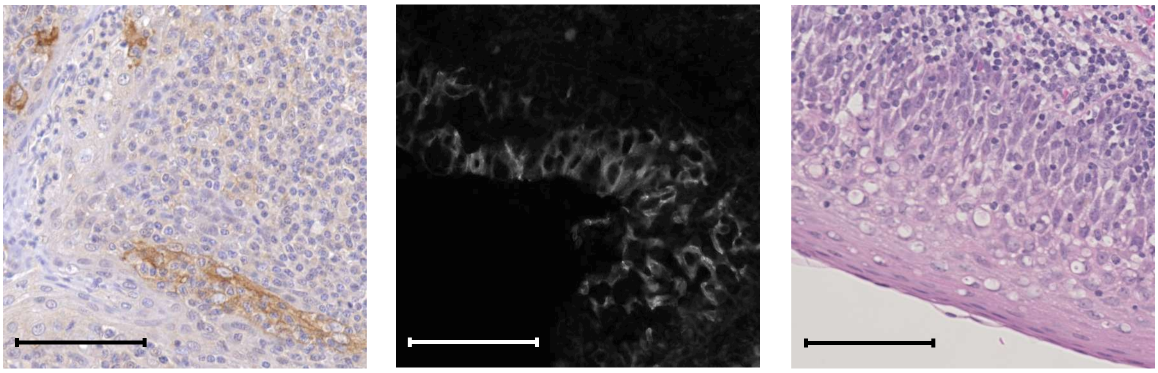

**Antigen: CD278 (ICOS)**
**Clone: D1K2T**

Company:
Cell Signaling Technology (custom)

Tissue: Tonsil

Dilution:
IHC: 1:200
CODEX: 1:20

CODEX oligo: 74-ATTO550

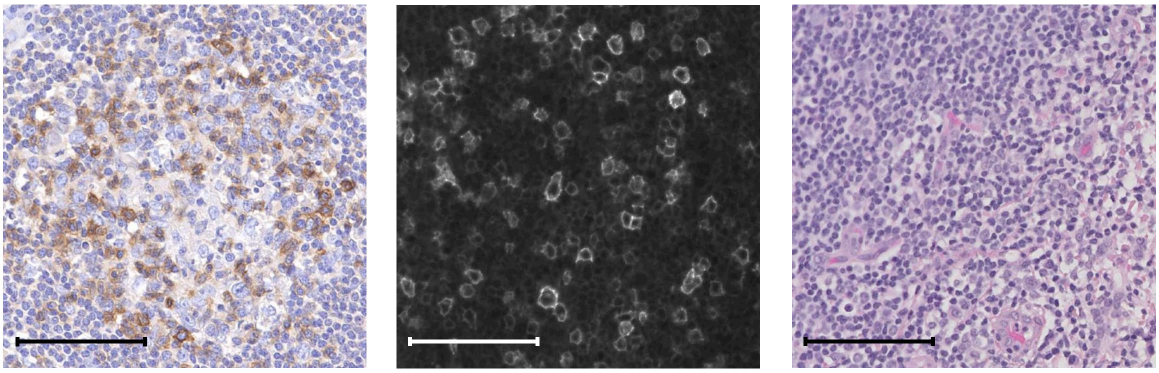

### Figure S1i

**Staining specifications**

**Antigen: CD279 (PD-1)**
**Clone: D1K2T**

Company:
Cell Signaling Technology (custom)

Tissue: Tonsil

Dilution:
IHC: 1:200
CODEX: 1:25

CODEX oligo: 23-ATTO550

**Antigen: α-SMA**
**Clone: polyclonal**

Company:
Abcam (ab5694)

Tissue: Tonsil

Dilution:
IHC: 1:200
CODEX: 1:200

CODEX oligo: 69-Alexa488

**Antigen: β-catenin**
**Clones: polyclonal or 14**

Company:
Novus Biologicals (AF1329)
Cell Marque (custom)

Tissue: Stomach

Dilution:
IHC: 1:100
CODEX: 1:25

CODEX oligo: 51-Alexa647

**Antigen: BCL-2**
**Clone: 124**

Company:
Cell Marque (custom)

Tissue: Tonsil

Dilution:
IHC: 1:200
CODEX: 1:25

CODEX oligo: 41-Alexa647

**Antigen: CDX2**
**Clone: CDX2/1690**

Company:
Novus Biologicals (NBP2-54472)

Tissue: Colorectal adenocarcinoma

Dilution:
IHC: 1:200
CODEX: 1:25

CODEX oligo: 53-Alexa647

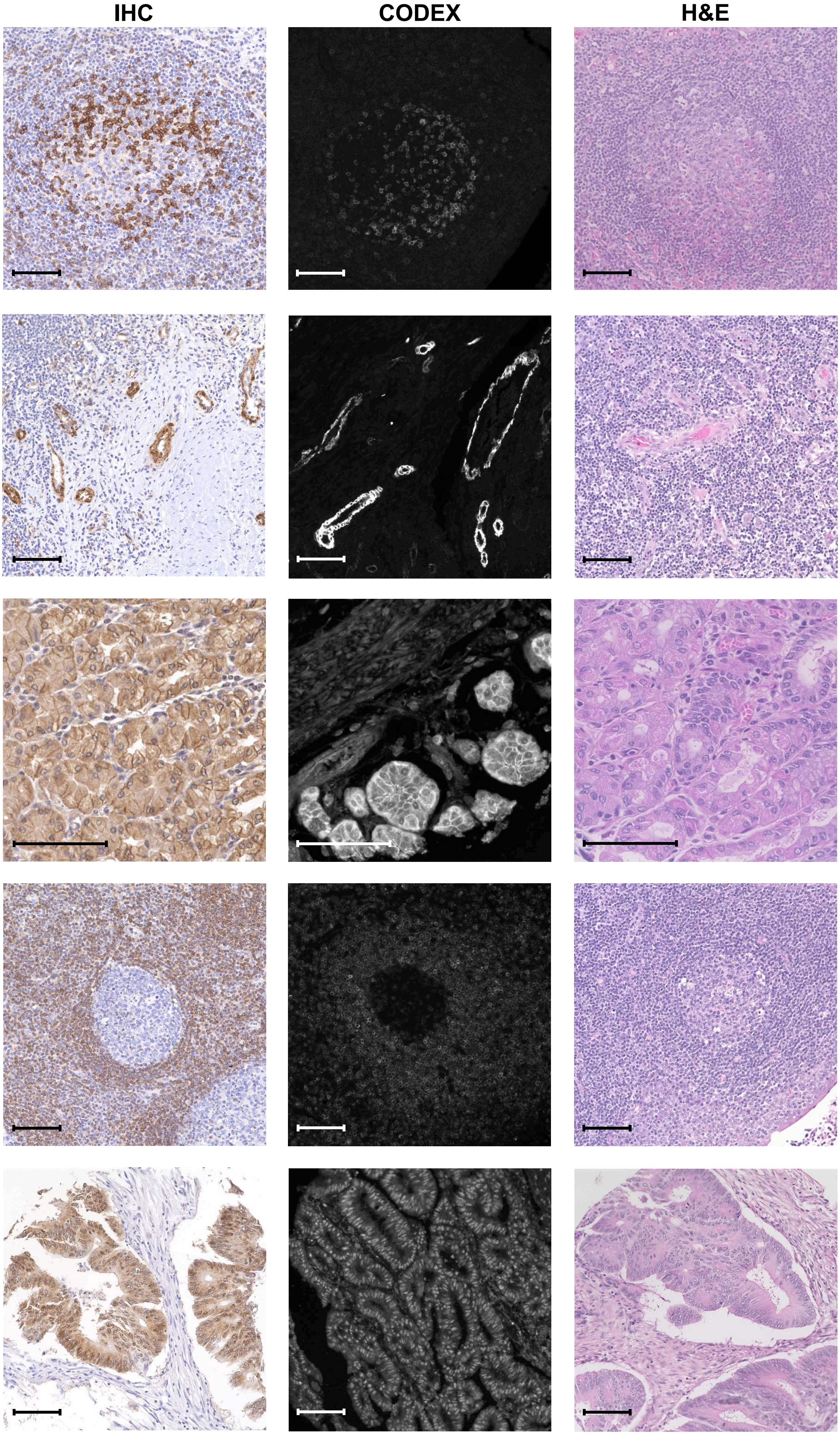

### Figure S1j

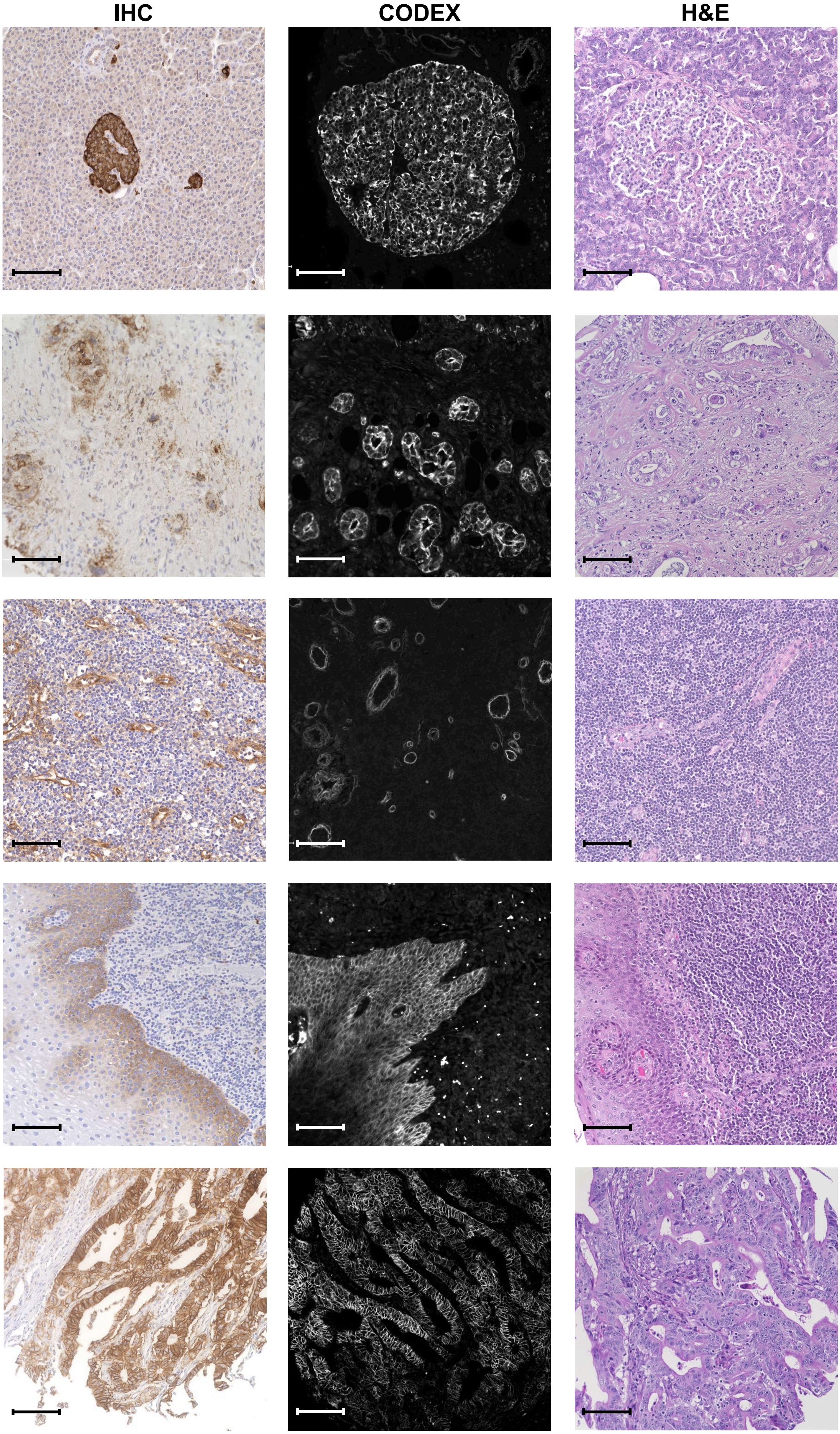

#### Staining specifications

**Antigen: Chromogranin A**
**Clones:**
**LK2H10 + PHE5 + CGA/414**

Company:
Novus Biologicals (NBP2-34674)

Tissue: Pancreas

Dilution:
IHC: 1:400
CODEX: 1:50

CODEX oligo: 43-Alexa488

**Antigen: CK7**
**Clone: OV-TL12/30**

Company:
Novus Biologicals (NBP2-47940)

Tissue: Cholangiocarcinoma

Dilution:
IHC: 1:100
CODEX: 1:50

CODEX oligo: 3-Alexa488

**Antigen: Collagen IV**
**Clone: polyclonal**

Company:
Abcam (ab6586)

Tissue: Tonsil

Dilution:
IHC: 1:200
CODEX: 1:200

CODEX oligo: 33-Alexa647

**Antigen: EGFR**
**Clone: D38B1**

Company:
Cell Signaling Technology (custom)

Tissue: Tonsil

Dilution:
IHC: 1:50
CODEX: 1:25

CODEX oligo: 58-Alexa488

**Antigen: EpCAM**
**Clone: Ber-EP4**

Company:
Cell Marque (custom)

Tissue: Gastric adenocarcinoma

Dilution:
IHC: 1:100
CODEX: 1:25

CODEX oligo: 70-Alexa647

### Figure S1k

#### Staining specifications

**Antigen: FOXP3**
**Clone: 236A/E7**

Company:
Invitrogen (14-4777-80)

Tissue: Tonsil

Dilution:
IHC: 1:20
CODEX: 1:100

CODEX oligo: 61-Alexa647

**Antigen: GATA3**
**Clone: L50-823**

Company:
Cell Marque (custom)

Tissue: Breast lobular carcinoma

Dilution:
IHC: 1:300
CODEX: 1:100

CODEX oligo: 60-Alexa647

**Antigen: GFAP**
**Clone: 2.2B10**

Company:
Invitrogen (130300)

Tissue: Glioblastoma

Dilution:
IHC: 1:100
CODEX: 1:25

CODEX oligo: 46-Alexa488

**Antigen: Granzyme B**
**Clone: EPR20129-217**

Company:
Abcam (ab219803)

Tissue: Tonsil

Dilution:
IHC: 1:200
CODEX: 1:100

CODEX oligo: 81-Alexa647

**Antigen: Hep-Par 1**
**Clone: OCH1E5**

Company:
Santa Cruz Bio (sc-58693)

Tissue: Hepatocellular carcinoma

Dilution:
IHC: 1:25
CODEX: 1:100

CODEX oligo: 28-Alexa488

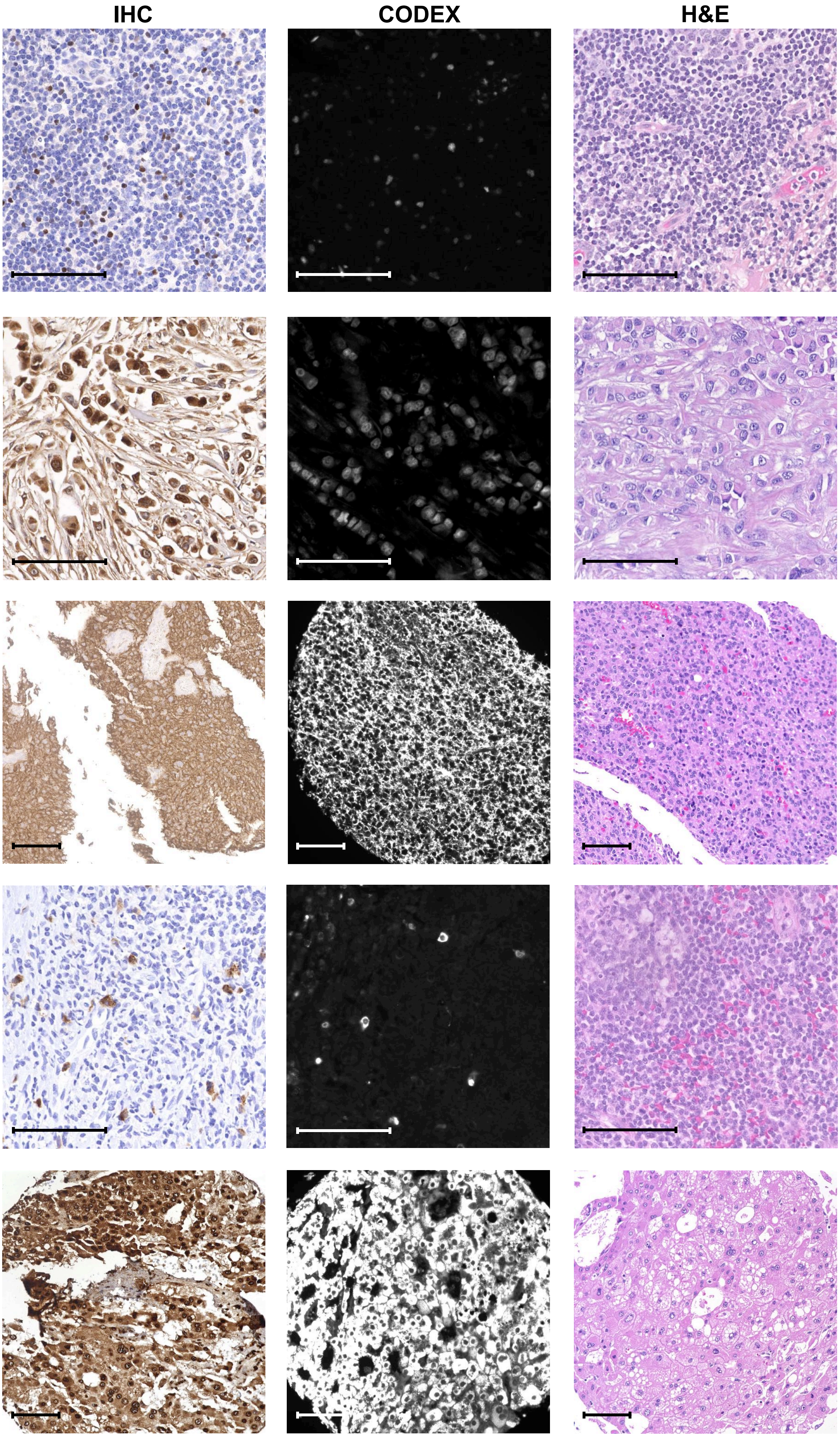

### Figure S1I

#### Staining specifications

**Antigen: Histone H3**
**Clone: D1H2**

**Company:**
**Cell Signaling Technology (custom)**

**Tissue: Tonsil**

**Dilution:**
**IHC: 1:100**
**CODEX: 1:50**

**CODEX oligo: 30-ATTO550**

**Antigen: HLA-DR**
**Clone: EPR3692**

**Company:**
**Abcam (ab215985)**

**Tissue: Tonsil**

**Dilution:**
**IHC: 1:100**
**CODEX: 1:50**

**CODEX oligo: 65-Alexa488**

**Antigen: IDO-1**
**Clone: D5J4E**

**Company:**
**Cell Signaling Technology (custom)**

**Tissue: Tonsil**

**Dilution:**
**IHC: 1:5000**
**CODEX: 1:20**

**CODEX oligo: 59-Alexa647**

**Antigen: IRF4**
**Clone: IRF4.3E4**

**Company:**
**Biolegend (646402)**

**Tissue: Plasmacytoma**

**Dilution:**
**IHC: 1:100**
**CODEX: 1:25**

**CODEX oligo: 51-Alexa647**

**Antigen: Ki-67**
**Clone: B56**

**Company:**
**BD Biosciences (556003)**

**Tissue: Tonsil**
**Dilution:**
**IHC: 1:100**
**CODEX: 1:100**

**CODEX oligo: 6-Alexa647**

**IHC** | **CODEX** | **H&E**

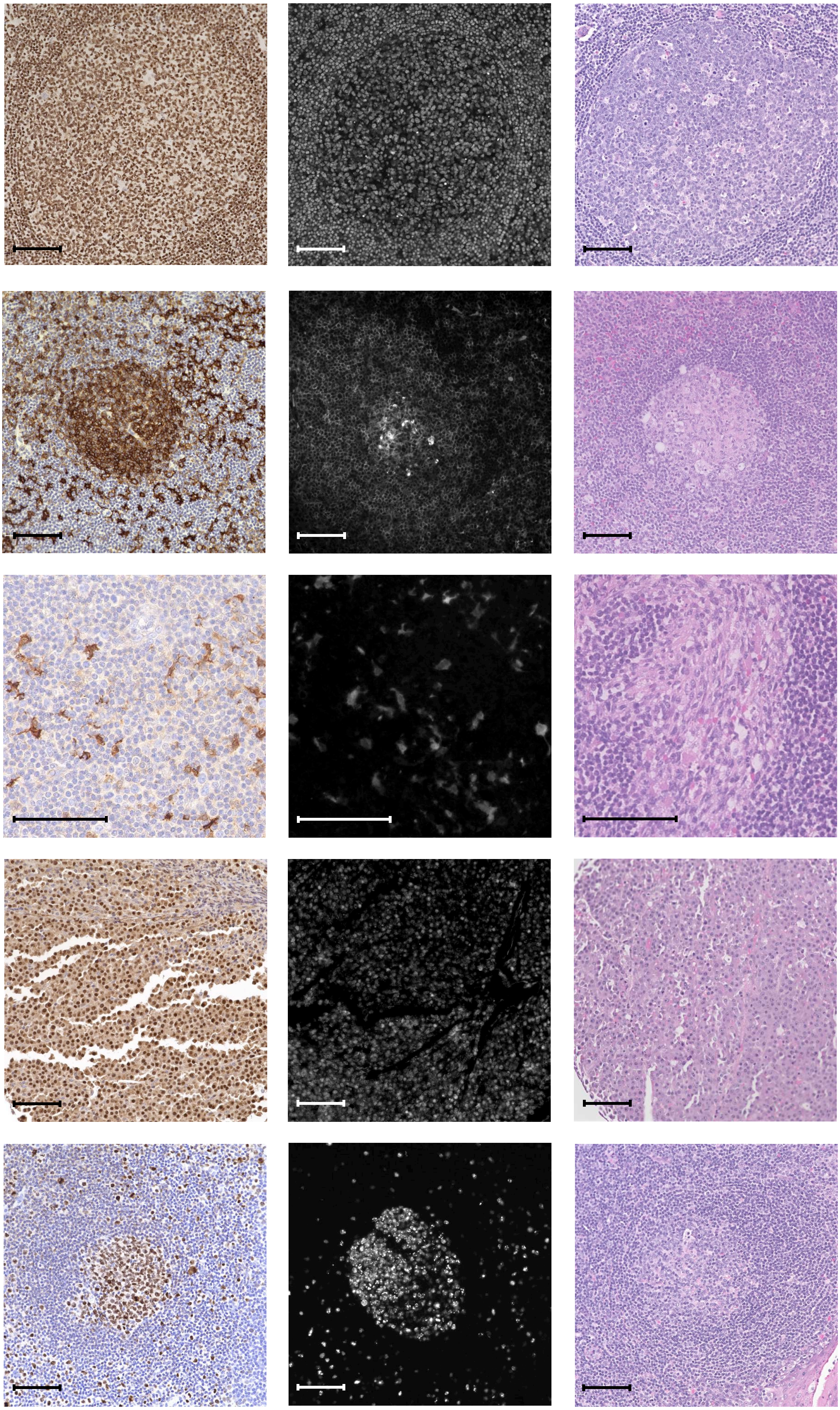

### Figure S1m

#### Staining specifications

**Antigen: Melan-A**
**Clones:**
**A103 + M2-7C10 + M2-9E3**

Company:
Novus Biologicals (NBP2-34546)

Tissue: Malignant melanoma

Dilution:
IHC: 1:250
CODEX: 1:50

CODEX oligo: 44-ATTO550

**Antigen: MMP9**
**Clone: L51/82**

Company:
Biolegend (819701)

Tissue: Bone marrow

Dilution:
IHC: 1:800
CODEX: 1:25

CODEX oligo: 65-Alexa488

**Antigen: MMP12**
**Clone: polyclonal**

Company:
Abcam (ab137444)

Tissue: Tonsil

Dilution:
IHC: 1:200
CODEX: 1:25

CODEX oligo: 80-Alexa647

**Antigen: Muc-1 (EMA)**
**Clone: 955**

Company:
NSJ Bioreagents (V2372SAF)

Tissue: Lung adenocarcinoma

Dilution:
IHC: 1:200
CODEX: 1:100

CODEX oligo: 15-Alexa488

**Antigen: Na-K-ATPase**
**Clone: EP1845Y**

Company:
Abcam (ab167390)

Tissue: Tonsil

Dilution:
IHC: 1:100
CODEX: 1:100

CODEX oligo: 36-Alexa488

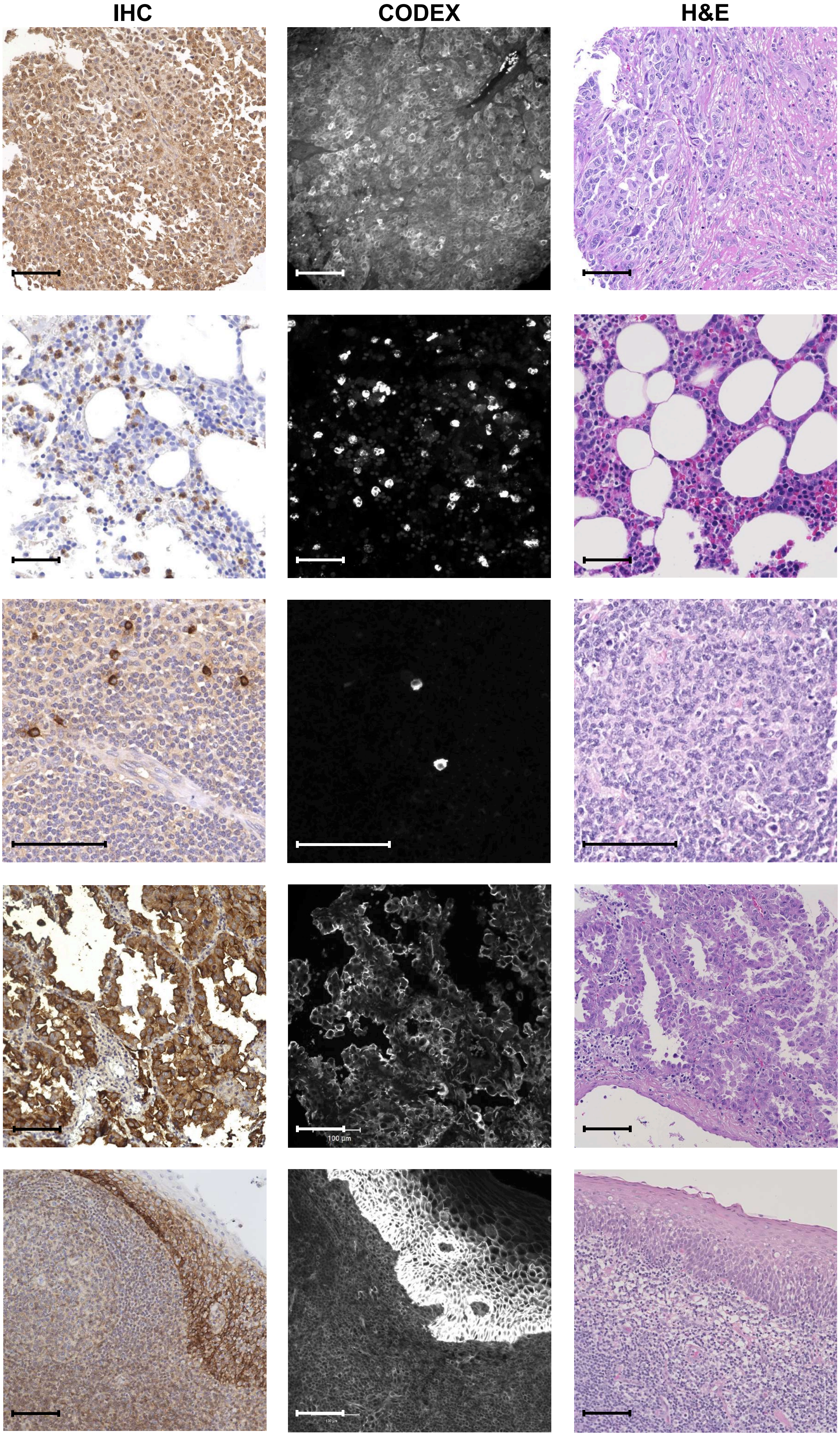

### Figure S1n

**Staining specifications** | **IHC** | **CODEX** | **H&E**

**Antigen: p53**
**Clone: DO-7**

Companies:
Cell Marque (custom)
Santa Cruz Bio (sc-47698)

Tissue: Breast cancer NST

Dilution:
IHC: 1:1000
CODEX: 1:25

CODEX oligo: 52-ATTO550

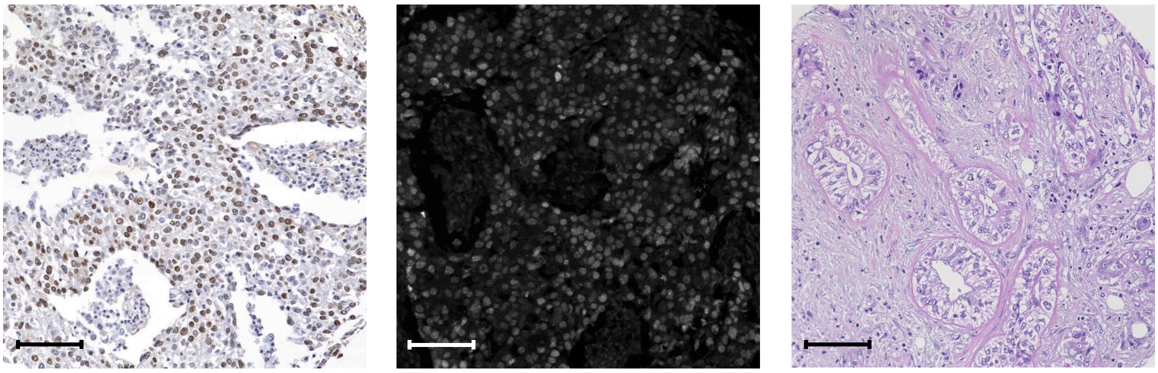

**Antigen: Pan-Cytokeratin**
**Clones: AE-1 + AE-3 or C-11**

Company:
Biolegend (914204 or 628602)

Tissue: Tonsil

Dilution:
IHC: 1:200
CODEX: 1:200

CODEX oligo: 67-ATTO550

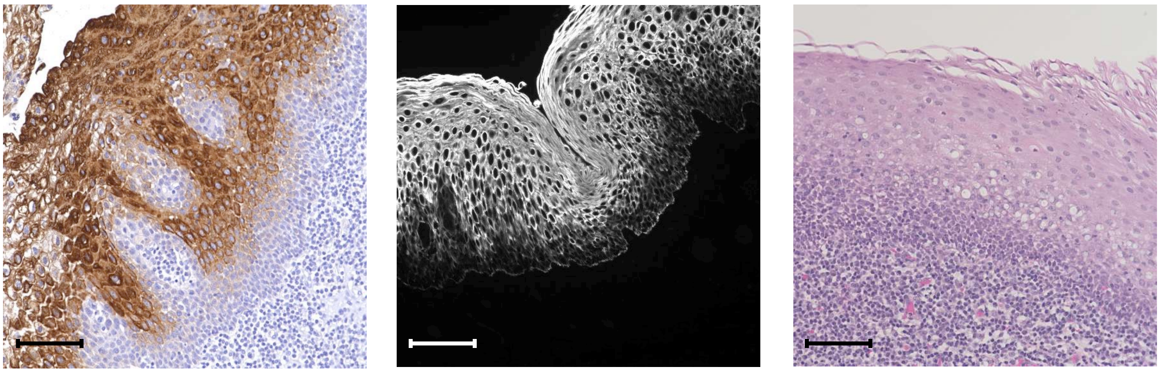

**Antigen: PAX5**
**Clone: D7H5X**

Company:
Cell Signaling Technology (custom)

Tissue: Tonsil

Dilution:
IHC: 1:200
CODEX: 1:25

CODEX oligo: 42-Alexa647

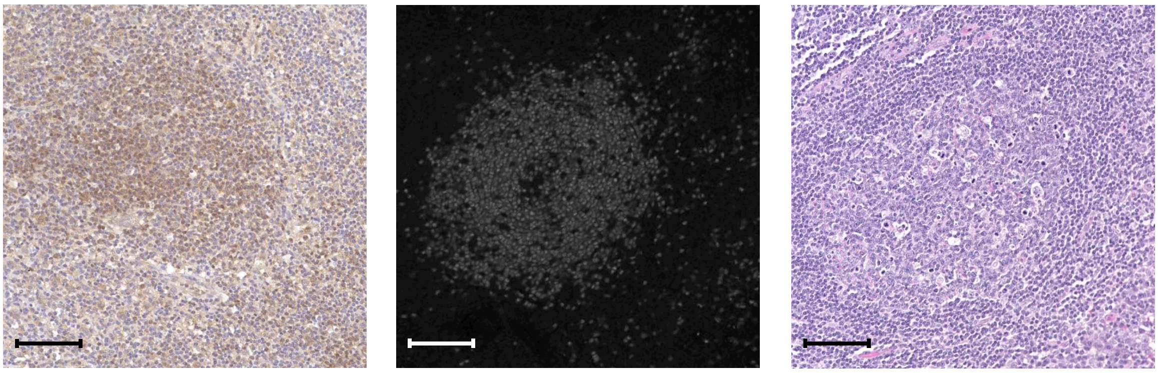

**Antigen: Podoplanin**
**Clone: D2-40 or NC-08**

Company:
Biolegend (916606 or 337002)

Tissue: Tonsil

Dilution:
IHC: 1:200
CODEX: 1:200

CODEX oligo: 32-Alexa647

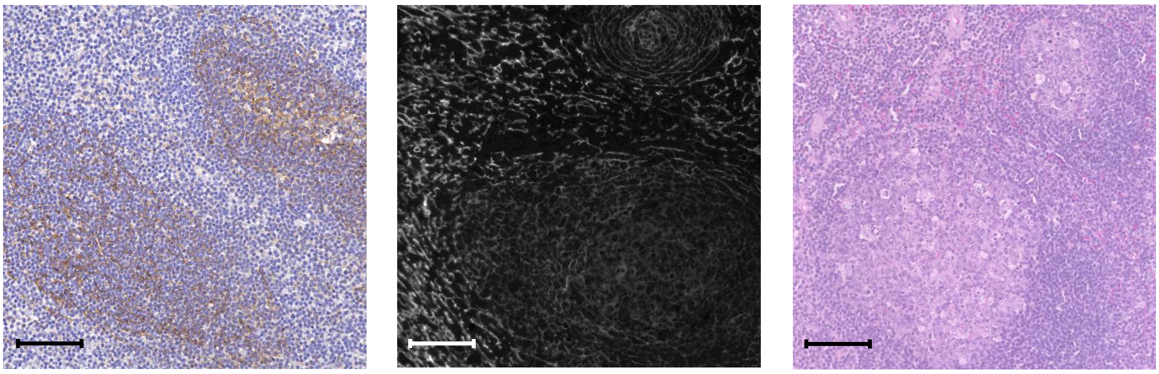

**Antigen: S100A6**
**Clone: 7D11**

Company:
Novus Biologicals (NB100-1765)

Tissue: Malignant melanoma

Dilution:
IHC: 1:100
CODEX: 1:20

CODEX oligo: 20-ATTO550

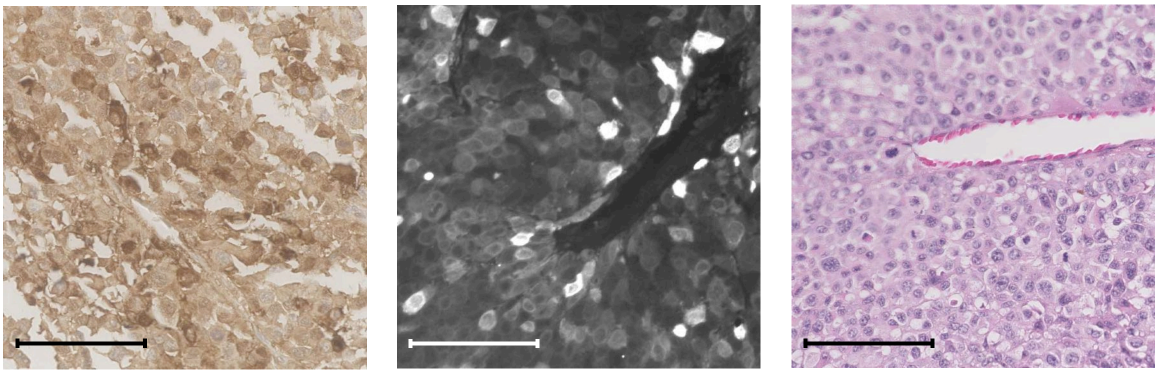

### Figure S1o

**Staining specifications** | **IHC** | **CODEX** | **H&E**

**Antigen: Synaptophysin**
**Clone: 7H12**

**Company:**
**Novus Biologicals (NBP1-47483)**

**Tissue: Pancreas**

**Dilution:**
**IHC: 1:200**
**CODEX: 1:100**

**CODEX oligo: 26-ATTO550**

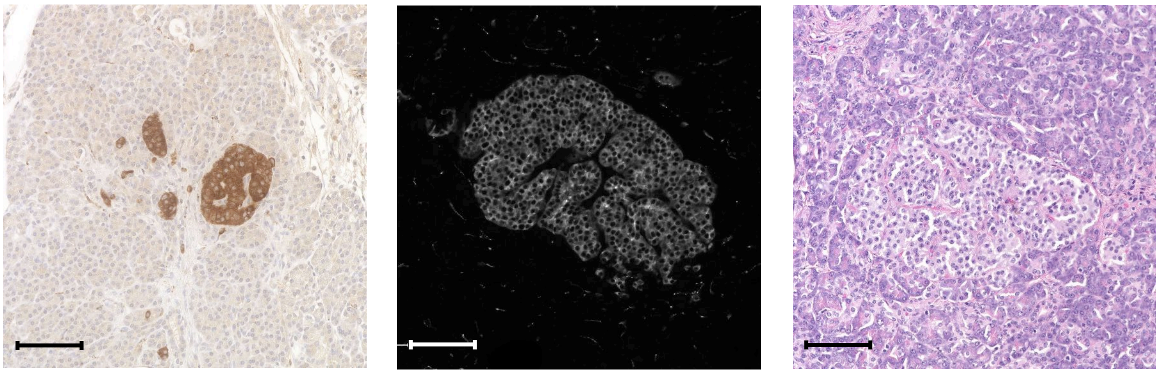

**Antigen: T-bet**
**Clones: D6N8B**

**Company:**
**Cell Signaling Technology (custom)**

**Tissue: Tonsil**

**Dilution:**
**IHC: 1:25**
**CODEX: 1:100**

**CODEX oligo: 5-ATTO550**

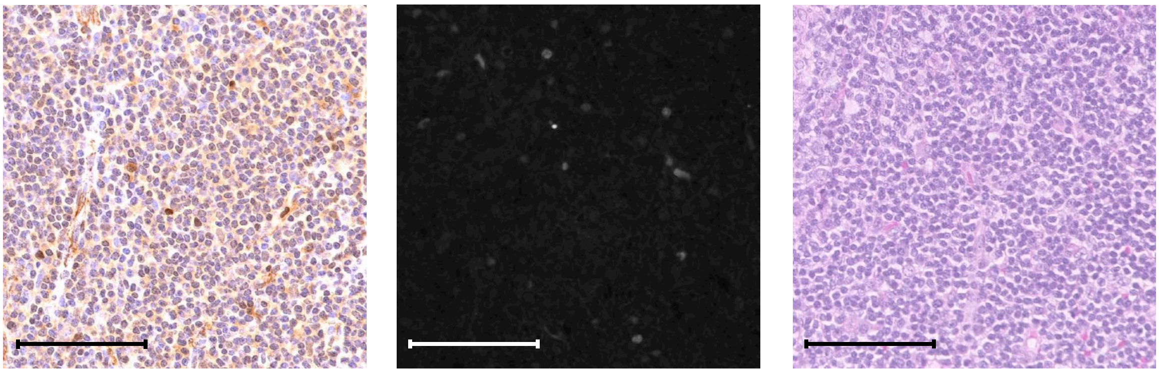

**Antigen: Tubulin β 3**
**Clone: TUJ1**

**Company:**
**Biolegend (801201)**

**Tissue: Pancreas**

**Dilution:**
**IHC: 1:100**
**CODEX: 1:10**

**CODEX oligo: 80-Alexa647**

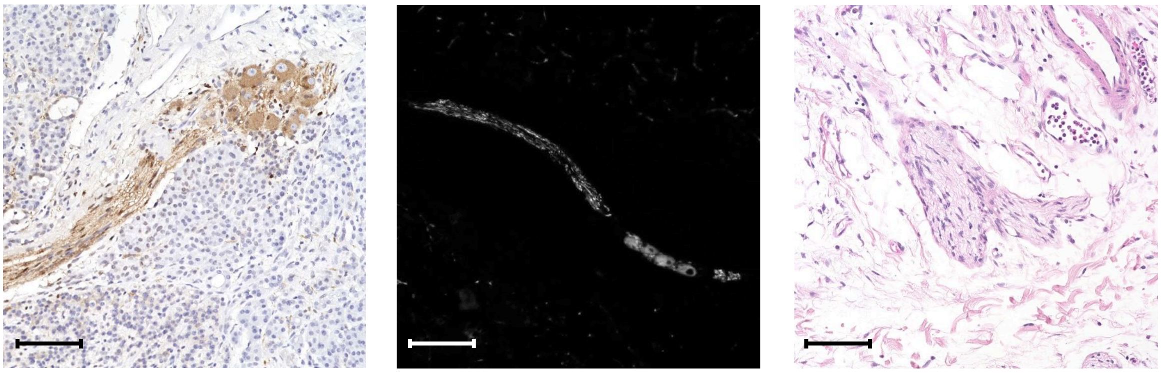

**Antigen: Vimentin**
**Clone: RV202**

**Company:**
**BD Biosciences (550513)**

**Tissue: Tonsil**

**Dilution:**
**IHC: 1:300**
**CODEX: 1:200**

**CODEX oligo: 7-Alexa488**

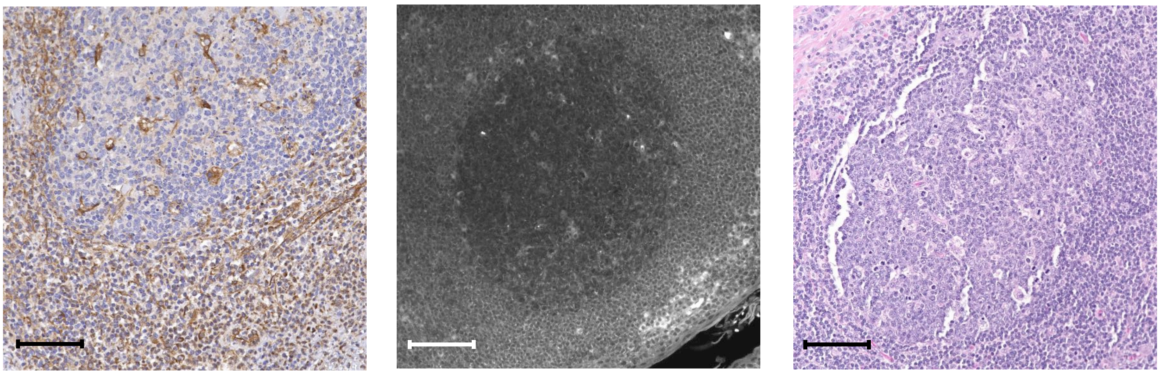

**Antigen: VISTA**
**Clone: D1L2G**

**Company:**
**Cell Signaling Technology (custom)**

**Tissue: Tonsil**

**Dilution:**
**IHC: 1:200**
**CODEX: 1:50**

**CODEX oligo: 79-Alexa647**

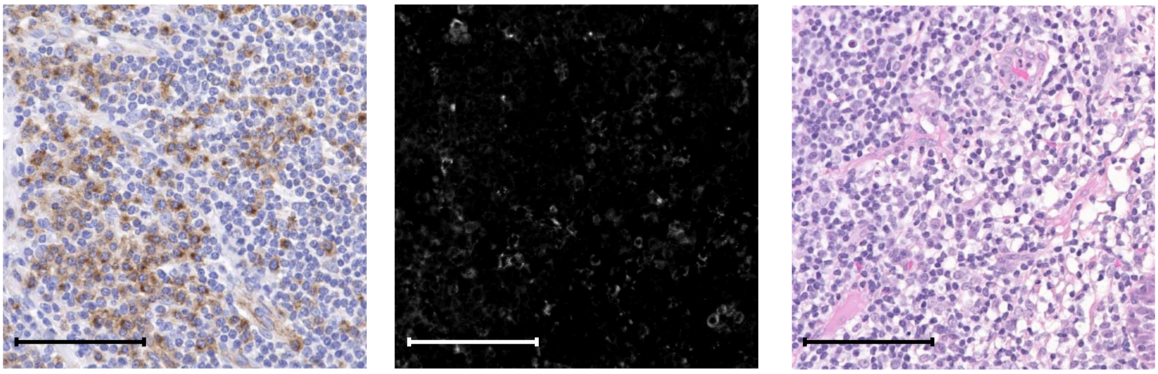

### Figure S2

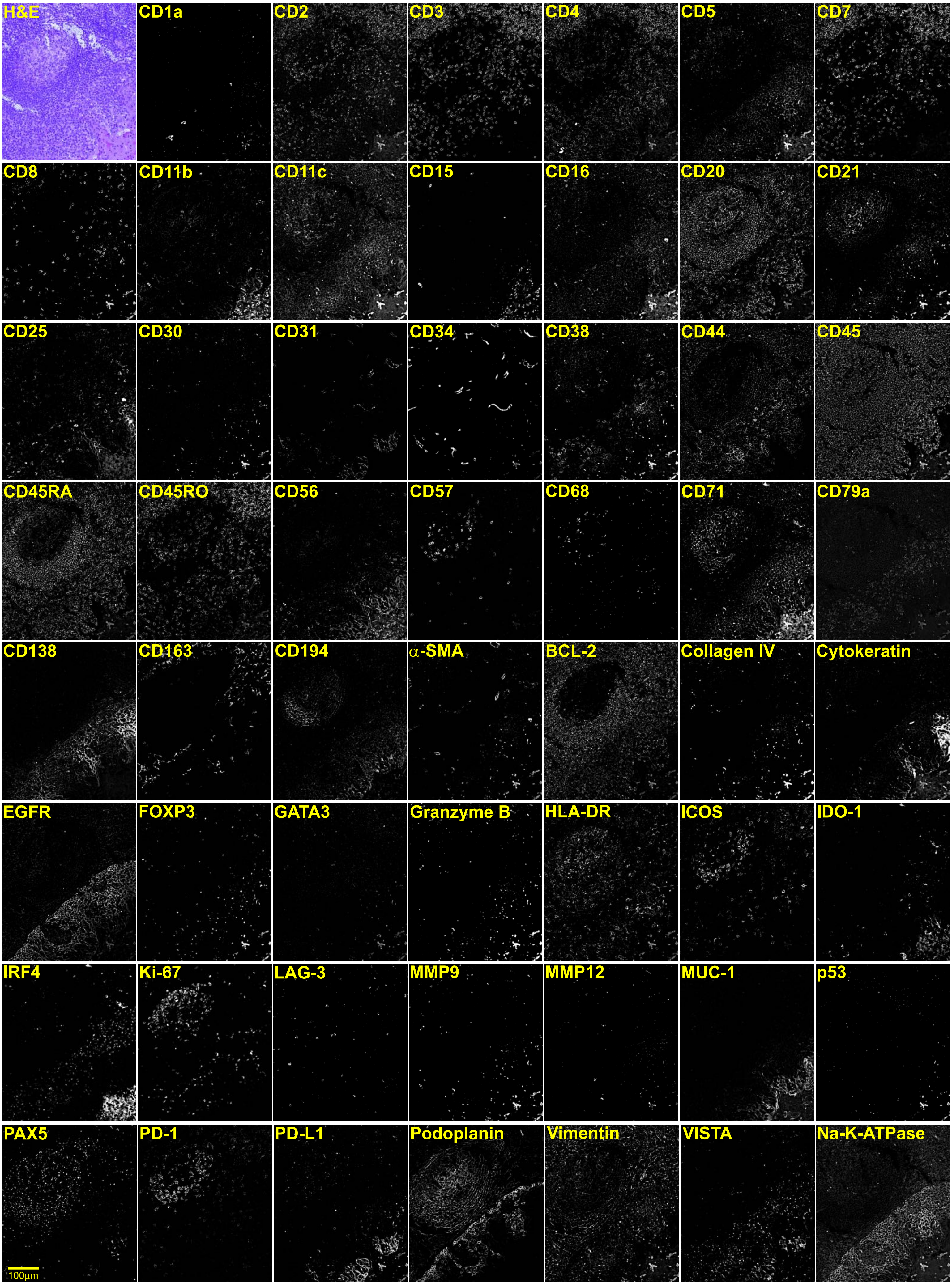
H&E
CD1a
CD2
CD3
CD4
CD5
CD7
CD8
CD11b
CD11c
CD15
CD16
CD20
CD21
CD25
CD30
CD31
CD34
CD38
CD44
CD45
CD45RA
CD45RO
CD56
CD57
CD68
CD71
CD79a
CD138
CD163
CD194
α-SMA
BCL-2
Collagen IV
Cytokeratin
EGFR
FOXP3
GATA3
Granzyme B
HLA-DR
ICOS
IDO-1
IRF4
Ki-67
LAG-3
MMP9
MMP12
MUC-1
p53
PAX5
PD-1
PD-L1
Podoplanin
Vimentin
VISTA
Na-K-ATPase
100μm

### Figure S3

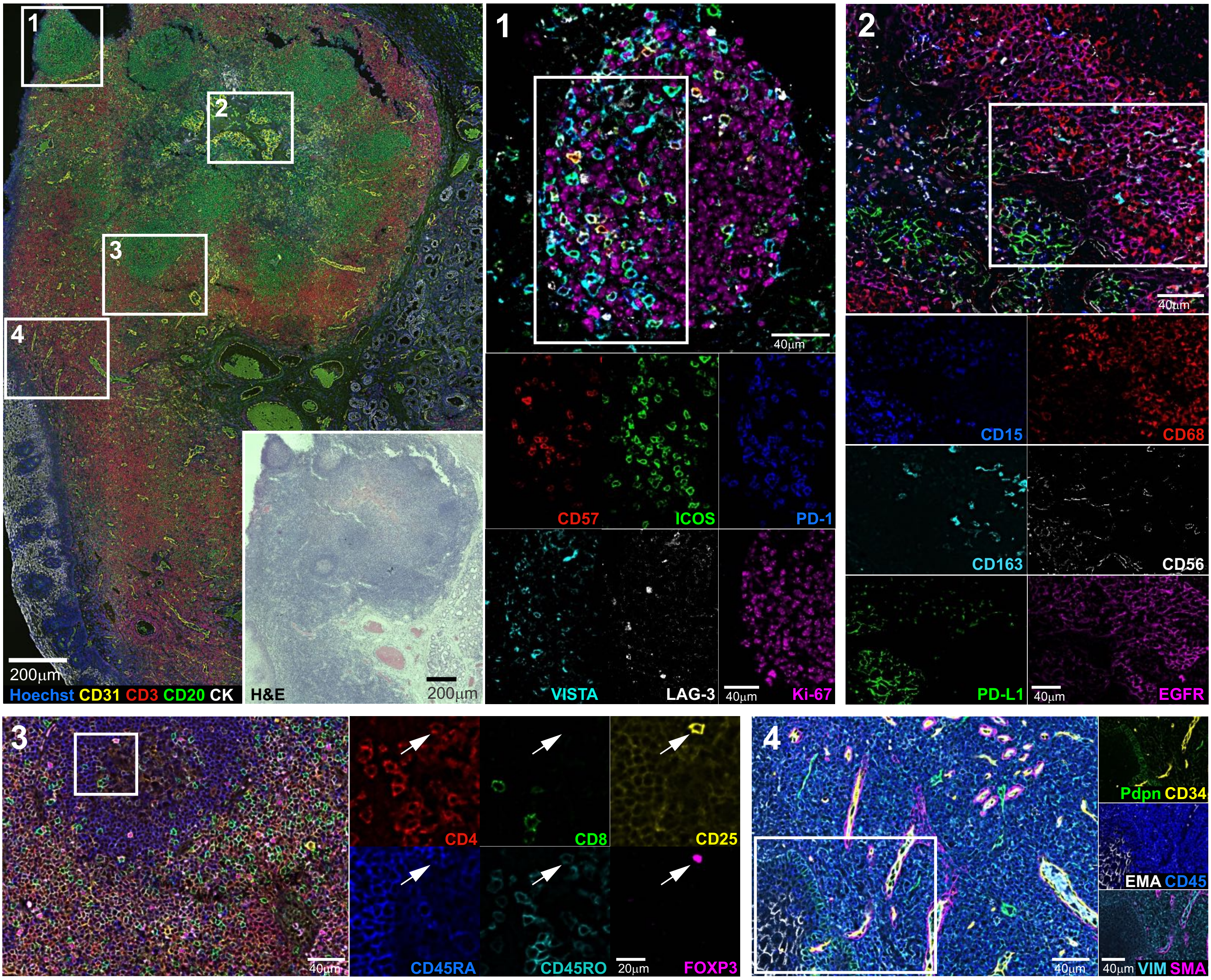
1
2
3
4
200µm
Hoechst CD31 CD3 CD20 CK
H&E
200µm
40µm
CD57
ICOS
PD-1
VISTA
LAG-3
40µm
Ki-67
40µm
CD15
CD68
CD163
CD56
PD-L1
40µm
EGFR
40µm
CD4
CD8
CD25
CD45RA
CD45RO
20µm
FOXP3
40µm
Pdpn CD34
EMA CD45
40µm VIM SMA

### Figure S4

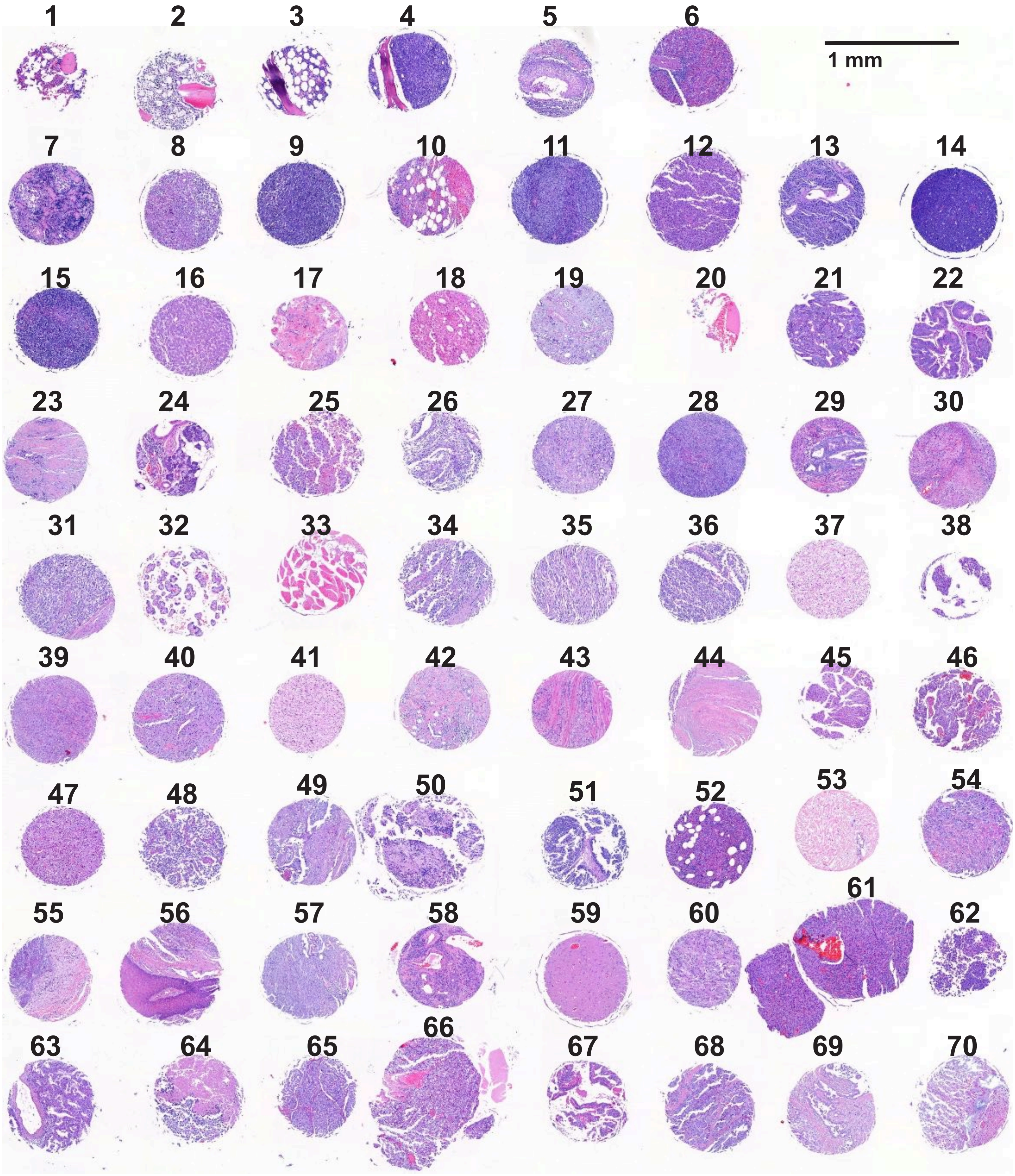
1 mm

### Figure S5

**A**

| Cycles | Channel 1 | Alexa488 | ATTO550 | Alexa647 |
|---|---|---|---|---|
| 1, 11, 21, 31, 32, 33 | HOECHST | - | - | - |
| 2, 5, 8, 12, 15, 18, 22, 25, 28 | HOECHST | HLA-DR | CD45 | FOXP3 |
| 3, 6, 9, 13, 16, 19, 23, 26, 29 | HOECHST | Na-K-ATPase | Pan-Cytokeratin | p53 |
| 4, 7, 10, 14, 17, 20, 24, 27, 30 | HOECHST | Ki-67 | CD20 | CD3 |

↻ repeat

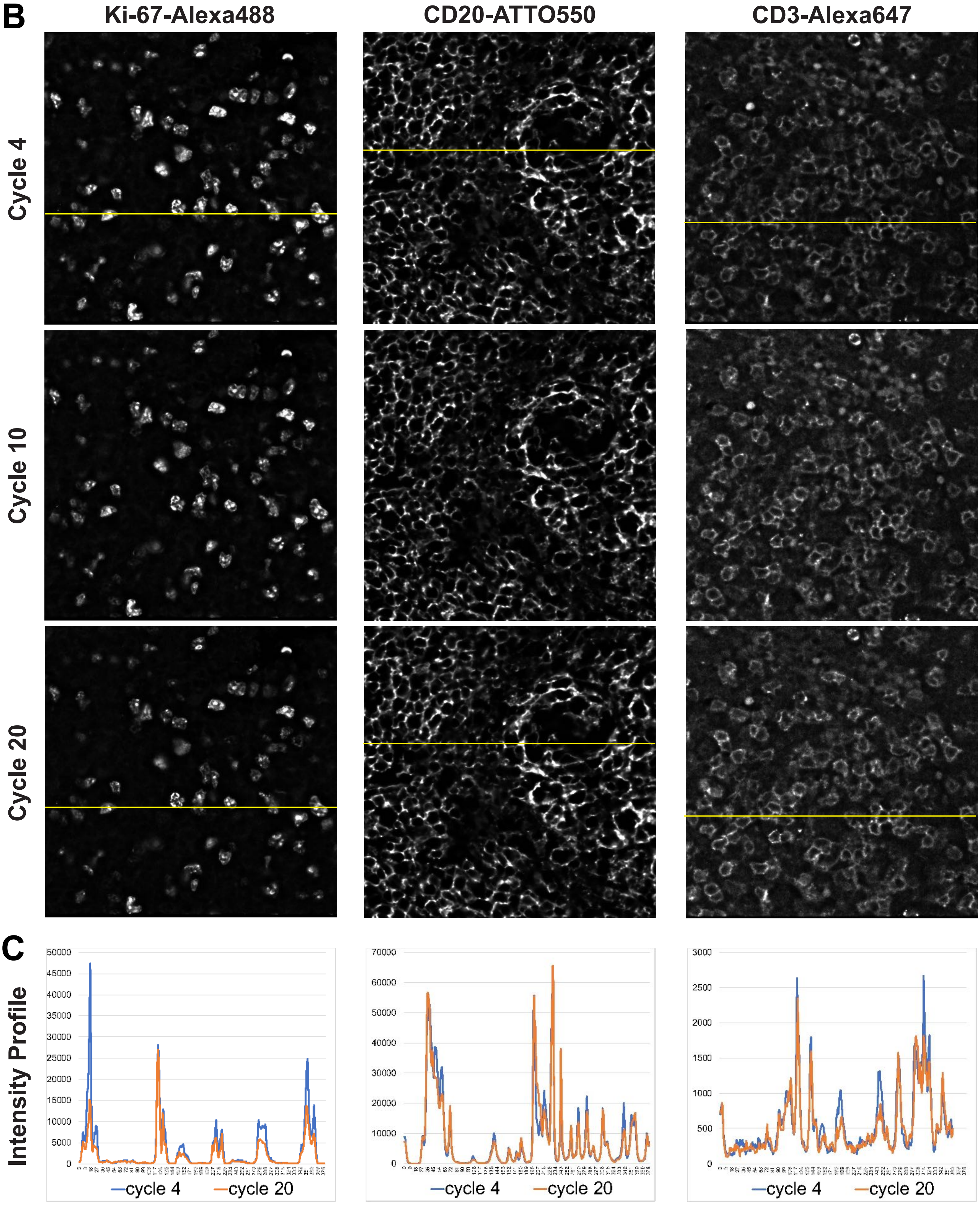

### Figure S6

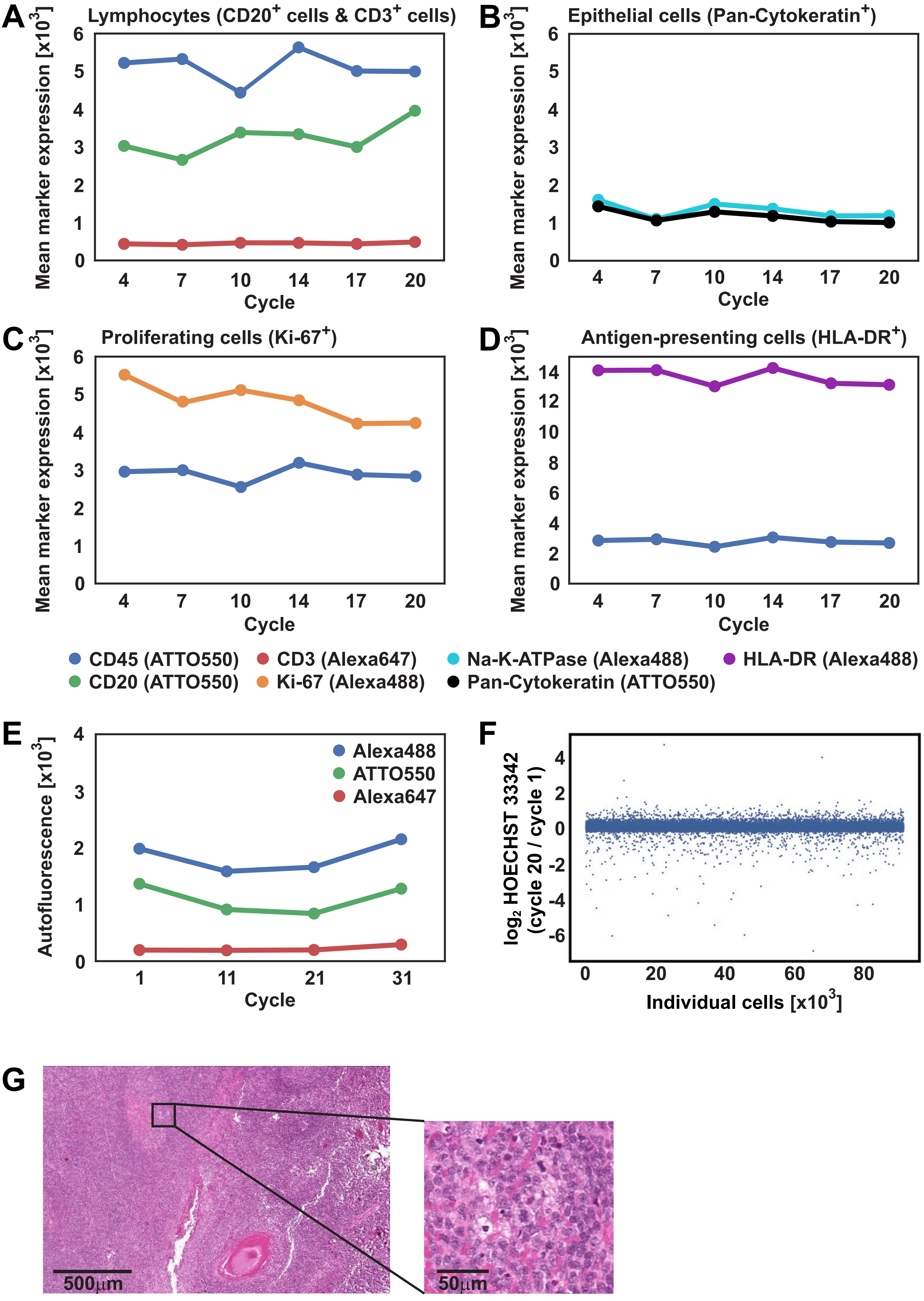

A
Lymphocytes (CD20+ cells & CD3+ cells)
Mean marker expression [x10^3]
Cycle
B
Epithelial cells (Pan-Cytokeratin+)
Mean marker expression [x10^3]
Cycle
C
Proliferating cells (Ki-67+)
Mean marker expression [x10^3]
Cycle
D
Antigen-presenting cells (HLA-DR+)
Mean marker expression [x10^3]
Cycle
CD45 (ATTO550)
CD3 (Alexa647)
Na-K-ATPase (Alexa488)
HLA-DR (Alexa488)
CD20 (ATTO550)
Ki-67 (Alexa488)
Pan-Cytokeratin (ATTO550)
E
Autofluorescence [x10^3]
Alexa488
ATTO550
Alexa647
Cycle
F
log2 HOECHST 33342 (cycle 20 / cycle 1)
Individual cells [x10^3]
G
500µm
50µm

### Figure S7

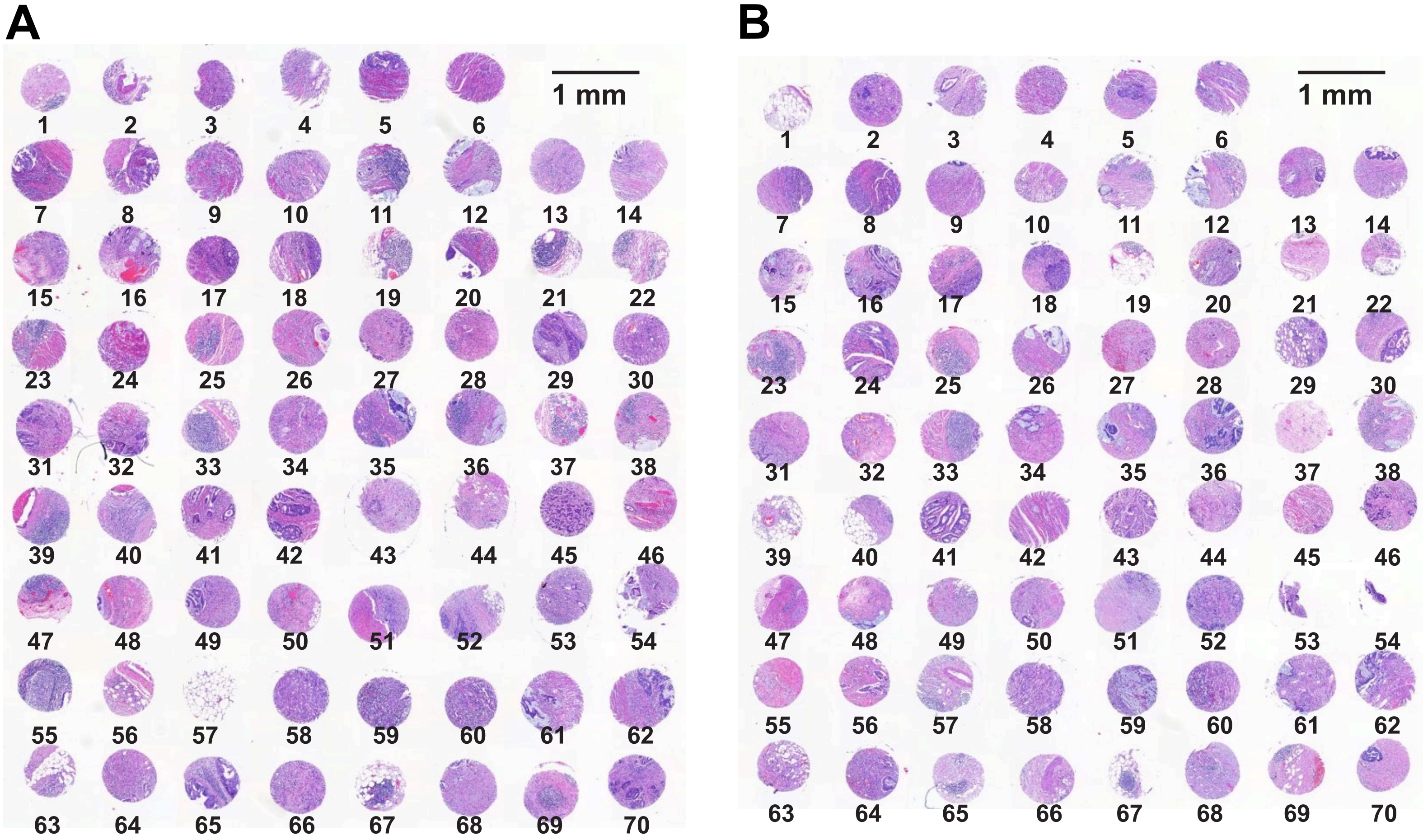

### Figure S8

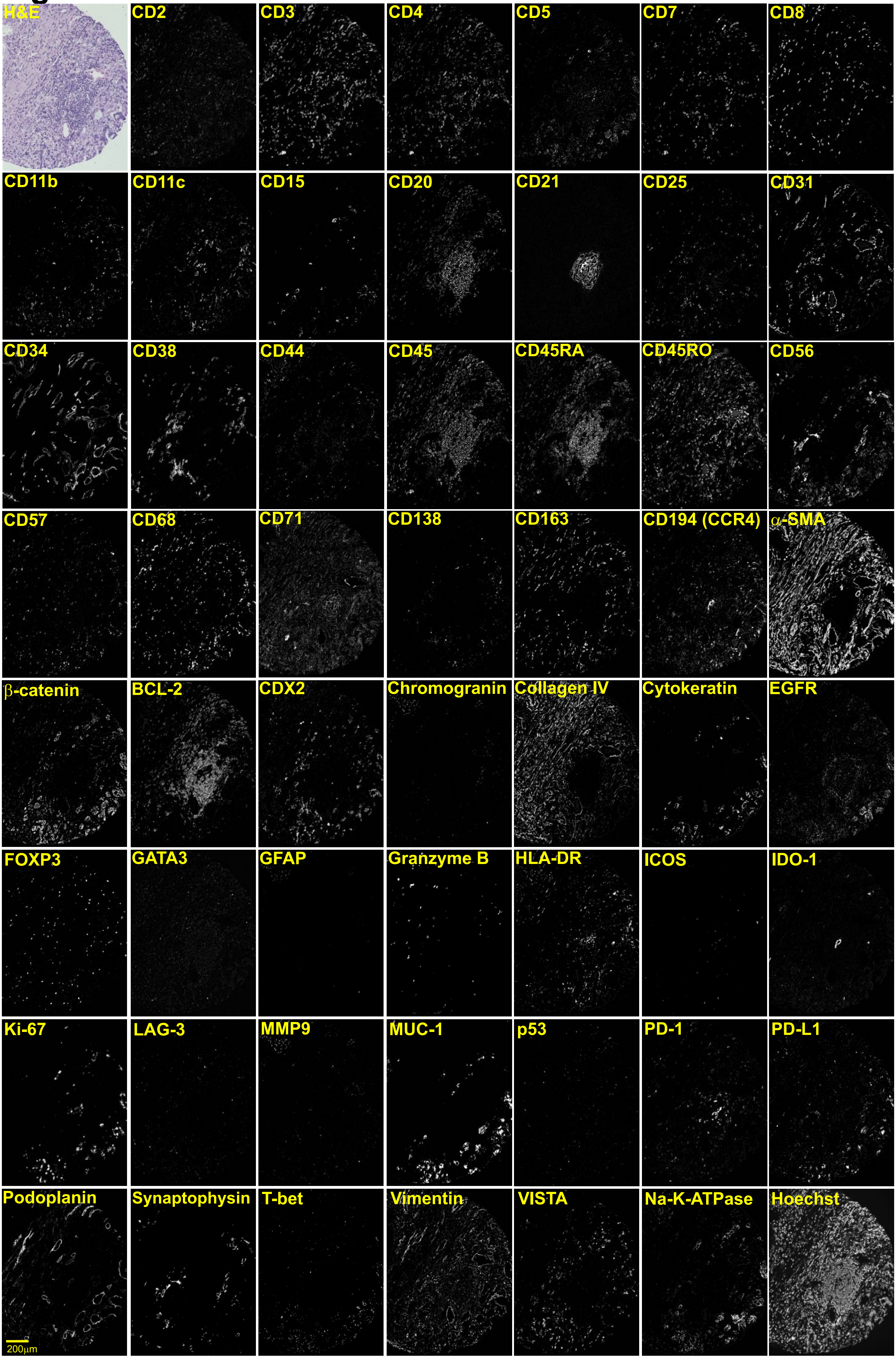
H&E
CD2
CD3
CD4
CD5
CD7
CD8
CD11b
CD11c
CD15
CD20
CD21
CD25
CD31
CD34
CD38
CD44
CD45
CD45RA
CD45RO
CD56
CD57
CD68
CD71
CD138
CD163
CD194 (CCR4)
α-SMA
β-catenin
BCL-2
CDX2
Chromogranin
Collagen IV
Cytokeratin
EGFR
FOXP3
GATA3
GFAP
Granzyme B
HLA-DR
ICOS
IDO-1
Ki-67
LAG-3
MMP9
MUC-1
p53
PD-1
PD-L1
Podoplanin
Synaptophysin
T-bet
Vimentin
VISTA
Na-K-ATPase
Hoechst
200μm

### Figure S9

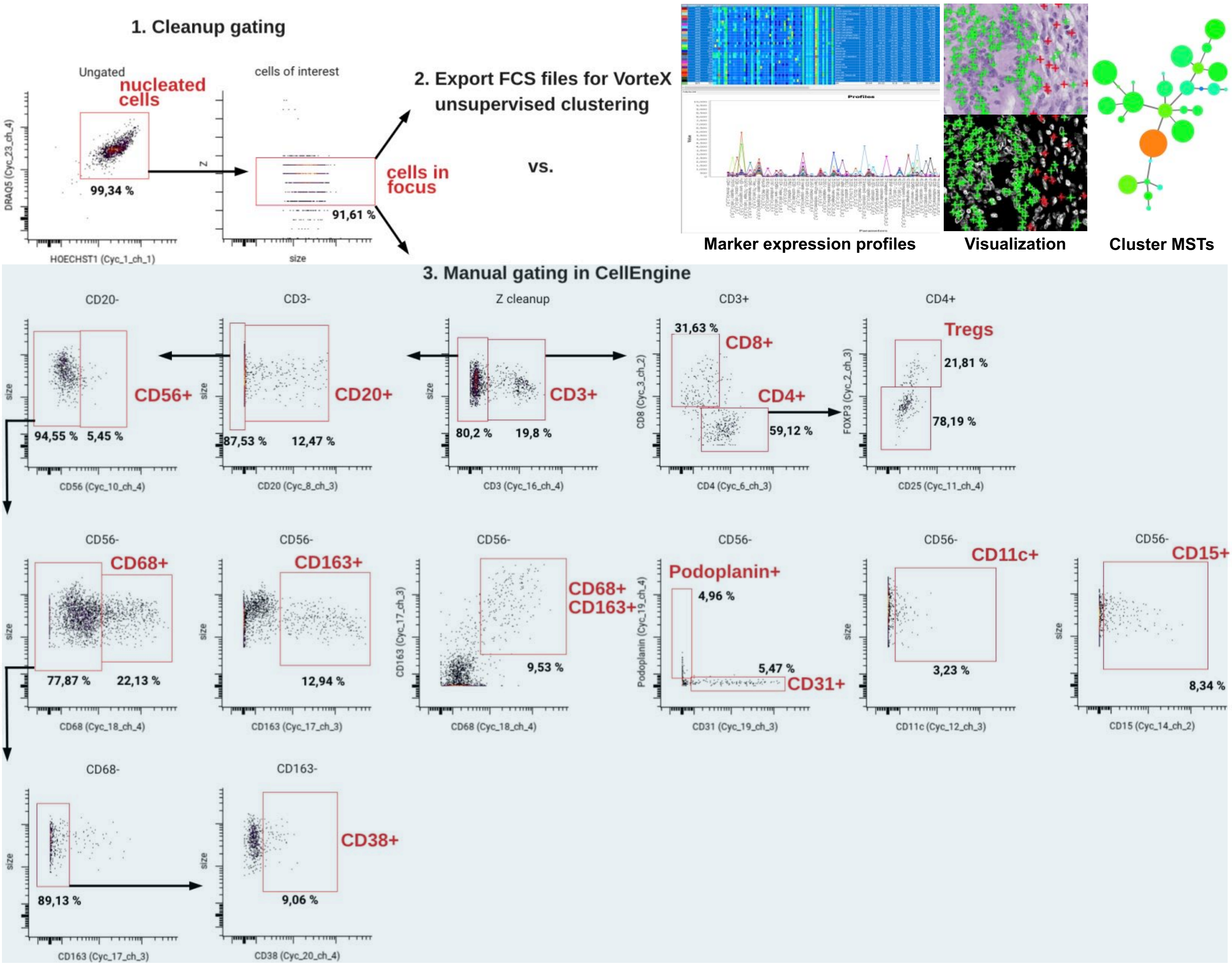
1. Cleanup gating
Ungated
nucleated cells
99,34 %
DRAQ5 (Cyc_23_ch_4)
HOECHST1 (Cyc_1_ch_1)
cells of interest
cells in focus
91,61 %
Z
size
2. Export FCS files for VorteX unsupervised clustering
vs.
Marker expression profiles
Visualization
Cluster MSTs
3. Manual gating in CellEngine
CD20-
CD56+
94,55 %
5,45 %
CD56 (Cyc_10_ch_4)
CD3-
CD20+
87,53 %
12,47 %
CD20 (Cyc_8_ch_3)
Z cleanup
CD3+
80,2 %
19,8 %
CD3 (Cyc_16_ch_4)
CD3+
31,63 %
CD8+
CD4+
59,12 %
CD8 (Cyc_3_ch_2)
CD4 (Cyc_6_ch_3)
CD4+
Tregs
21,81 %
78,19 %
FOXP3 (Cyc_2_ch_3)
CD25 (Cyc_11_ch_4)
CD56-
CD68+
77,87 %
22,13 %
CD68 (Cyc_18_ch_4)
CD56-
CD163+
12,94 %
CD163 (Cyc_17_ch_3)
CD56-
CD68+ CD163+
9,53 %
CD163 (Cyc_17_ch_3)
CD68 (Cyc_18_ch_4)
CD56-
Podoplanin+
4,96 %
5,47 %
CD31+
Podoplanin (Cyc_19_ch_4)
CD31 (Cyc_19_ch_3)
CD56-
CD11c+
3,23 %
CD11c (Cyc_12_ch_3)
CD56-
CD15+
8,34 %
CD15 (Cyc_14_ch_2)
CD68-
89,13 %
CD163 (Cyc_17_ch_3)
CD163-
CD38+
9,06 %
CD38 (Cyc_20_ch_4)
size

### Figure S10

A. Tumor cell cluster

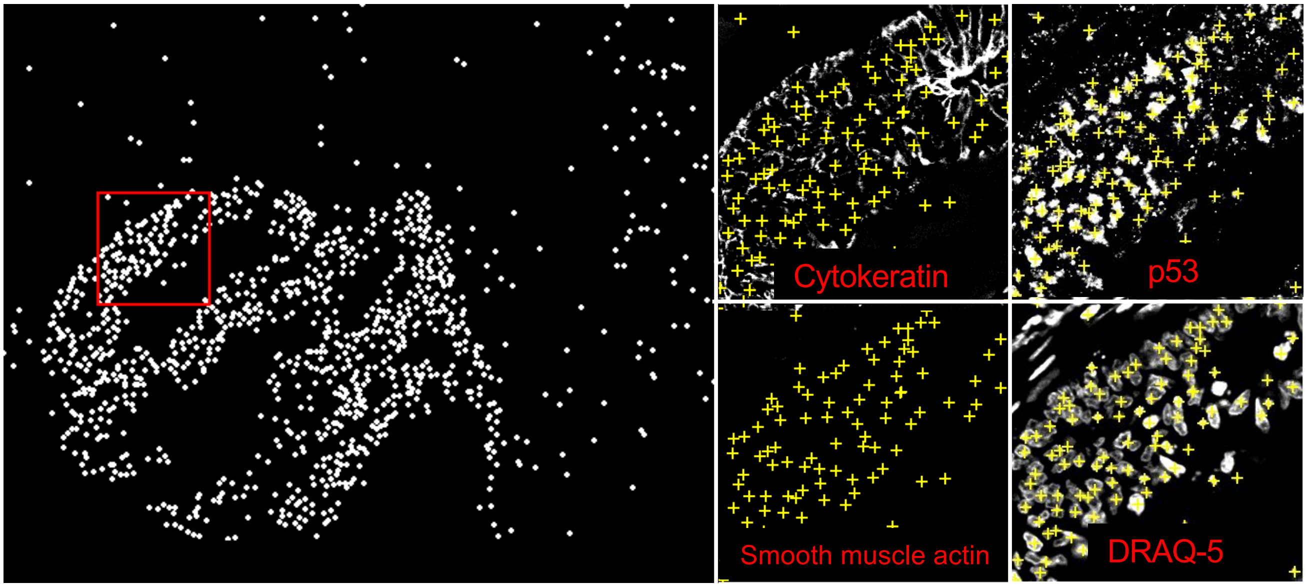

B. CD68+CD163+ macrophage cluster

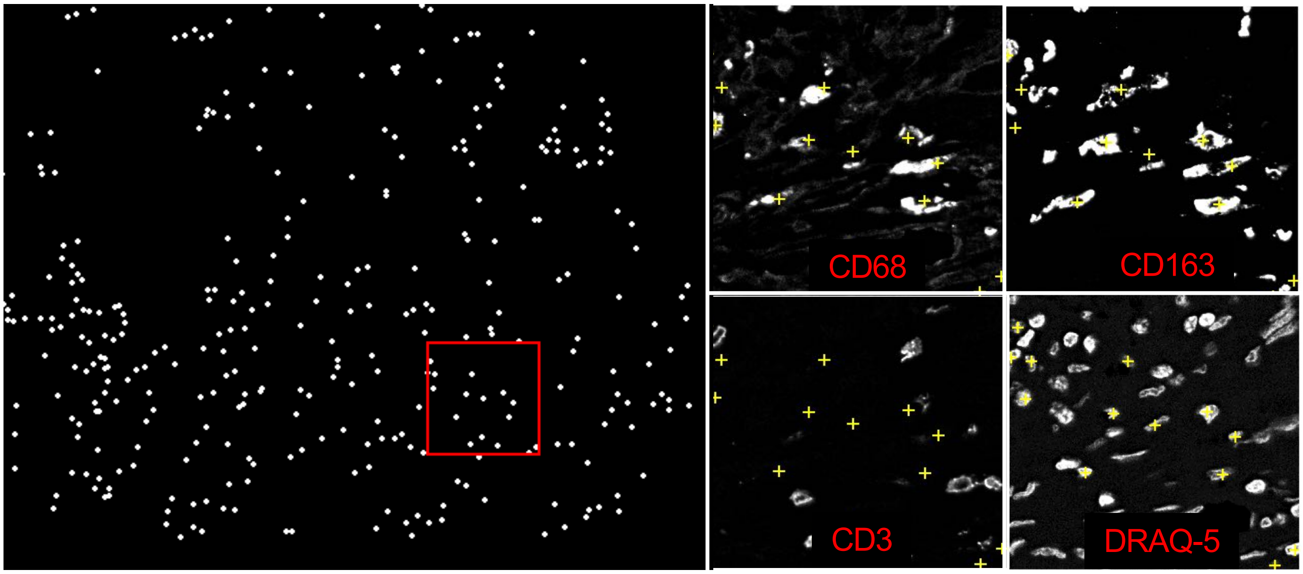

C. Smooth muscle cluster

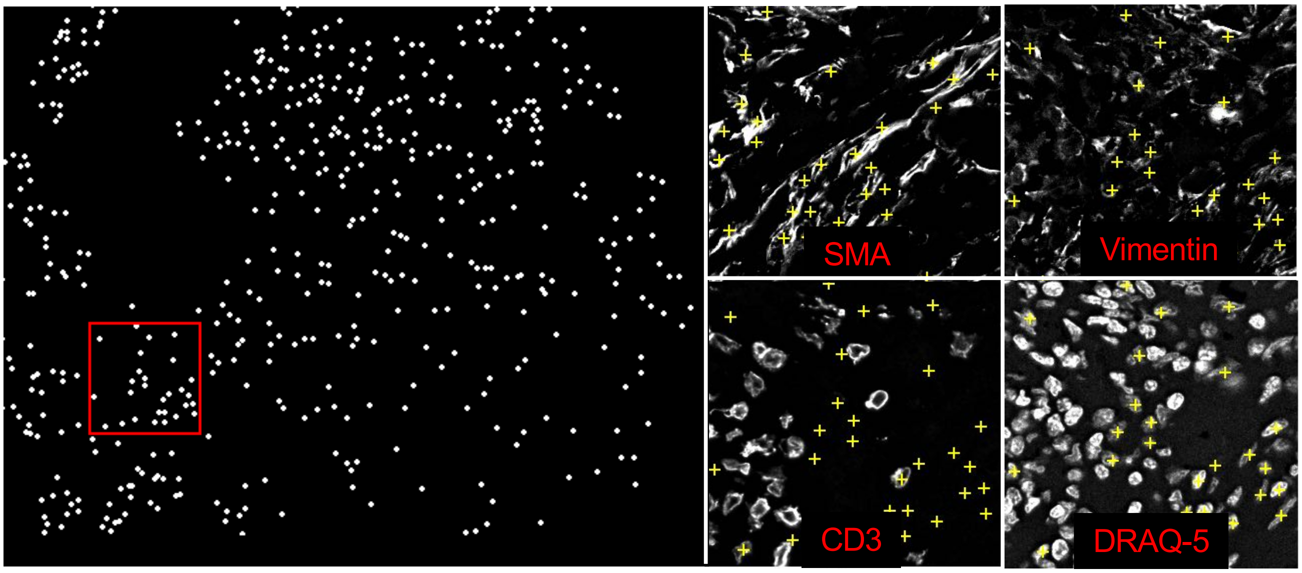

D. Granulocyte cluster

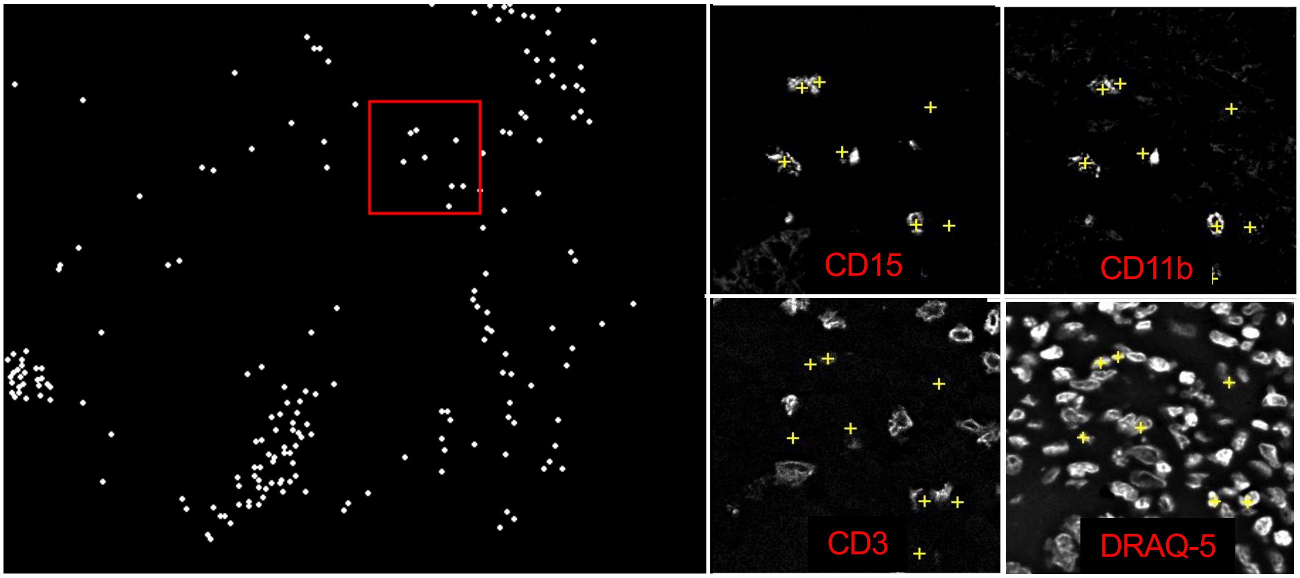

E. Stroma cluster

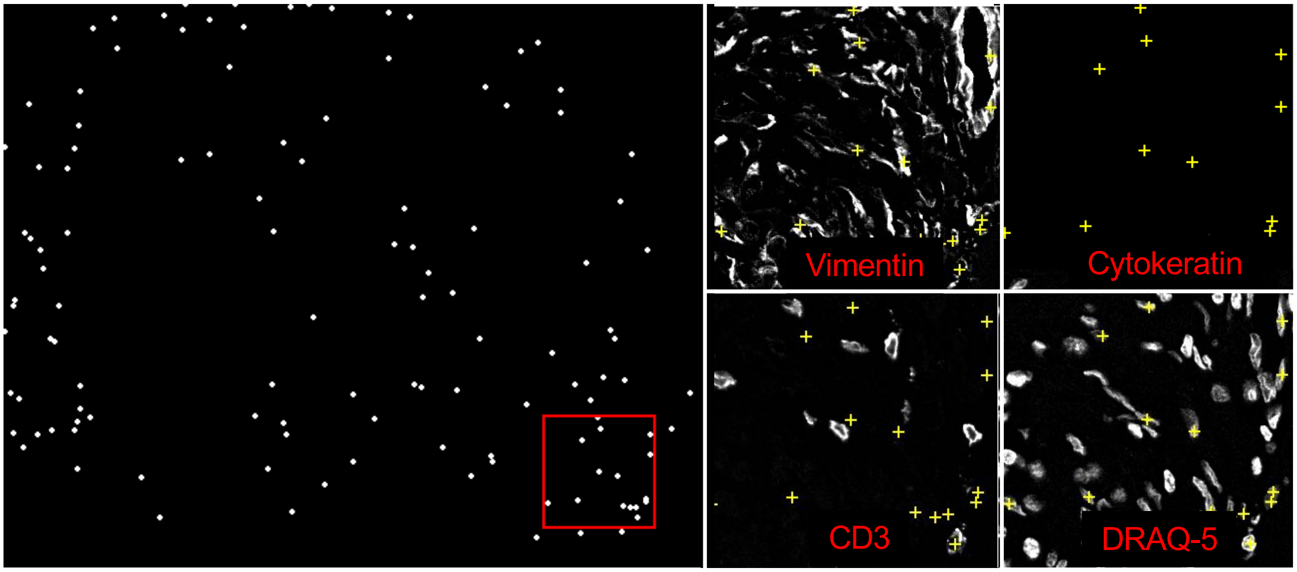

F. CD4+CD45RO+ T cell cluster

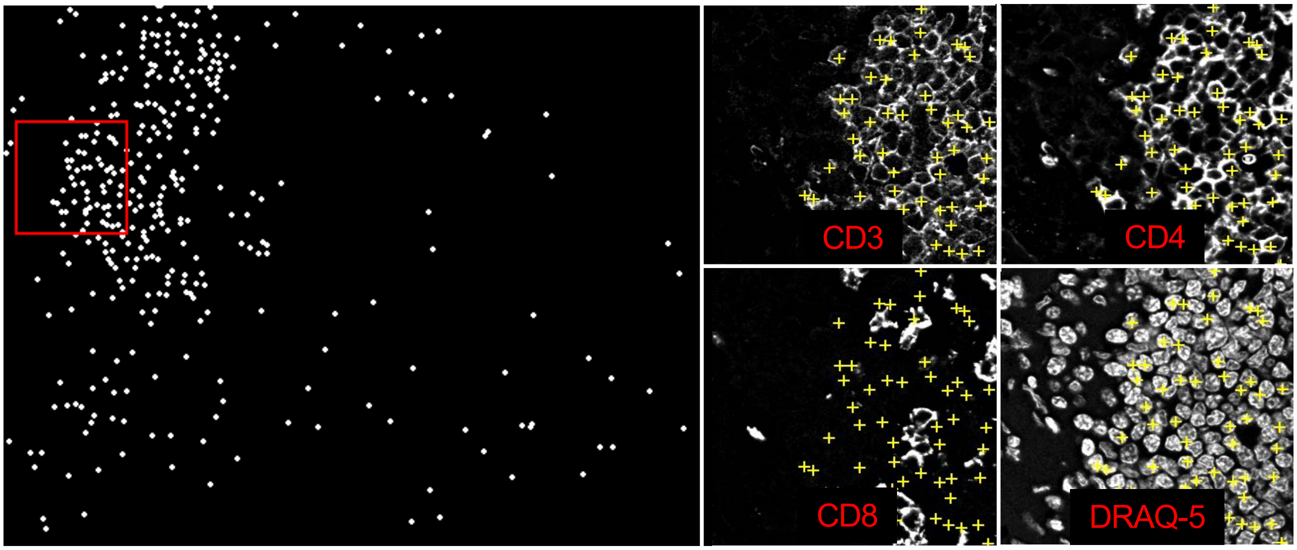

G. CD8+ T cell cluster

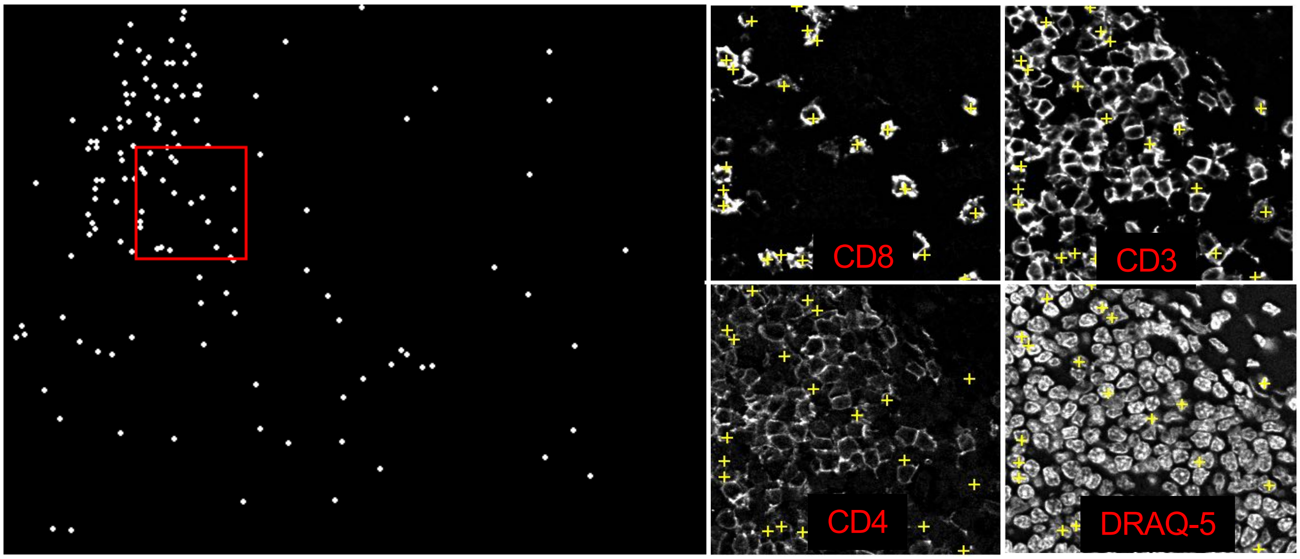

H. B cell cluster

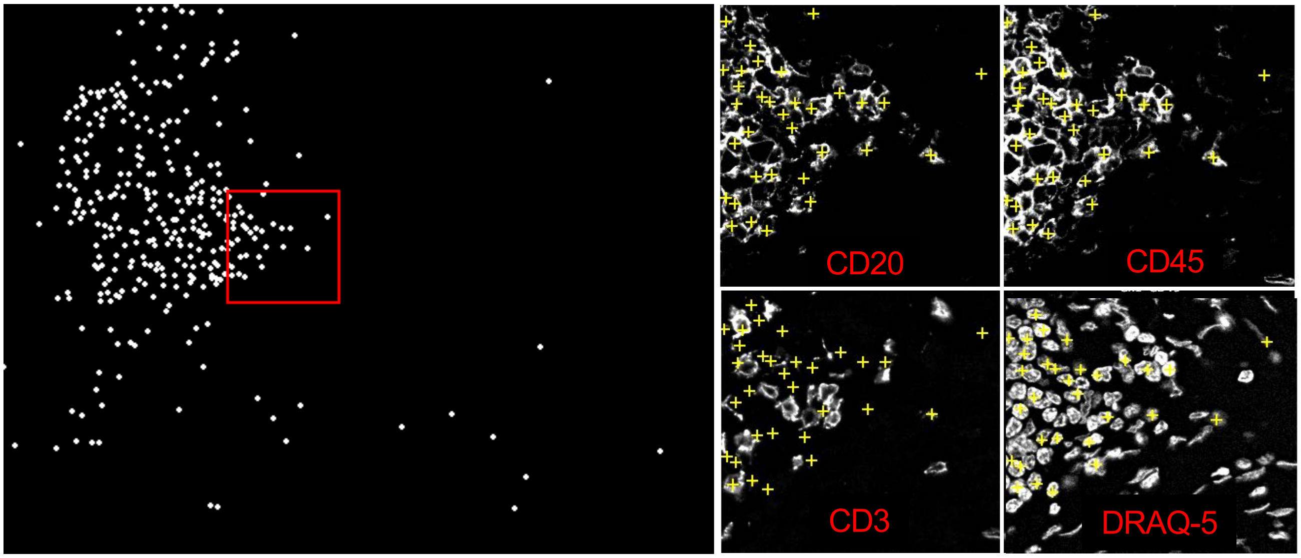

I. Vasculature cluster

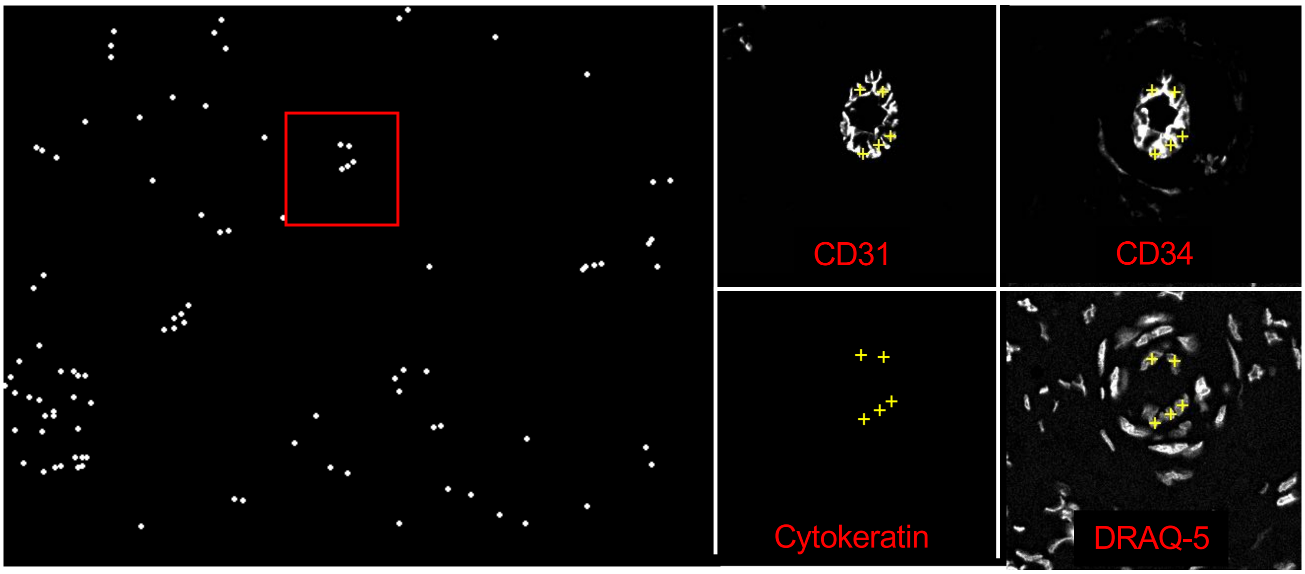

J. Plasma cell cluster

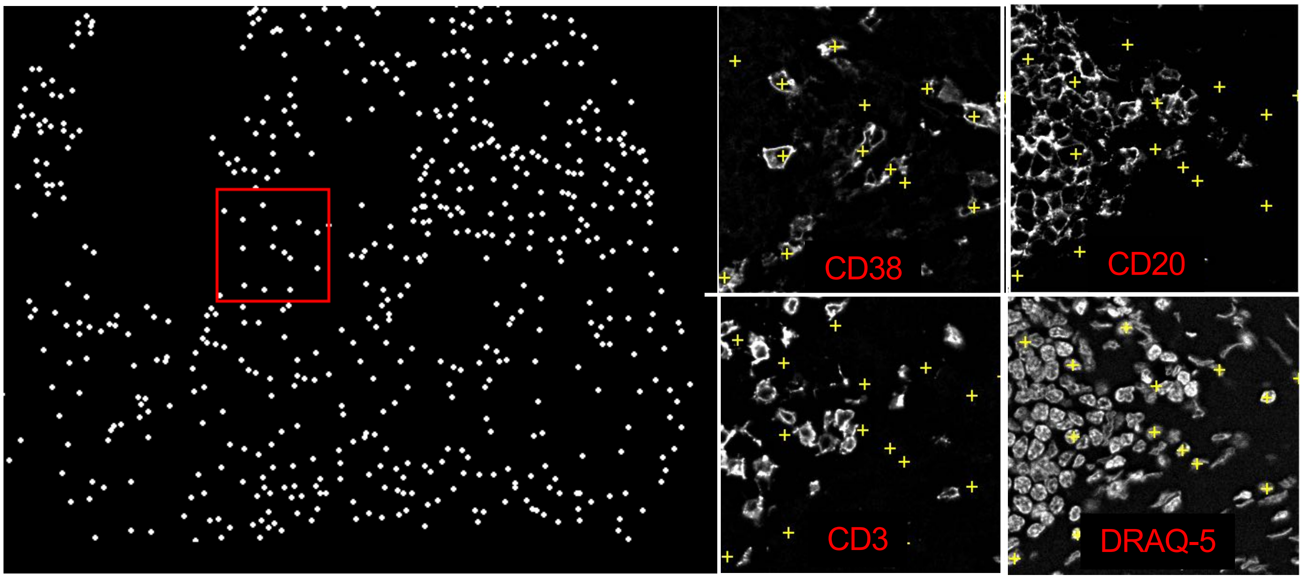

K. Treg cluster

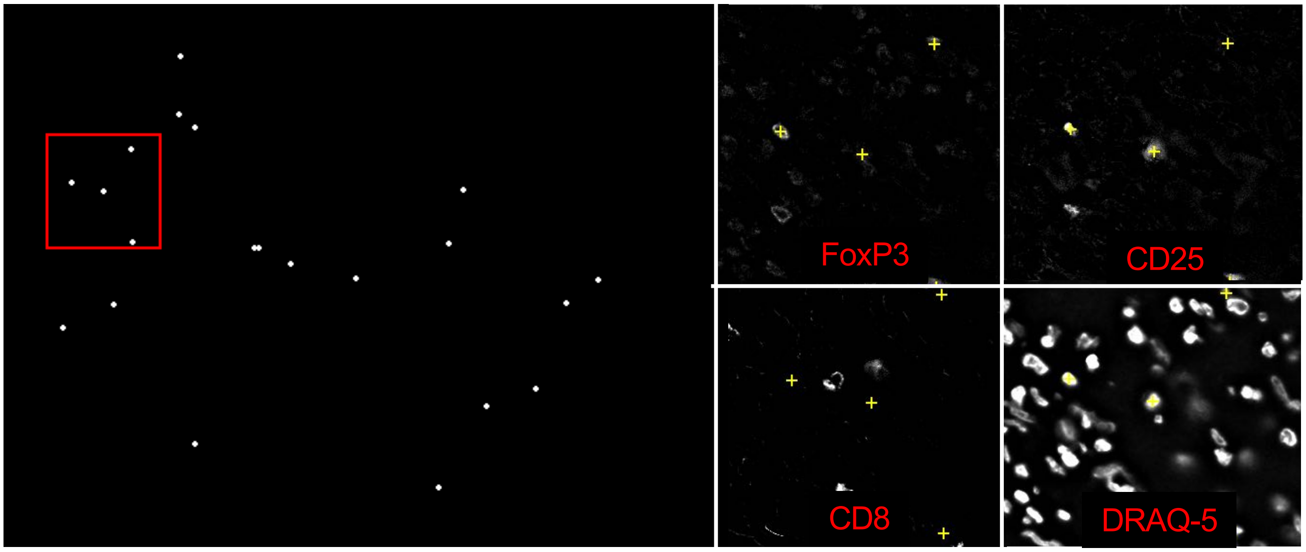

L. CD4+ T cell cluster

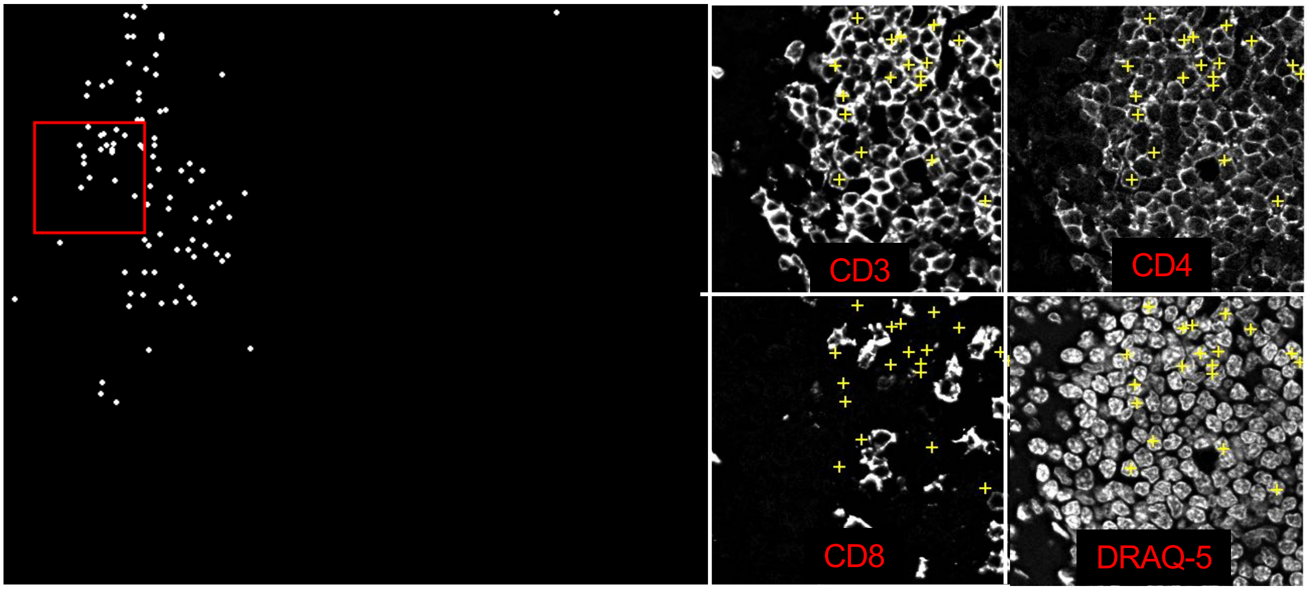

M. Adipocyte cluster

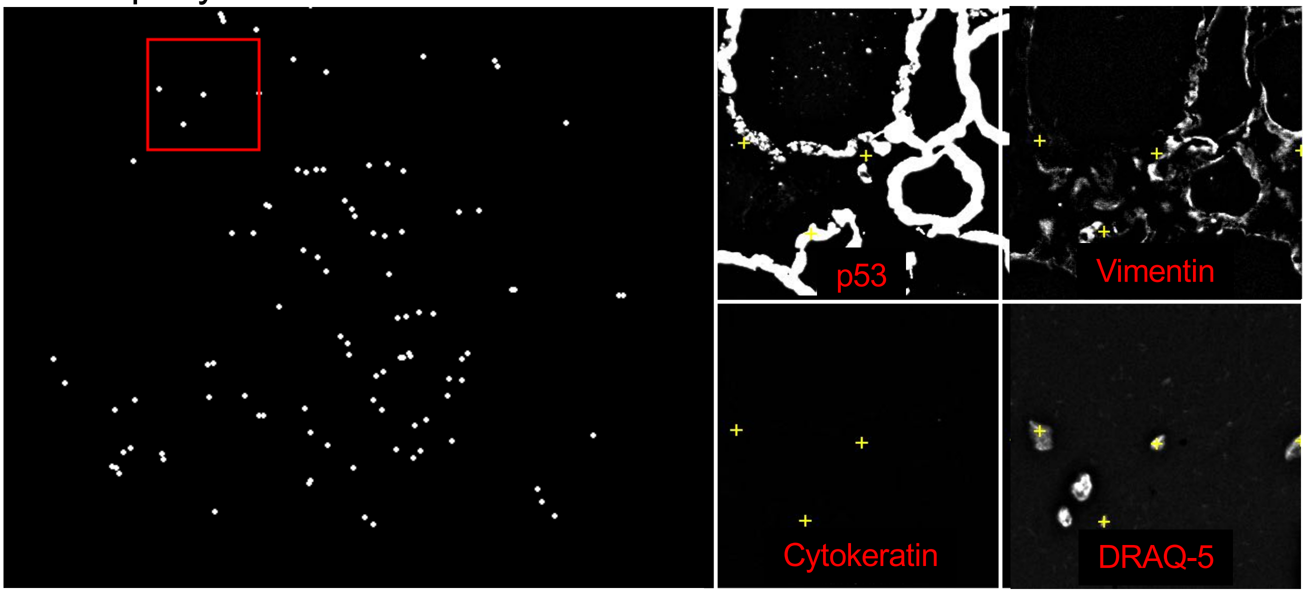

N. CD68+ macrophage cluster

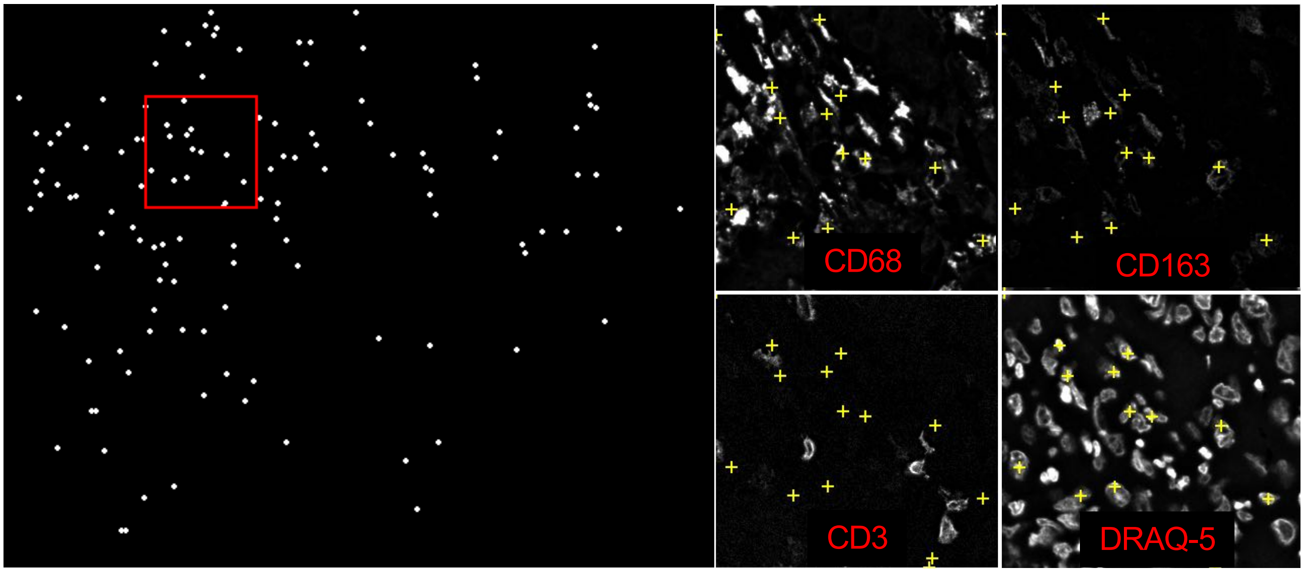

O. CD11b+CD68+ macrophage cluster

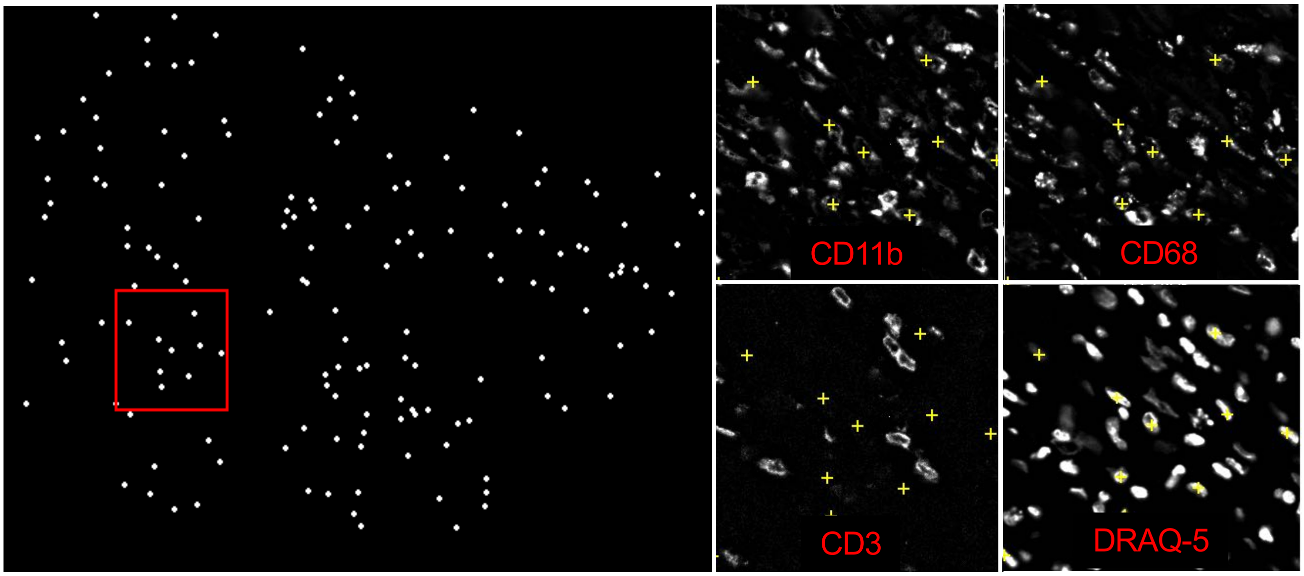

P. CD11c dendritic cell cluster

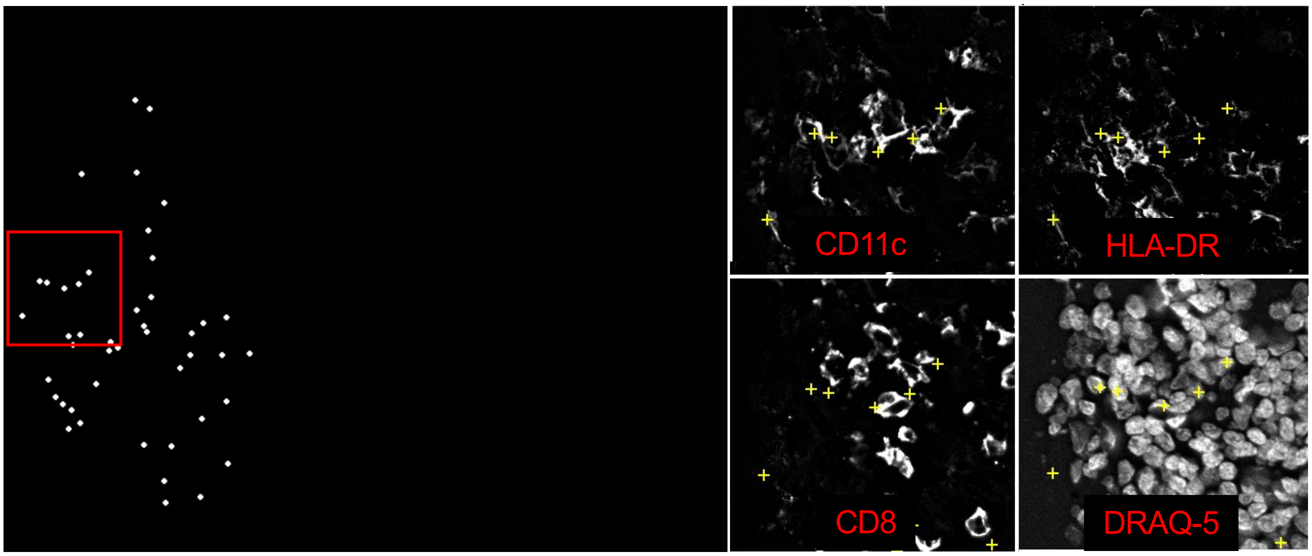

Q. NK cell cluster

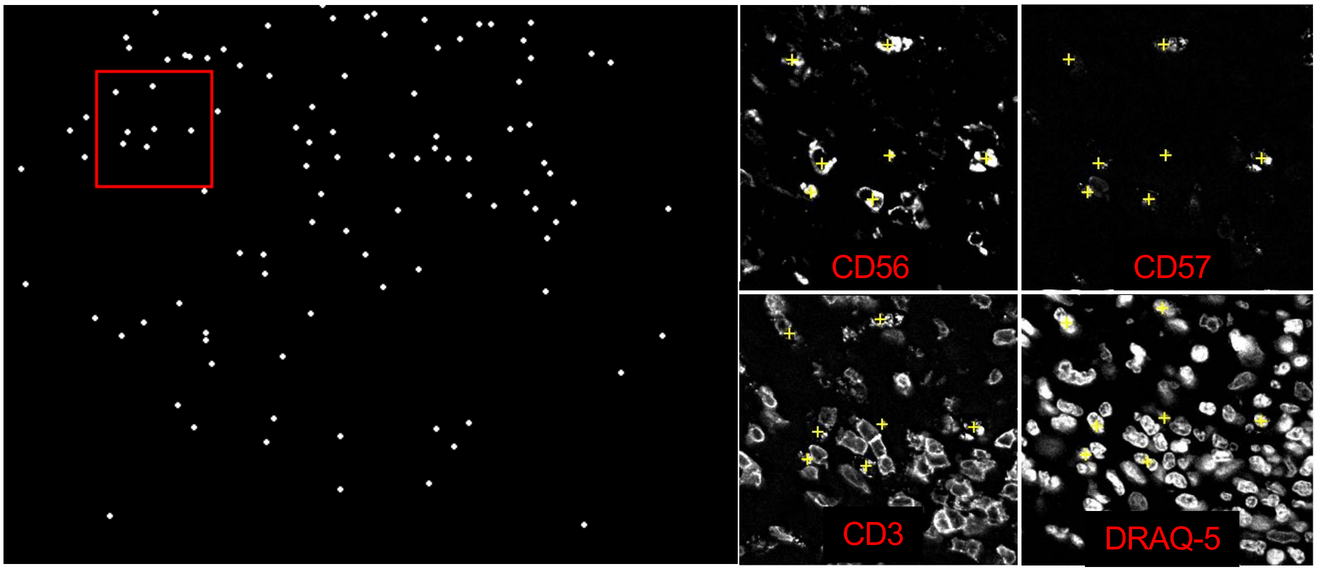

R. CD3+ T cell cluster

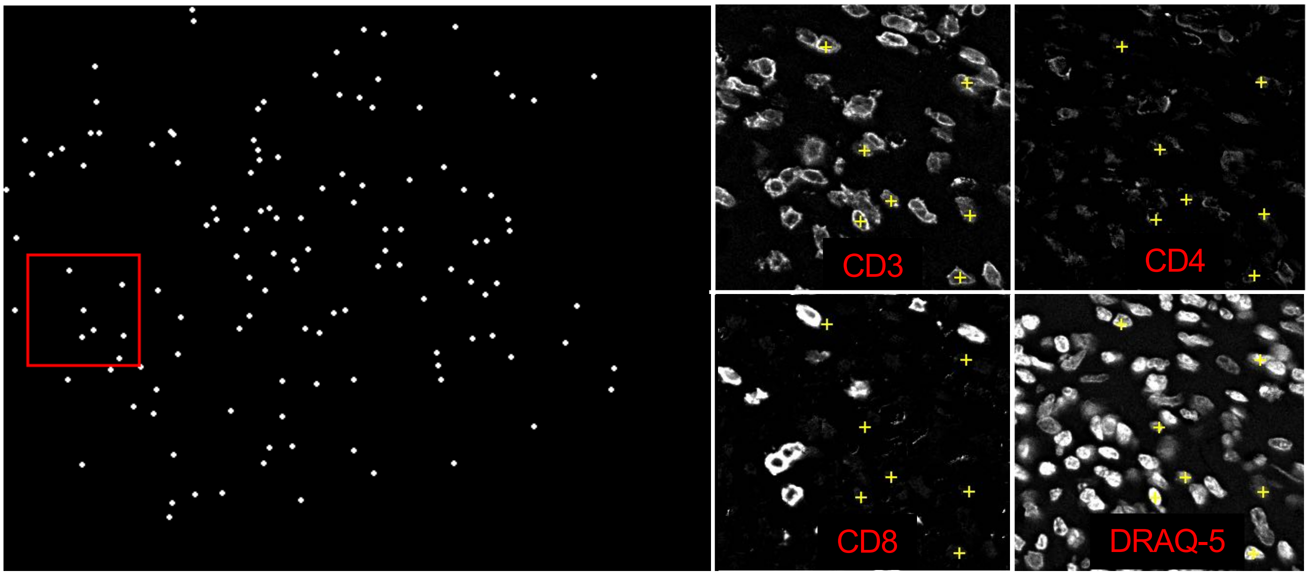

#### Figure S11

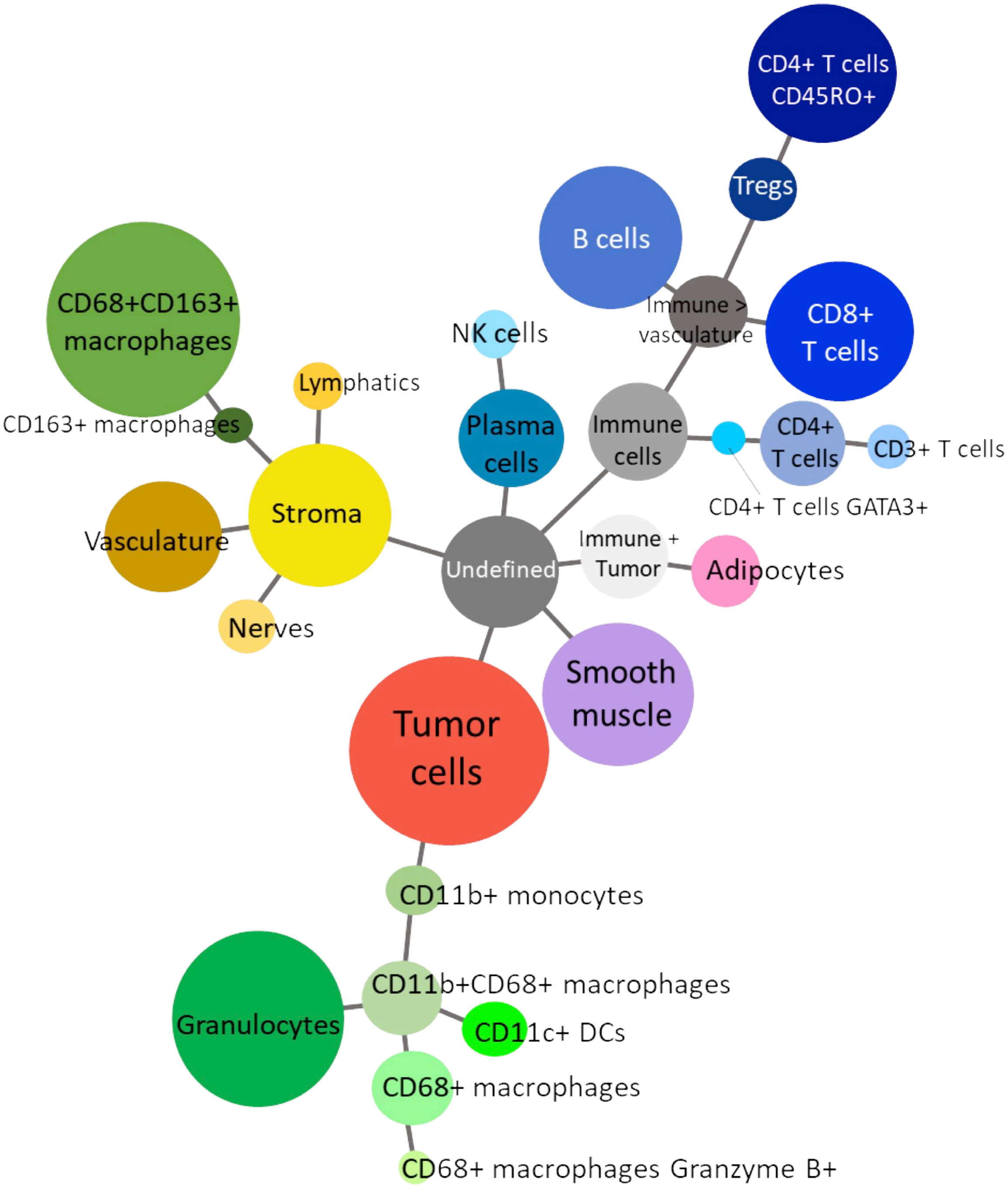

28 unique clusters

# CD2

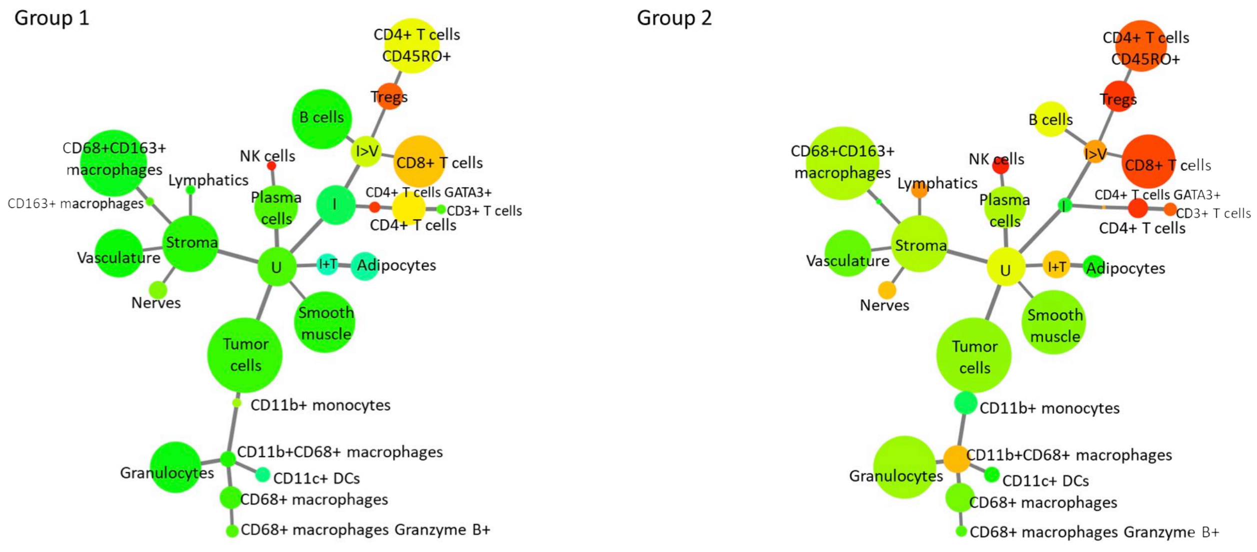

Group 1
Group 2
CD4+ T cells CD45RO+
Tregs
B cells
I>V
CD8+ T cells
CD68+CD163+ macrophages
NK cells
Lymphatics
Plasma cells
I
CD4+ T cells GATA3+
CD3+ T cells
CD4+ T cells
CD163+ macrophages
Stroma
Vasculature
U
I+T
Adipocytes
Nerves
Smooth muscle
Tumor cells
CD11b+ monocytes
CD11b+CD68+ macrophages
Granulocytes
CD11c+ DCs
CD68+ macrophages
CD68+ macrophages Granzyme B+

# CD3

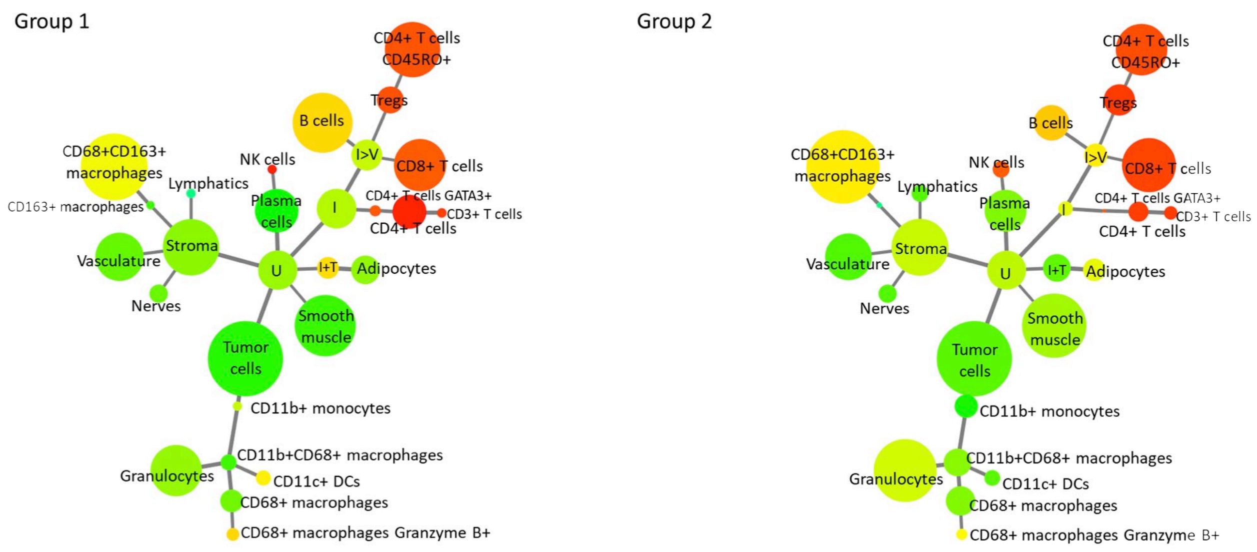

Group 1
Group 2
CD4+ T cells CD45RO+
Tregs
B cells
I>V
CD8+ T cells
CD68+CD163+ macrophages
NK cells
Lymphatics
Plasma cells
I
CD4+ T cells GATA3+
CD3+ T cells
CD4+ T cells
CD163+ macrophages
Stroma
Vasculature
U
I+T
Adipocytes
Nerves
Smooth muscle
Tumor cells
CD11b+ monocytes
CD11b+CD68+ macrophages
Granulocytes
CD11c+ DCs
CD68+ macrophages
CD68+ macrophages Granzyme B+

# CD4

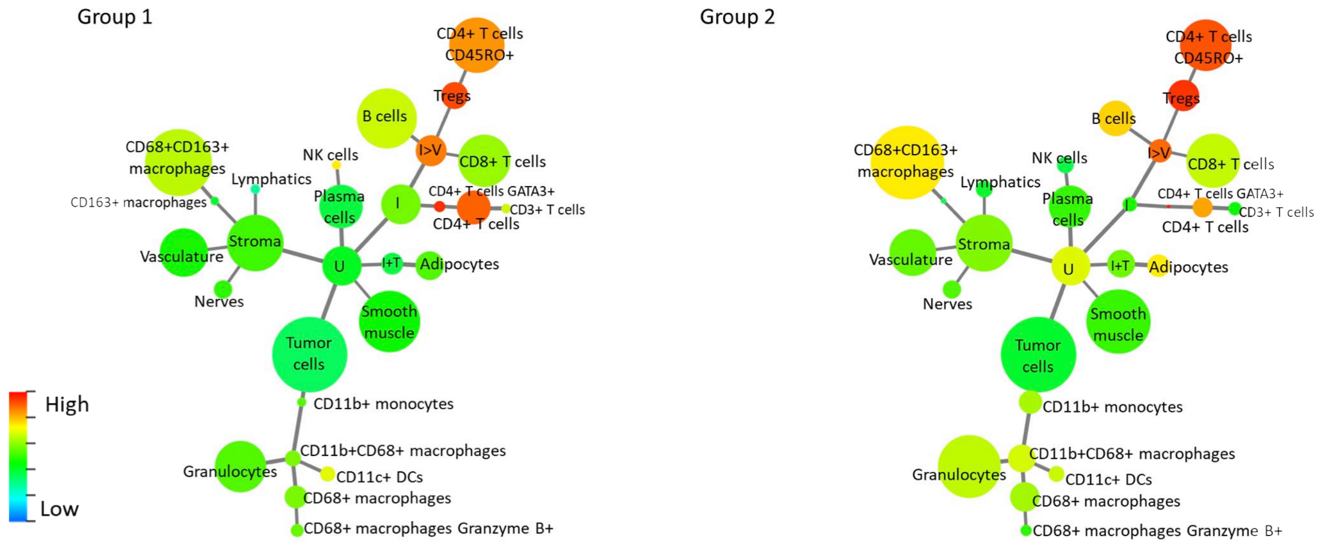

Group 1
Group 2
CD4+ T cells CD45RO+
Tregs
B cells
I>V
CD8+ T cells
CD68+CD163+ macrophages
NK cells
Lymphatics
Plasma cells
I
CD4+ T cells GATA3+
CD3+ T cells
CD4+ T cells
CD163+ macrophages
Stroma
Vasculature
U
I+T
Adipocytes
Nerves
Smooth muscle
Tumor cells
CD11b+ monocytes
CD11b+CD68+ macrophages
Granulocytes
CD11c+ DCs
CD68+ macrophages
CD68+ macrophages Granzyme B+
High
Low

# CD5

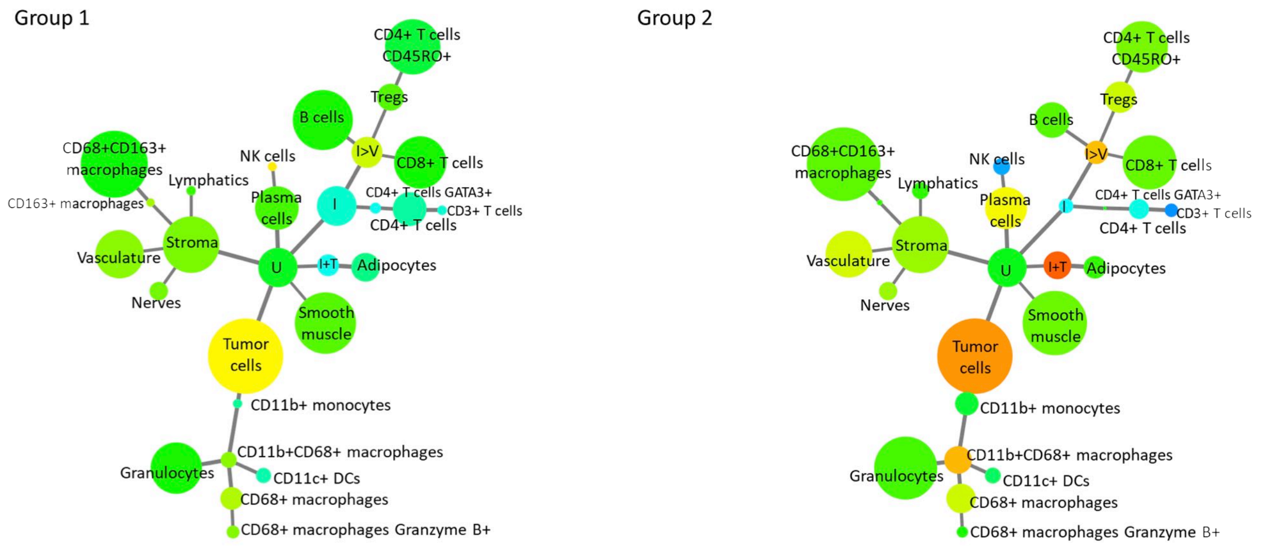

Group 1
Group 2
CD4+ T cells CD45RO+
Tregs
B cells
I>V
CD8+ T cells
CD68+CD163+ macrophages
NK cells
Lymphatics
Plasma cells
CD163+ macrophages
I
CD4+ T cells GATA3+
CD3+ T cells
CD4+ T cells
Stroma
Vasculature
U
I+T
Adipocytes
Nerves
Smooth muscle
Tumor cells
CD11b+ monocytes
CD11b+CD68+ macrophages
Granulocytes
CD11c+ DCs
CD68+ macrophages
CD68+ macrophages Granzyme B+

# CD7

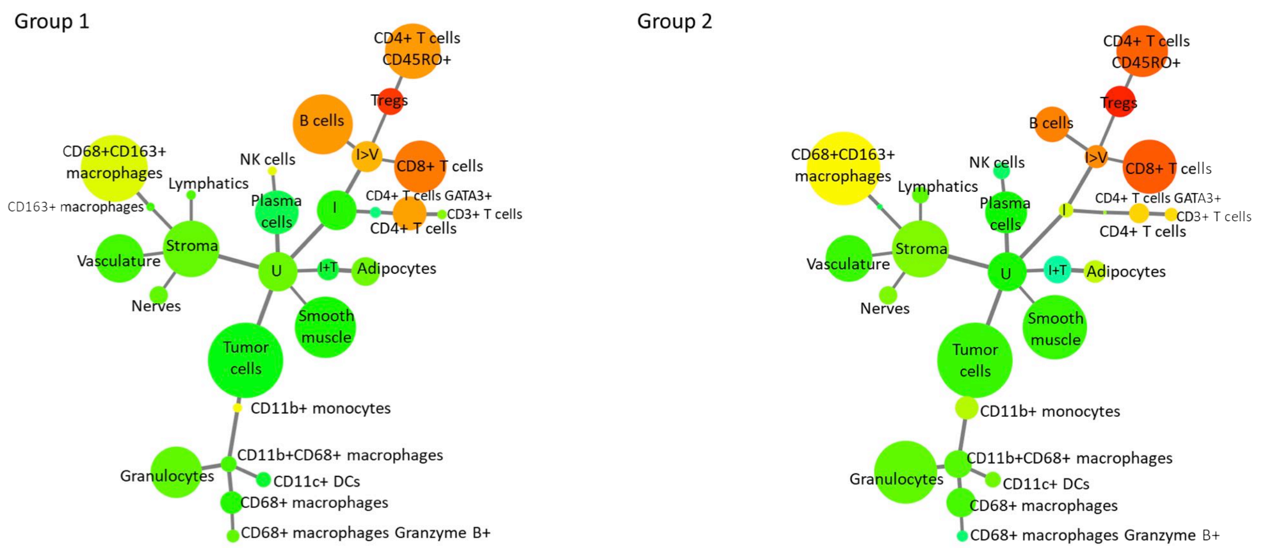

Group 1
Group 2
CD4+ T cells CD45RO+
Tregs
B cells
I>V
CD8+ T cells
CD68+CD163+ macrophages
NK cells
Lymphatics
Plasma cells
CD163+ macrophages
I
CD4+ T cells GATA3+
CD3+ T cells
CD4+ T cells
Stroma
Vasculature
U
I+T
Adipocytes
Nerves
Smooth muscle
Tumor cells
CD11b+ monocytes
CD11b+CD68+ macrophages
Granulocytes
CD11c+ DCs
CD68+ macrophages
CD68+ macrophages Granzyme B+

# CD8

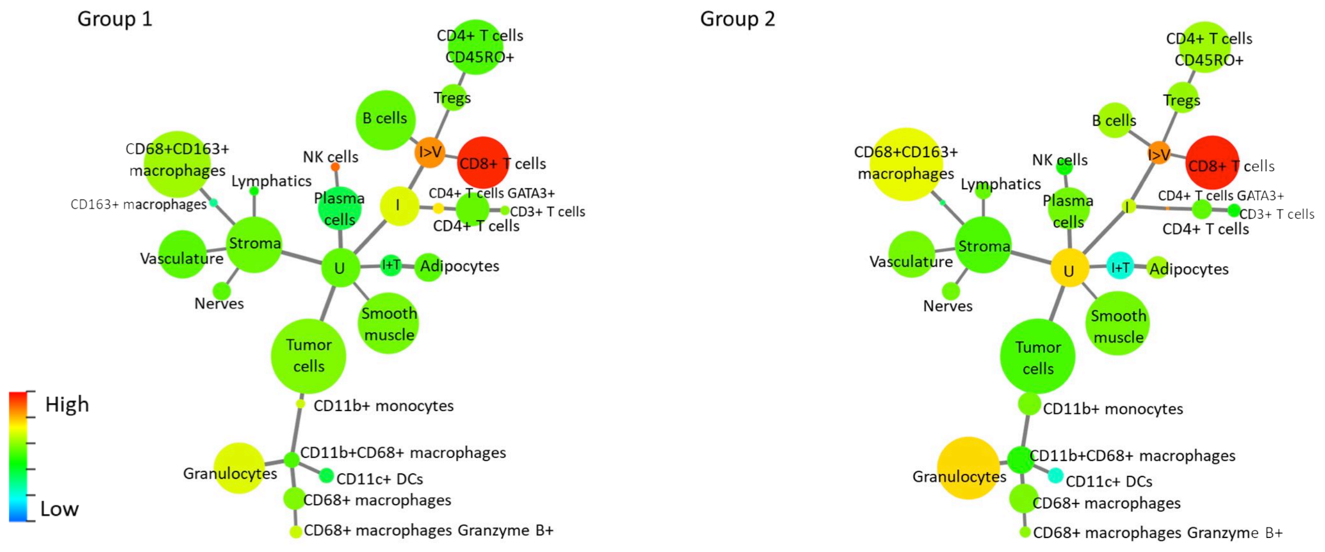

Group 1
Group 2
CD4+ T cells CD45RO+
Tregs
B cells
I>V
CD8+ T cells
CD68+CD163+ macrophages
NK cells
Lymphatics
Plasma cells
CD163+ macrophages
I
CD4+ T cells GATA3+
CD3+ T cells
CD4+ T cells
Stroma
Vasculature
U
I+T
Adipocytes
Nerves
Smooth muscle
Tumor cells
CD11b+ monocytes
CD11b+CD68+ macrophages
Granulocytes
CD11c+ DCs
CD68+ macrophages
CD68+ macrophages Granzyme B+
High
Low

# CD11b

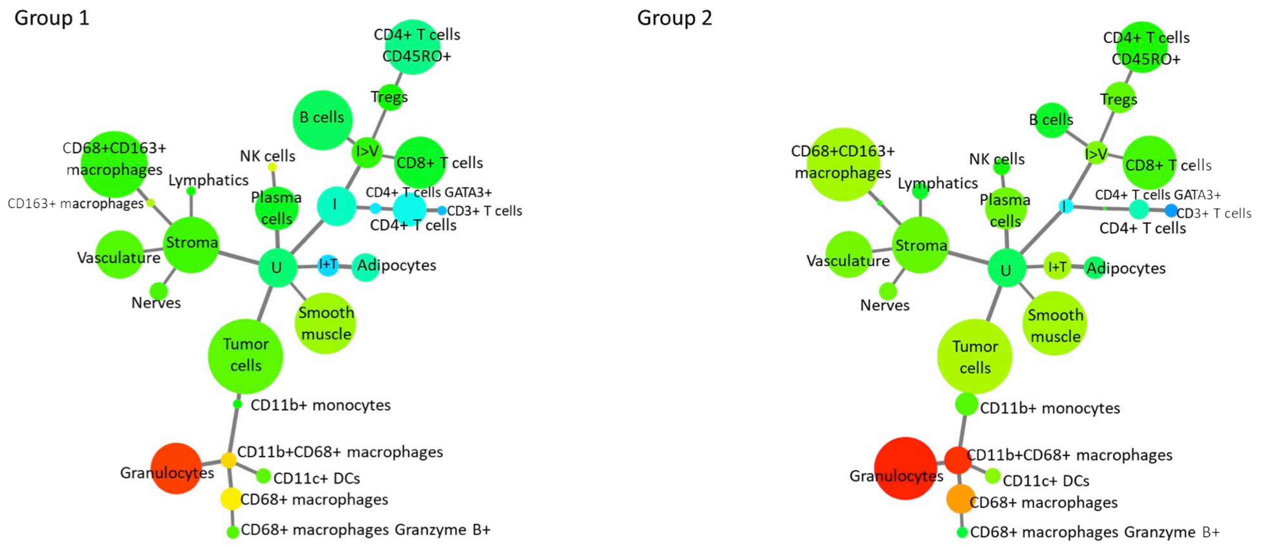

# CD11c

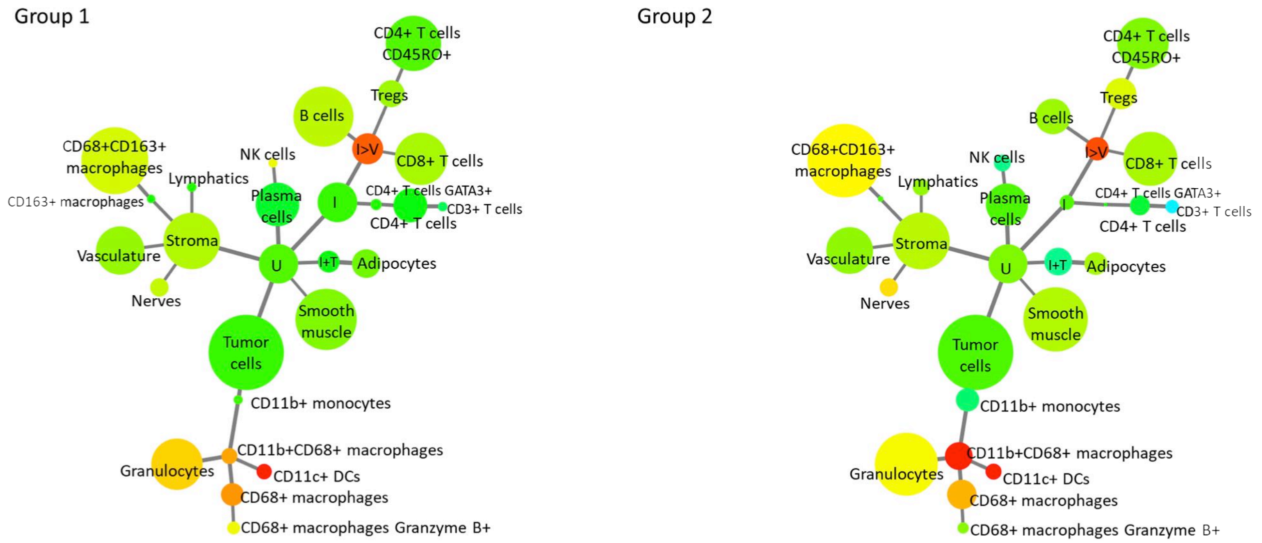

# CD15

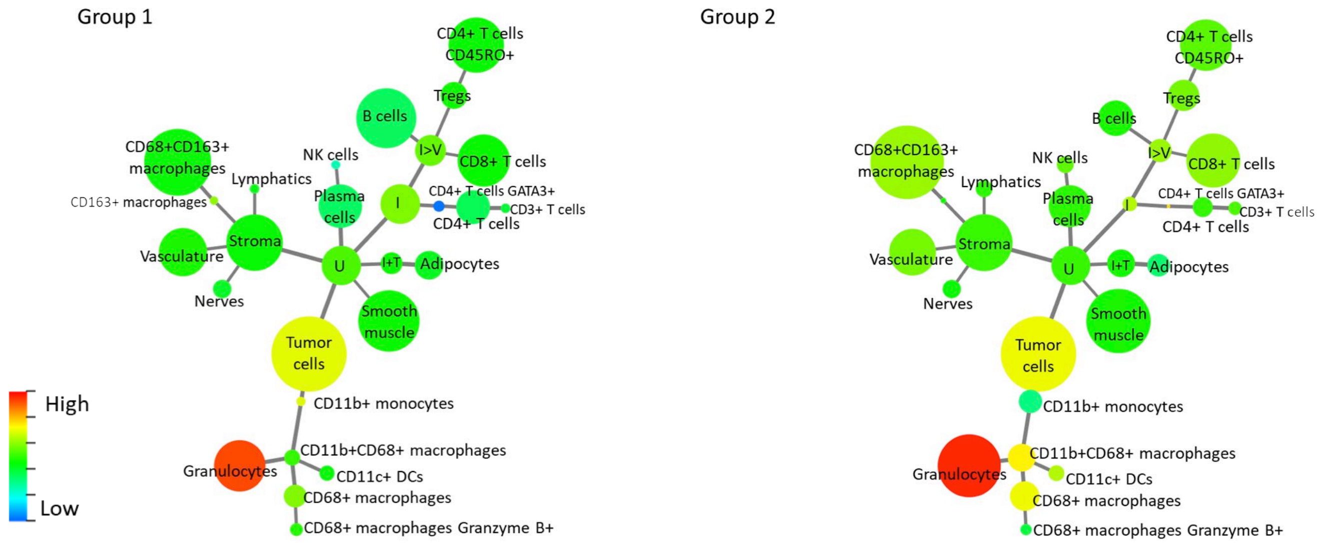

# CD20

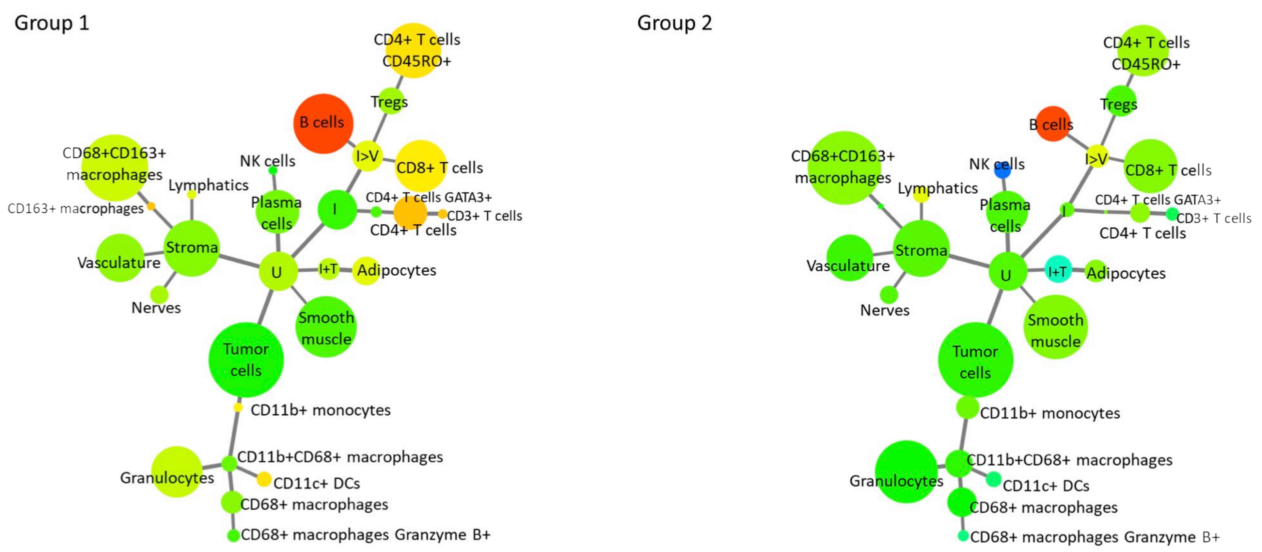

# CD21

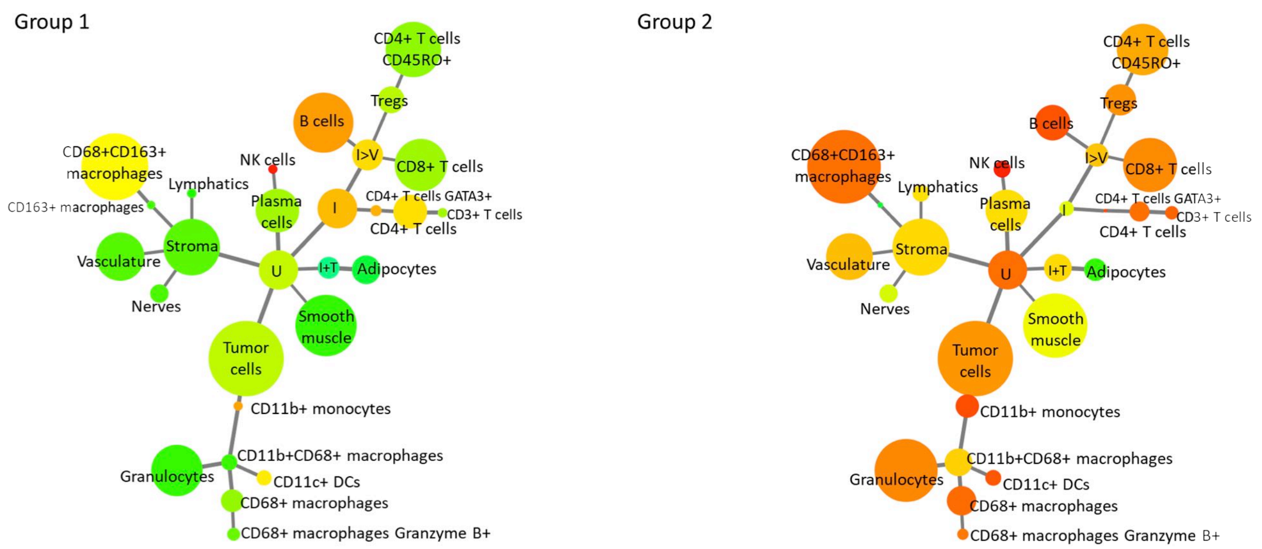

# CD25

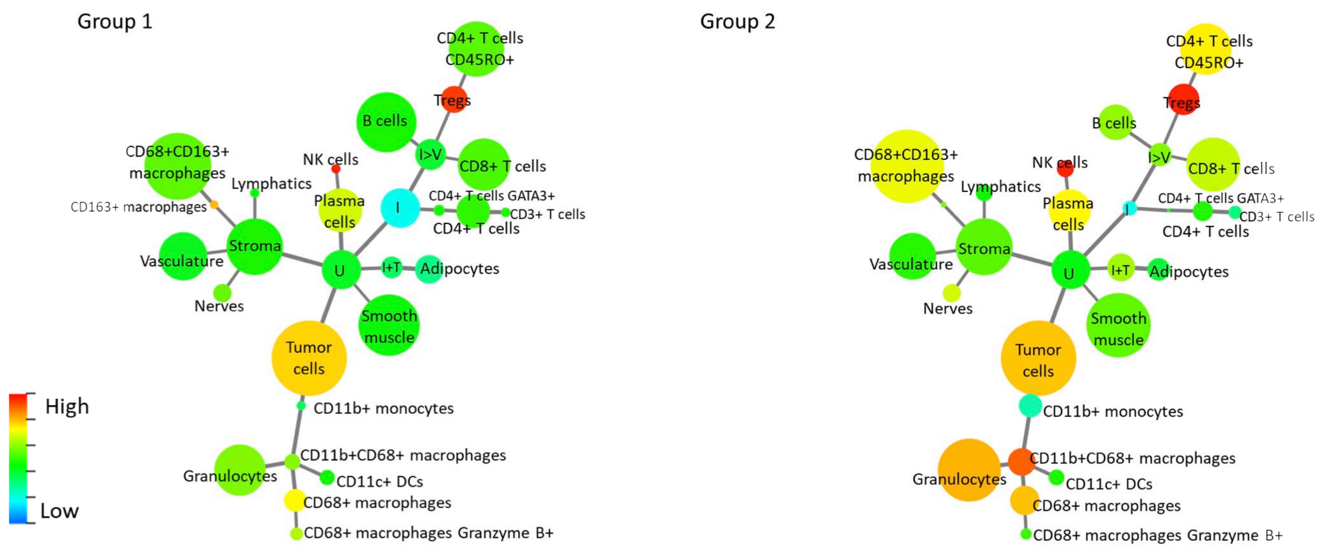

# CD30

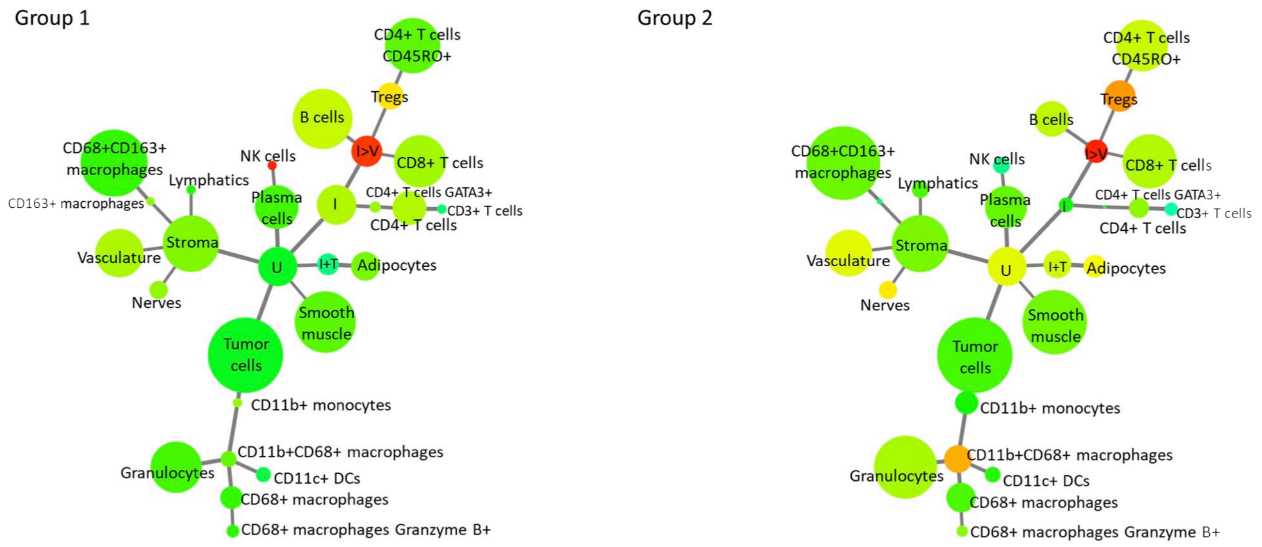

# CD31

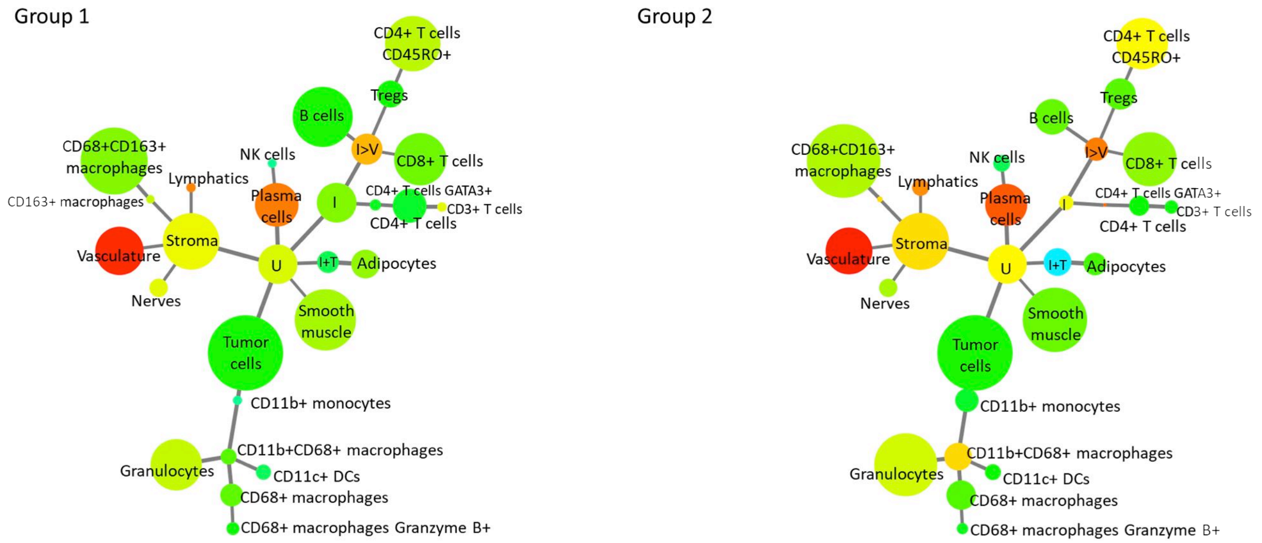

# CD34

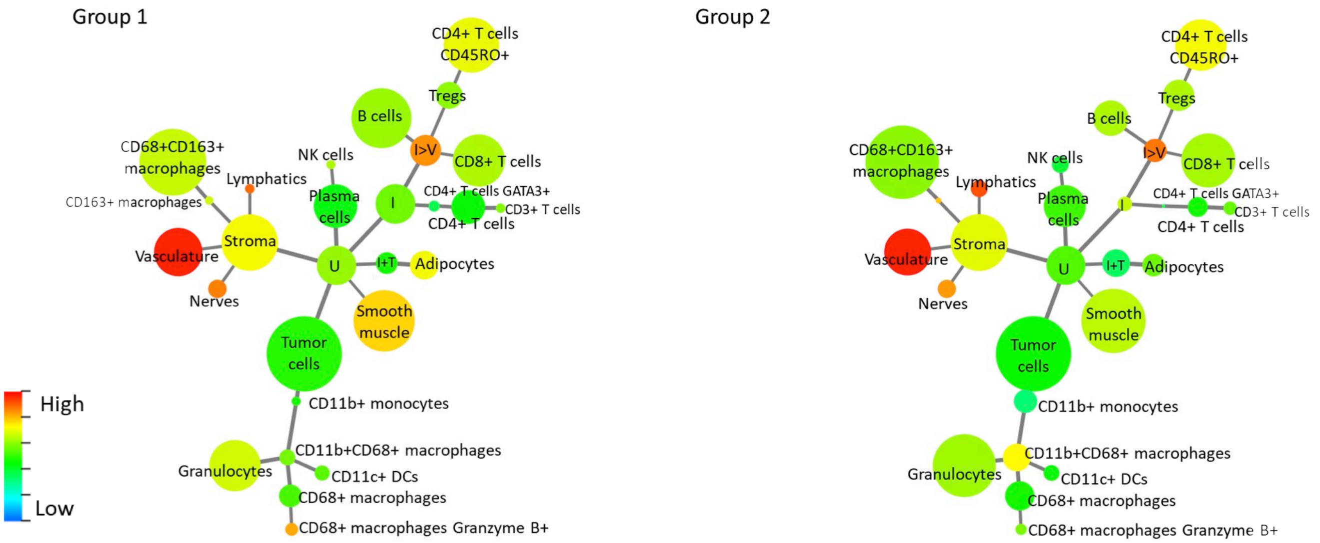

# CD38

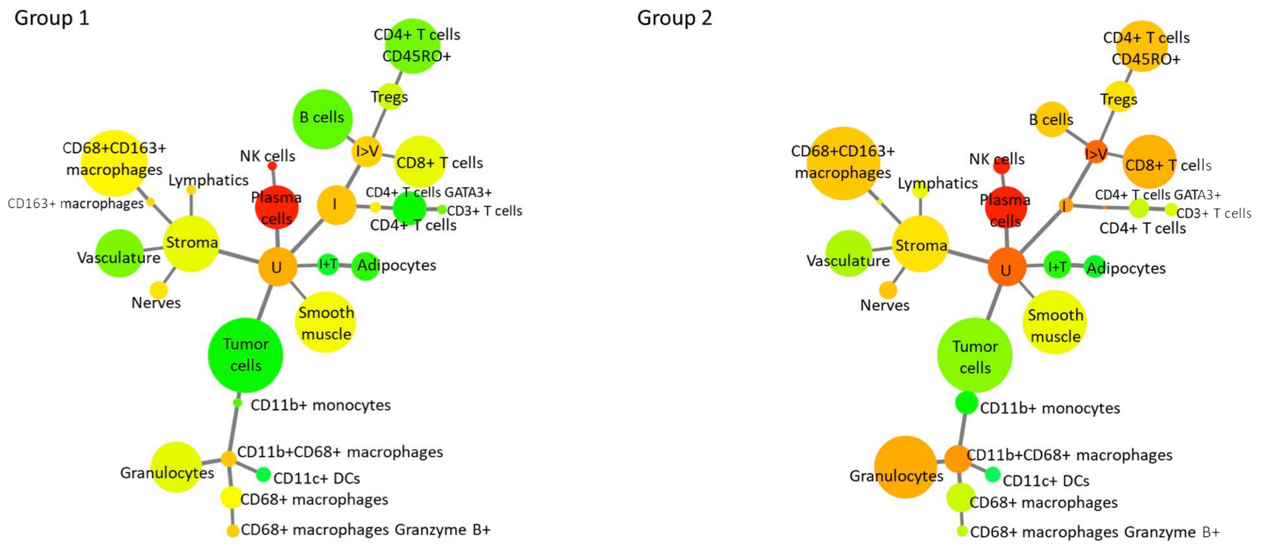

# CD44

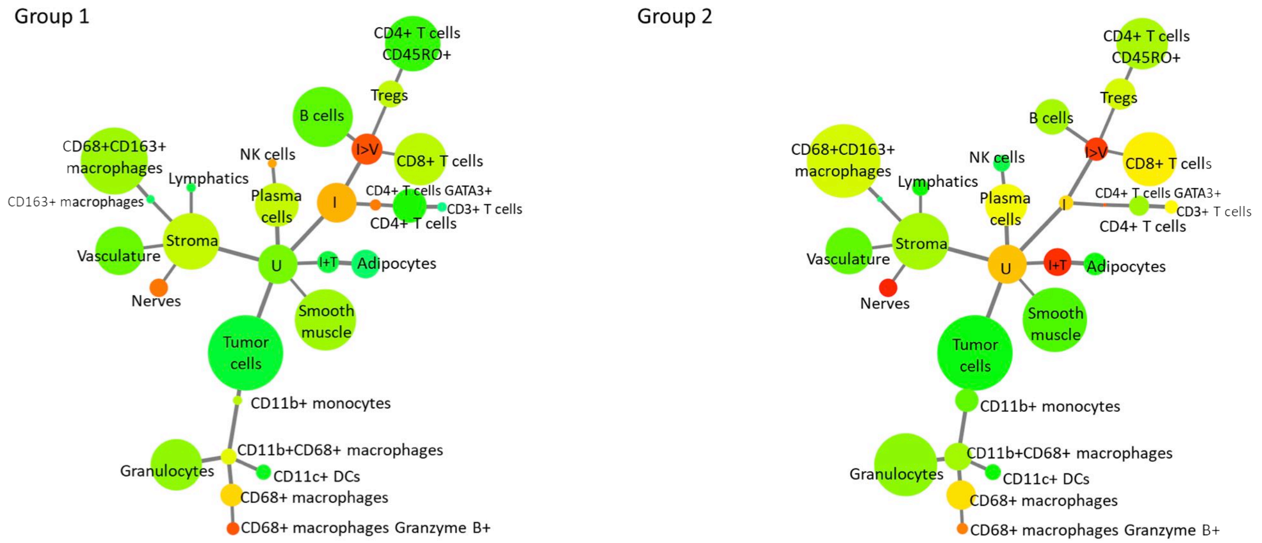

# CD45

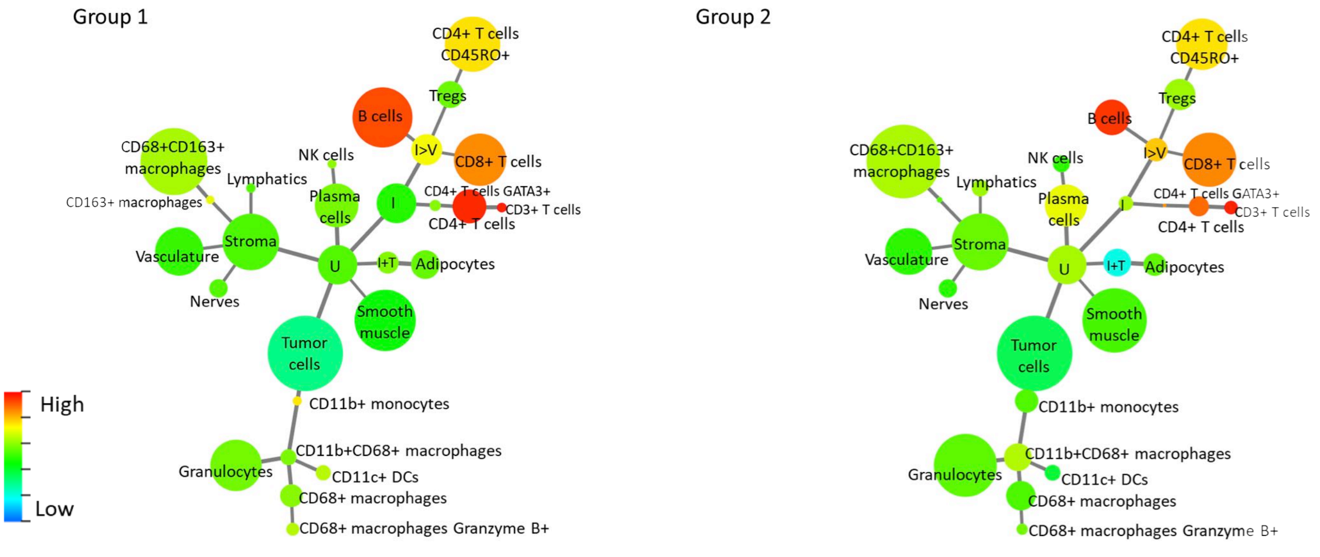

# CD45RA

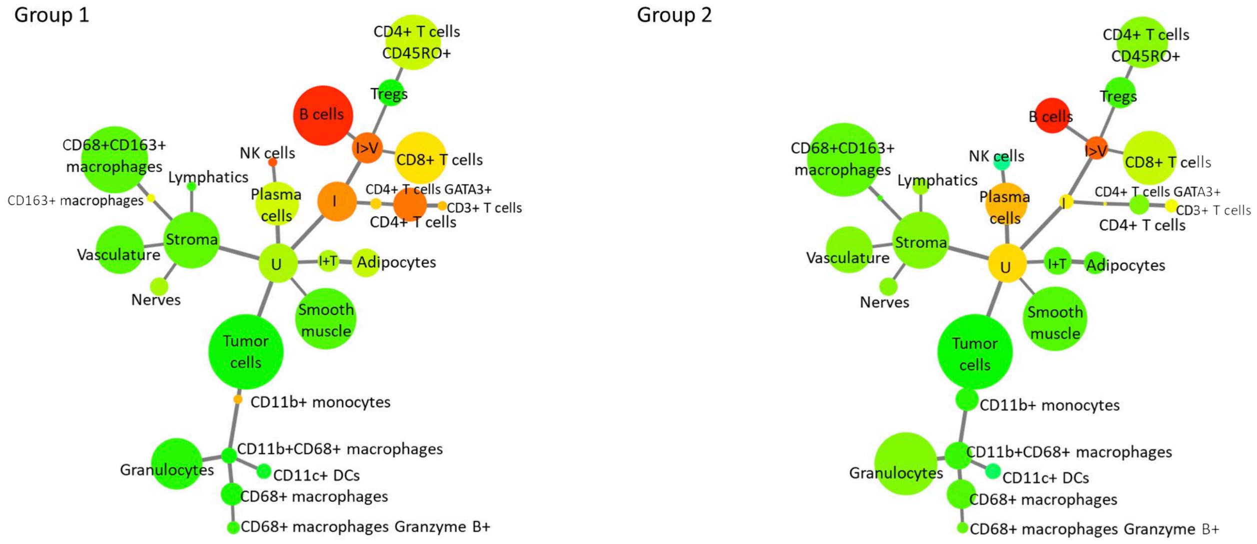

# CD45RO

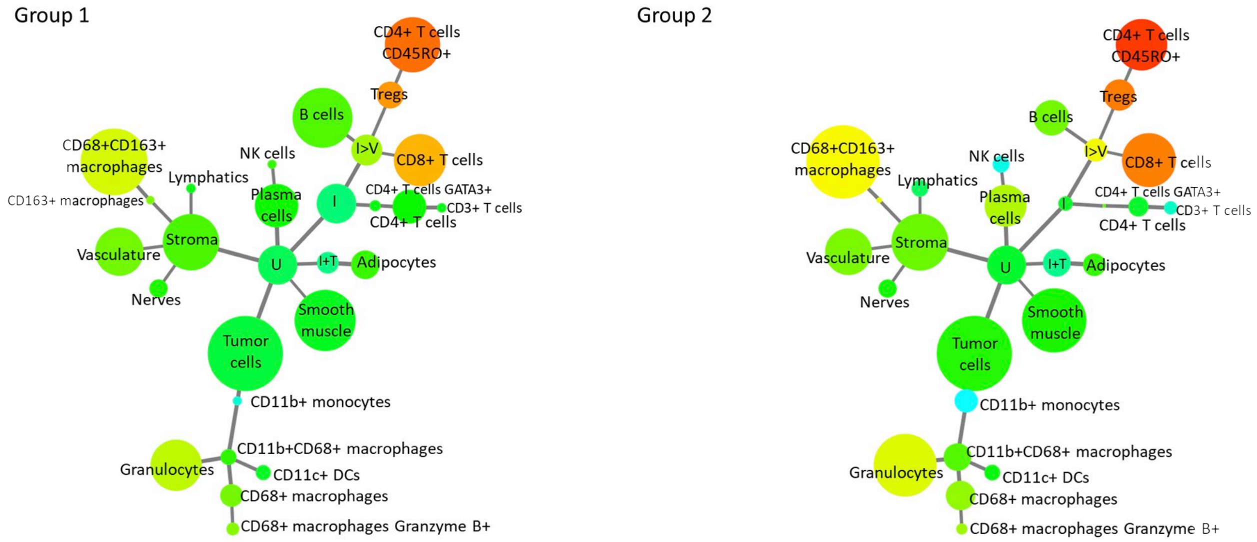

# CD56

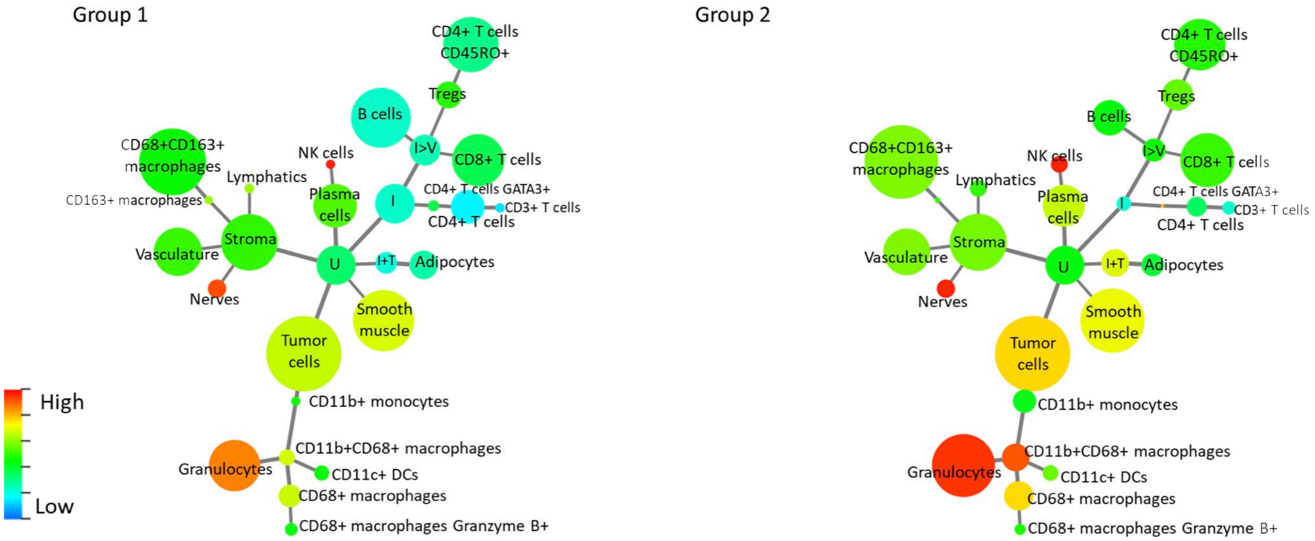

# CD57

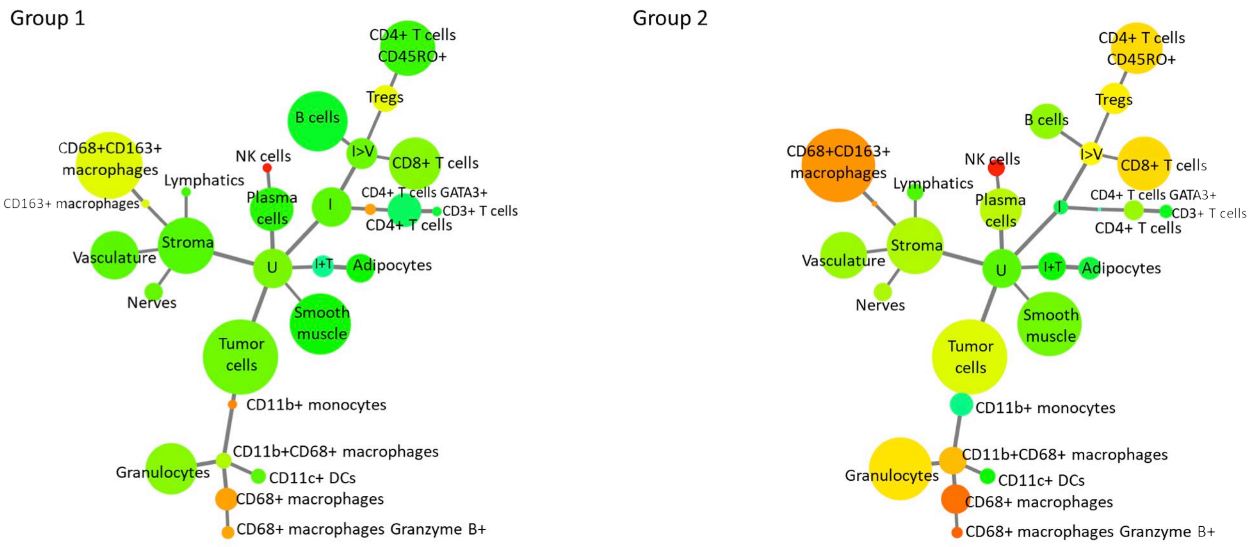

# CD68

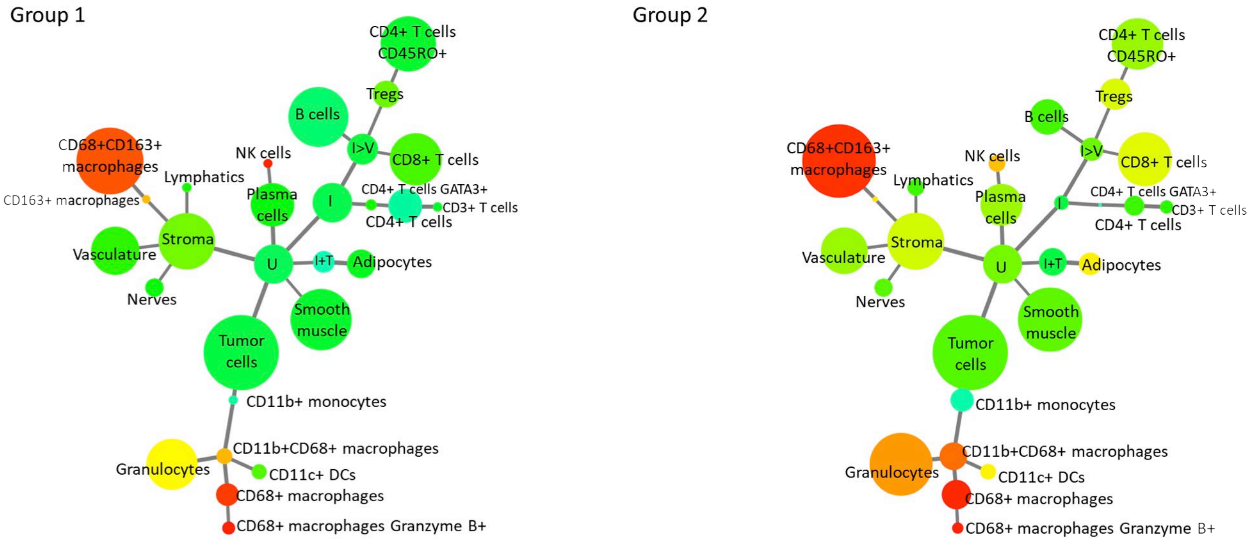

# CD71

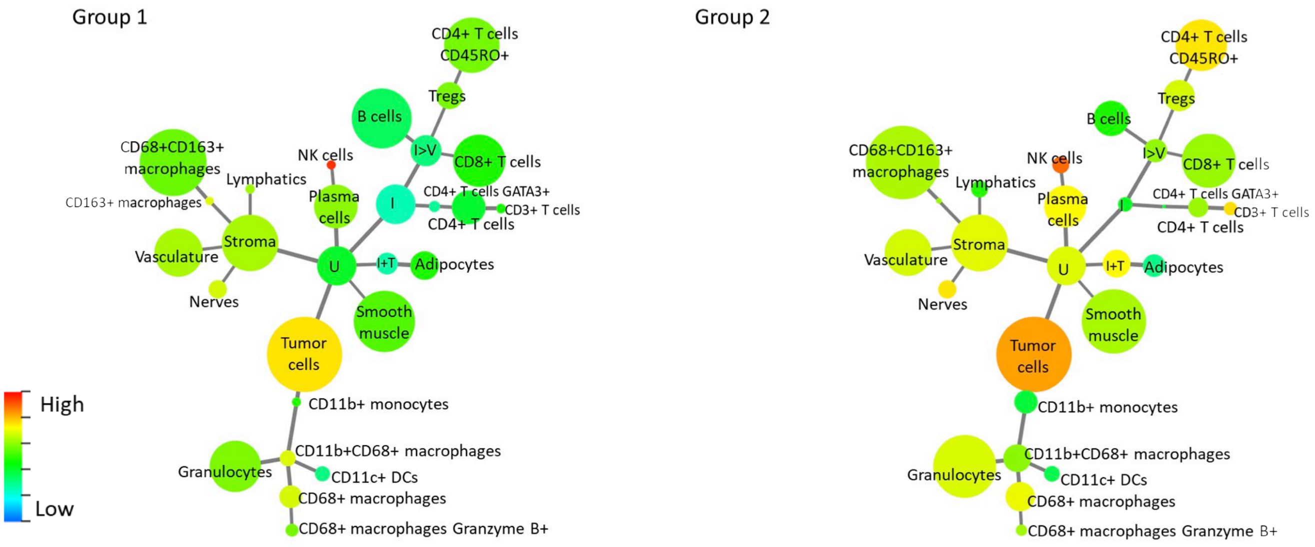

# CD138

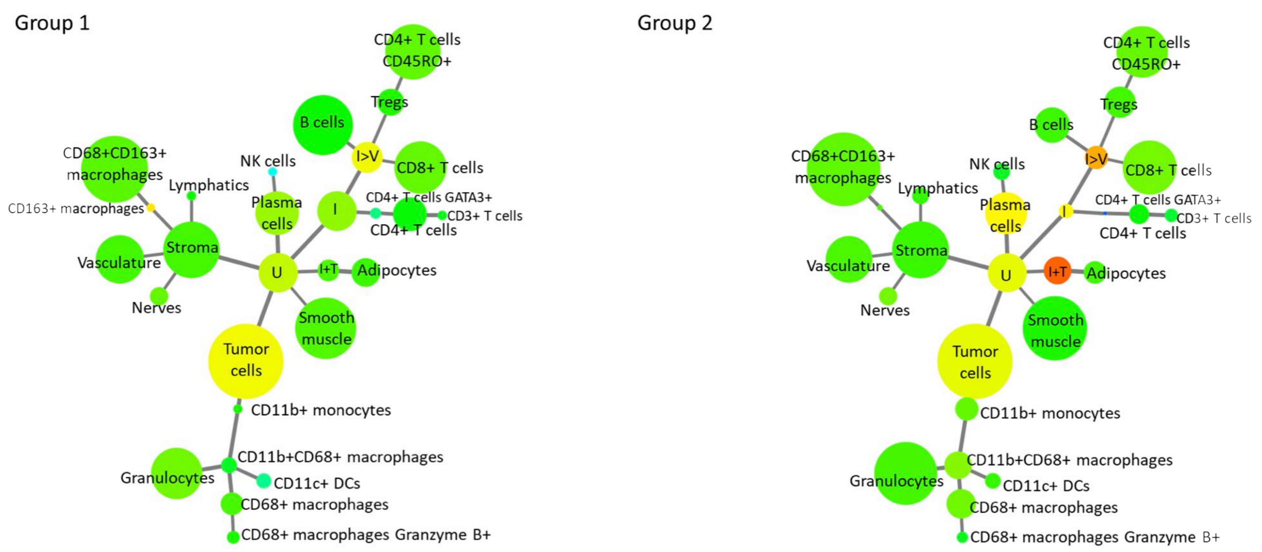

# CD163

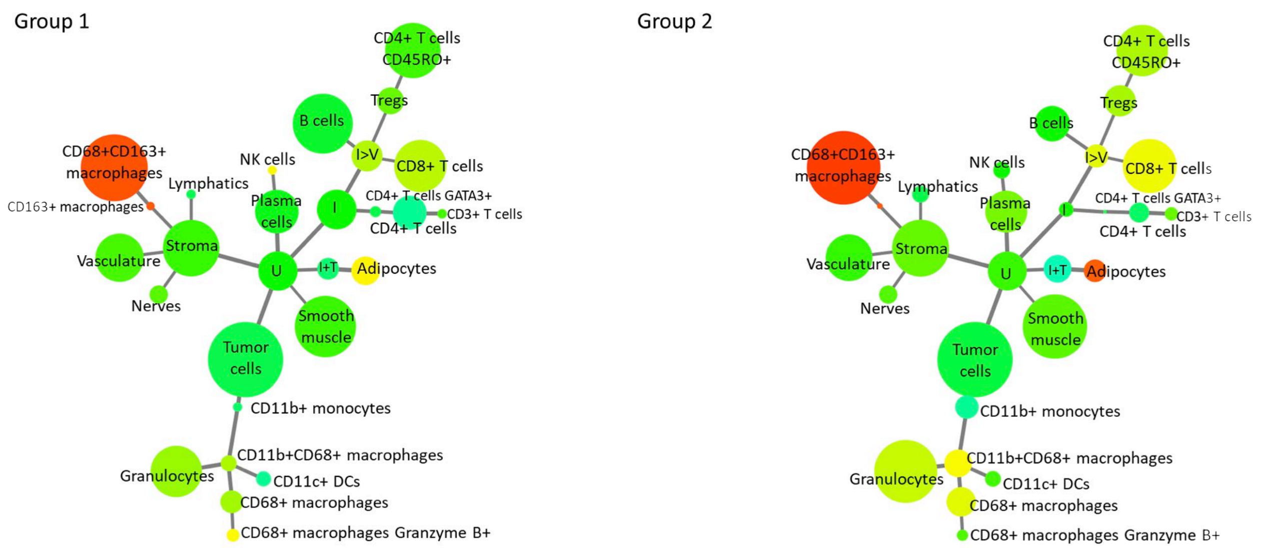

# CD194

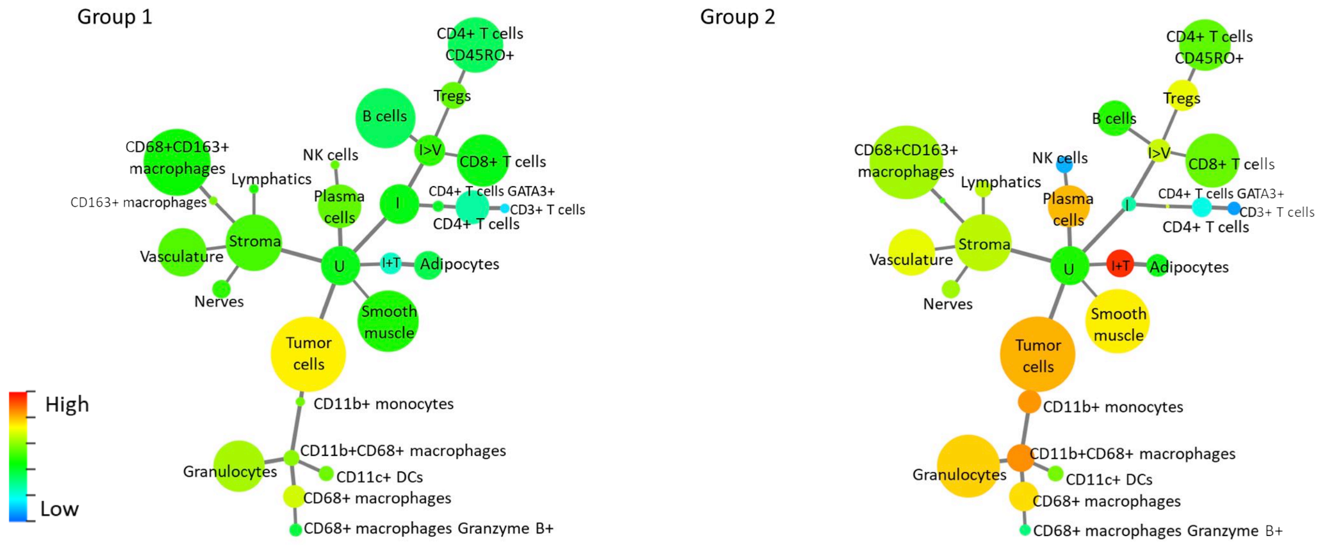

### BCL-2

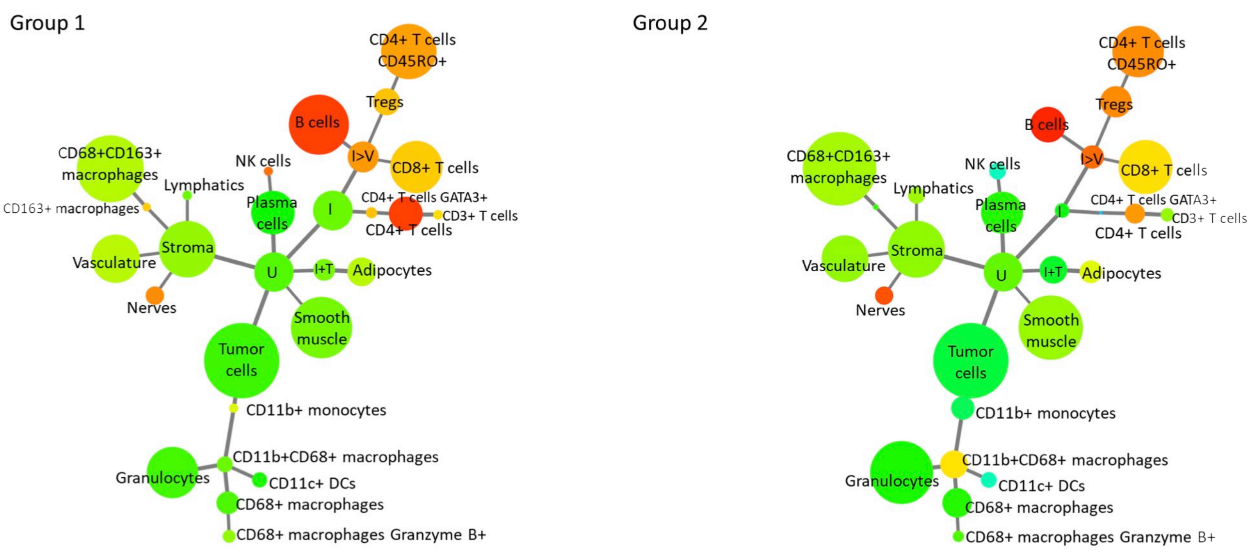

### Beta-catenin

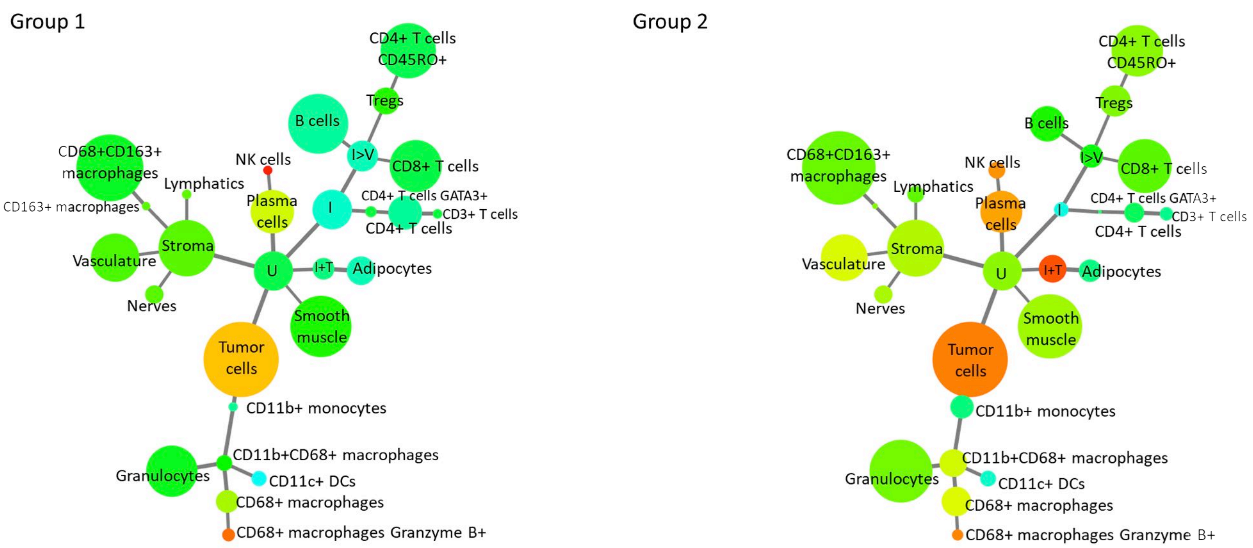

### CDX2

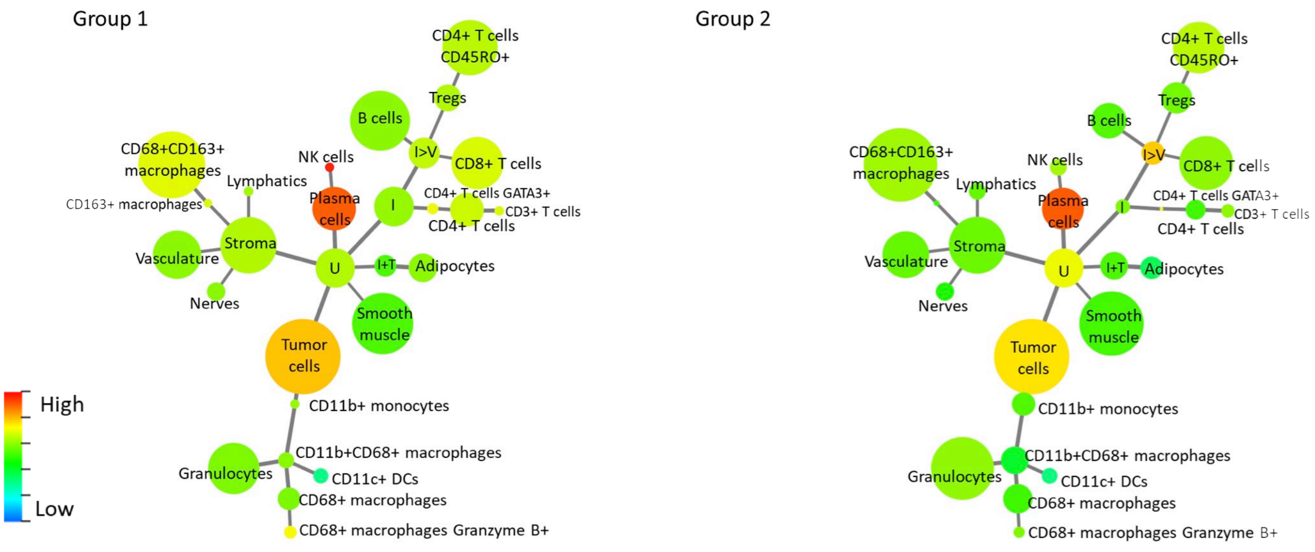

### Chromogranin A

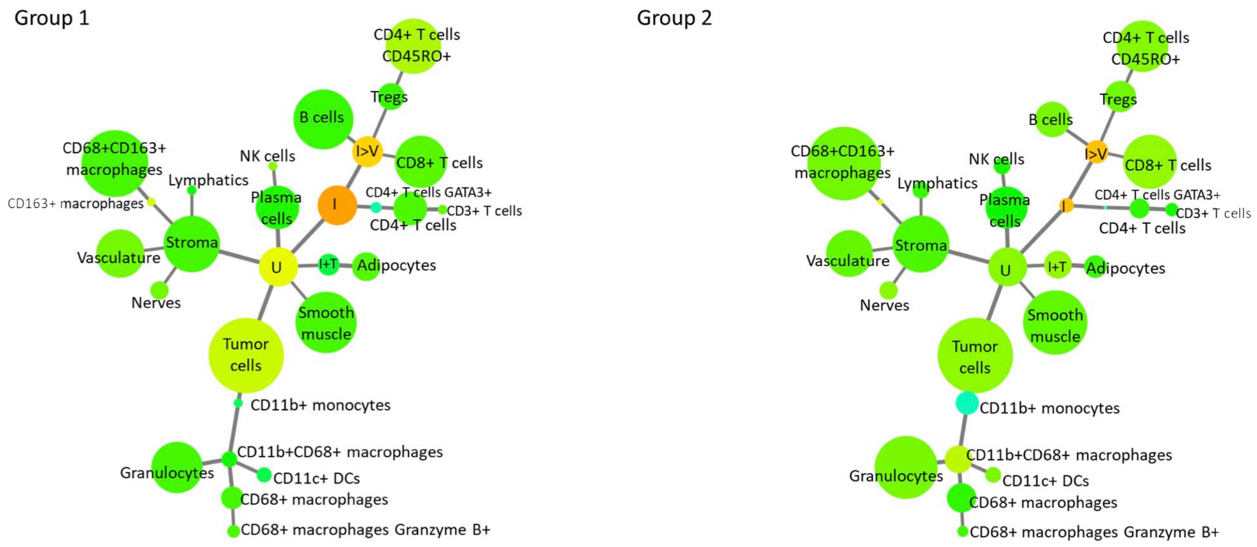

### Collagen IV

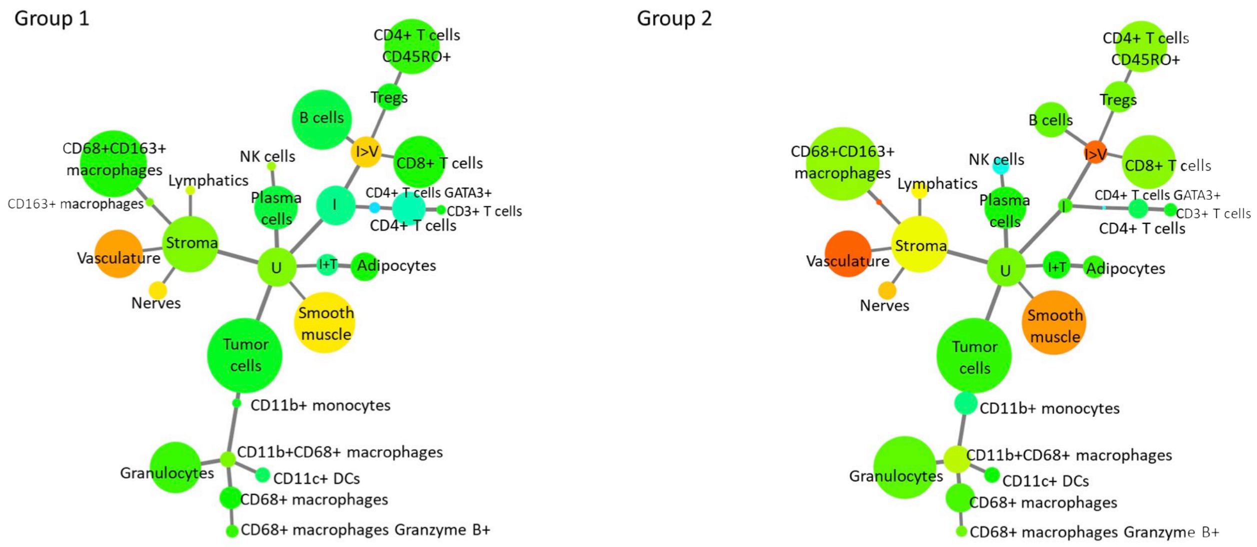

### Cytokeratin

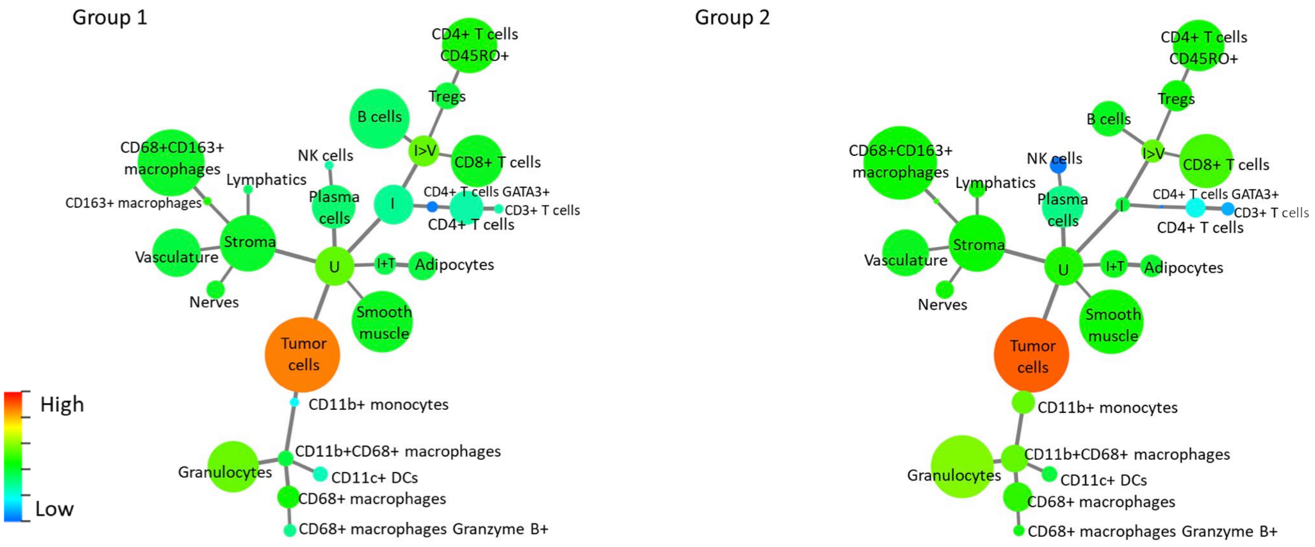

### EGFR

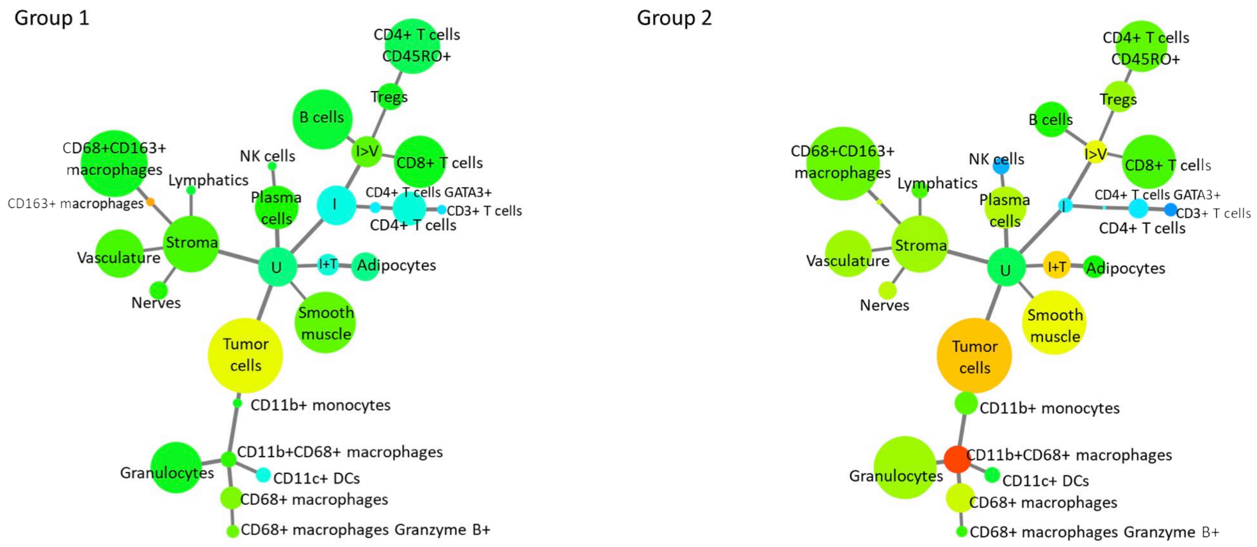

### FoxP3

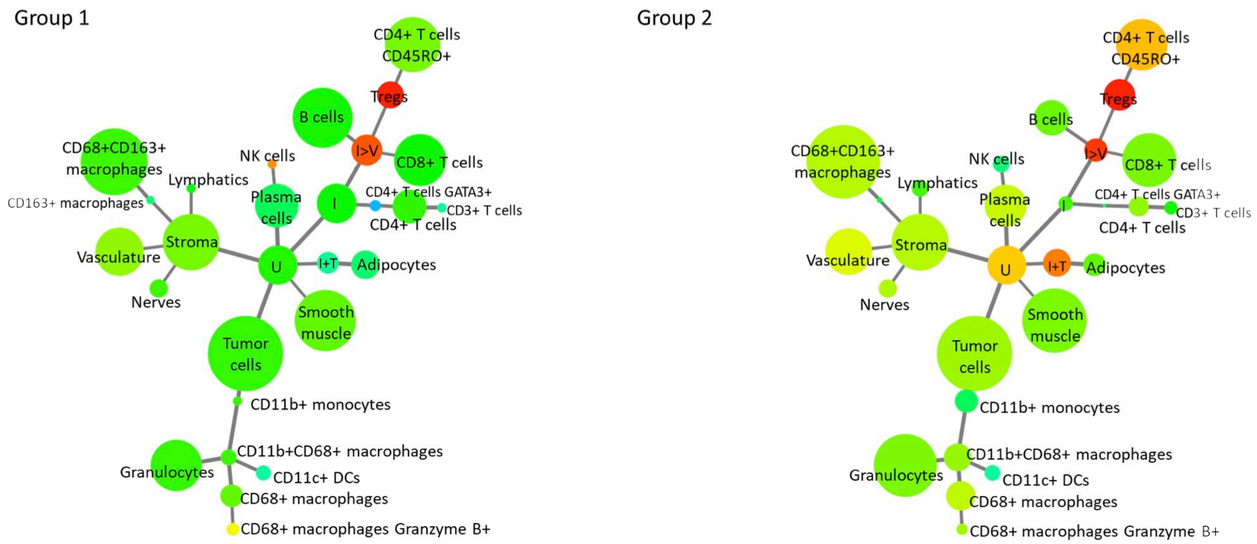

### GATA3

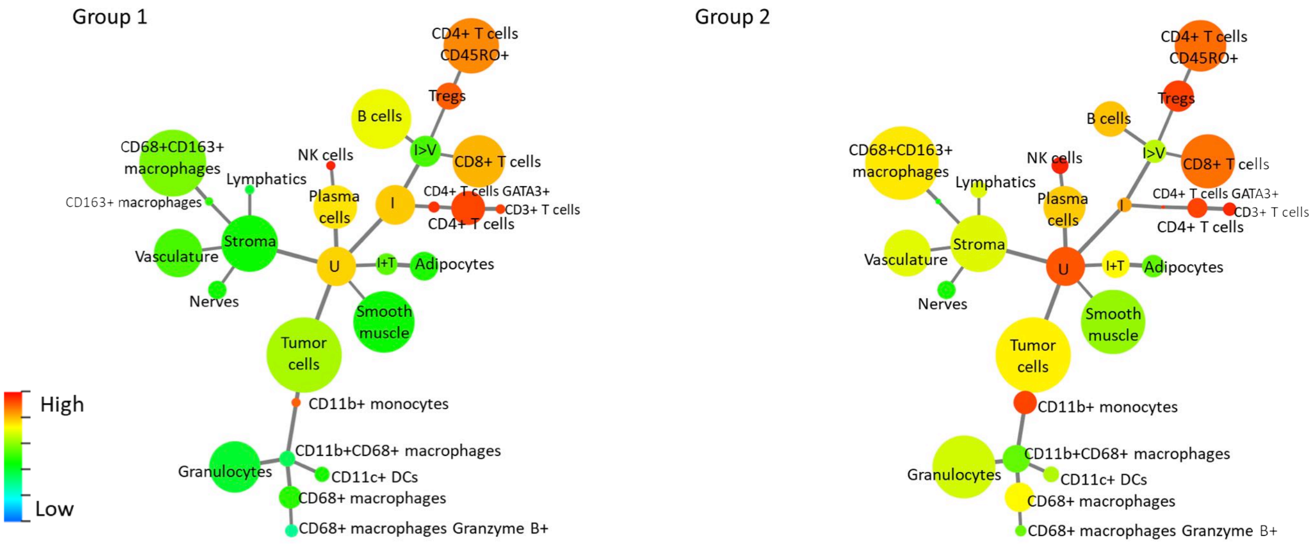

### GFAP

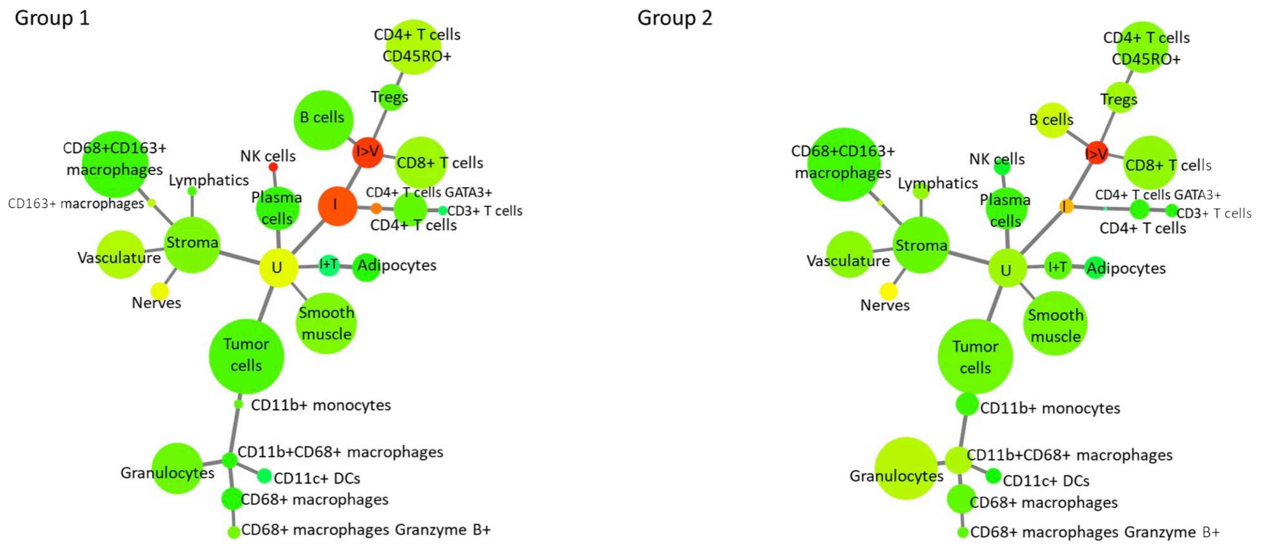

### Granzyme B

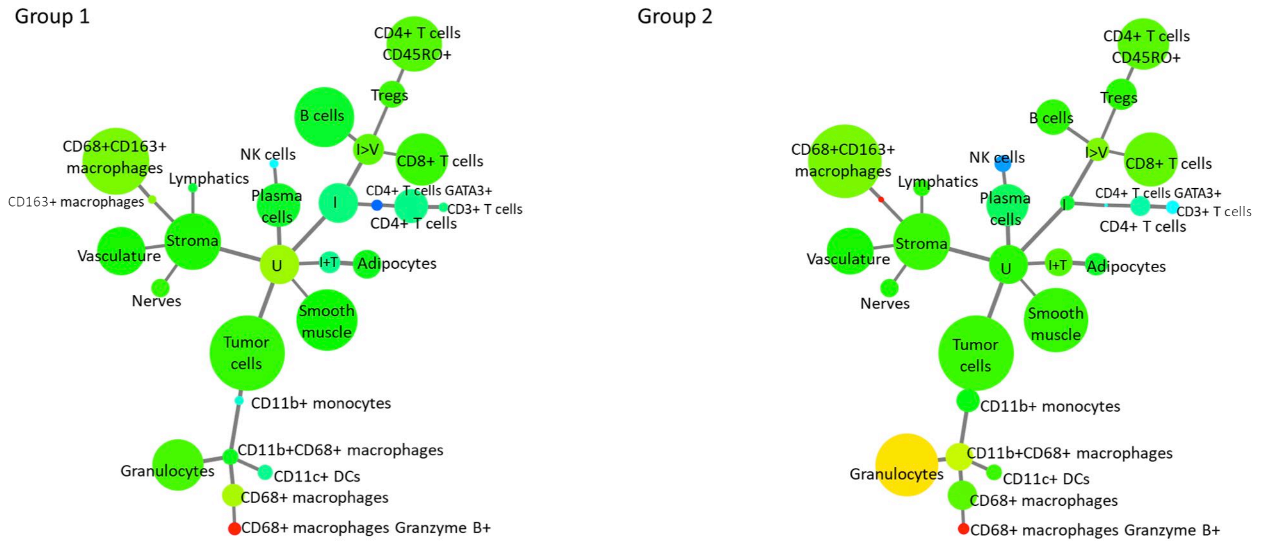

### HLA-DR

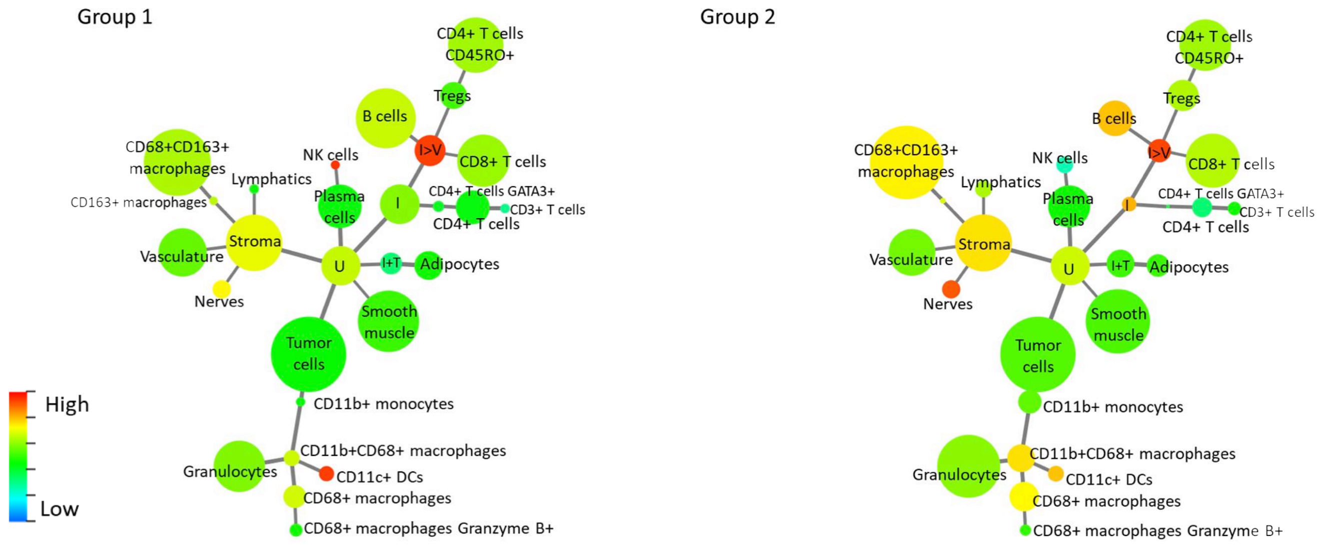

### ICOS

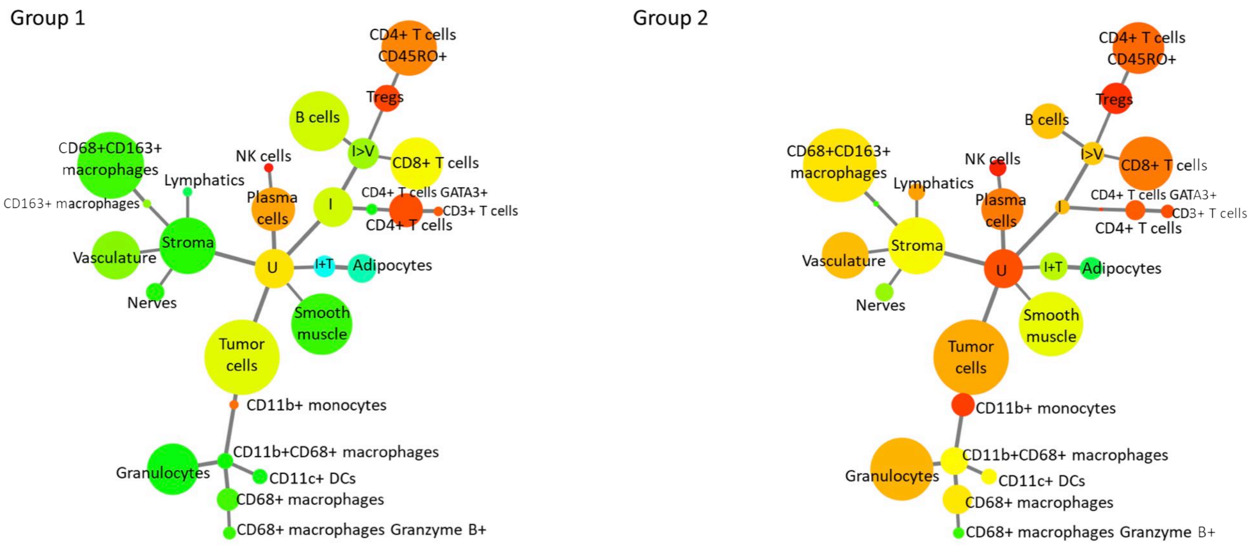

### IDO-1

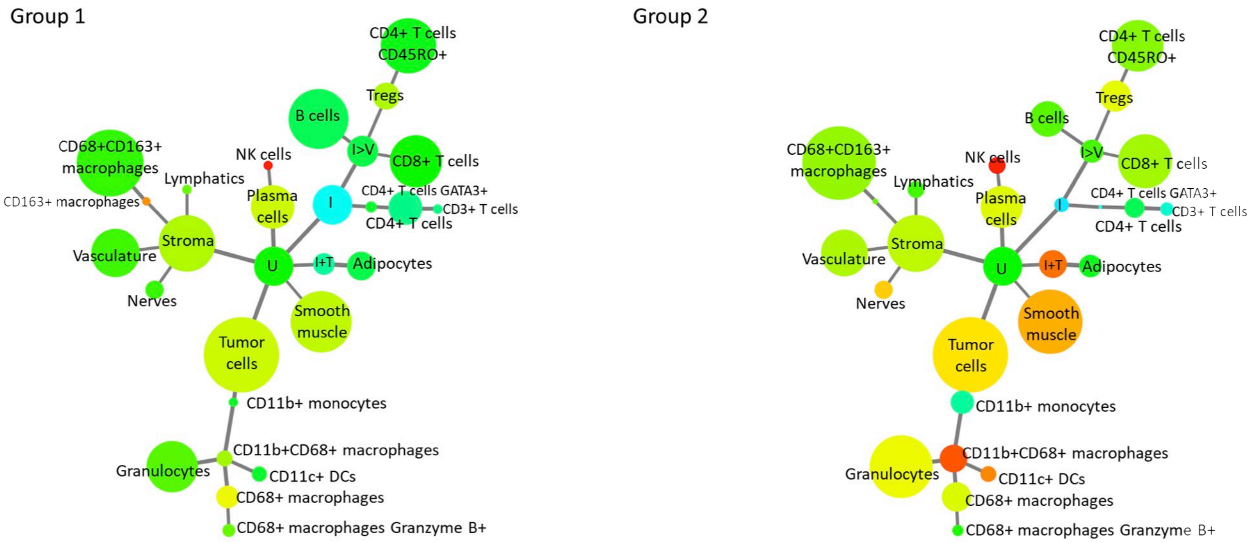

# Ki-67

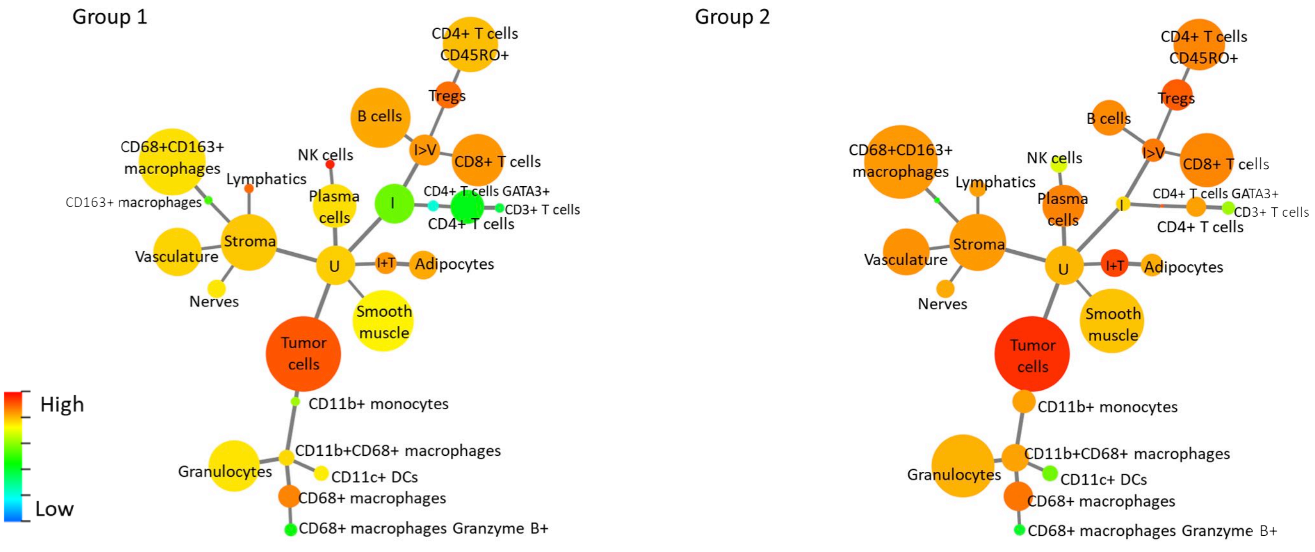

### LAG-3

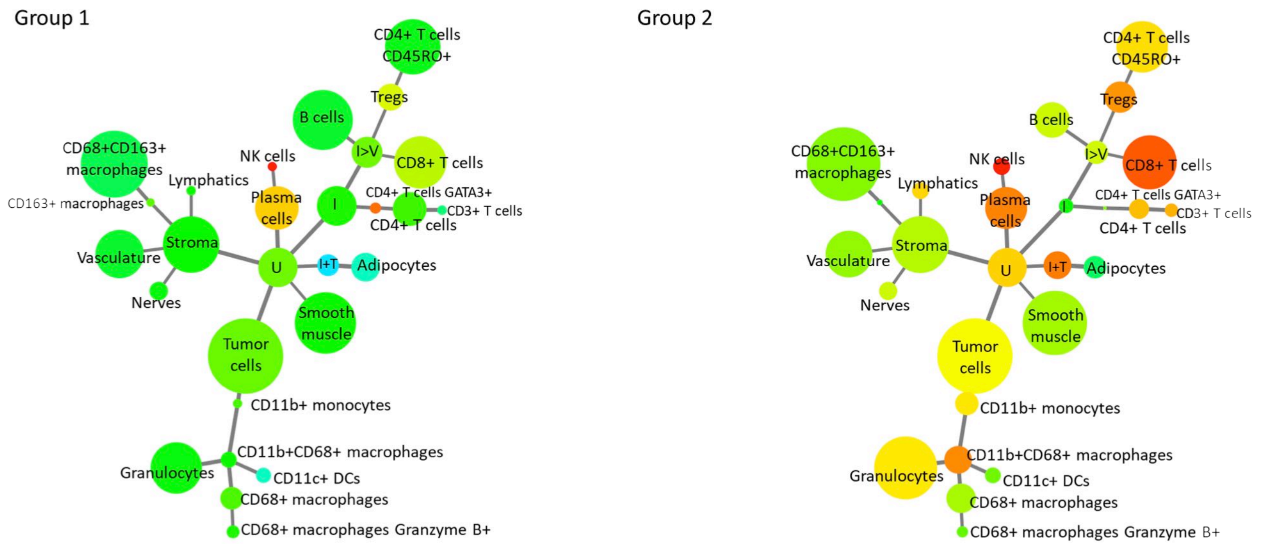

### MMP-9

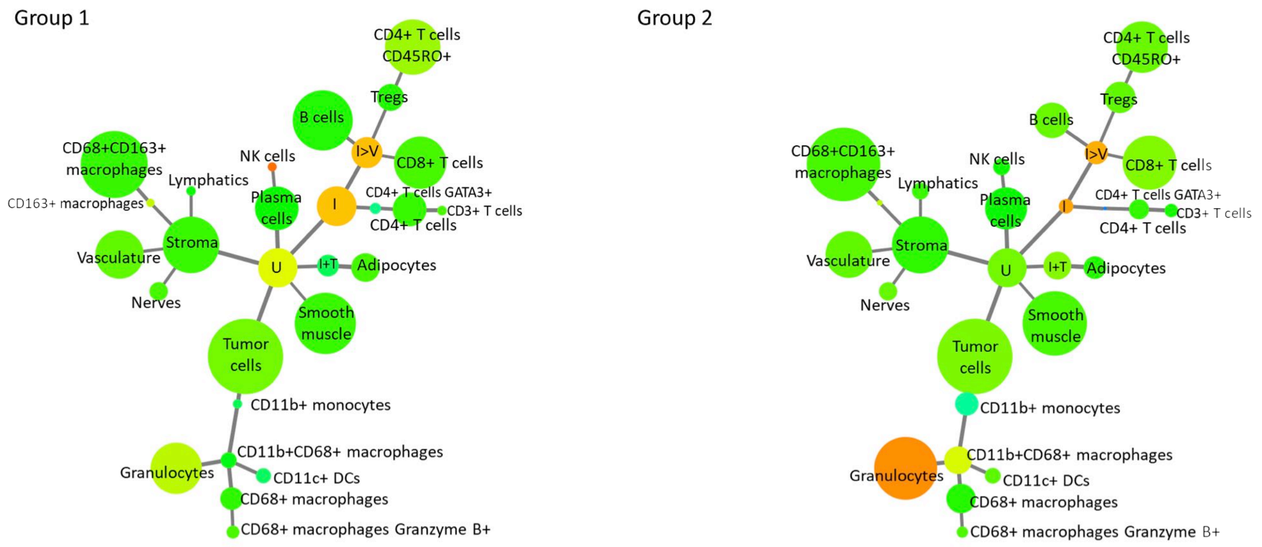

### MUC-1

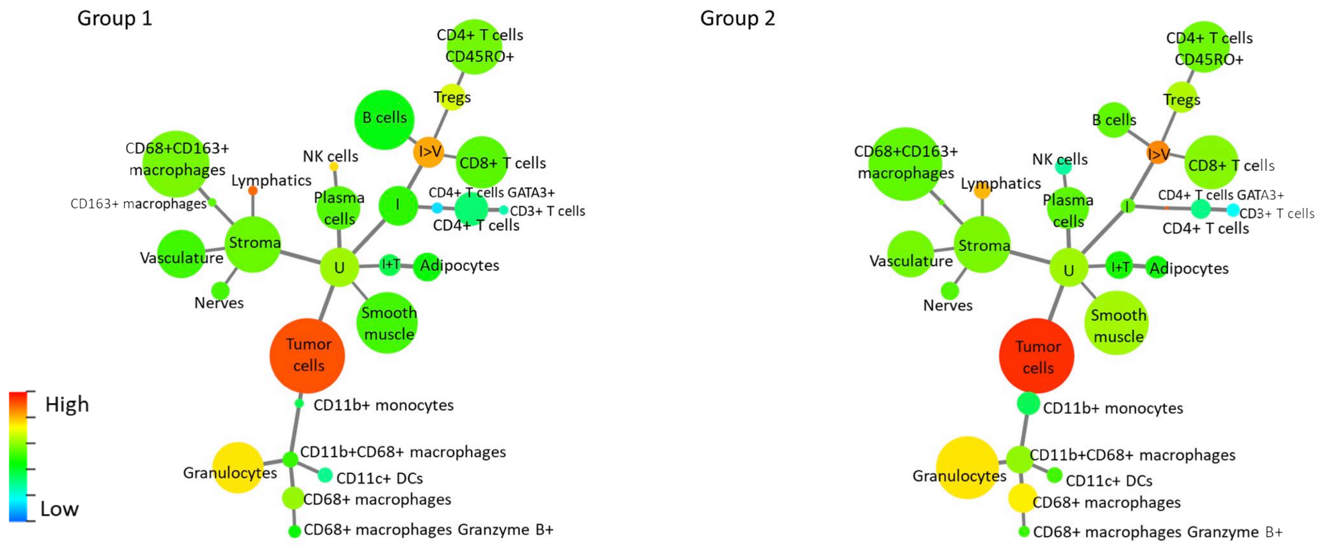

### Na-K-ATPase

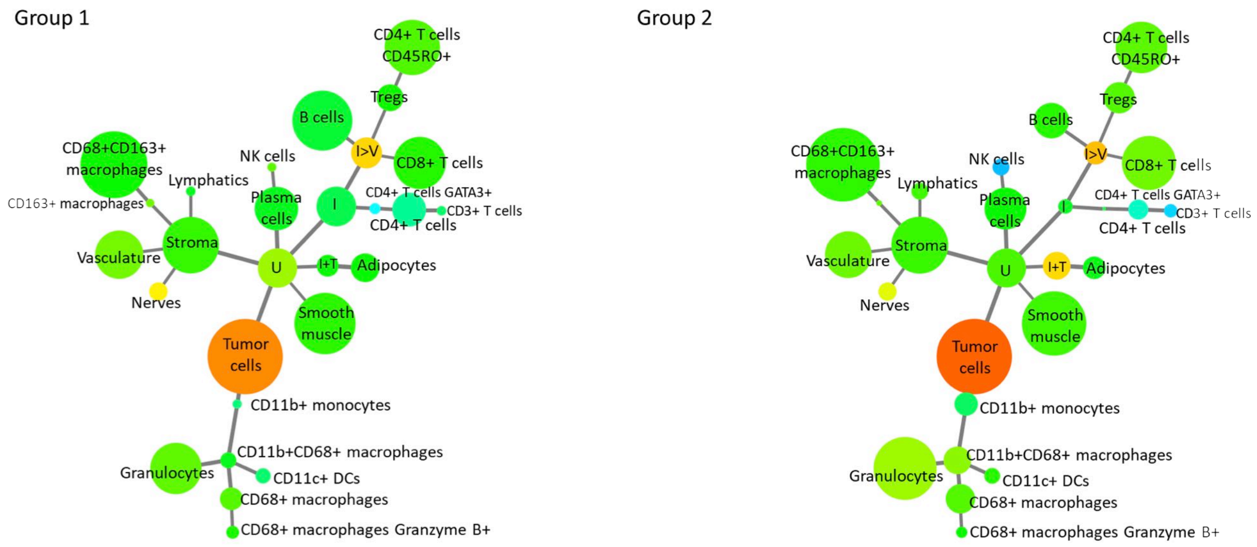

# p53

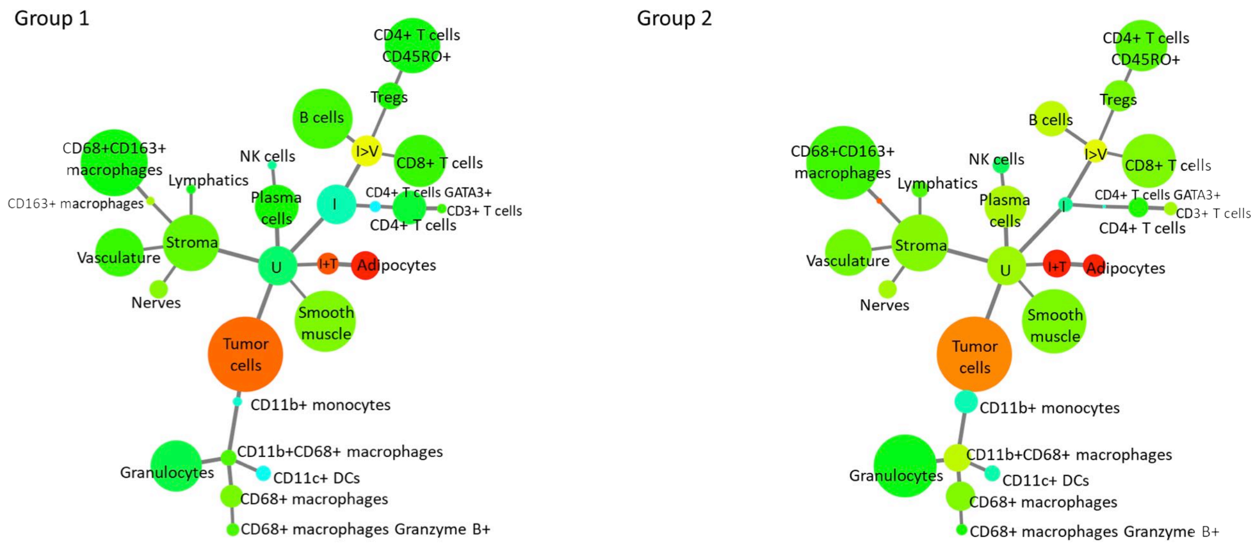

# PD-1

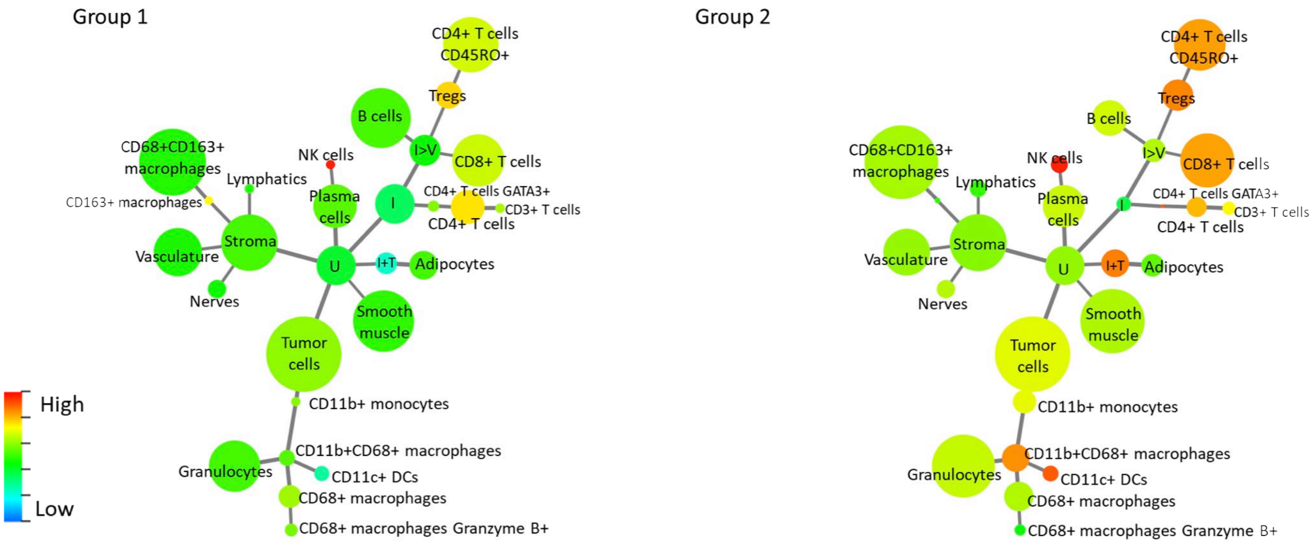

# PD-L1

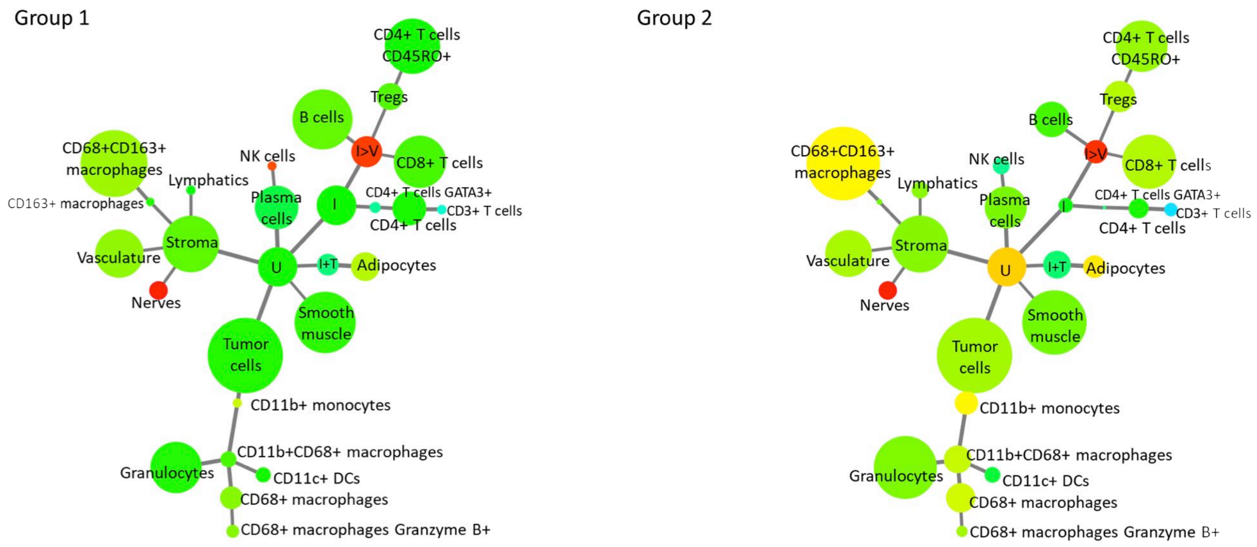

### Podoplanin

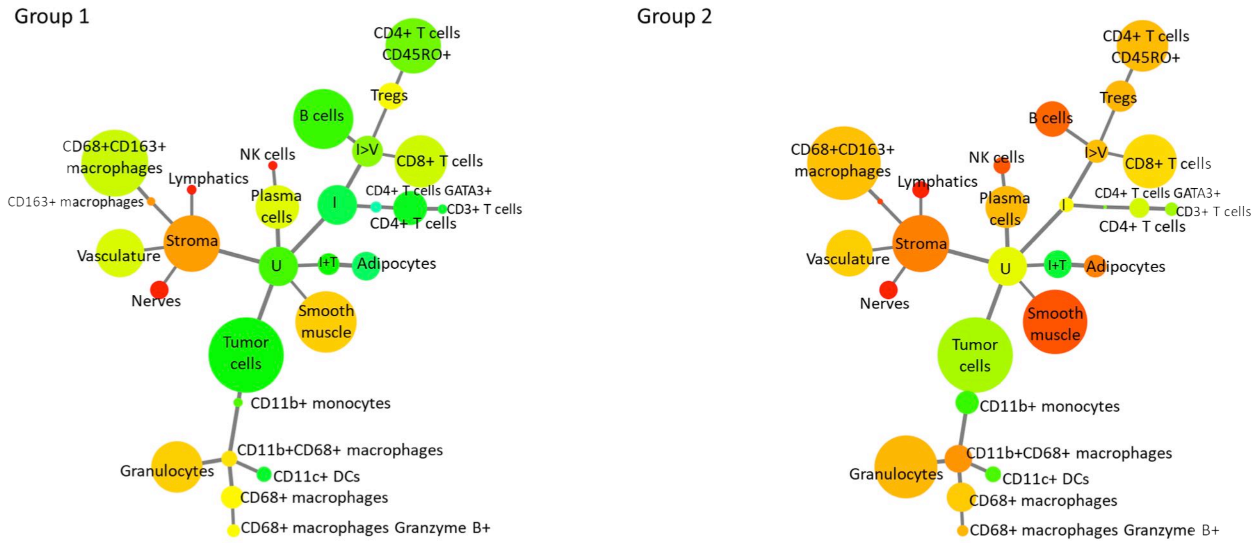

### Smooth Muscle Actin (SMA)

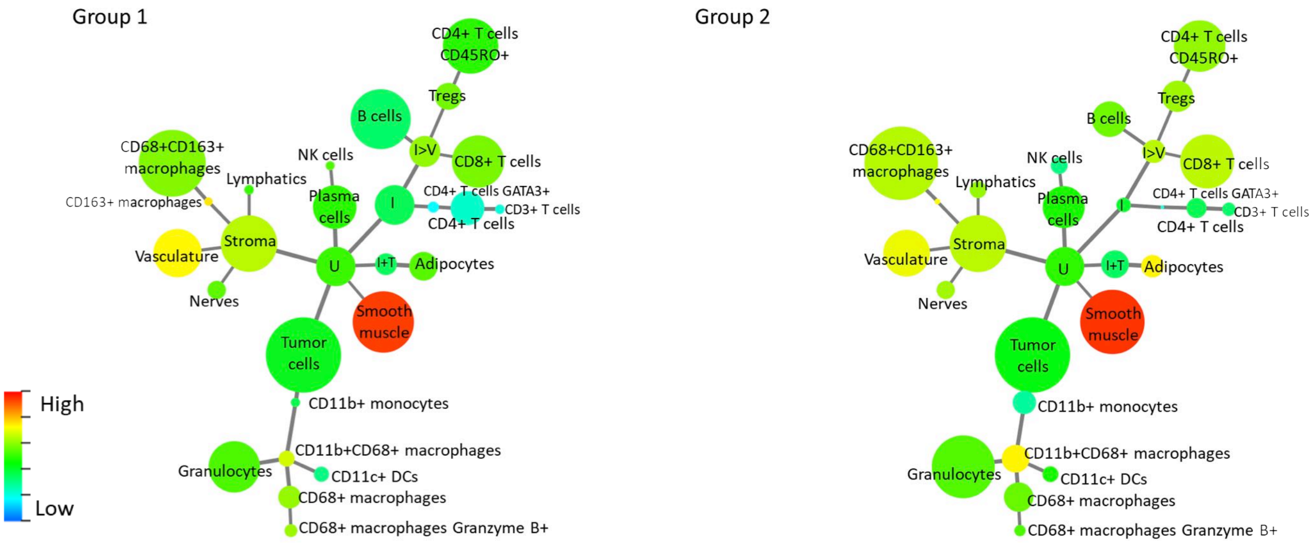

### Synaptophysin

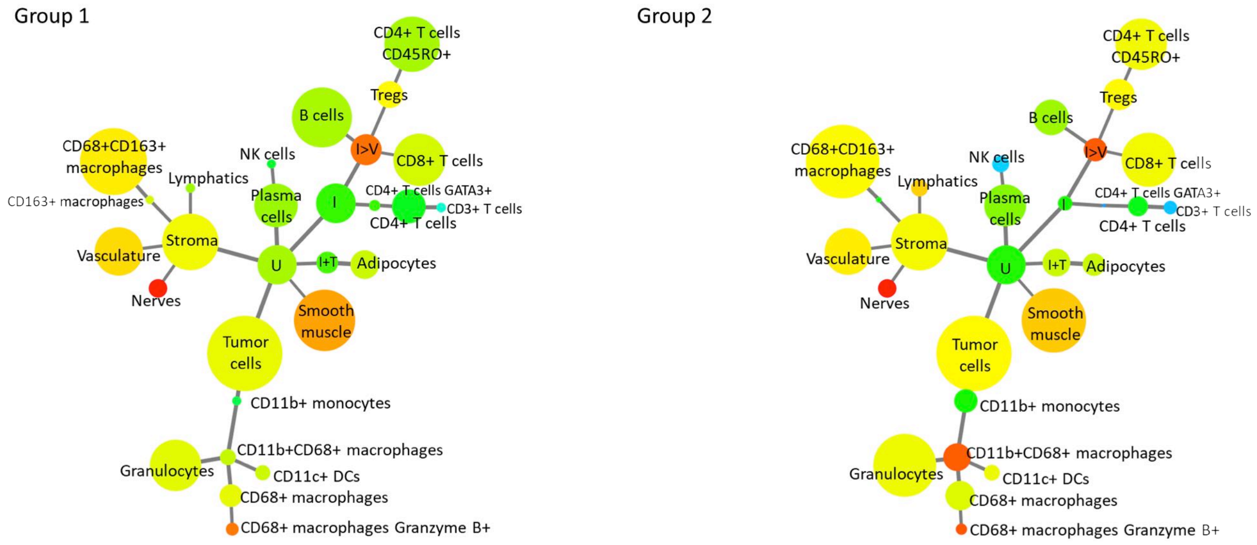

### T-bet

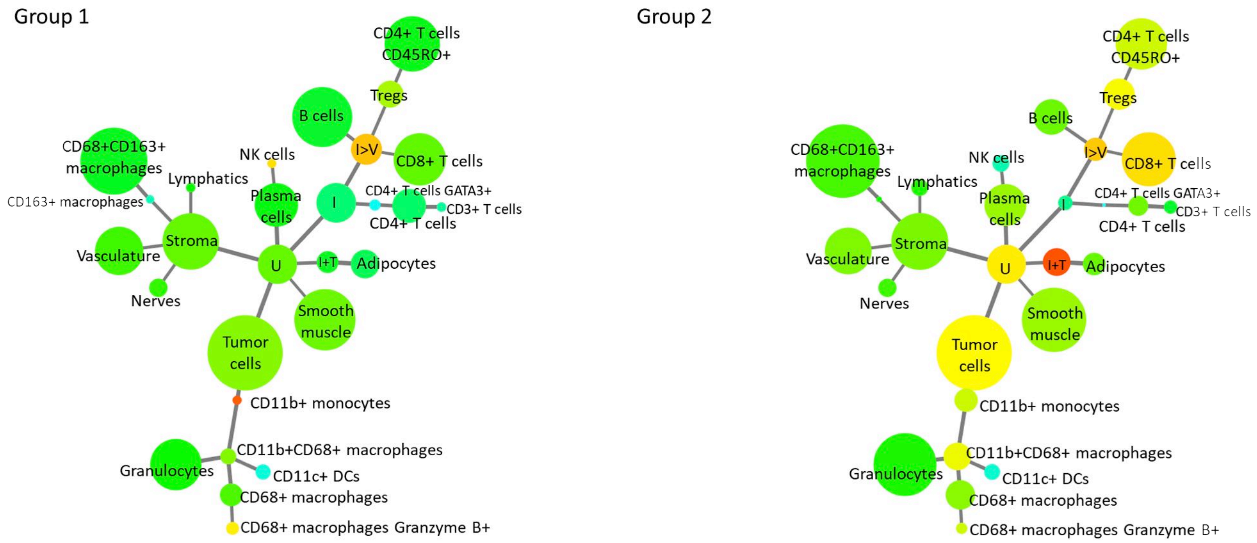

### Vimentin

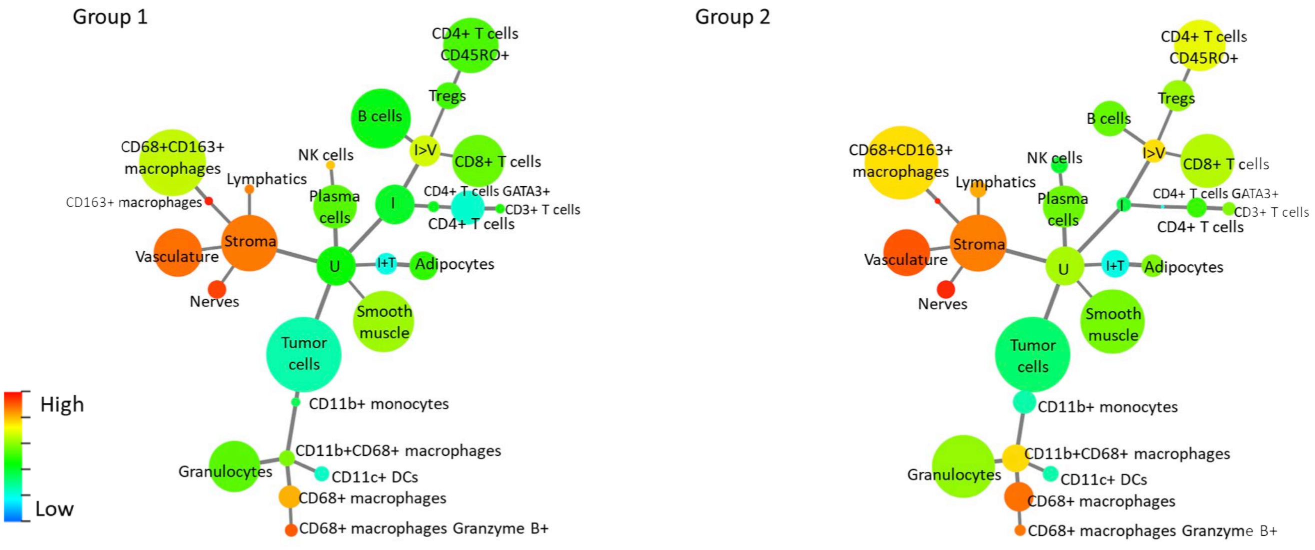

### VISTA

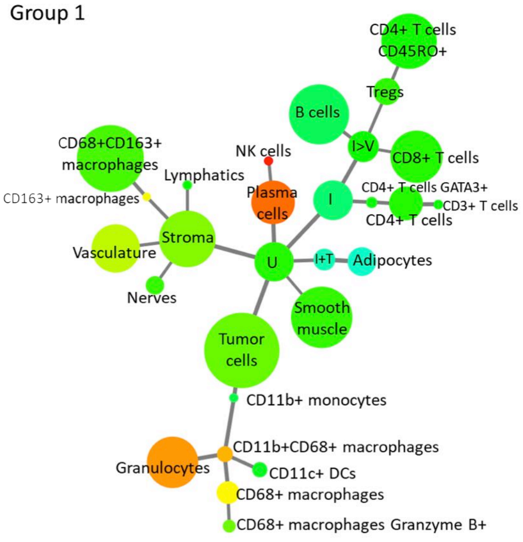

Group 1
CD4+ T cells CD45RO+
Tregs
B cells
I>V
CD8+ T cells
CD68+CD163+ macrophages
NK cells
Lymphatics
Plasma cells
I
CD4+ T cells GATA3+
CD3+ T cells
CD4+ T cells
CD163+ macrophages
Stroma
Vasculature
U
I+T
Adipocytes
Nerves
Smooth muscle
Tumor cells
CD11b+ monocytes
CD11b+CD68+ macrophages
Granulocytes
CD11c+ DCs
CD68+ macrophages
CD68+ macrophages Granzyme B+

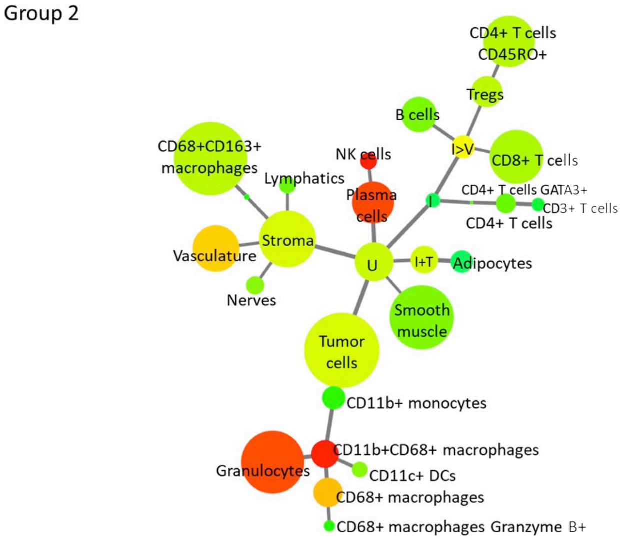

Group 2
CD4+ T cells CD45RO+
Tregs
B cells
I>V
CD68+CD163+ macrophages
NK cells
CD8+ T cells
Lymphatics
Plasma cells
I
CD4+ T cells GATA3+
CD3+ T cells
CD4+ T cells
Vasculature
Stroma
U
I+T
Adipocytes
Nerves
Smooth muscle
Tumor cells
CD11b+ monocytes
CD11b+CD68+ macrophages
Granulocytes
CD11c+ DCs
CD68+ macrophages
CD68+ macrophages Granzyme B+

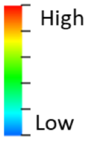

High
Low

**Figure S12**

**A**

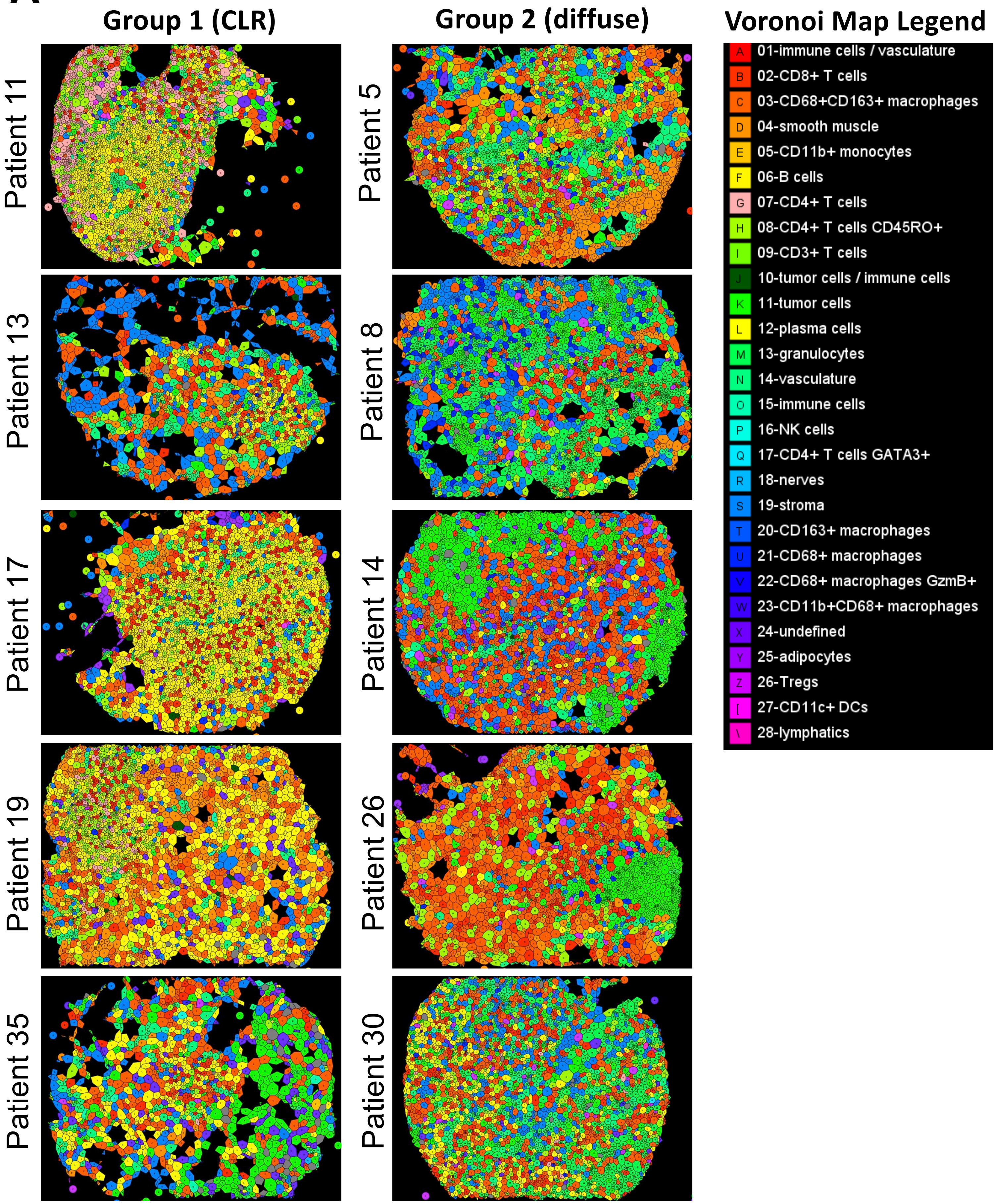
Group 1 (CLR)
Group 2 (diffuse)
Voronoi Map Legend
Patient 11
Patient 13
Patient 17
Patient 19
Patient 35
Patient 5
Patient 8
Patient 14
Patient 26
Patient 30
A 01-immune cells / vasculature
B 02-CD8+ T cells
C 03-CD68+CD163+ macrophages
D 04-smooth muscle
E 05-CD11b+ monocytes
F 06-B cells
G 07-CD4+ T cells
H 08-CD4+ T cells CD45RO+
I 09-CD3+ T cells
J 10-tumor cells / immune cells
K 11-tumor cells
L 12-plasma cells
M 13-granulocytes
N 14-vasculature
O 15-immune cells
P 16-NK cells
Q 17-CD4+ T cells GATA3+
R 18-nerves
S 19-stroma
T 20-CD163+ macrophages
U 21-CD68+ macrophages
V 22-CD68+ macrophages GzmB+
W 23-CD11b+CD68+ macrophages
X 24-undefined
Y 25-adipocytes
Z 26-Tregs
[ 27-CD11c+ DCs
\ 28-lymphatics

**B**

| | All CRC Patients (n=251,028 cells) Mean % ± SD | Group 1 (CLR) (n=107,966 cells) Mean % ± SD | Group 2 (Diffuse) (n=143,062 cells) Mean % ± SD | Mann-Whitney Test p-value |
|---|---|---|---|---|
| Tumor cells | 19.3 ± 14.4% | 19.3 ± 18.9% | 19.3 ± 9.1% | 0.3141 |
| CD68+CD163+ macrophages | 15.6 ± 7.1% | 13.4 ± 6.4% | 17.6 ± 7.2% | 0.0831 |
| Smooth muscle | 10.2 ± 8.4% | 9.5 ± 7.0% | 10.9 ± 9.8% | 0.9605 |
| **Granulocytes** | **8.5 ± 8.2%** | **5.8 ± 5.3%** | **11.0 ± 9.7%** | **0.0496** |
| Stroma | 8.0 ± 4.3% | 7.9 ± 4.7% | 8.0 ± 4.1% | 0.9080 |
| CD4+ T cells CD45RO+ | 7.0 ± 5.0% | 7.5 ± 5.5% | 6.6 ± 4.6% | 0.7043 |
| CD8+ T cells | 7.0 ± 4.3% | 6.0 ± 2.9% | 7.9 ± 5.3% | 0.6322 |
| **B cells** | **5.1 ± 7.5%** | **8.8 ± 9.4%** | **1.7 ± 2.3%** | **0.0009** |
| Vasculature | 4.5 ± 2.2% | 4.8 ± 2.4% | 4.2 ± 1.9% | 0.4780 |
| Plasma cells | 3.3 ± 4.0% | 3.8 ± 4.0% | 2.9 ± 4.2% | 0.2038 |
| Undefined | 2.9 ± 3.8% | 2.9 ± 2.9% | 3.0 ± 4.6% | 0.2283 |
| **Immune cells (mixed)** | **1.5 ± 4.5%** | **3.0 ± 6.2%** | **0.1 ± 0.2%** | **0.0306** |
| **Tregs** | **1.1 ± 0.7%** | **0.9 ± 0.6%** | **1.4 ± 0.7%** | **0.0218** |
| **CD4+ T cells** | **1.0 ± 2.5%** | **1.8 ± 3.4%** | **0.3 ± 0.7%** | **0.0167** |
| Immmune cells / vasculature | 1.0 ± 1.8% | 1.5 ± 2.5% | 0.5 ± 0.8% | 0.1923 |
| **Adipocytes** | **0.8 ± 1.1%** | **1.2 ± 1.0%** | **0.5 ± 1.1%** | **0.0004** |
| **CD68+ macrophages** | **0.8 ± 1.0%** | **0.5 ± 0.5%** | **1.1 ± 1.3%** | **0.0333** |
| **Tumor cells / immune cells** | **0.6 ± 2.0%** | **0.5 ± 1.0%** | **0.8 ± 2.6%** | **0.0127** |
| **CD11b+CD68+ macrophages** | **0.5 ± 1.4%** | **0.2 ± 0.1%** | **0.9 ± 2.0%** | **0.0459** |
| CD11b+ monocytes | 0.3 ± 1.5% | 0.0 ± 0.1% | 0.6 ± 2.1% | 0.4485 |
| CD11c+ DCs | 0.2 ± 0.4% | 0.2 ± 0.2% | 0.2 ± 0.5% | 0.1326 |
| Nerves | 0.2 ± 0.3% | 0.2 ± 0.3% | 0.2 ± 0.3% | 0.9605 |
| NK cells | 0.1 ± 0.4% | 0.0 ± 0.1% | 0.2 ± 0.5% | 0.5323 |
| Lymphatics | 0.1 ± 0.3% | 0.0 ± 0.1% | 0.2 ± 0.4% | 0.2452 |
| CD3+ T cells | 0.1 ± 0.2% | 0.0 ± 0.1% | 0.1 ± 0.3% | 0.6111 |
| CD68+ macrophages GzmB+ | 0.1 ± 0.1% | 0.1 ± 0.1% | 0.1 ± 0.1% | 0.7965 |
| CD4+ T cells GATA3+ | 0.0 ± 0.2% | 0.1 ± 0.2% | 0.0 ± 0.0% | 0.3029 |
| CD163+ macrophages | 0.0 ± 0.0% | 0.0 ± 0.1% | 0.0 ± 0.0% | 0.4905 |

C

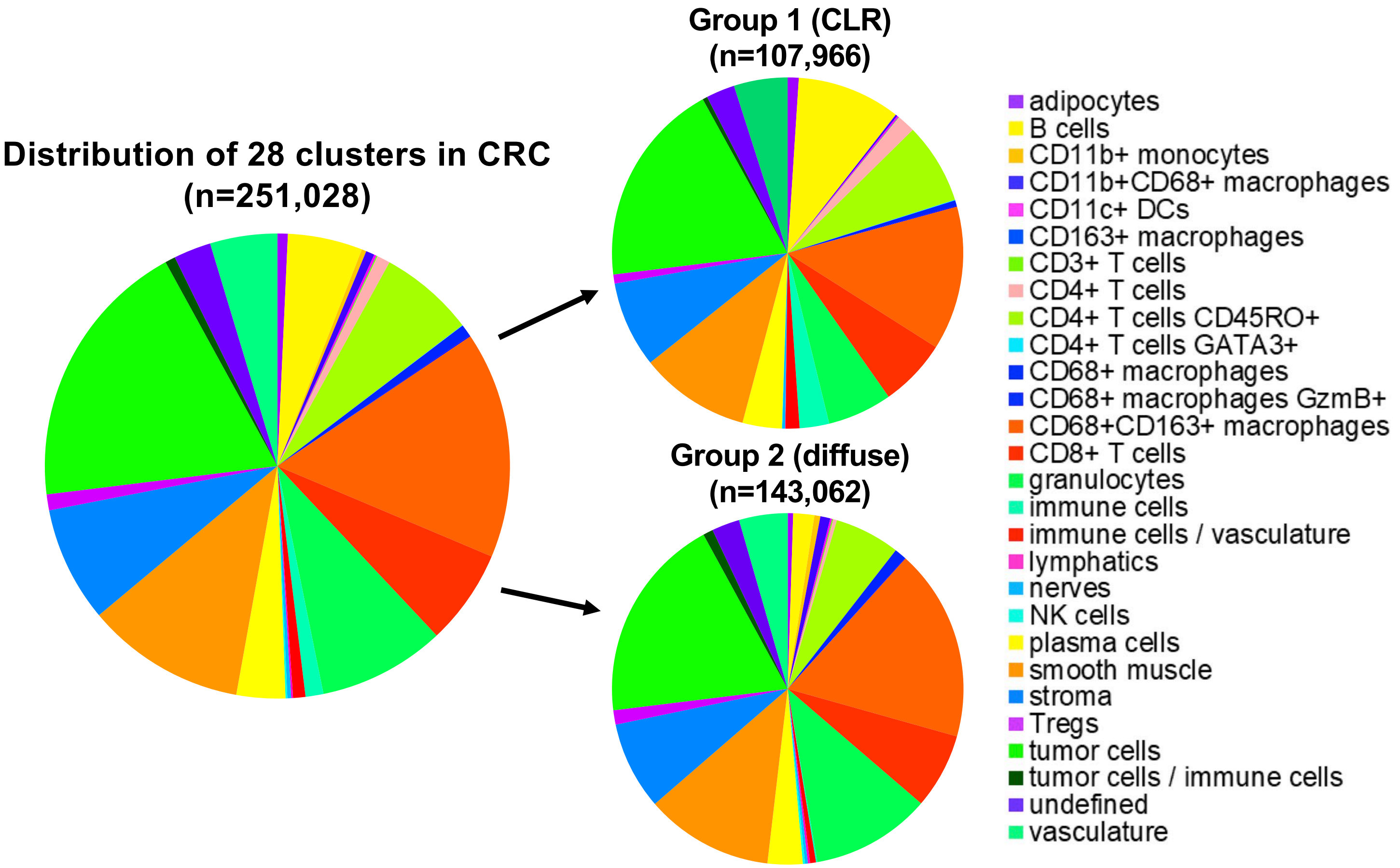

Distribution of 28 clusters in CRC
(n=251,028)
Group 1 (CLR)
(n=107,966)
Group 2 (diffuse)
(n=143,062)
adipocytes
B cells
CD11b+ monocytes
CD11b+CD68+ macrophages
CD11c+ DCs
CD163+ macrophages
CD3+ T cells
CD4+ T cells
CD4+ T cells CD45RO+
CD4+ T cells GATA3+
CD68+ macrophages
CD68+ macrophages GzmB+
CD68+CD163+ macrophages
CD8+ T cells
granulocytes
immune cells
immune cells / vasculature
lymphatics
nerves
NK cells
plasma cells
smooth muscle
stroma
Tregs
tumor cells
tumor cells / immune cells
undefined
vasculature

D

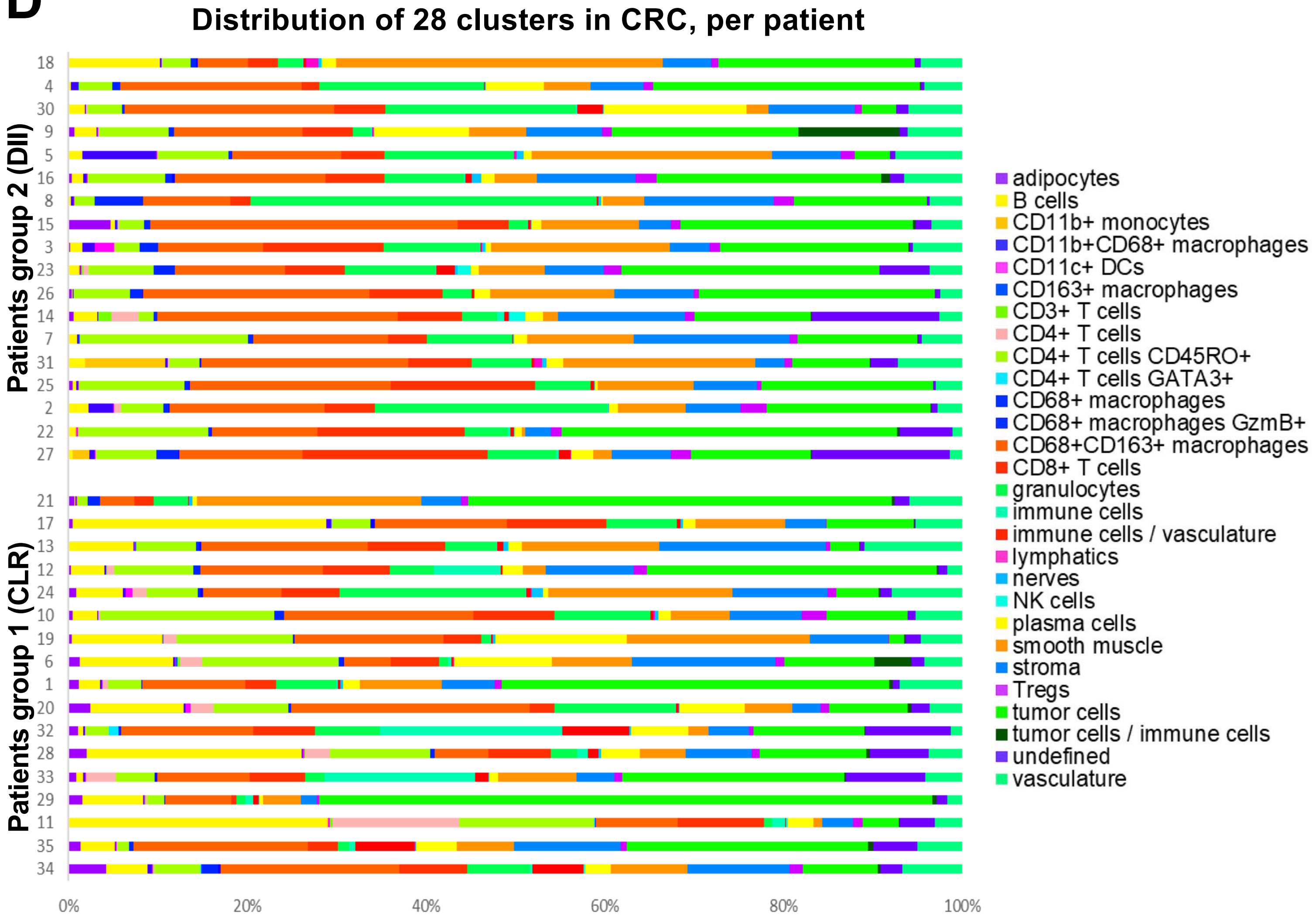

Distribution of 28 clusters in CRC, per patient
Patients group 2 (DII)
Patients group 1 (CLR)
18
4
30
9
5
16
8
15
3
23
26
14
7
31
25
2
22
27
21
17
13
12
24
10
19
6
1
20
32
28
33
29
11
35
34
0%
20%
40%
60%
80%
100%
adipocytes
B cells
CD11b+ monocytes
CD11b+CD68+ macrophages
CD11c+ DCs
CD163+ macrophages
CD3+ T cells
CD4+ T cells
CD4+ T cells CD45RO+
CD4+ T cells GATA3+
CD68+ macrophages
CD68+ macrophages GzmB+
CD68+CD163+ macrophages
CD8+ T cells
granulocytes
immune cells
immune cells / vasculature
lymphatics
nerves
NK cells
plasma cells
smooth muscle
stroma
Tregs
tumor cells
tumor cells / immune cells
undefined
vasculature

### Figure S13

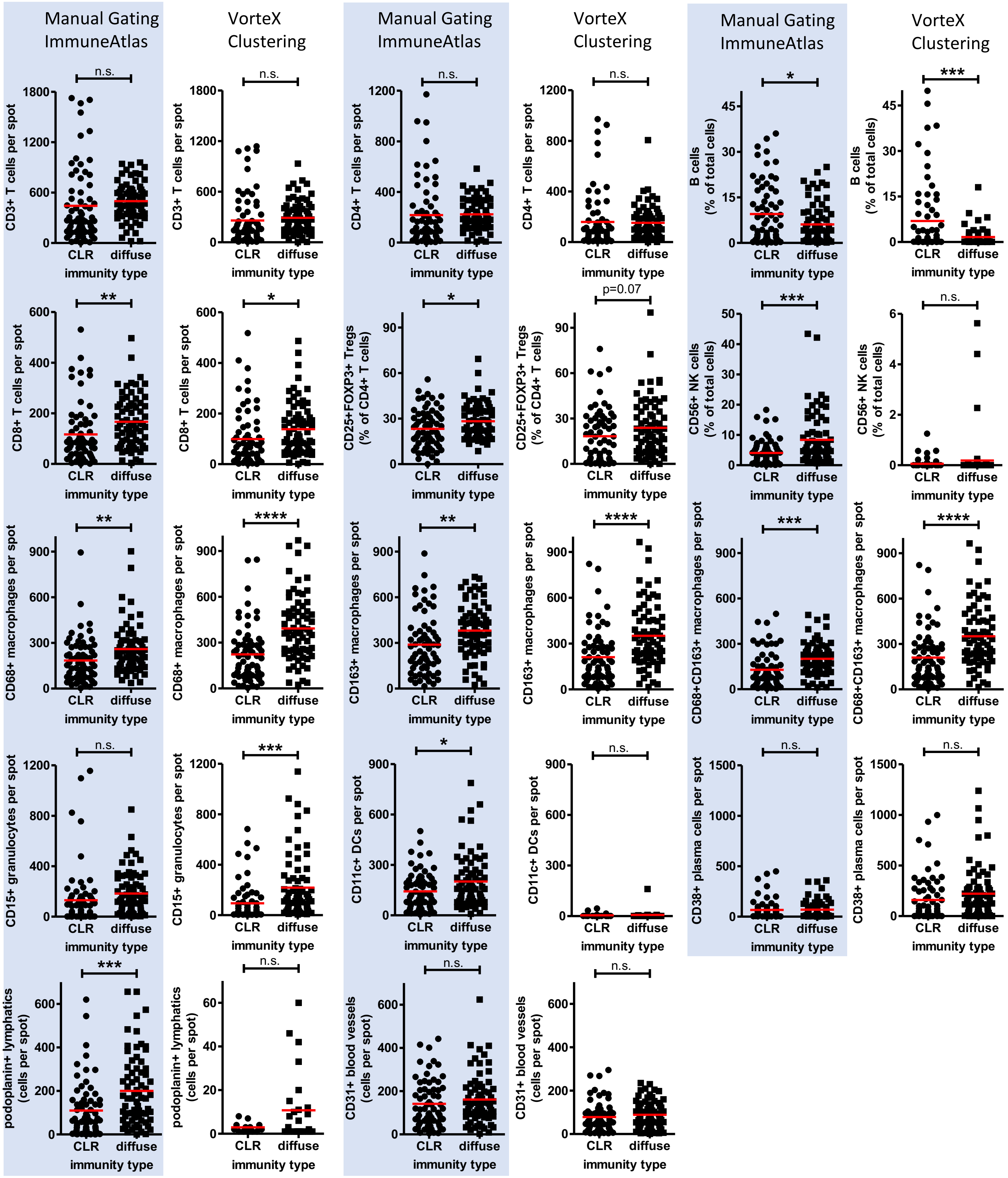

Manual Gating ImmuneAtlas
VorteX Clustering
Manual Gating ImmuneAtlas
VorteX Clustering
Manual Gating ImmuneAtlas
VorteX Clustering
CD3+ T cells per spot
CD4+ T cells per spot
B cells (% of total cells)
CD8+ T cells per spot
CD25+FOXP3+ Tregs (% of CD4+ T cells)
CD56+ NK cells (% of total cells)
CD68+ macrophages per spot
CD163+ macrophages per spot
CD68+CD163+ macrophages per spot
CD15+ granulocytes per spot
CD11c+ DCs per spot
CD38+ plasma cells per spot
podoplanin+ lymphatics (cells per spot)
CD31+ blood vessels (cells per spot)
CLR
diffuse
immunity type
n.s.
*
**
***
****
p=0.07

### Figure S14

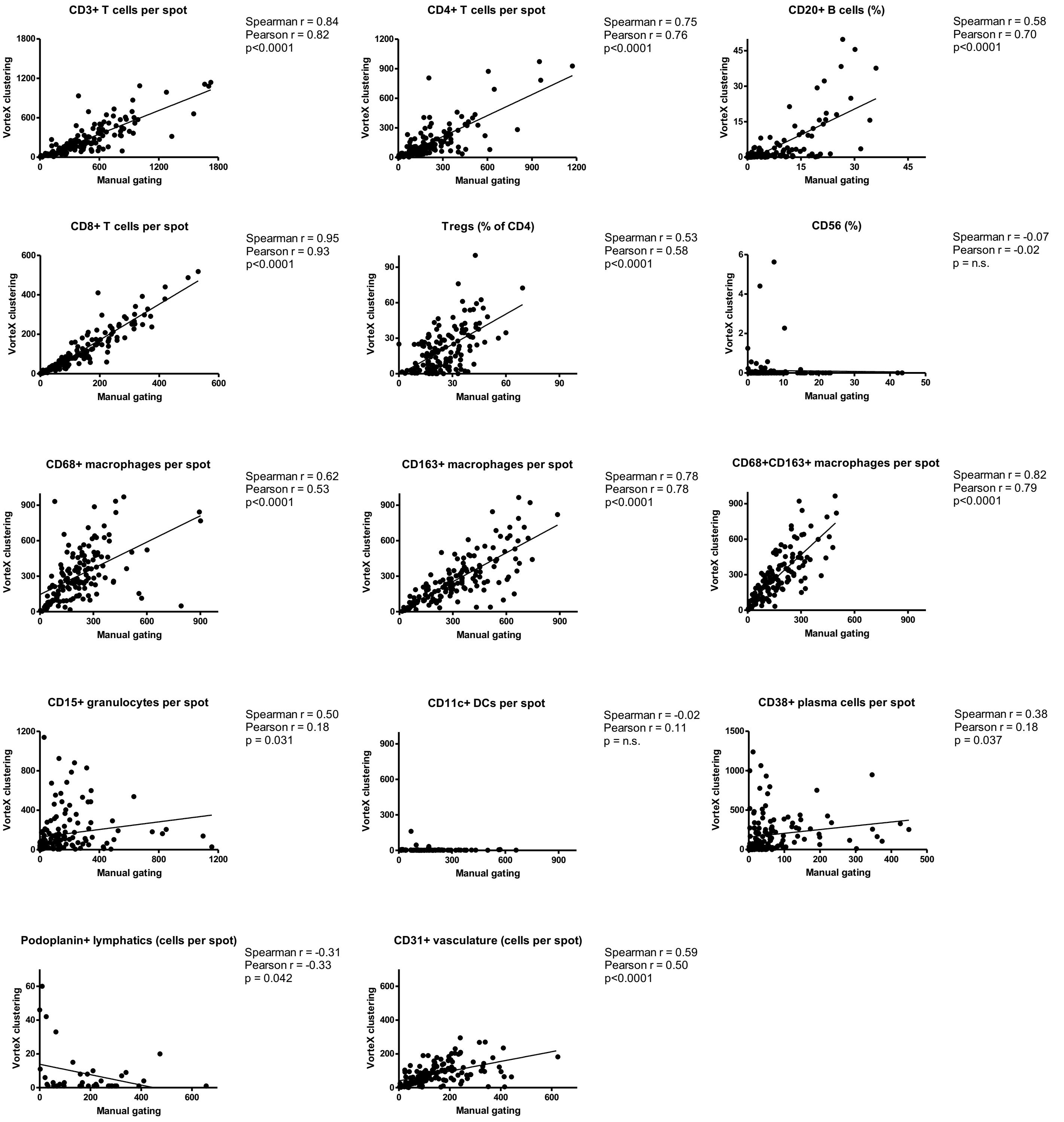

CD3+ T cells per spot
Spearman r = 0.84
Pearson r = 0.82
p<0.0001
CD4+ T cells per spot
Spearman r = 0.75
Pearson r = 0.76
p<0.0001
CD20+ B cells (%)
Spearman r = 0.58
Pearson r = 0.70
p<0.0001
CD8+ T cells per spot
Spearman r = 0.95
Pearson r = 0.93
p<0.0001
Tregs (% of CD4)
Spearman r = 0.53
Pearson r = 0.58
p<0.0001
CD56 (%)
Spearman r = -0.07
Pearson r = -0.02
p = n.s.
CD68+ macrophages per spot
Spearman r = 0.62
Pearson r = 0.53
p<0.0001
CD163+ macrophages per spot
Spearman r = 0.78
Pearson r = 0.78
p<0.0001
CD68+CD163+ macrophages per spot
Spearman r = 0.82
Pearson r = 0.79
p<0.0001
CD15+ granulocytes per spot
Spearman r = 0.50
Pearson r = 0.18
p = 0.031
CD11c+ DCs per spot
Spearman r = -0.02
Pearson r = 0.11
p = n.s.
CD38+ plasma cells per spot
Spearman r = 0.38
Pearson r = 0.18
p = 0.037
Podoplanin+ lymphatics (cells per spot)
Spearman r = -0.31
Pearson r = -0.33
p = 0.042
CD31+ vasculature (cells per spot)
Spearman r = 0.59
Pearson r = 0.50
p<0.0001
VorteX clustering
Manual gating

#### Supplemental Figure 15

**A**

| | All CRC Patients (n=131,329 immune cells) Mean % ± SD | Group 1 (CLR) (n=56,343 immune cells) Mean % ± SD | Group 2 (Diffuse) (n=74,986 immune cells) Mean % ± SD | Mann-Whitney Test p-value |
|---|---|---|---|---|
| **B cells** | **9.5 ± 12.2%** | **15.3 ± 13.5%** | **4.0 ± 7.9%** | **0.0008** |
| **Macrophages** | **33.5 ± 13.2%** | **28.4 ± 11.2%** | **38.4 ± 13.5%** | **0.0496** |
| Tregs | 2.2 ± 1.3% | 1.8 ± 1.3% | 2.6 ± 1.2% | 0.0535 |
| CD11c+ DCs | 0.4 ± 0.8% | 0.4 ± 0.4% | 0.4 ± 1.1% | 0.0512 |
| CD4+ T cells | 14.7 ± 9.0% | 16.2 ± 9.5% | 13.3 ± 8.6% | 0.3641 |
| CD8+ T cells | 13.1 ± 7.3% | 11.2 ± 3.9% | 15.0 ± 9.2% | 0.4187 |
| NK cells | 0.4 ± 1.0% | 0.1 ± 0.1% | 0.8 ± 1.4% | 0.7197 |
| Other immune cells | 26.3 ± 14.4% | 26.7 ± 13.3% | 25.9 ± 15.7% | 0.6322 |

**B**

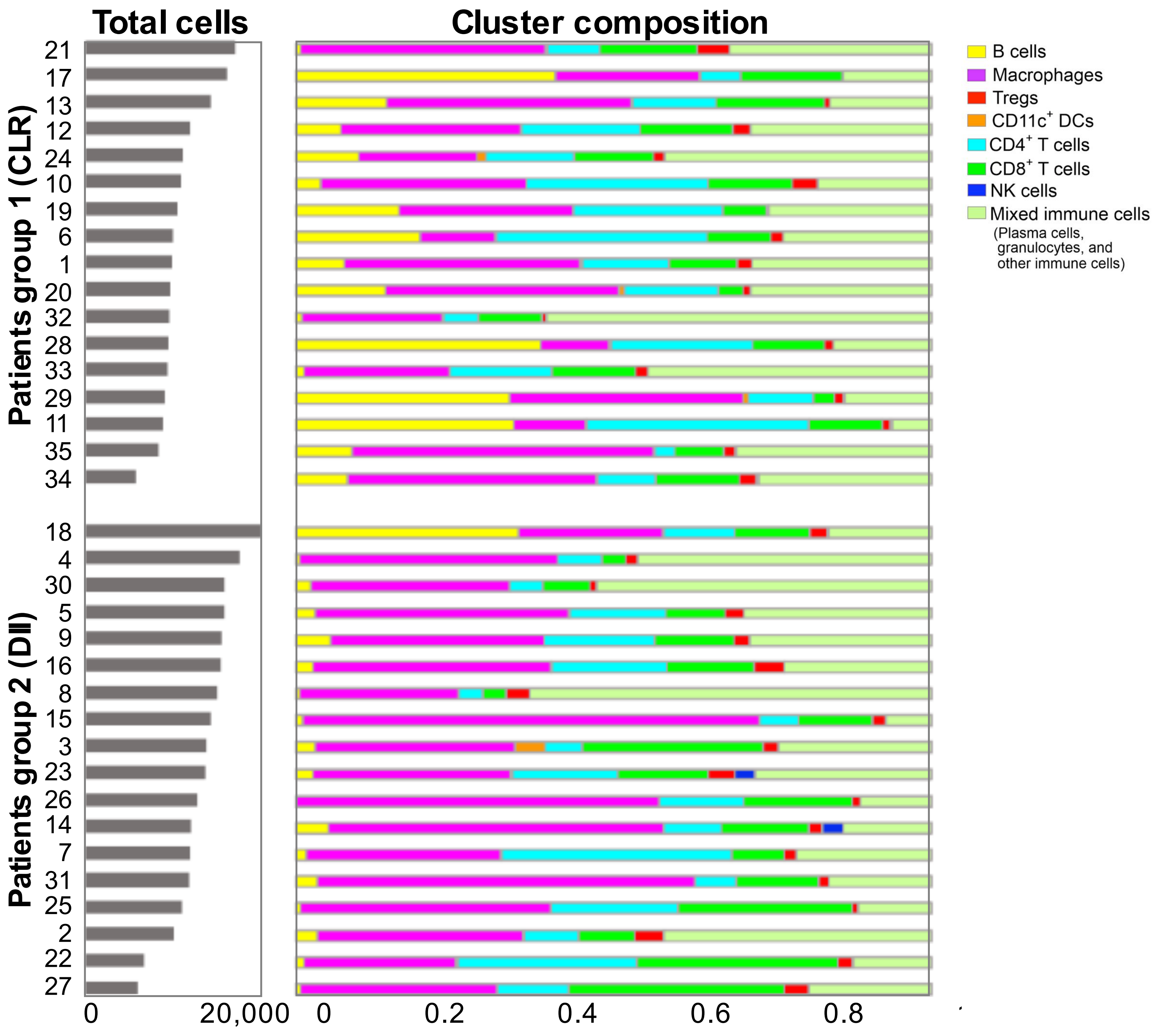

### Figure S16

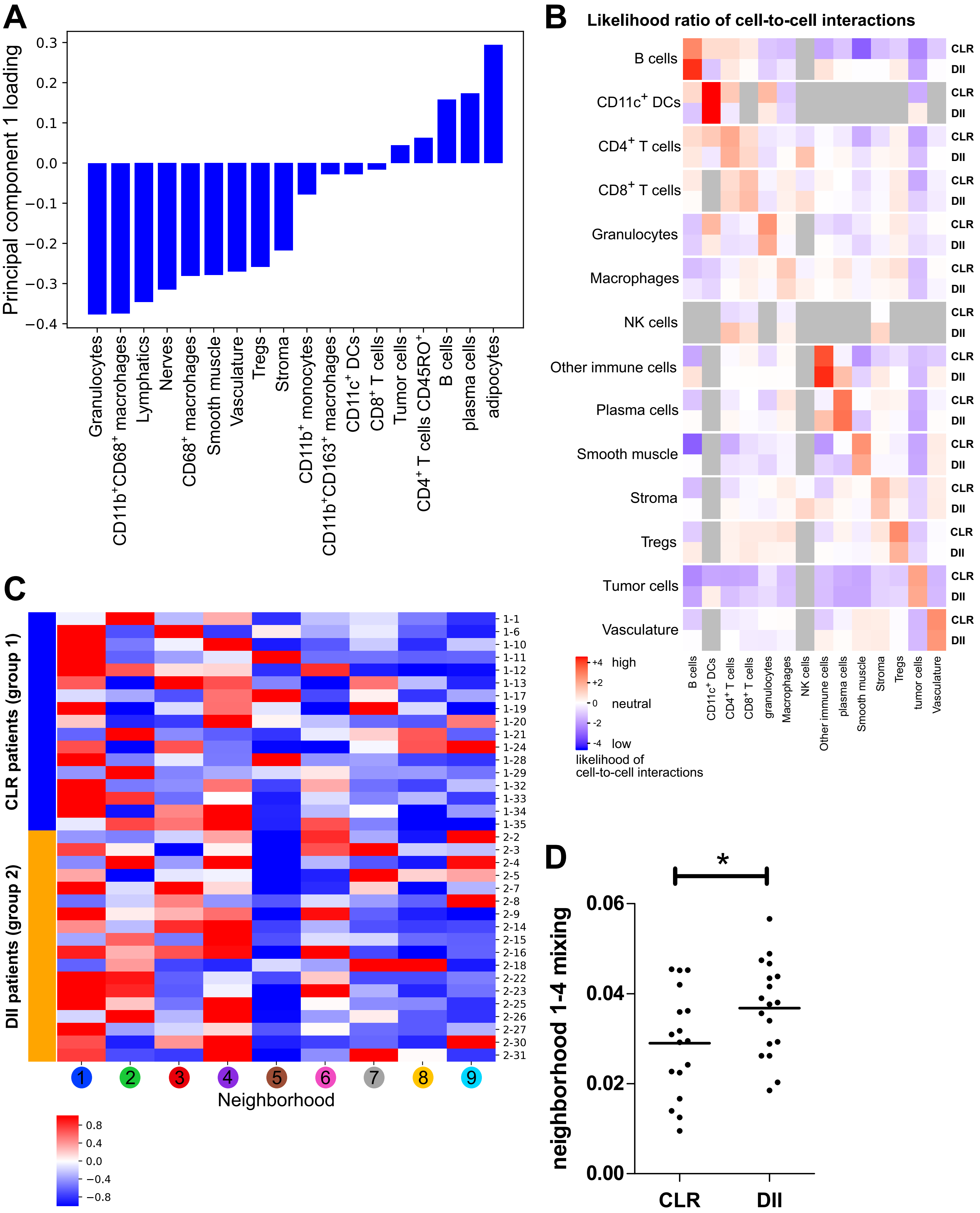
A
Principal component 1 loading
Granulocytes
CD11b+CD68+ macrophages
Lymphatics
Nerves
CD68+ macrophages
Smooth muscle
Vasculature
Tregs
Stroma
CD11b+ monocytes
CD11b+CD163+ macrophages
CD11c+ DCs
CD8+ T cells
Tumor cells
CD4+ T cells CD45RO+
B cells
plasma cells
adipocytes
B
Likelihood ratio of cell-to-cell interactions
B cells
CD11c+ DCs
CD4+ T cells
CD8+ T cells
Granulocytes
Macrophages
NK cells
Other immune cells
Plasma cells
Smooth muscle
Stroma
Tregs
Tumor cells
Vasculature
CLR
DII
high
neutral
low
likelihood of cell-to-cell interactions
C
CLR patients (group 1)
DII patients (group 2)
Neighborhood
D
neighborhood 1-4 mixing
CLR
DII

**Figure S17**

**A**

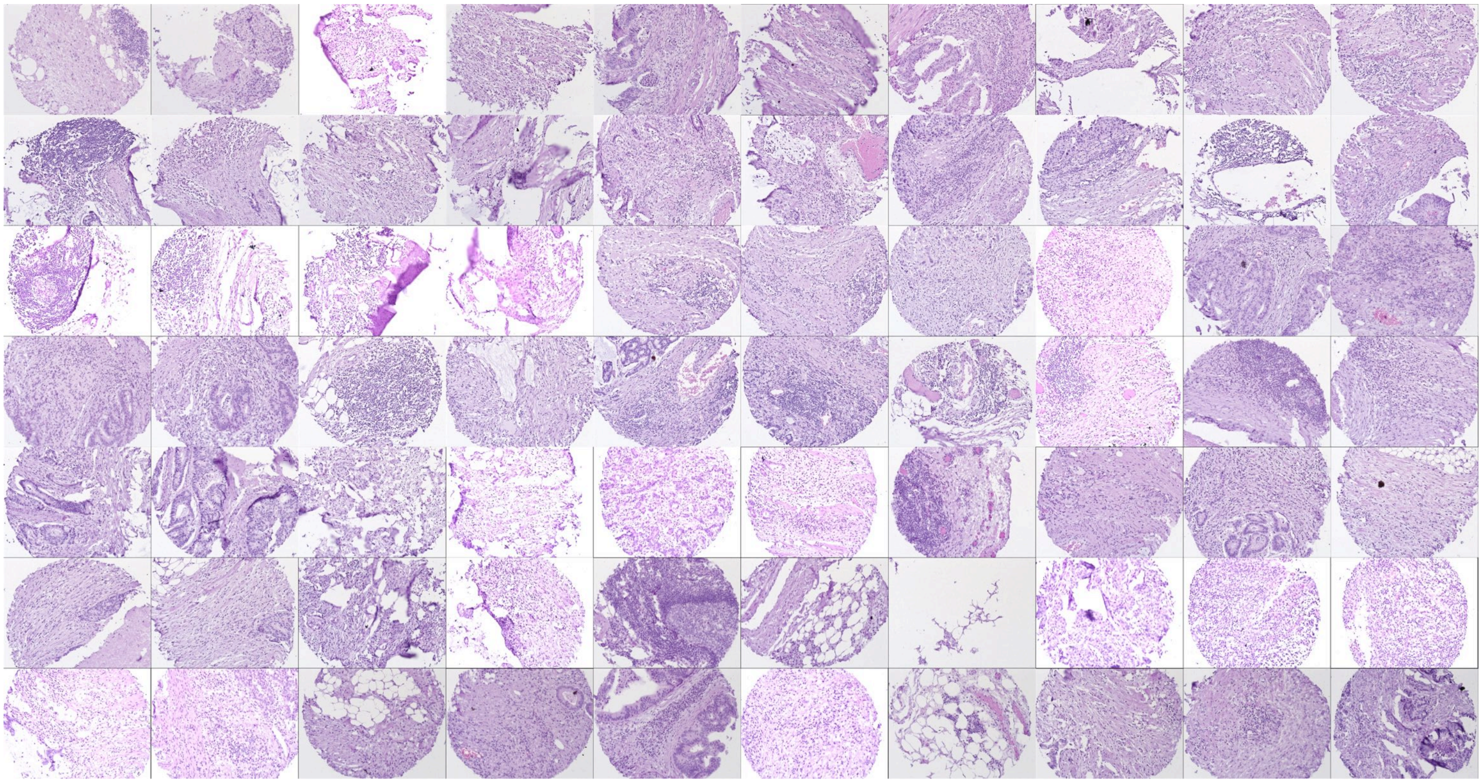

**B**

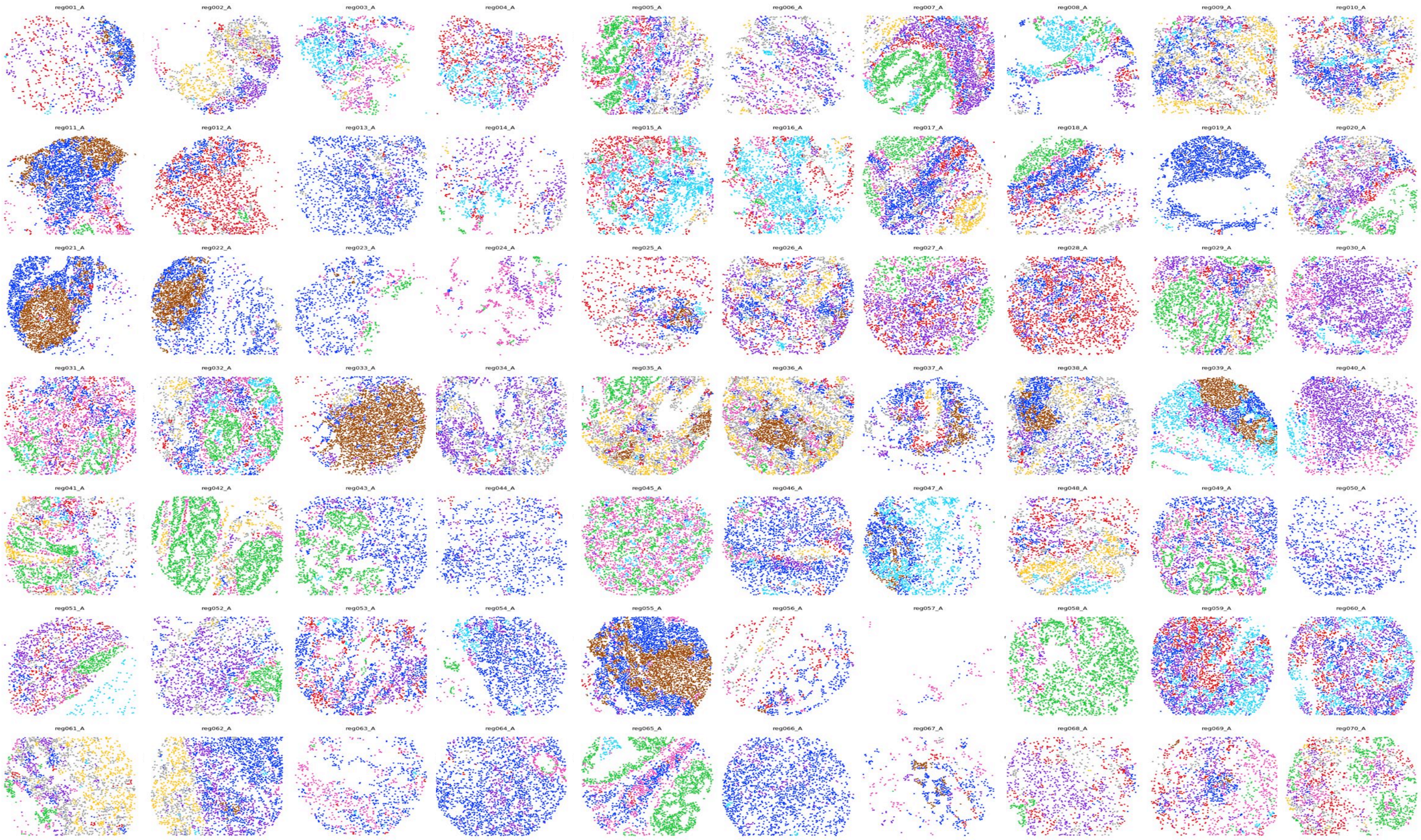

Neighborhood Legend

- **1** T cell enriched
- **2** main tumor
- **3** immune-infiltrated stroma
- **4** macrophage enriched
- **5** follicle
- **6** tumor boundary
- **7** vascular smooth muscle
- **8** smooth muscle
- **9** granulocyte enriched

**C**

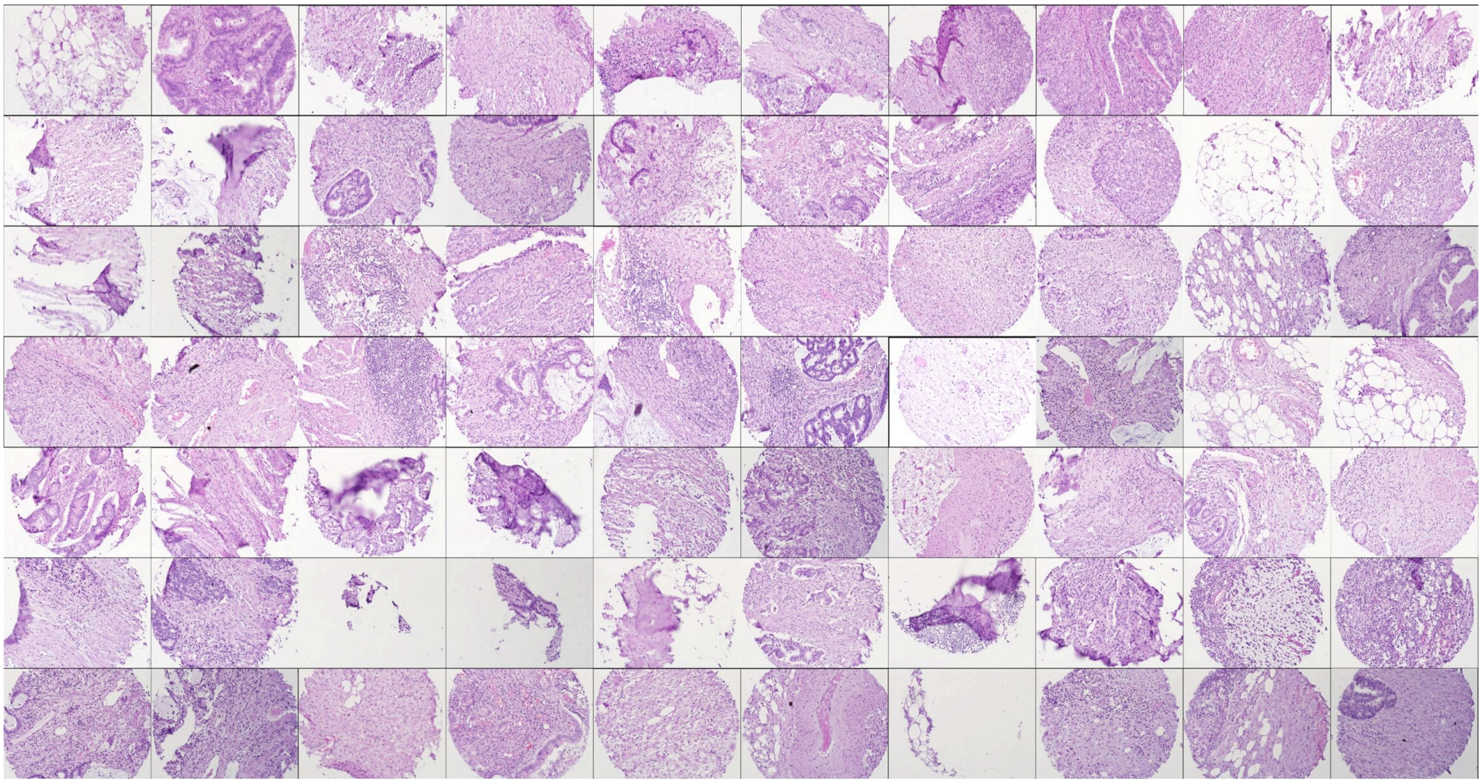

**D**

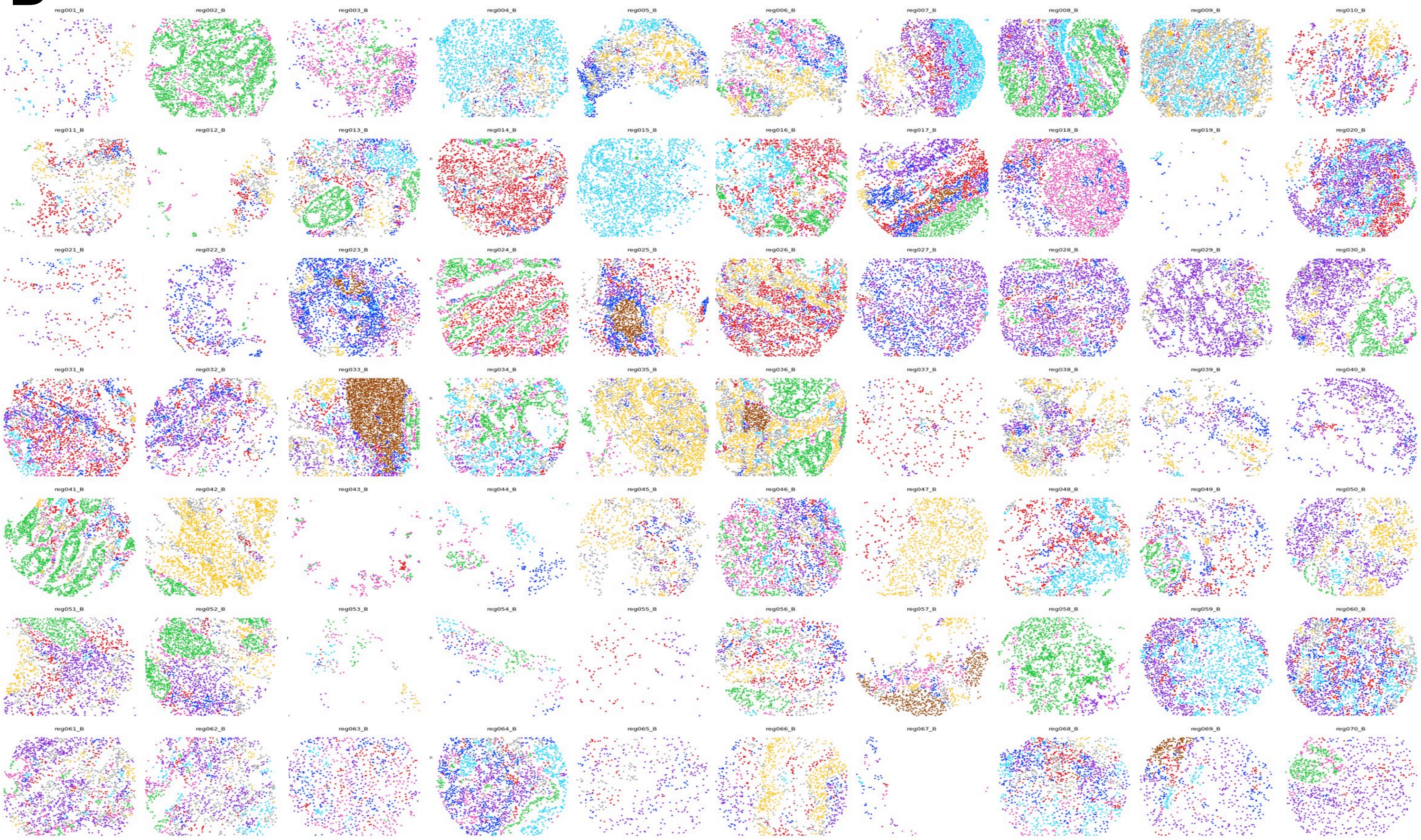

Neighborhood Legend

- **1** T cell enriched
- **2** main tumor
- **3** immune-infiltrated stroma
- **4** macrophage enriched
- **5** follicle
- **6** tumor boundary
- **7** vascular smooth muscle
- **8** smooth muscle
- **9** granulocyte enriched

### Figure S18

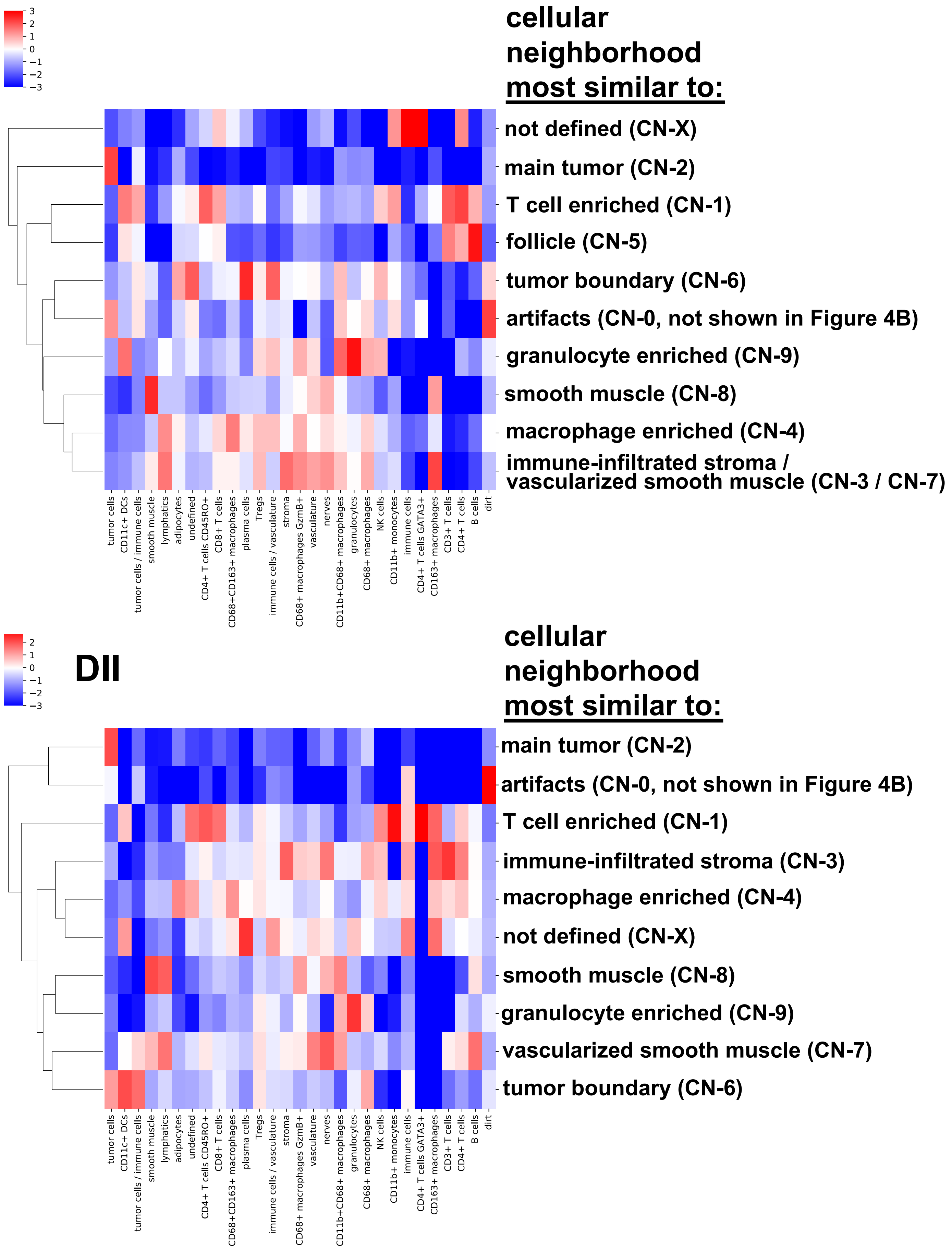

3
2
1
0
−1
−2
−3
cellular neighborhood most similar to:
not defined (CN-X)
main tumor (CN-2)
T cell enriched (CN-1)
follicle (CN-5)
tumor boundary (CN-6)
artifacts (CN-0, not shown in Figure 4B)
granulocyte enriched (CN-9)
smooth muscle (CN-8)
macrophage enriched (CN-4)
immune-infiltrated stroma / vascularized smooth muscle (CN-3 / CN-7)
tumor cells
CD11c+ DCs
tumor cells / immune cells
smooth muscle
lymphatics
adipocytes
undefined
CD4+ T cells CD45RO+
CD8+ T cells
CD68+CD163+ macrophages
plasma cells
Tregs
immune cells / vasculature
stroma
CD68+ macrophages GzmB+
vasculature
nerves
CD11b+CD68+ macrophages
granulocytes
CD68+ macrophages
NK cells
CD11b+ monocytes
immune cells
CD4+ T cells GATA3+
CD163+ macrophages
CD3+ T cells
CD4+ T cells
B cells
dirt
2
1
0
−1
−2
−3
DII
cellular neighborhood most similar to:
main tumor (CN-2)
artifacts (CN-0, not shown in Figure 4B)
T cell enriched (CN-1)
immune-infiltrated stroma (CN-3)
macrophage enriched (CN-4)
not defined (CN-X)
smooth muscle (CN-8)
granulocyte enriched (CN-9)
vascularized smooth muscle (CN-7)
tumor boundary (CN-6)
tumor cells
CD11c+ DCs
tumor cells / immune cells
smooth muscle
lymphatics
adipocytes
undefined
CD4+ T cells CD45RO+
CD8+ T cells
CD68+CD163+ macrophages
plasma cells
Tregs
immune cells / vasculature
stroma
CD68+ macrophages GzmB+
vasculature
nerves
CD11b+CD68+ macrophages
granulocytes
CD68+ macrophages
NK cells
CD11b+ monocytes
immune cells
CD4+ T cells GATA3+
CD163+ macrophages
CD3+ T cells
CD4+ T cells
B cells
dirt

### Figure S19

#### CLR

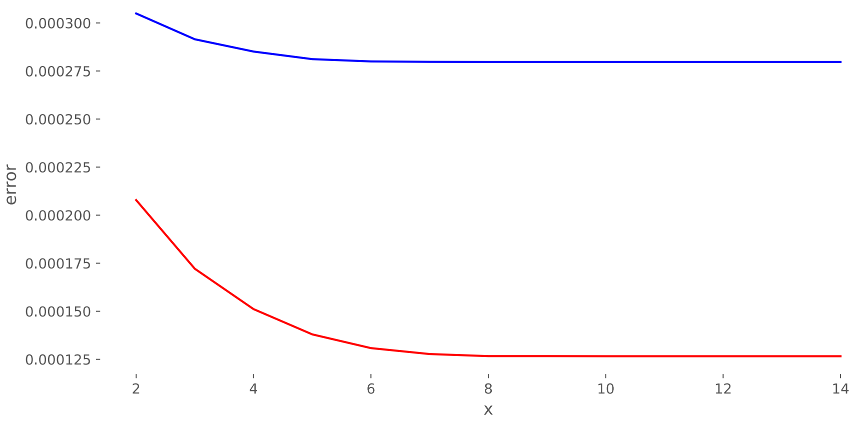

#### DII

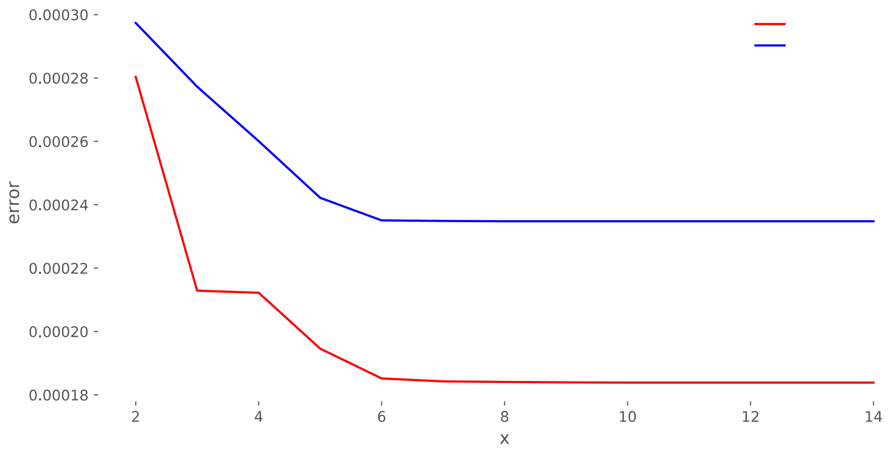

### Figure S20

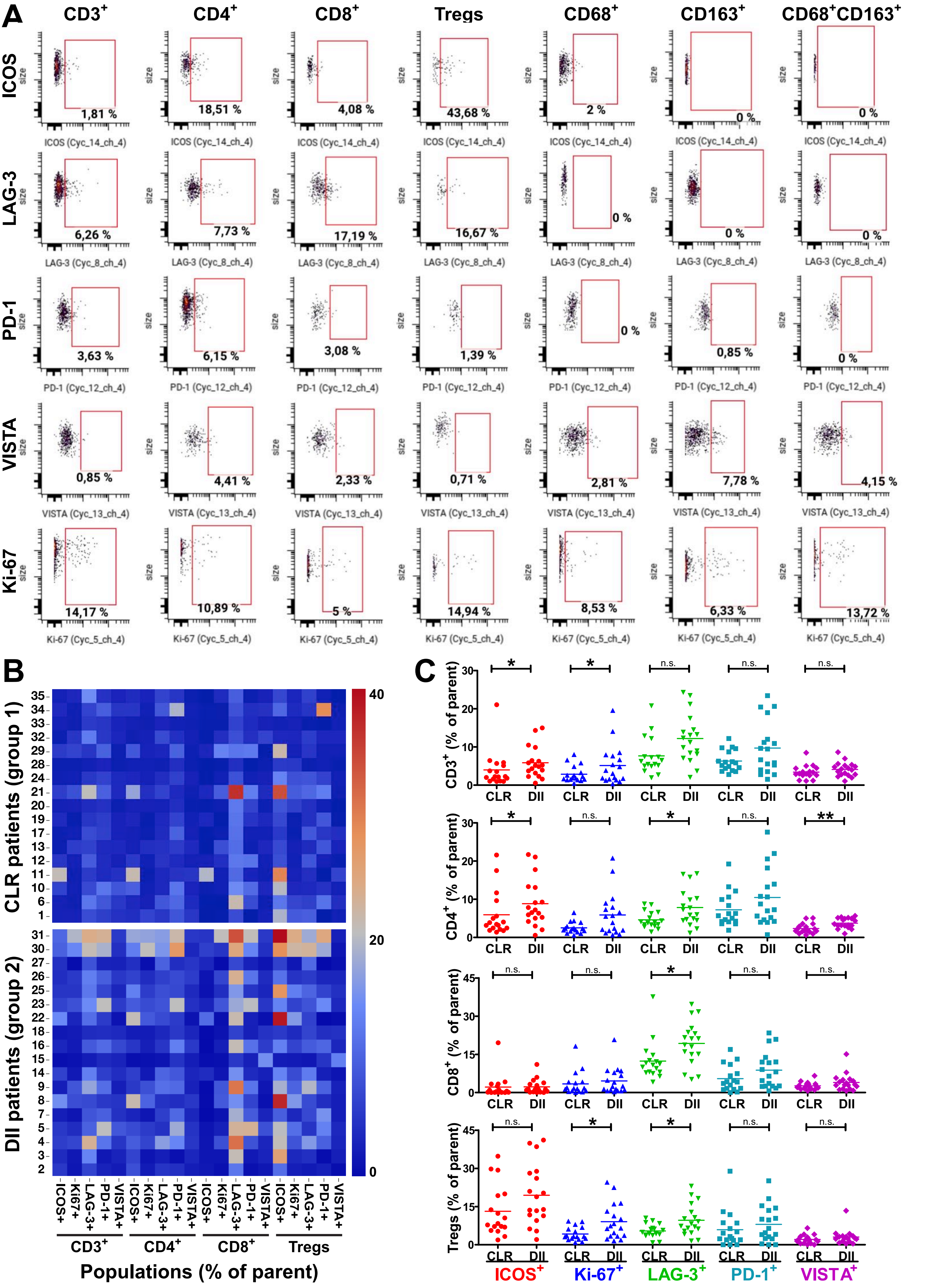
A
CD3+
CD4+
CD8+
Tregs
CD68+
CD163+
CD68+CD163+
ICOS
LAG-3
PD-1
VISTA
Ki-67
ICOS (Cyc_14_ch_4)
LAG-3 (Cyc_8_ch_4)
PD-1 (Cyc_12_ch_4)
VISTA (Cyc_13_ch_4)
Ki-67 (Cyc_5_ch_4)
size
1,81 %
18,51 %
4,08 %
43,68 %
2 %
0 %
0 %
6,26 %
7,73 %
17,19 %
16,67 %
0 %
0 %
0 %
3,63 %
6,15 %
3,08 %
1,39 %
0 %
0,85 %
0 %
0,85 %
4,41 %
2,33 %
0,71 %
2,81 %
7,78 %
4,15 %
14,17 %
10,89 %
5 %
14,94 %
8,53 %
6,33 %
13,72 %
B
CLR patients (group 1)
DII patients (group 2)
Populations (% of parent)
ICOS+
Ki67+
LAG-3+
PD-1+
VISTA+
CD3+
CD4+
CD8+
Tregs
40
20
0
C
CD3+ (% of parent)
CD4+ (% of parent)
CD8+ (% of parent)
Tregs (% of parent)
CLR
DII
n.s.
*
**
ICOS+
Ki-67+
LAG-3+
PD-1+
VISTA+

### Figure S21

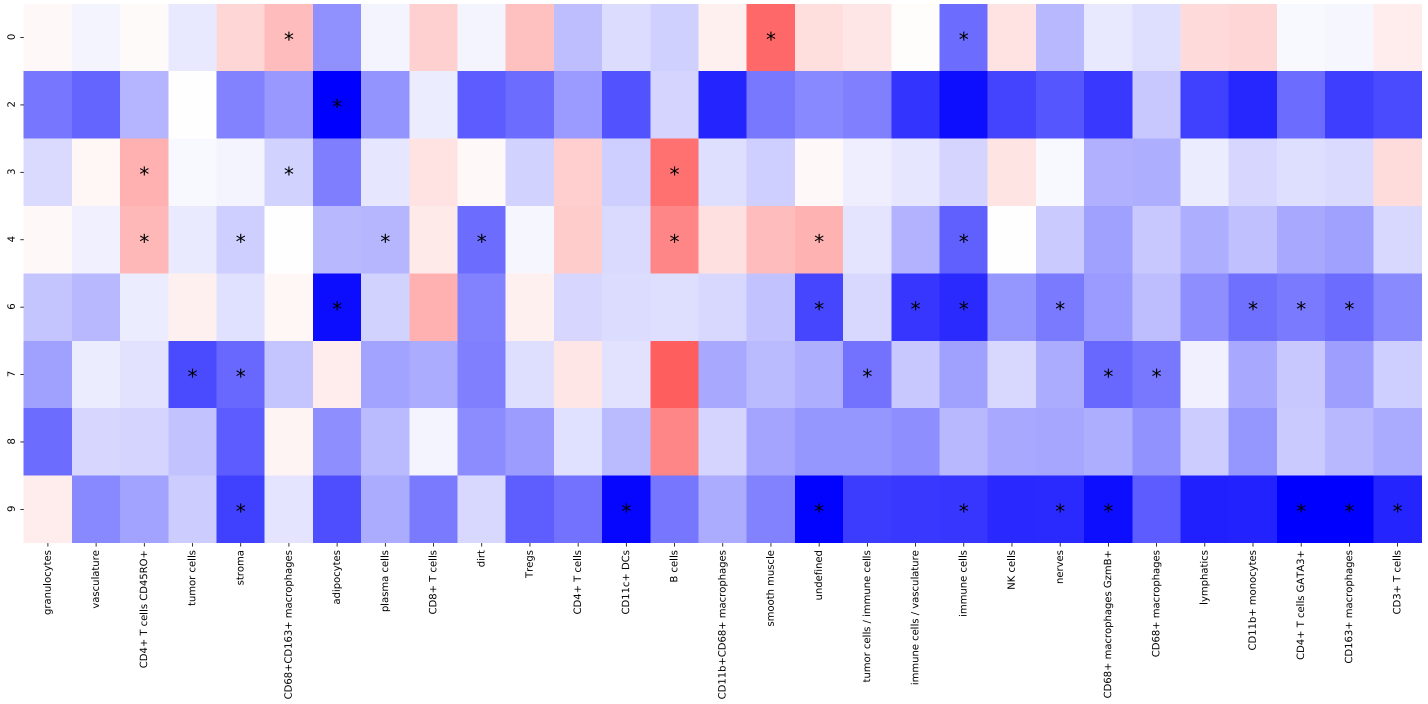
0
2
3
4
6
7
8
9
granulocytes
vasculature
CD4+ T cells CD45RO+
tumor cells
stroma
CD68+CD163+ macrophages
adipocytes
plasma cells
CD8+ T cells
dirt
Tregs
CD4+ T cells
CD11c+ DCs
B cells
CD11b+CD68+ macrophages
smooth muscle
undefined
tumor cells / immune cells
immune cells / vasculature
immune cells
NK cells
nerves
CD68+ macrophages GzmB+
CD68+ macrophages
lymphatics
CD11b+ monocytes
CD4+ T cells GATA3+
CD163+ macrophages
CD3+ T cells

#### Supplemental Figure 22

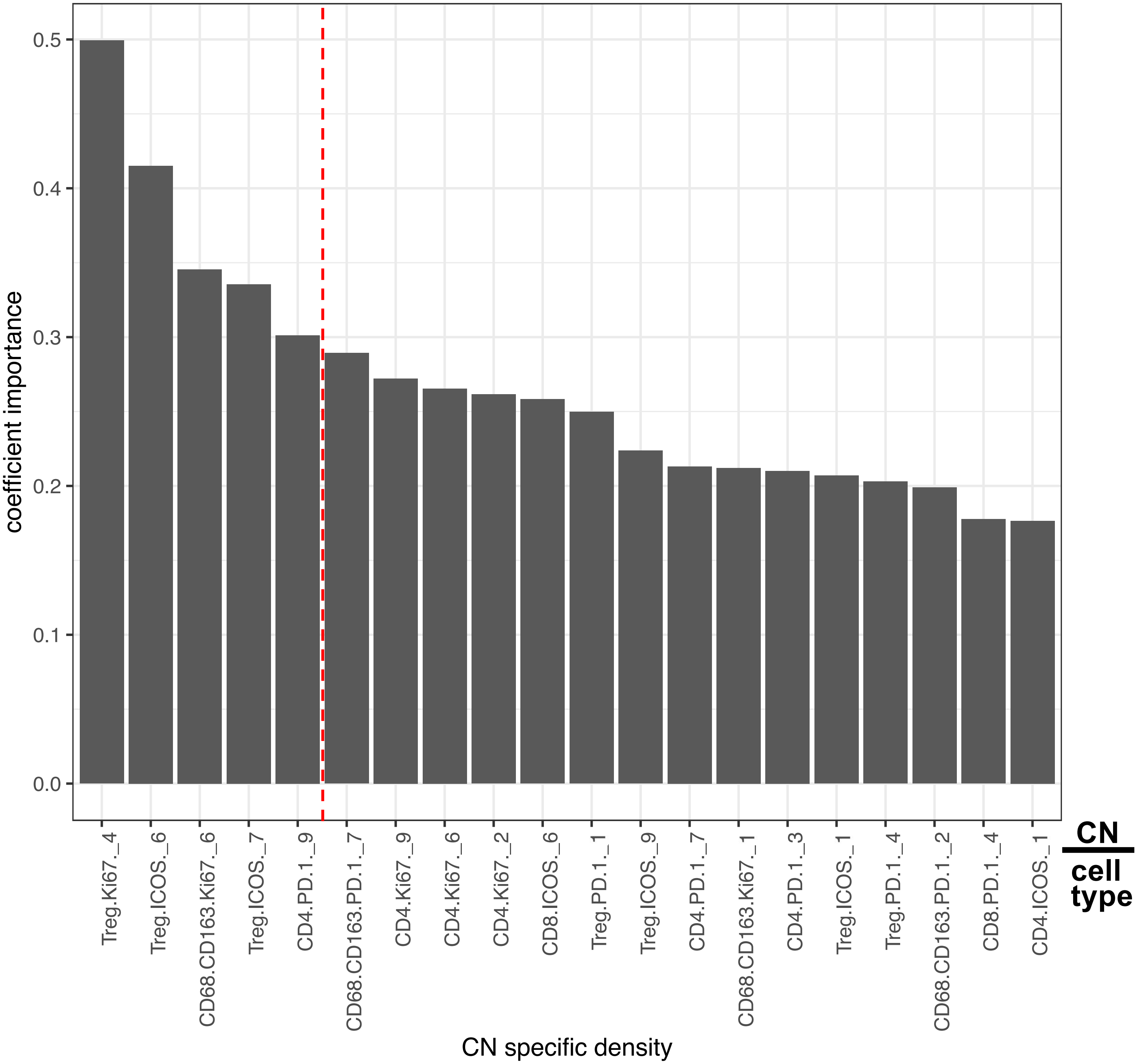

#### TABLE S2

| Antigen | Clone(s) | Manufacturer | Catalog No. | CODEX oligo | Working dilution | Exposure time |
|---|---|---|---|---|---|---|
| $\alpha$-SMA | polyclonal | Abcam | ab5694 | 69 | 1:200 | 1/4s |
| $\beta$-catenin | polyclonal | Novus Biologicals | AF1329 | 51 | 1:25 | 1/2s |
| BCL-2 | 124 | Cell Marque | custom | 41 | 1:50 | 1/2s |
| CD11b | EPR1344 | Abcam | ab216445 | 28 | 1:25 | 1/2s |
| CD11c | EP1347Y | Abcam | ab216655 | 49 | 1:50 | 1/2s |
| CD134 (OX40) | Ber-ACT35 | Biolegend | 350002 | 71 | 1:100 | 1/2s |
| CD138 | B-A38 | Thermo Fisher Scientific | MA1-10091 | 76 | 1:100 | 1/8.5s |
| CD15 | MMA | BD Biosciences | 559045 | 14 | 1:200 | 1/8.5s |
| CD163 | EDHu-1 | Novus Biologicals | NB110-40686 | 45 | 1:50 | 1/2s |
| CD194 (CCR4) | L291H4 | Biolegend | 359402 | 55 | 1:25 | 1/2s |
| CD2 | RPA-2.10 | Biolegend | 300202 | 25 | 1:25 | 1/2s |
| CD20 | rIGEL/773 | Novus Biologicals | NBP2-54591 | 48 | 1:200 | 1/4s |
| CD21 | Bu32 | Biolegend | 354902 | 21 | 1:50 | 1/2s |
| CD223 (LAG3) | D2G4O | Cell Signaling Technology | custom | 42 | 1:25 | 1/2s |
| CD25 | 4C9 | Cell Marque | custom | 24 | 1:100 | 1/2s |
| CD274 (PD-L1) | E1L3N | Cell Signaling Technology | custom | 11 | 1:50 | 1/2s |
| CD278 (ICOS) | D1K2T | Cell Signaling Technology | custom | 74 | 1:100 | 1/4s |
| CD279 (PD-1) | D4W2J | Cell Signaling Technology | custom | 23 | 1:50 | 1/2s |
| CD3 | MRQ-39 | Cell Marque | custom | 77 | 1:100 | 1/2s |
| CD30 | BerH2 | Cell Marque | custom | 57 | 1:25 | 1/2s |
| CD31 | C31.3 + C31.7 + C31.10 | Novus Biologicals | NBP2-47785 | 68 | 1:200 | 1/5s |
| CD34 | QBEnd/10 | Novus Biologicals | NBP2-34713 | 38 | 1:100 | 1/2s |
| CD38 | EPR4106 | Abcam | ab176886 | 66 | 1:100 | 1/2s |
| CD4 | EPR6855 | Abcam | ab181724 | 20 | 1:100 | 1/2s |
| CD44 | IM-7 | Biolegend | 103002 | 44 | 1:25 | 1/2s |
| CD45 | 2B11 + PD7/26 | Novus Biologicals | NBP2-34528 | 56 | 1:100 | 1/4s |
| CD45RA | HI100 | BD Biosciences | 555486 | 72 | 1:50 | 1/2s |
| CD45RO | UCH-L1 | Santa Cruz Biotechnology | sc-1183 | 2 | 1:25 | 1/2s |
| CD5 | UCHT2 | BD Biosciences | 555350 | 75 | 1:25 | 1/2s |
| CD56 | MRQ-42 | Cell Marque | custom | 29 | 1:50 | 1/2s |
| CD57 | HCD57 | Biolegend | 322325 | 30 | 1:200 | 1/2s |
| CD68 | KP-1 | Biolegend | 916104 | 70 | 1:200 | 1/3s |

| | | | | | | |
|---|---|---|---|---|---|---|
| CD68 | D4B9C | Cell Signaling Technology | custom | 5 | 1:100 | 1/2s |
| CD7 | MRQ-56 | Cell Marque | custom | 63 | 1:100 | 1/4s |
| CD71 | MRQ-48 | Cell Marque | custom | 3 | 1:100 | 1/2s |
| CD79a | JBC117 | Cell Marque | custom | 46 | 1:25 | 1/2s |
| CD8 | C8/144B | Cell Marque | custom | 8 | 1:50 | 1/2s |
| CDX2 | CDX2/1690 | Novus Biologicals | NBP2-54472 | 53 | 1:25 | 1/2s |
| Chromogranin A | LK2H10 + PHE5 + CGA/414 | Novus | NBP2-34674 | 43 | 1:400 | 1/4s |
| Collagen IV | polyclonal | Abcam | ab6586 | 33 | 1:100 | 1/4s |
| Cytokeratin 7 | OV-TL12/30 | Novus | NBP2-47940 | 3 | 1:200 | 1/4s |
| EGFR | D38B1 | Cell Signaling Technology | custom | 58 | 1:25 | 1/2s |
| EpCAM | BerEp4 | Cell Marque | custom | 70 | 1:25 | 1/2s |
| FoxP3 | 236A/E7 | Invitrogen | 14-4777-80 | 61 | 1:100 | 1/2s |
| GATA3 | L50-823 | Cell Marque | custom | 60 | 1:100 | 1/2s |
| GFAP | 2.2B10 | Thermo Fisher Scientific | 13-0300 | 46 | 1:25 | 1/2s |
| Granzyme B | EPR20129-217 | Abcam | ab219803 | 81 | 1:200 | 1/4s |
| Hep-Par-1 | OCH1E5 | Santa Cruz Biotechnology | sc-58693 | 28 | 1:100 | 1/2s |
| HLA-DR | EPR3692 | Abcam | ab215985 | 65 | 1:100 | 1/4s |
| IDO-1 | D5J4E | Cell Signaling Technology | custom | 59 | 1:25 | 1/2s |
| IRF4 | IRF4.3E4 | Biolegend | 646402 | 51 | 1:25 | 1/2s |
| Ki-67 | B56 | BD Biosciences | 556003 | 6 | 1:100 | 1/8.5s |
| Melan-A | A103 + M2-7C10 + M2-9E3 | Novus Biologicals | NBP2-34546 | 44 | 1:50 | 1/2s |
| MMP12 | polyclonal | Abcam | ab137444 | 80 | 1:200 | 1/4s |
| MMP9 | L51/82 | Biolegend | 819701 | 62 | 1:400 | 1/4s |
| MUC-1 (EMA) | 955 | NSJ Bioreagents | V2372SAF | 15 | 1:100 | 1/4s |
| Na-K-ATPase | EP1845Y | Abcam | ab167390 | 36 | 1:100 | 1/4s |
| p53 | D07 | Cell Marque | custom | 52 | 1:25 | 1/2s |
| Pan-Cytokeratin | C11 | Biolegend | 628602 | 67 | 1:200 | 1/8.5s |
| PAX5 | D7H5X | Cell Signaling Technology | custom | 42 | 1:25 | 1/2s |
| Podoplanin | D2-40 | Biolegend | 916606 | 32 | 1:200 | 1/4s |
| Synaptophysin | 7H12 | Novus | NBP1-47483 | 26 | 1:100 | 1/4s |
| T-bet | D6N8B | Cell Signaling Technology | custom | 5 | 1:100 | 1/2s |
| Vimentin | RV202 | BD Biosciences | 550513 | 7 | 1:200 | 1/2s |
| VISTA | D1L2G | Cell Signaling Technology | custom | 79 | 1:50 / 1:100 | 1/2s |

### TABLE S3

| Oligo No. | Antibody oligo sequence (5'-3') | Fluorescent oligonucleotide sequences (5'-3') | | | Blocking component 4 (BC4) oligos (5'-3') |
|---|---|---|---|---|---|
| 2 | /mal/ATGGTTTAGGACTAC | /5Alex488N/GTAGTCCTAAACCAT | /5ATTO550N/GTAGTCCTAAACCAT | | GTAGTCCTAAACCAT |
| 3 | /mal/TACTCCTCGCCG | /5Alex488N/CGGCGAGGAGTA | /5ATTO550N/CGGCGAGGAGTA | /5Alex647N/CGGCGAGGAGTA | CGGCGAGGAGTA |
| 5 | /mal/TCTCCCATTAGTCGG | /5Alex488N/CCGACTAATGGGAGA | /5ATTO550N/CCGACTAATGGGAGA | | CCGACTAATGGGAGA |
| 6 | /mal/TGGATGTGTTACGAT | /5Alex488N/ATCGTAACACATCCA | /5ATTO550N/ATCGTAACACATCCA | /5Alex647N/ATCGTAACACATCCA | ATCGTAACACATCCA |
| 7 | /mal/CGCTAAGATATTCTAAG | /5Alex488N/CTTAGAATATCTTAGCG | /5ATTO550N/CTTAGAATATCTTAGCG | | CTTAGAATATCTTAGCG |
| 8 | /mal/CGCAGATGAATATTC | /5Alex488N/GAATATTCATCTGCG | /5ATTO550N/GAATATTCATCTGCG | /5Alex647N/GAATATTCATCTGCG | GAATATTCATCTGCG |
| 11 | /mal/GGGTTATACACTCGT | /5Alex488N/ACGAGTGTATAACCC | /5ATTO550N/ACGAGTGTATAACCC | | ACGAGTGTATAACCC |
| 14 | /mal/AGATTCAGTCTCTCG | /5Alex488N/CGAGAGACTGAATCT | /5ATTO550N/CGAGAGACTGAATCT | | CGAGAGACTGAATCT |
| 15 | /mal/CTGTAATAGGCACTA | /5Alex488N/TAGTGCCTATTACAG | /5ATTO550N/TAGTGCCTATTACAG | /5Alex647N/TAGTGCCTATTACAG | TAGTGCCTATTACAG |
| 20 | /mal/ATGTAGGATGGTCTC | | /5ATTO550N/GAGACCATCCTACAT | /5Alex647N/GAGACCATCCTACAT | GAGACCATCCTACAT |
| 21 | /mal/CGTGCCGTTTCAC | /5Alex488N/GTGAAACGGCACG | /5ATTO550N/GTGAAACGGCACG | /5Alex647N/GTGAAACGGCACG | GTGAAACGGCACG |
| 23 | /mal/GGTTTCCTCAGACAC | /5Alex488N/GTGTCTGAGGAAACC | /5ATTO550N/GTGTCTGAGGAAACC | /5Alex647N/GTGTCTGAGGAAACC | GTGTCTGAGGAAACC |
| 24 | /mal/ATAAGGGCTCATTGT | /5Alex488N/ACAATGAGCCCTTAT | /5ATTO550N/ACAATGAGCCCTTAT | /5Alex647N/ACAATGAGCCCTTAT | ACAATGAGCCCTTAT |
| 25 | /mal/GCACGCCCTTTTA | | /5ATTO550N/TAAAAGGGCGTGC | /5Alex647N/TAAAAGGGCGTGC | TAAAAGGGCGTGC |
| 26 | /mal/CACTTGTCTAACCAA | /5Alex488N/TTGGTTAGACAAGTG | /5ATTO550N/TTGGTTAGACAAGTG | /5Alex647N/TTGGTTAGACAAGTG | TTGGTTAGACAAGTG |
| 28 | /mal/TGGTCCACTAACGTA | /5Alex488N/TACGTTAGTGGACCA | /5ATTO550N/TACGTTAGTGGACCA | /5Alex647N/TACGTTAGTGGACCA | TACGTTAGTGGACCA |
| 29 | /mal/ATAGGGCATTTGAAG | /5Alex488N/CTTCAAATGCCCTAT | /5ATTO550N/CTTCAAATGCCCTAT | /5Alex647N/CTTCAAATGCCCTAT | CTTCAAATGCCCTAT |
| 30 | /mal/CACATGAGCGAATCA | | /5ATTO550N/TGATTCGCTCATGTG | /5Alex647N/TGATTCGCTCATGTG | TGATTCGCTCATGTG |
| 32 | /mal/TACCAAATCCTGATG | /5Alex488N/CATCAGGATTTGGTA | /5ATTO550N/CATCAGGATTTGGTA | /5Alex647N/CATCAGGATTTGGTA | CATCAGGATTTGGTA |
| 33 | /mal/TTATCATGAGGAGCG | /5Alex488N/CGCTCCTCATGATAA | | /5Alex647N/CGCTCCTCATGATAA | CGCTCCTCATGATAA |
| 36 | /mal/ACCTACACAATGCTA | /5Alex488N/TAGCATTGTGTAGGT | /5ATTO550N/TAGCATTGTGTAGGT | | TAGCATTGTGTAGGT |
| 38 | /mal/GCGTCTACTTATAAG | /5Alex488N/CTTATAAGTAGACGC | /5ATTO550N/CTTATAAGTAGACGC | | CTTATAAGTAGACGC |
| 41 | /mal/TGTATGAGTAGTAATCT | /5Alex488N/AGATTACTACTCATACA | /5ATTO550N/AGATTACTACTCATACA | /5Alex647N/AGATTACTACTCATACA | AGATTACTACTCATACA |
| 42 | /mal/TCTAAGTCAGAGAGC | | /5ATTO550N/GCTCTCTGACTTAGA | /5Alex647N/GCTCTCTGACTTAGA | GCTCTCTGACTTAGA |
| 43 | /mal/GACATTATCCGTGAT | /5Alex488N/ATCACGGATAATGTC | /5ATTO550N/ATCACGGATAATGTC | /5Alex647N/ATCACGGATAATGTC | ATCACGGATAATGTC |
| 44 | /mal/TCACTACTATTAGTACT | /5Alex488N/AGTACTAATAGTAGTGA | /5ATTO550N/AGTACTAATAGTAGTGA | | AGTACTAATAGTAGTGA |
| 45 | /mal/GCCAGAATGCCA | | /5ATTO550N/TGGCATTCTGGC | /5Alex647N/TGGCATTCTGGC | TGGCATTCTGGC |
| 46 | /mal/GACTCGAACCTGAG | /5Alex488N/CTCAGGTTCGAGTC | | /5Alex647N/CTCAGGTTCGAGTC | CTCAGGTTCGAGTC |
| 48 | /mal/GCACGGCAAAGTG | /5Alex488N/CACTTTGCCGTGC | /5ATTO550N/CACTTTGCCGTGC | /5Alex647N/CACTTTGCCGTGC | CACTTTGCCGTGC |
| 49 | /mal/ATAACGCCTCGTATC | /5Alex488N/GATACGAGGCGTTAT | /5ATTO550N/GATACGAGGCGTTAT | | GATACGAGGCGTTAT |
| 51 | /mal/CGTGCGGGAAAAT | /5Alex488N/ATTTTCCCGCACG | | /5Alex647N/ATTTTCCCGCACG | ATTTTCCCGCACG |
| 52 | /mal/ACCAGGTGATTGCAT | | /5ATTO550N/ATGCAATCACCTGGT | /5Alex647N/ATGCAATCACCTGGT | ATGCAATCACCTGGT |
| 53 | /mal/TATCCCGTAAACTTC | | /5ATTO550N/GAAGTTTACGGGATA | /5Alex647N/GAAGTTTACGGGATA | GAAGTTTACGGGATA |
| 55 | /mal/AGGTCAACTCGCAC | /5Alex488N/GTGCGAGTTGACCT | /5ATTO550N/GTGCGAGTTGACCT | | GTGCGAGTTGACCT |
| 56 | /mal/GGTCACATGGTCGTT | /5Alex488N/AACGACCATGTGACC | /5ATTO550N/AACGACCATGTGACC | /5Alex647N/AACGACCATGTGACC | AACGACCATGTGACC |
| 57 | /mal/GCGGATTTCGTATTT | /5Alex488N/AAATACGAAATCCGC | /5ATTO550N/AAATACGAAATCCGC | /5Alex647N/AAATACGAAATCCGC | AAATACGAAATCCGC |
| 58 | /mal/CGTCAGTACTTTCAG | /5Alex488N/CTGAAAGTACTGACG | /5ATTO550N/CTGAAAGTACTGACG | | CTGAAAGTACTGACG |
| 59 | /mal/GCTTATTATGGACTTC | | /5ATTO550N/GAAGTCCATAATAAGC | /5Alex647N/GAAGTCCATAATAAGC | GAAGTCCATAATAAGC |
| 60 | /mal/ACAAAACTGCTGTCG | | /5ATTO550N/CGACAGCAGTTTTGT | /5Alex647N/CGACAGCAGTTTTGT | CGACAGCAGTTTTGT |
| 61 | /mal/TCTTATTCCCGAATA | | /5ATTO550N/TATTCGGGAATAAGA | /5Alex647N/TATTCGGGAATAAGA | TATTCGGGAATAAGA |
| 62 | /mal/TAGGGGAACAGGTTG | /5Alex488N/CAACCTGTTCCCCTA | /5ATTO550N/CAACCTGTTCCCCTA | | CAACCTGTTCCCCTA |
| 63 | /mal/AATTAGCTTAAGAGAGT | /5Alex488N/ACTCTCTTAAGCTAATT | /5ATTO550N/ACTCTCTTAAGCTAATT | | ACTCTCTTAAGCTAATT |
| 65 | /mal/GATAAATATTTTACAGAGT | /5Alex488N/ACTCTGTAAAATATTTAT( | /5ATTO550N/ACTCTGTAAAATATTTATC | | ACTCTGTAAAATATTTATC |
| 66 | /mal/TACGTGCTTTGGTT | /5Alex488N/AACCAAAGCACGTA | /5ATTO550N/AACCAAAGCACGTA | /5Alex647N/AACCAAAGCACGTA | AACCAAAGCACGTA |
| 67 | /mal/GACGACGAAGGC | /5Alex488N/GCCTTCGTCGTC | /5ATTO550N/GCCTTCGTCGTC | /5Alex647N/GCCTTCGTCGTC | GCCTTCGTCGTC |
| 68 | /mal/CTTCTTGTGGAACC | /5Alex488N/GGTTCCACAAGAAG | /5ATTO550N/GGTTCCACAAGAAG | | GGTTCCACAAGAAG |
| 69 | /mal/CCCGGCAGTT | /5Alex488N/AACTGCCGGG | | /5Alex647N/AACTGCCGGG | AACTGCCGGG |
| 70 | /mal/AACCAAACTGACCG | /5Alex488N/CGGTCAGTTTGGTT | | /5Alex647N/CGGTCAGTTTGGTT | CGGTCAGTTTGGTT |
| 71 | /mal/TCACCCCCAGC | | /5ATTO550N/GCTGGGGGTGA | /5Alex647N/GCTGGGGGTGA | GCTGGGGGTGA |
| 72 | /mal/AACGCGACGGAT | /5Alex488N/ATCCGTCGCGTT | /5ATTO550N/ATCCGTCGCGTT | /5Alex647N/ATCCGTCGCGTT | ATCCGTCGCGTT |
| 74 | /mal/ATTTGCTTCGACGA | | /5ATTO550N/TCGTCGAAGCAAAT | /5Alex647N/TCGTCGAAGCAAAT | TCGTCGAAGCAAAT |
| 75 | /mal/CGCTTGGGTGTTTA | /5Alex488N/TAAACACCCAAGCG | /5ATTO550N/TAAACACCCAAGCG | | TAAACACCCAAGCG |
| 76 | /mal/CTGTCGGTCGTCA | | /5ATTO550N/TGACGACCGACAG | /5Alex647N/TGACGACCGACAG | TGACGACCGACAG |
| 77 | /mal/ATTTCAACAAATATTGTT | | /5ATTO550N/AACAATATTTGTTGAAA | /5Alex647N/AACAATATTTGTTGAAAT | AACAATATTTGTTGAAAT |
| 79 | /mal/GCCGGAAGTGGT | | /5ATTO550N/ACCACTTCCGGC | /5Alex647N/ACCACTTCCGGC | ACCACTTCCGGC |
| 80 | /mal/CGGGGCACCA | | /5ATTO550N/TGGTGCCCCG | /5Alex647N/TGGTGCCCCG | TGGTGCCCCG |
| 81 | /mal/CAAGGAACTACCGA | /5Alex488N/TCGGTAGTTCCTTG | /5ATTO550N/TCGGTAGTTCCTTG | /5Alex647N/TCGGTAGTTCCTTG | TCGGTAGTTCCTTG |

### TABLE S4

#### CRC panel

| Cycle | Channel 1 | Exposure Time | Alexa488 | Oligo | Clone | Ab dilution | Exposure Time | ATTO550 | Oligo | Clone | Ab dilution | Exposure Time | Alexa647 | Oligo | Clone | Ab dilution | Exposure Time |
|---|---|---|---|---|---|---|---|---|---|---|---|---|---|---|---|---|---|
| 1 | Hoechst 33342 | 1/175s | blank | | | | 1/2s | blank | | | | 1/2s | blank | | | | 1/2s |
| 2 | Hoechst 33342 | 1/175s | CD44 | 44 | IM-7 | 1:25 | 1/2s | FoxP3 | 61 | 236A/E7 | 1:100 | 1/2s | CDX2 | 53 | CDX2/1690 | 1:25 | 1/2s |
| 3 | Hoechst 33342 | 1/175s | CD8 | 8 | C8/144B | 1:50 | 1/2s | p53 | 52 | D07 | 1:25 | 1/2s | GATA3 | 60 | L50-823 | 1:100 | 1/2s |
| 4 | Hoechst 33342 | 1/175s | CD45 | 56 | 2B11+PD7/26 | 1:100 | 1/4s | T-bet | 5 | D6N8B | 1:100 | 1/2s | β-catenin | 51 | polyclonal | 1:25 | 1/2s |
| 5 | Hoechst 33342 | 1/175s | HLA-DR | 65 | EPR3692 | 1:100 | 1/4s | CD274 (PD-L1) | 11 | E1L3N | 1:50 | 1/2s | Ki-67 | 6 | B56 | 1:100 | 1/8.5s |
| 6 | Hoechst 33342 | 1/175s | CD45RA | 72 | HI100 | 1:50 | 1/2s | CD4 | 20 | EPR6855 | 1:100 | 1/2s | CD21 | 21 | Bu32 | 1:50 | 1/2s |
| 7 | Hoechst 33342 | 1/175s | MUC-1 (EMA) | 15 | 955 | 1:100 | 1/4s | CD30 | 57 | BerH2 | 1:25 | 1/2s | CD2 | 25 | RPA-2.10 | 1:25 | 1/2s |
| 8 | Hoechst 33342 | 1/175s | Vimentin | 7 | RV202 | 1:200 | 1/2s | CD20 | 48 | rIGEL/773 | 1:200 | 1/4s | CD223 (LAG3) | 42 | D2G4O | 1:25 | 1/2s |
| 9 | Hoechst 33342 | 1/175s | Na-K-ATPase | 36 | EP1845Y | 1:100 | 1/4s | CD5 | 75 | UCHT2 | 1:25 | 1/2s | IDO-1 | 59 | D5J4E | 1:25 | 1/2s |
| 10 | Hoechst 33342 | 1/175s | Pan-Cytokeratin | 67 | C11 | 1:200 | 1/8.5s | CD11b | 28 | EPR1344 | 1:25 | 1/2s | CD56 | 29 | MRQ-42 | 1:50 | 1/2s |
| 11 | Hoechst 33342 | 1/175s | α-SMA | 69 | polyclonal | 1:200 | 1/4s | BCL-2 | 41 | 124 | 1:50 | 1/2s | CD25 | 24 | 4C9 | 1:100 | 1/2s |
| 12 | Hoechst 33342 | 1/175s | Collagen IV | 33 | polyclonal | 1:100 | 1/4s | CD11c | 49 | EP1347Y | 1:50 | 1/2s | CD279 (PD-1) | 23 | D4W2J | 1:50 | 1/2s |
| 13 | Hoechst 33342 | 1/175s | Granzyme B | 81 | EPR20129-217 | 1:200 | 1/8.5s | EGFR | 58 | D38B1 | 1:25 | 1/2s | VISTA | 79 | D1L2G | 1:50 | 1/2s |
| 14 | Hoechst 33342 | 1/175s | CD15 | 14 | MMA | 1:200 | 1/8.5s | CD194 (CCR4) | 55 | L291H4 | 1:25 | 1/2s | CD278 (ICOS) | 74 | D1K2T | 1:100 | 1/4s |
| 15 | Hoechst 33342 | 1/175s | MMP9 | 62 | L51/82 | 1:400 | 1/4s | Synaptophysin | 26 | 7H12 | 1:100 | 1/4s | CD71 | 3 | MRQ-48 | 1:100 | 1/2s |
| 16 | Hoechst 33342 | 1/175s | GFAP | 46 | 2.2B10 | 1:25 | 1/2s | CD7 | 63 | MRQ-56 | 1:100 | 1/4s | CD3 | 77 | MRQ-39 | 1:100 | 1/2s |
| 17 | Hoechst 33342 | 1/175s | Chromogranin A | 43 | LK2H10+PHE5+CGA/414 | 1:400 | 1/4s | CD163 | 45 | EDHu-1 | 1:50 | 1/4s | CD57 | 30 | HCD57 | 1:200 | 1/2s |
| 18 | Hoechst 33342 | 1/175s | blank | | | | 1/600s | CD45RO | 2 | UCH-L1 | 1:25 | 1/2s | CD68 | 70 | KP-1 | 1:200 | 1/3s |
| 19 | Hoechst 33342 | 1/175s | blank | | | | 1/600s | CD31 | 68 | C31.3+C31.7+C31.10 | 1:200 | 1/5s | Podoplanin | 32 | D2-40 | 1:200 | 1/4s |
| 20 | Hoechst 33342 | 1/175s | blank | | | | 1/600s | CD34 | 38 | QBEnd/10 | 1:100 | 1/2s | CD38 | 66 | EPR4106 | 1:100 | 1/2s |
| 21 | Hoechst 33342 | 1/175s | blank | | | | 1/600s | CD138 | 76 | B-A38 | 1:100 | 1/8.5s | MMP12 | 80 | polyclonal | 1:200 | 1/4s |
| 22 | Hoechst 33342 | 1/175s | blank | | | | 1/8.5s | blank | | | | 1/8.5s | blank | | | | 1/8.5s |
| 23 | Hoechst 33342 | 1/175s | blank | | | | 1/600s | blank | | | | 1/600s | DRAQ5 | | | 1:100 | 1/8.5s |
| 24 | H&E (brightfield) | | | | | | | | | | | | | | | | |

#### Multi-tumor TMA panel

| Cycle | Channel 1 | Exposure Time | Alexa488 | Oligo | Clone | Ab dilution | Exposure Time | ATTO550 | Oligo | Clone | Ab dilution | Exposure Time | Alexa647 | Oligo | Clone | Ab dilution | Exposure Time |
|---|---|---|---|---|---|---|---|---|---|---|---|---|---|---|---|---|---|
| 1 | Hoechst 33342 | 1/175s | blank | | | | 1/2s | blank | | | | 1/2s | blank | | | | 1/2s |
| 2 | Hoechst 33342 | 1/175s | CD79a | 46 | JBC117 | 1:25 | 1/2s | FoxP3 | 61 | 236A/E7 | 1:100 | 1/2s | CDX2 | 53 | CDX2/1690 | 1:25 | 1/2s |
| 3 | Hoechst 33342 | 1/175s | CD8 | 8 | C8/144B | 1:50 | 1/2s | p53 | 52 | D07 | 1:25 | 1/2s | GATA3 | 60 | L50-823 | 1:100 | 1/2s |
| 4 | Hoechst 33342 | 1/175s | CD21 | 21 | Bu32 | 1:50 | 1/2s | CD274 (PD-L1) | 11 | E1L3N | 1:50 | 1/2s | Ki-67 | 6 | B56 | 1:100 | 1/8.5s |
| 5 | Hoechst 33342 | 1/175s | CD45 | 56 | 2B11+PD7/26 | 1:100 | 1/4s | CD30 | 57 | BerH2 | 1:25 | 1/2s | PAX5 | 42 | D7H5X | 1:25 | 1/2s |
| 6 | Hoechst 33342 | 1/175s | HLA-DR | 65 | EPR3692 | 1:100 | 1/4s | CD5 | 75 | UCHT2 | 1:50 | 1/2s | CD2 | 25 | RPA-2.10 | 1:25 | 1/2s |
| 7 | Hoechst 33342 | 1/175s | CD45RA | 72 | HI100 | 1:50 | 1/2s | CD4 | 20 | EPR6855 | 1:100 | 1/2s | CD279 (PD-1) | 23 | D4W2J | 1:50 | 1/2s |
| 8 | Hoechst 33342 | 1/175s | MUC-1 (EMA) | 15 | 955 | 1:100 | 1/4s | BCL-2 | 41 | 124 | 1:50 | 1/2s | CD56 | 29 | MRQ-42 | 1:100 | 1/2s |
| 9 | Hoechst 33342 | 1/175s | Cytokeratin 7 | 3 | OV-TL12/30 | 1:200 | 1/4s | CD25 | 24 | 4C9 | 1:100 | 1/2s | VISTA | 79 | D1L2G | 1:100 | 1/2s |
| 10 | Hoechst 33342 | 1/175s | Hep-Par-1 | 28 | OCH1E5 | 1:100 | 1/2s | CD11c | 49 | EP1347Y | 1:50 | 1/2s | IRF4 | 51 | IRF4.3E4 | 1:25 | 1/2s |
| 11 | Hoechst 33342 | 1/175s | CD20 | 48 | rIGEL/773 | 1:400 | 1/8.5s | EGFR | 58 | D38B1 | 1:25 | 1/2s | IDO-1 | 59 | D5J4E | 1:25 | 1/2s |
| 12 | Hoechst 33342 | 1/175s | Granzyme B | 81 | EPR20129-217 | 1:200 | 1/4s | Melan-A | 44 | A103 + M2-7C10 + M2 | 1:50 | 1/2s | - | | | | |
| 13 | Hoechst 33342 | 1/175s | Vimentin | 7 | RV202 | 1:200 | 1/2s | Synaptophysin | 26 | 7H12 | 1:100 | 1/4s | - | | | | |
| 14 | Hoechst 33342 | 1/175s | Na-K-ATPase | 36 | EP1845Y | 1:100 | 1/4s | CD194 (CCR4) | 55 | L291H4 | 1:25 | 1/2s | CD57 | 30 | HCD57 | 1:200 | 1/2s |
| 15 | Hoechst 33342 | 1/175s | Pan-Cytokeratin | 67 | C11 | 1:200 | 1/8.5s | CD68 | 5 | D4B9C | 1:100 | 1/2s | CD3 | 77 | MRQ-39 | 1:100 | 1/2s |
| 16 | Hoechst 33342 | 1/175s | α-SMA | 69 | polyclonal | 1:200 | 1/4s | CD34 | 38 | QBEnd/10 | 1:50 | 1/2s | EpCAM | 70 | BerEp4 | 1:25 | 1/2s |
| 17 | Hoechst 33342 | 1/175s | Collagen IV | 33 | polyclonal | 1:200 | 1/4s | CD45RO | 2 | UCH-L1 | 1:25 | 1/2s | Podoplanin | 32 | D2-40 | 1:200 | 1/4s |
| 18 | Hoechst 33342 | 1/175s | CD15 | 14 | MMA | 1:200 | 1/8.5s | CD7 | 63 | MRQ-56 | 1:25 | 1/2s | CD163 | 45 | EDHu-1 | 1:50 | 1/2s |
| 19 | Hoechst 33342 | 1/175s | Chromogranin A | 43 | LK2H10+PHE5+CGA/414 | 1:400 | 1/4s | CD31 | 68 | C31.3+C31.7+C31.10 | 1:200 | 1/5s | - | | | | |
| 20 | Hoechst 33342 | 1/175s | MMP9 | 62 | L51/82 | 1:400 | 1/4s | CD138 | 76 | B-A38 | 1:100 | 1/8.5s | CD38 | 66 | EPR4106 | 1:25 | 1/2s |
| 21 | Hoechst 33342 | 1/175s | blank | | | | 1/2s | blank | | | | 1/2s | blank | | | | 1/2s |
| 22 | Hoechst 33342 | 1/175s | blank | | | | 1/600s | Hyaluronan-BP-bi | Streptavidin-PE | | 1:500 each | 1/4s | DRAQ5 | | | 1:100 | 1/8.5s |
| 23 | H&E (brightfield) | | | | | | | | | | | | | | | | |

#### Tonsil panel

| Cycle | Channel 1 | Exposure Time | Alexa488 | Oligo | Clone | Ab dilution | Exposure Time | ATTO550 | Oligo | Clone | Ab dilution | Exposure Time | Alexa647 | Oligo | Clone | Ab dilution | Exposure Time |
|---|---|---|---|---|---|---|---|---|---|---|---|---|---|---|---|---|---|
| 1 | Hoechst 33342 | 1/175s | blank | | | | 1/2s | blank | | | | 1/2s | blank | | | | 1/2s |
| 2 | Hoechst 33342 | 1/175s | CD79a | 46 | JBC117 | 1:25 | 1/2s | FoxP3 | 61 | 236A/E7 | 1:100 | 1/4 | GATA3 | 60 | L50-823 | 1:100 | 1/2s |
| 3 | Hoechst 33342 | 1/175s | CD8 | 8 | C8/144B | 1:50 | 1/2s | p53 | 52 | D07 | 1:50 | 1/2s | PAX5 | 42 | D7H5X | 1:25 | 1/2s |
| 4 | Hoechst 33342 | 1/175s | CD21 | 21 | Bu32 | 1:50 | 1/2s | CD274 (PD-L1) | 11 | E1L3N | 1:50 | 1/2s | Ki-67 | 6 | B56 | 1:100 | 1/8.5s |
| 5 | Hoechst 33342 | 1/175s | CD45 | 56 | 2B11+PD7/263 | 1:100 | 1/4s | CD30 | 57 | BerH2 | 1:25 | 1/2s | CD2 | 25 | RPA-2.10 | 1:25 | 1/2s |
| 6 | Hoechst 33342 | 1/175s | HLA-DR | 65 | EPR3692 | 1:100 | 1/4s | CD5 | 75 | UCHT2 | 1:50 | 1/2s | CD279 (PD-1) | 23 | D4W2J | 1:50 | 1/2s |
| 7 | Hoechst 33342 | 1/175s | CD45RA | 72 | HI100 | 1:50 | 1/2s | CD4 | 20 | EPR6855 | 1:50 | 1/2s | CD56 | 29 | MRQ-42 | 1:50 | 1/2s |
| 8 | Hoechst 33342 | 1/175s | MUC-1 (EMA) | 15 | 955 | 1:100 | 1/4s | BCL-2 | 41 | 124 | 1:50 | 1/2s | CD223 (LAG3) | 53 | 17B4 | 1:25 | 1/2s |
| 9 | Hoechst 33342 | 1/175s | CD71 | 3 | MRQ-48 | 1:100 | 1/8.5s | CD25 | 24 | 4C9 | 1:100 | 1/2s | VISTA | 79 | D1L2G | 1:50 | 1/2s |
| 10 | Hoechst 33342 | 1/175s | CD11c | 49 | EP1347Y | 1:100 | 1/4s | EGFR | 58 | D38B1 | 1:25 | 1/2s | CD1a | 43 | O10 + CA1/7 | 1:25 | 1/2s |
| 11 | Hoechst 33342 | 1/175s | Na-K-ATPase | 36 | EP1845Y | 1:100 | 1/4s | CD44 | 44 | IM-7 | 1:25 | 1/2s | IDO-1 | 59 | D5J4E | 1:25 | 1/2s |
| 12 | Hoechst 33342 | 1/175s | CD38 | 66 | EPR4106 | 1:100 | 1/2s | CD16 | 26 | D1N9L | 1:50 | 1/2s | IRF4 | 51 | IRF4.3E4 | 1:25 | 1/2s |
| 13 | Hoechst 33342 | 1/175s | CD20 | 48 | rIGEL/773 | 1:200 | 1/8.5s | CD7 | 63 | MRQ-56 | 1:50 | 1/2s | CD3 | 77 | MRQ-39 | 1:100 | 1/2s |
| 14 | Hoechst 33342 | 1/175s | Granzyme B | 81 | EPR20129-217 | 1:200 | 1/4s | CD194 (CCR4) | 55 | L291H4 | 1:50 | 1/2s | CD57 | 30 | HCD57 | 1:100 | 1/2s |
| 15 | Hoechst 33342 | 1/175s | Vimentin | 7 | RV202 | 1:200 | 1/2s | CD68 | 5 | D4B9C | 1:100 | 1/8.5s | CD268 (ICOS) | 74 | D1K2T | 1:100 | 1/2s |
| 16 | Hoechst 33342 | 1/175s | Pan-Cytokeratin | 67 | C11 | 1:200 | 1/8.5s | CD34 | 38 | QBEnd/10 | 1:100 | 1/2s | CD163 | 45 | EDHu-1 | 1:50 | 1/2s |
| 17 | Hoechst 33342 | 1/175s | SMA | 69 | polyclonal | 1:200 | 1/8.5s | CD45RO | 2 | UCH-L1 | 1:25 | 1/2s | CD11b | 28 | EPR1344 | 1:25 | 1/2s |
| 18 | Hoechst 33342 | 1/175s | Collagen IV | 33 | polyclonal | 1:200 | 1/4s | CD31 | 68 | C31.3+C31.7+C31.10 | 1:200 | 1/5s | - | | | | |
| 19 | Hoechst 33342 | 1/175s | CD15 | 14 | MMA | 1:200 | 1/4s | CD138 | 76 | B-A38 | 1:100 | 1/8.5s | - | | | | |
| 20 | Hoechst 33342 | 1/175s | MMP9 | 62 | L51/82 | 1:400 | 1/4s | MMP12 | 80 | polyclonal | 1:200 | 1/4s | Podoplanin | 32 | D2-40 | 1:100 | 1/2s |
| 21 | Hoechst 33342 | 1/175s | blank | | | | 1/2s | blank | | | | 1/2s | blank | | | | 1/2s |
| 22 | Hoechst 33342 | 1/175s | blank | | | | 1/600s | blank | | | | 1/600s | DRAQ5 | | | 1:100 | 1/8.5s |
| 23 | H&E (brightfield) | | | | | | | | | | | | | | | | |

#### Table S5: Tissue composition of the multi-tumor TMA

| # | tissue | normal / neoplasia | diagnosis |
|---|---|---|---|
| 1 | bone marrow | normal | Normal bone marrow (patient with diffuse large B cell lymphoma) |
| 2 | bone marrow | normal | Normal bone marrow (pediatric patient post chemotherapy for neuroblastoma) |
| 3 | bone marrow | cancer | Acute myeloid leukemia (80% blasts) |
| 4 | bone marrow | cancer | B lymphoblastic leukemia (B-ALL) |
| 5 | tonsil | normal | Normal tonsil - follicular hyperplasia |
| 6 | spleen | normal | Normal spleen (patient with mesenterial infarction) |
| 7 | lymph node | normal | Normal lymph node (iliacal) |
| 8 | lymph node | cancer | Classic Hodgkin's lymphoma - nodular sclerosis |
| 9 | lymph node | cancer | Chronic lymphocytic leukemia / small lymphocytic lymphoma (CLL / SLL) |
| 10 | lymph node | cancer | Diffuse large B cell lymphoma (DLBCL), NOS |
| 11 | lymph node | cancer | Follicular lymphoma (FL), low-grade (G 1-2) |
| 12 | lymph node | cancer | Plasmacytoma / Plasma cell myeloma |
| 13 | nasopharyngeal | cancer | Extranodal NK/T cell lymphoma, nasal type |
| 14 | lymph node | cancer | T lymphoblastic lymphoma (T-LBL) |
| 15 | thymus | cancer | Thymoma type AB with predominant type B component |
| 16 | liver | normal | Normal liver |
| 17 | appendix | normal | Normal appendix |
| 18 | liver | cancer | Hepatocellular carcinoma, G3 |
| 19 | biliary system | cancer | Cholangiocellular carcinoma, G3 |
| 20 | stomach | normal | Normal stomach (sleeve gastrectomy for obesity) |
| 21 | stomach | cancer | Gastric Adenocarcinoma, intestinal type Laurén, G3 |
| 22 | colon | cancer | Colonic Adenocarcinoma, G2 |
| 23 | pancreas | cancer | Neuroendocrine tumor, G1, insulinoma |
| 24 | salivary gland | normal | Normal seromucous salivary gland (submandibular) |
| 25 | pancreas | cancer | Anaplastic pancreatic carcinoma with osteoclast-like giant cells |
| 26 | breast | cancer | Breast invasive carcinoma, no special type (NST) |
| 27 | breast | cancer | Breast invasive carcinoma, lobular |
| 28 | ovary | cancer | Serous high-grade ovarian carcinoma |
| 29 | uterus | cancer | Endometrioid endometrial carcinoma |
| 30 | cervix | cancer | Cervical squamous cell carcinoma |
| 31 | uterus | cancer | Leiomyosarcoma |
| 32 | placenta | normal | Normal placenta |
| 33 | muscle | normal | Normal muscle |
| 34 | muscle | cancer | Alveolar rhabdomyosarcoma |
| 35 | muscle | cancer | Embryonal rhabdomyosarcoma |
| 36 | kidney | cancer | Ewing sarcoma / PNET |
| 37 | nerve | non-malignant tumor | Neurofibroma |
| 38 | soft tissue | cancer | Monophasic synovial sarcoma |
| 39 | tendon | non-malignant tumor | Giant cell tumor of tendon sheath, localized |
| 40 | bone | cancer | Telangiectatic osteosarcoma, G3 |
| 41 | nerve | cancer | Malignant peripheral nerve sheath tumor (MPNST) |
| 42 | skin | cancer | Angiosarcoma |
| 43 | prostate | cancer | Prostate acinar adenocarcinoma |
| 44 | kidney | cancer | Clear cell renal cell carcinoma, G4 |
| 45 | kidney | cancer | Papillary urothelial carcinoma, minimally invasive, of the renal pelvis |
| 46 | adrenal gland | cancer | Pheochromocytoma |
| 47 | adrenal gland | cancer | Adrenocortical carcinoma |
| 48 | testis | cancer | Seminoma |
| 49 | testis | cancer | Embryonal carcinoma |
| 50 | adrenal gland | cancer | Neuroblastoma |
| 51 | kidney | cancer | Wilms tumor / Nephroblastoma |
| 52 | adrenal gland | non-malignant tumor | Myelolipoma |
| 53 | skin | normal | "Normal skin" (chronic inflammation) |
| 54 | skin | cancer | Malignant melanoma, nodular |
| 55 | skin | cancer | Basal cell carcinoma |
| 56 | skin | cancer | Kaposi's sarcoma |
| 57 | skin | cancer | Dermatofibrosarcoma protuberans (DFSP) |
| 58 | skin | non-malignant tumor | Glomangioma |
| 59 | brain | normal | Normal brain |
| 60 | meninges | non-malignant tumor | Meningioma |
| 61 | brain | cancer | Glioblastoma multiforme, IDH-mutant |
| 62 | pituitary | non-malignant tumor | Pituitary adenoma |
| 63 | lung | cancer | Lung Adenocarcinoma |
| 64 | lung | cancer | Lung small cell carcinoma |
| 65 | lung | cancer | Lung carcinoid, typical |
| 66 | parathyroid | non-malignant tumor | Parathyroid adenoma |
| 67 | thyroid | cancer | Papillary thyroid carcinoma, cribriform-morular variant |
| 68 | thyroid | cancer | Medullary thyroid carcinoma, lymph node metastasis |
| 69 | pleura | cancer | Malignant pleural mesothelioma |
| 70 | musculosceletal | cancer | Chordoma, sacral |

### TABLE S6. KEY RESOURCES

| REAGENT or RESOURCE | SOURCE | IDENTIFIER |
|---|---|---|
| Antibodies and proteins | | |
| Purified antibodies, see **Table S2** | various | various |
| Mouse IgG | Sigma | I5381 |
| Rat IgG | Sigma | I4131 |
| Biotinylated VG1 hyaluronan-detection reagent | Bollyky Lab, Stanford U | (Clark et al., 2011) |
| Oligonucleotides | | |
| CODEX oligonucleotides, see **Table S3** | TriLink Biotechnologies and Integrated DNA Technologies | N/A |
| Biological Samples | | |
| FFPE tissue blocks | Institute of Pathology, University of Bern | N/A |
| Chemicals and Reagents | | |
| Streptavidin-PE | Biolegend | 405203 |
| PBS | Thermo Fisher Scientific | 14190-250 |
| NaCl | Thermo Fisher Scientific | S271-10 |
| $Na_2HPO_4$ | Sigma | S7907 |
| $NaH_2PO_4 \cdot 7\ H_2O$ | Sigma | S9390 |
| $MgCl_2 \cdot 6\ H_2O$ | Sigma | M2670 |
| $NaN_3$ | Sigma | S8032 |
| EDTA | Sigma | 93302 |
| TCEP | Sigma | C4706 |
| NaOH | Sigma | S8263 |
| BS3 | Thermo Fisher Scientific | 21580 |
| DMSO | Thermo Fisher Scientific | D128-4 |
| DMSO | Sigma | 472301 |
| DMSO ampoules | Sigma | D2650 |
| Paraformaldehyde ampoules, 16% | Thermo Fisher Scientific | 50-980-487 |
| BSA | Sigma | A3059 |
| Tris 1 M, pH 8.0 | Teknova | T1080 |
| Candor PBS antibody stabilizer solution | Thermo Fisher Scientific | NC0436689 |
| Salmon sperm DNA, sheared | Thermo Fisher Scientific | AM9680 |
| Triton™ X-100 | Sigma | T8787 |
| Ethanol, 100% | Sigma | E7023 |
| Acetone, 100% | Thermo Fisher Scientific | A929-4 |
| Methanol, 100% | Thermo Fisher Scientific | A412-4 |
| Trizma® HCl | Sigma | T3253 |
| Trizma® Base | Sigma | T1503 |
| Drierite indicating desiccant | Thermo Fisher Scientific | 07-578-3A |
| Bondic polyacrylamide gel | Amazon | B018IBEHQU |
| Dako target retrieval solution, pH 9.0 | Agilent | S236784-2 |
| TBS IHC wash buffer with Tween® 20 | Cell Marque | 935B-09 |
| Antibody diluent | Agilent | S080981-2 |
| Protein block, serum-free | Agilent | X090930-2 |
| Dual endogenous enzyme-blocking reagent | Agilent | S200380-2 |
| Liquid DAB+ substrate chromogen system | Agilent | K346711-2 |
| EnVision+ HRP Mouse | Agilent | K400011-2 |
| EnVision+ HRP Rabbit | Agilent | K400211-2 |
| Hematoxylin, ready-to-use | Agilent | S330930-2 |
| Eosin Y solution | Sigma | HT110116 |
| Cytoseal XYL | Thermo Fisher Scientific | 8312-4 |
| Sally Hansen Nail Polish, clear | Amazon | B00CMFMYEG |
| Critical Commercial Instruments, Consumables, Kits and Assays | | |
| LTS filter tips, 10 µl | Rainin | 30389225 |
| LTS filter tips, 200 µl | Rainin | 30389239 |
| LTS filter tips, 1000 µl | Rainin | 30389212 |
| Amicon™ Ultra Centrifugal Filters, 50kDa | Thermo Fisher Scientific | UFC505096 |
| Nalgene™ Rapid Flow 500 ml filter, 0.2 µm | Thermo Fisher Scientific | 09-740-28C |
| Glass coverslips, 22x22 mm, # 1 1/2 | Electron Microscopy Sciences | 72204-01 |
| Frosted microscope slides | Thermo Fisher Scientific | 12-550-343 |

| | | |
|---|---|---|
| Glass coverslip storage box | Qintay | CS-22 |
| 22x22 mm coverslip mounting gaskets | Qintay | TMG-22 |
| Wheaton™ Coverslip glass jars | Thermo Fisher Scientific | 02-912-637 |
| Dumont #5/45 cover slip forceps | Fine Science Tools | 11251-33 |
| ST4020 small linear stainer | Leica | 14050946425 |
| Labconco™ Fast-Freeze™ Flasks, Complete Assembly, 900ml | Thermo Fisher Scientific | 10-269-63 |
| FreeZone® 4.5 Plus Cascade Benchtop Freeze Dry System | Labconco | 7386030 |
| 8-strip tubes, 0.2 ml | E&K Scientific | 280008 |
| 8-strip caps, flat top | E&K Scientific | 491008 |
| 8-strip caps, dome top | E&K Scientific | 491018 |
| CODEX acrylic plates | Bayview Plastic Solutions | custom made |
| BZ-X710 fluorescence microscope | Keyence | N/A |
| Hoechst 33342 | Thermo Fisher Scientific | 62249 |
| DRAQ5 | Cell Signaling Technology | 4084L |
| Vectabond™ | Vector Labs | SP-1800 |
| Vacuum Desiccators, 23L | Thermo Fisher Scientific | 08-648-112 |
| Corning™ black 96-well plates | Thermo Fisher Scientific | 07-200-762 |
| Axygen aluminum sealing film | VWR Scientific | 47734-817 |
| TMA Grand Master | 3DHistech | N/A |
| Pannoramic P250 digital slide scanner | 3DHistech | N/A |
| CODEX System | Akoya Biosciences | N/A |
| Deposited Data | | |
| Software and Algorithms | | |
| BZ-X viewer | Keyence | N/A |
| CODEX driver | Akoya Biosciences | N/A |
| CODEX Toolkit, version 1.3.5 | https://github.com /nolanlab/CODEX | (Goltsev et al., 2018) |
| Microvolution software for deconvolution | www.microvolution.com | N/A |
| ImageJ (Fiji version 2.0.0) | http://imagej.net | N/A |
| VorteX (X-shift clustering algorithm) | https://github.com /nolanlab/VORTEX | (Samusik et al., 2016) |
| CellEngine | www.cellengine.com | (Bjornson-Hooper et al., 2019) |
| R, version 3.4.3 | www.r-project.org | N/A |
| R studio desktop, version 1.1.423 | www.rstudio.com | N/A |
| Neighborhood analysis notebooks | This paper | N/A |
| Tensorly Python package | http://tensorly.org/ | (Kossaifi et al., 2019) |
| Statsmodel Python package | https://www.statsmodels.org/ | (Seabold and Perktold, 2010) |
| Scikit learn Python package | https://scikit-learn.org/ | (Pedregosa et al., 2011) |
| Survival R package | https://cran.r-project.org/web/packages/survival/index.html | (Therneau, 2015) |
| Glmnet R package | https://cran.r-project.org/web/packages/glmnet/index.html | (Friedman et al., 2010) |
| Visreg R package | https://cran.r-project.org/web/packages/visreg/index.html | (Breheny and Burchett, 2013) |
| Deldir R package | https://cran.r-project.org/web/packages/deldir/index.html | N/A |
| ComplexHeatmap R package | https://bioconductor.org/packages/release/bioc/html/ComplexHeatmap.html | (Gu et al., 2016) |
| The Human Protein Atlas | www.proteinatlas.org | N/A |
| Pathology Outlines | www.pathologyoutlines.com | N/A |
